# SPOROS: A pipeline to analyze DISE/6mer seed toxicity

**DOI:** 10.1101/2021.07.01.450720

**Authors:** Elizabeth T. Bartom, Masha Kocherginsky, Bidur Paudel, Aparajitha Vaidyanathan, Ashley Haluck-Kangas, Monal Patel, Kaitlyn L. O’Shea, Andrea E. Murmann, Marcus E. Peter

**Author notes:** Corresponding authors: Marcus Peter, and Elizabeth Bartom.

## Abstract

micro(mi)RNAs are (18-22nt long) noncoding short (s)RNAs that suppress gene expression by targeting the 3’ untranslated region of target mRNAs. This occurs through the seed sequence located in position 2-7/8 of the miRNA guide strand, once it is loaded into the RNA induced silencing complex (RISC). G-rich 6mer seed sequences can kill cells by targeting C-rich 6mer seed matches located in genes that are critical for cell survival. This results in induction of Death Induced by Survival gene Elimination (DISE), also referred to as 6mer seed toxicity. miRNAs are often quantified in cells by aligning the reads from small (sm)RNA sequencing to the genome. However, the analysis of any smRNA Seq data set for 6mer seed toxicity requires an advanced workflow, solely based on the exact position 2-7 of any sRNA that can enter the RISC. Therefore, we developed SPOROS, an automated pipeline that produces multiple useful outputs to compare 6mer seed toxicity of all cellular sRNAs, regardless of their nature, between different samples. We provide two examples to illustrate the capabilities of SPOROS: Example one involves the analysis of RISC-bound sRNAs in a cancer cell line (either wild-type or two mutant lines unable to produce most miRNAs). Example two is based on a publicly available smRNA Seq data set from postmortem brains (either from normal or Alzheimer’s patients). Our methods are designed to be used to analyze a variety of smRNA Seq data in various normal and disease settings.

## 1. Introduction

micro(mi)RNAs are short (18-22nt long) noncoding RNAs that negatively regulate gene expression [1]. They are generated as double stranded (ds)RNA duplexes. Their activity involves only a very short region of complete complementarity between the ‘seed’, at position 2-7/8 of the guide strand of the miRNA [2, 3] and ‘seed matches’ predominantly located in the 3’ untranslated region (3’ UTR) of targeted mRNAs [4, 5]. This targeting results in gene silencing [6]. miRNA biogenesis begins in the nucleus with the transcription of a primary miRNA precursor [7]. The Drosha/DGCR8 microprocessor complex first processes them into pre-miRNAs [8], which are then exported by Exportin-5 from the nucleus to the cytoplasm [9]. Once in the cytoplasm, Dicer/TRBP processes the pre-miRNAs further [10, 11], and these mature dsRNA duplexes are then loaded onto argonaute (Ago) proteins forming the RNA-induced silencing complex (RISC) [12]. The active miRNA guide strand incorporates into the RISC [12], while the inactive passenger strand is degraded [13].

We previously discovered a powerful new cell death mechanism (6mer seed toxicity) that is based on a 6mer seed embedded in miRNAs. Any si-, sh, or miRNA that carries a 6mer seed of a certain nucleotide composition, kills all cancer cells by targeting the mRNAs of hundreds of genes that are critical for cell survival [14]. An arrayed high throughput screen of all 4096 possible 6mer seeds in a neutral backbone with a chemically inactivated passenger strand revealed that the most toxic seeds were G-rich followed by seeds rich in Cs [15]. A consensus seed among the 100 most toxic seeds for human cells was identified as GGGGGC and we demonstrated that it is toxic by targeting GCCCCC seed matches present in the 3’ UTR of numerous survival genes [16].

The latest number of putative human miRNAs has been estimated to be >8,300 [17]. The most widely established approach to study the role of miRNAs focuses on only the miRNAs that are significantly deregulated when comparing two states (e.g., tumor versus normal tissue, or two developmental stages of an embryo). Hence, most methods to normalize and analyze miRNAs are aimed at allowing investigators to identify individual miRNAs or groups of miRNAs. This makes the depiction of the relevant miRNAs more manageable as there is no need to visually display hundreds of miRNAs at the same time. However, there are two major drawbacks to this approach: First, the detected fold change in relative expression of a deregulated miRNA does not allow one to conclude that a miRNA is significantly expressed, and second, miRNAs that belong to different families but function in similar ways in different tissues or in a disease context in different patients, are hard to identify.

The 6mer Seed Tox concept requires analysis of short (s)RNAs, including miRNAs, in a different way. Rather than aligning all reads to the genome and finding the ones coding for miRNA genes, the only relevant information needed of any sRNA that is bound to the RISC and active in RNA interference (RNAi) function, is the precise knowledge of its 6mer seed (position 2-7 from the 5’ end). The nature of the sRNA is secondary and can be determined later. 6mer Seed Tox can be best determined by analyzing sRNAs that are loaded into the RISC, but total small RNA Seq data can also be useful, as long as the reads are in the range of 18 and 25 nt long. It has become clear that this activity is not only found in miRNAs, but in any abundant sRNA such as tRNA or ribosomal (r)RNA fragments that can be loaded into the RISC and exert RNAi [18, 19]. We now describe SPOROS (Greek for seed), an automated pipeline that allows for the analysis of sRNAs in a new way, focused not on individual miRNAs and their targets, but on the 6mer seed of any sRNA that is loaded into the RISC. SPOROS generates multiple output files that allow one to assess both composition and seed toxicity changes in miRNAs in any small RNA Seq data set. The focus in developing SPOROS was to provide simple and robust analysis tools that can be used without requiring advanced programming knowledge.

## 2. Methods and applications

In the following, we detail different steps to analyze sRNAs based on their 6mer Seed Tox (**Figure 1**). We analyze two examples to illustrate the power and utility of the pipeline. The first example is a smRNA Seq dataset of RISC-bound sRNAs in a wild-type (wt) human cancer cell line and two mutant cell lines lacking expression of either Drosha or Dicer, resulting in a fundamental reduction in miRNA expression (**Figure 2**). The second example is based on a publicly available small RNA Seq data set derived from postmortem normal and Alzheimer’s disease (AD) patient brains (**Figure 3**). We also provide a comparison of the groups of miRNAs that exist in both human and mouse cells with conserved seed toxicities in the two species (**Figure 4** and **Table S1**). **Figure 4A** shows a Venn diagram with the overlap of 406 mature miRNAs that are conserved with the same seed between human and mouse and **Figure 4B** compares the seed toxicities between the two species in a regression analysis with a number of toxic miRNAs, mostly tumor suppressive, and some miRNAs with nontoxic seeds labeled in red and green, respectively.

**Figure 1.**
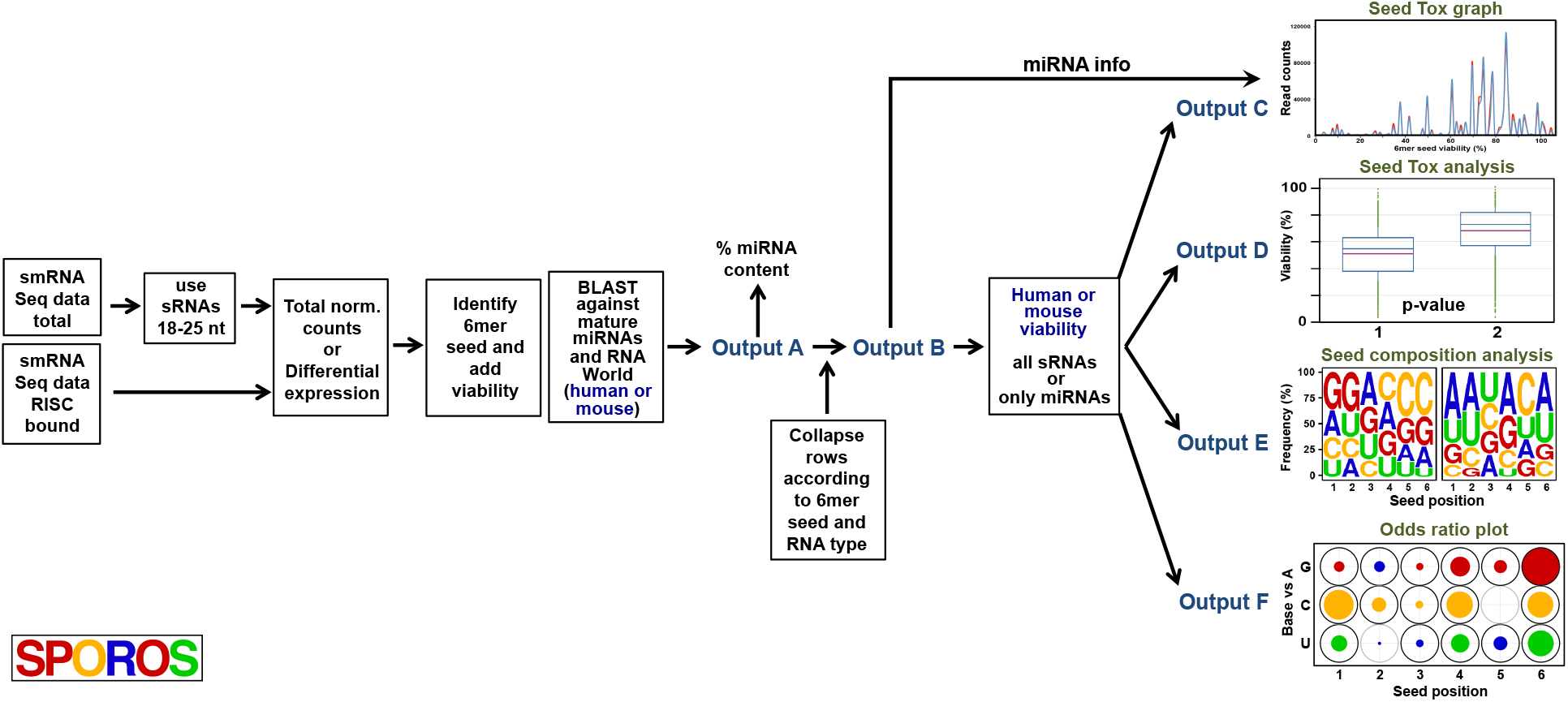
SPOROS workflow developed to analyze seed toxicity of smRNA Seq data. From left to right: smRNA seq data, either total or RISC-bound are normalized to 1 million reads/sample. After elimination of residual adapter sequences 6mer seeds and matching seed viabilities are added for each read and sequences are blasted against all mature miRNAs or RNA World data sets of all small RNAs (either human or mouse). At this point the miRNA content (%) can be determined. Reads of this Output A file are collapsed according to 6mer seed and RNA type resulting in Output B. At this point all short RNAs can be analyzed (sRNA) or just the miRNA fraction. Output B is fed into four scripts generating four output files: C: A Seed Tox graph that depicts all miRNAs as peaks according to their seed viability; D: Average seed toxicity of all reads in a samples depicted as box and whisker plot; E: Weblogo plot showing the average seed composition in positions 1-6 of the 6mer seed in each sample; F: The result of a multinomial mixed model odds ratio analysis allowing to compare both different 6mer seeds as well as differences in each position of different seeds.

**Figure 2.**
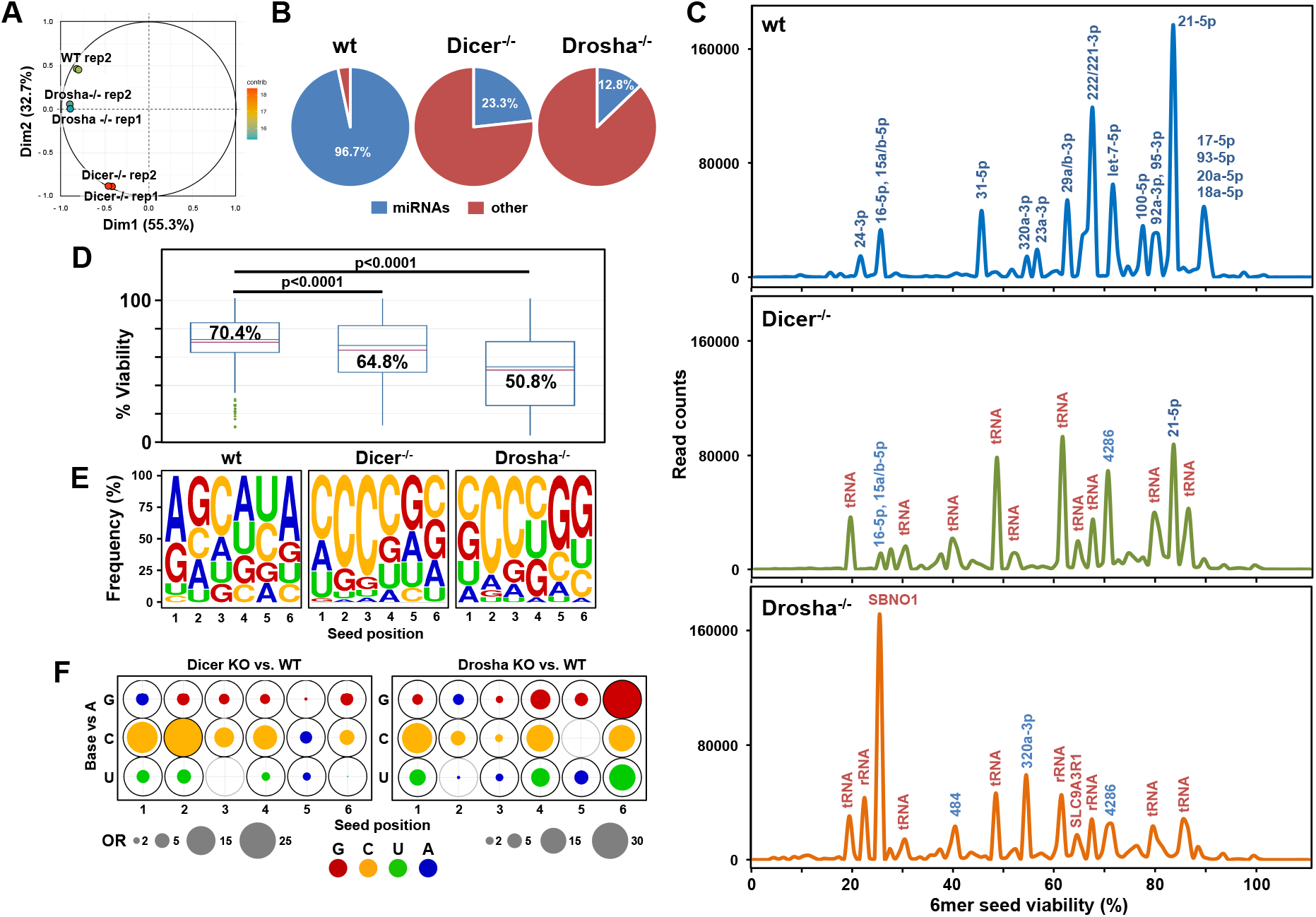
The RISC in HCT116 Dicer and Drosha k.o. cells is enriched in sRNAs with toxic 6mer seeds compared to wt cells. (A) Principal component analysis (PCA) plot illustrating the differences in RISC composition of Dicer and Drosha k.o. cells compared to HCT116 wt cells, and the reproducibility of technical and biological replicates. The x-axis represents dimension 1 (dim1) and explains 55.3% of the variance, while the y-axis represents dimension 2 (dim2) and explains 32.7% of the variance. Each cell type was analysed as two biological replicates. Each spot represents a single replicate sample from each cell type. Green-HCT116 wt, blue-Drosha k.o. and red-Dicer k.o. (B) Pie charts showing RISC composition of HCT116 wt, Dicer and Drosha k.o. cells. Abundance of miRNAs is shown in blue and all other sRNAs in red. (C) RISC-bound sRNA Seed Tox graphs of HCT116 wt, Dicer k.o., and Drosha k.o. cells. When a peak is labeled with multiple miRNAs (blue), the most abundant one is listed first. RNAs are only labeled if they account for 1000 reads or more. miRNAs are labeled in blue, other sRNAs in red. (D) Average Seed Tox of all RISC-bound sRNAs enriched in cells in C. p values were calculated using a Wilcoxon rank test. (E) Seed composition of all RISC-bound sRNAs enriched in cells in C. (F) Filled circles at each position represent the odds ratio (OR) estimates comparing the odds of observing G, C and U vs A between genotypes, based on the multinomial mixed effects model. The outer circle corresponds to the largest observed OR for each pairwise genotype comparison (e.g., Dicer k.o. vs. wt), with bold black circles denoting statistical significance based on p-values adjusted using Tukey’s method. OR<1 estimates are represented with blue circles with area OR*=1/OR, indicating that A is more likely in this position. Circle area is scaled to be proportional to the OR.

**Figure 3.**
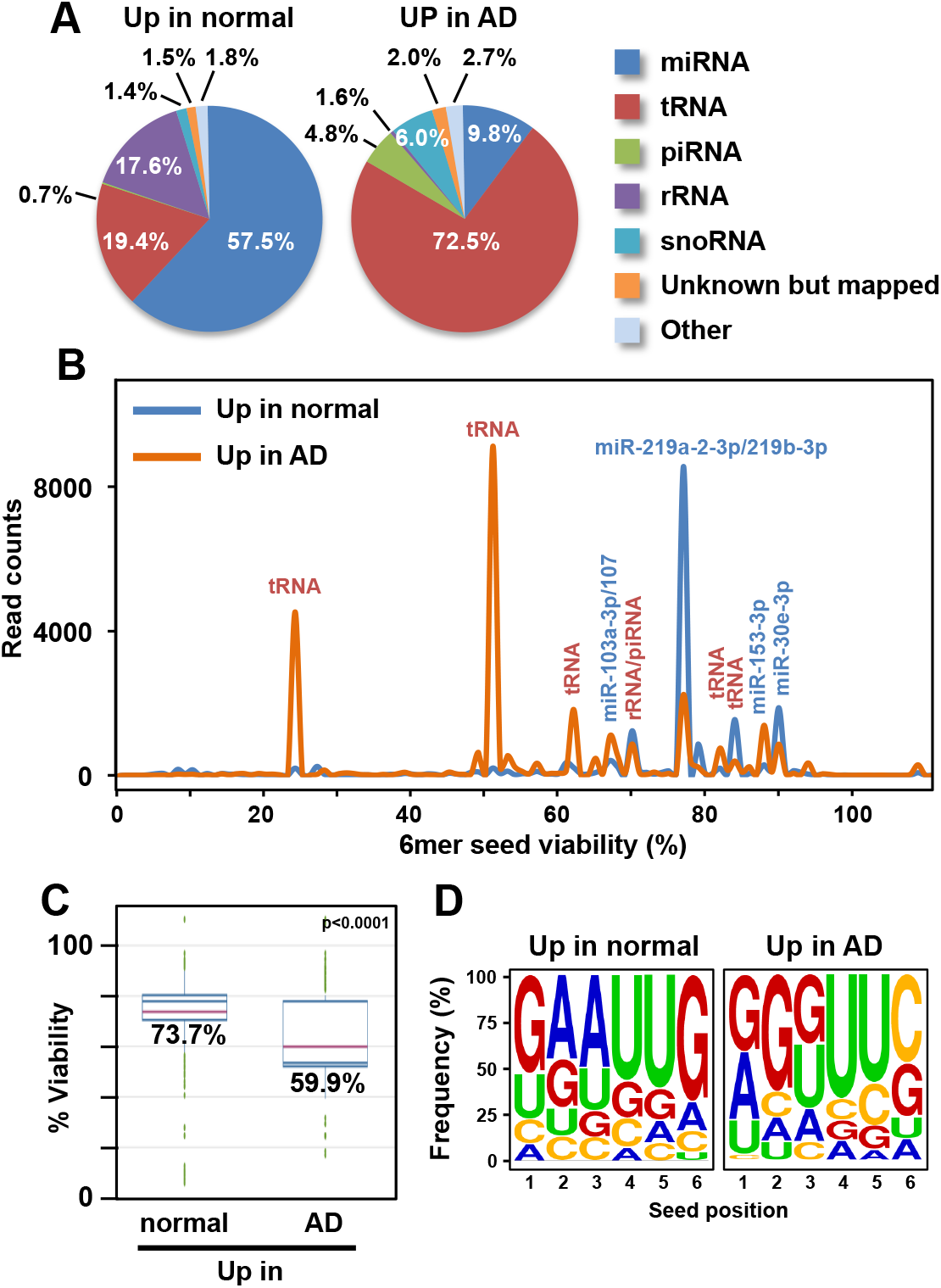
sRNAs enriched in AD brains contain more toxic seeds than in control brains. (A) Pie charts showing composition of sRNAs expressed in normal and AD patient brains. (B) Seed Tox plots of sRNAs differentially expressed in brain samples in A. When a peak is labeled with multiple miRNAs (blue), the most abundant one is listed first. RNAs are only labeled if they account for 1000 reads or more. miRNAs are labeled in blue, other sRNAs in red. (C) Average Seed Tox of all differentially regulated sRNAs in brain samples in A. p-value was calculated using a Wilcoxon rank test. (D) Seed composition of all RISC-bound differentially regulated sRNAs in brain samples in A

**Figure 4.**
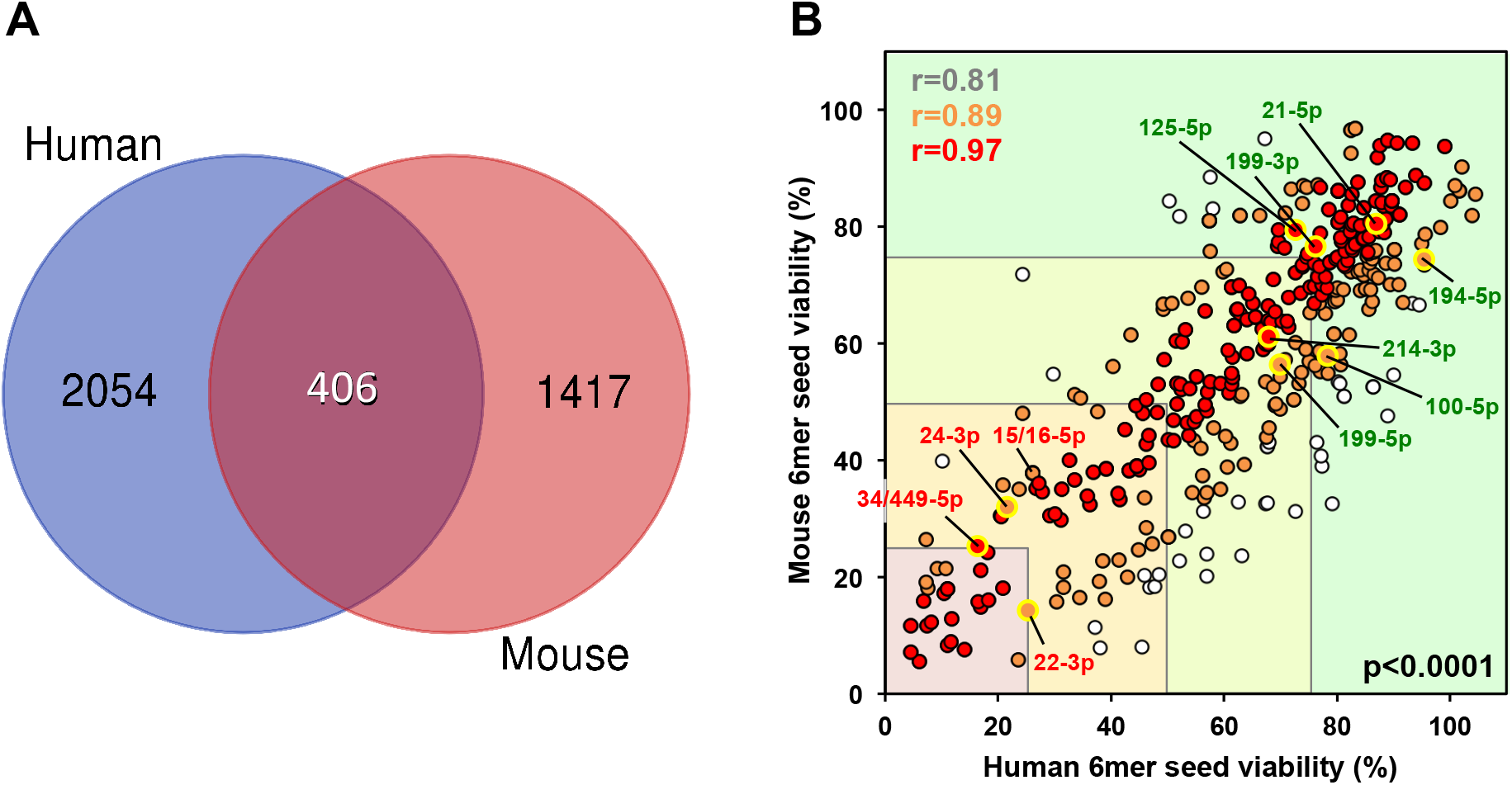
The miRNAs conserved between human and mouse and their seed viabilities. (A) Venn diagram of the overlap of mature miRNAs present in both human and mouse genomes (see **Tables S2** and **S3**). (B) The 406 miRNAs overlapping between human or mouse were grouped into brackets based on the average seed viability between human mouse (0-25%, pink; >25-50%, orange; >50% - 75%, light green; >75% green). Three Pearson regression analyses were performed: 1-All miRNAs (grey dots), 2-miRNAs with a seed viability difference of less than 25% (orange dots), and 3-miRNAs with a seed viability difference of less than 10% (red dots). r-values are given for each of the three analyses in matching colors. p-values of all three analyses were <0.0001. Selected miRNAs with either toxic (red) or nontoxic (green) seed we previously identified [15, 16, 22, 34] and/or validated are labeled.

## 3. Description of the example data sets

The first smRNA Seq data set of RISC-bound sRNAs was generated by performing an Ago pull down experiment followed by smRNA Seq as described before [15]. In brief, 10^6^ HCT116 wt, Drosha knock-out (k.o.) or Dicer k.o. cells [20] were subjected to an AGO pull down (in duplicate) using a bead bound GW182 protein [21]. After smRNA library preparation, the samples were subjected to 50 nt single end smRNA Seq on an Illumina HiSEQ4000. Raw read sequences were trimmed with trim_galore to remove all standard Illumina adapters, leaving reads at least 6 bp in length, as described previously [22]. The second smRNA Seq data set was obtained at GEO (accession number GSE63501) and contains sRNAs (16-25 nt in length) derived from 7 control brains and 6 AD brains [23].

## 4. The SPOROS pipeline

The goal of the SPOROS pipeline is to display and analyze abundance of sRNAs according to their 6mer Seed Tox (**Figure 1**). It can be accessed at https://github.com/ebartom/SPOROS. At heart, SPOROS is a Perl-based decision tree. Given a few essential arguments (location of the input data, organism, experimental parameters if relevant), SPOROS will generate a commented shell script listing each step in the pipeline. This script can be run on the command line to carry out the pipeline in its entirety. Any small (sm)RNA Seq data set (either total smRNA or smRNAs bound to the RISC) can be used, either starting from raw fastq files, or from a table of read counts. While the 6mer Seed Tox concept is based mostly on the activity of miRNAs, it can be applied to any sRNA that enters the RISC as a guide, and hence the relevant activity of an sRNA is determined by its position 2-7 (the 6mer seed). This allows one to display all sRNAs in a graph as a function of only the predicted toxicity of its 6mer seed. While the 3’ end of the RNAs is not that relevant for their activity, for this analysis to succeed the knowledge of the exact 5’ start is critical. Consequently, the first step of the analysis is to de-multiplex the samples and remove any Unique Molecular Identifier (UMI) and adapter sequences. Trim_galore is used to identify and remove standard Illumina adapters. 5’ adapter sequences are removed first, followed by the removal of the 3’ adapters which in the case of our libraries is the substring GTCCGACGATC. When analyzing total smRNA Seq data, all reads that are longer than 25 or shorter than 18 nt are removed, as shorter and longer reads have a reduced chance of entering the RISC. Once any extraneous sequences are removed, a table of all unique reads observed in all samples, and their counts in each sample is generated. If desired, such a table can be the start of the pipeline. SPOROS then extracts the 6mer seed sequence for each read and adds its predicted 6mer Seed Tox (which is the % viability determined by transfecting three human and three mouse cell lines with siRNAs carrying all of the 4096 possible 6mer seeds ([15], 6merdb.org). By default, the average seed viabilities of the three human and three mouse cell lines are added. SPOROS then automatically creates the following output files that can be used to create display figures (**Figure 1**):

### Output A

(file name starts with “A_normCounts ”) contains all normalized data (each sample normalized to 1 million reads) with seeds, seed toxicities, miRNA, and RNA World information added. The information on miRNAs is added by blasting each read against a curated list of either human or mouse mature miRNAs (human and mouse list are in **Table S2** and **S3**). For this analysis, we set the stringency so that only hits with at least 18 nt of complete identity between the queried read and the mature miRNA, were counted. These BLAST results can be used to determine % miRNA content. In RISC-associated sRNAs, the percent miRNA of a good pull-down is expected to be >95%. The second BLAST search is performed against a list of small RNAs (data sets for human and mouse available at https://rnaworld.rockefeller.edu, can be obtained from Dr. Thomas Tuschl and are also available upon request). In this case, we allowed a 95% identity for the search.

### Output B

(file name starts with “B_collapsed ”) is generated by adding up and collapsing all rows that contain the same 6mer seed and either the same name in the miRNA or the RNA World blast columns. Counts are added for each seed and miRNA combination. An intermediary file (file name starts with “Int_seedKeyed”) is also created, with each seed listed only once, even if it originates from multiple sRNAs. Depending on the species, subsequent steps will be done either with human or with the mouse seed viability data. Of note, in the analysis we use viability as a measure of Seed Tox. Thus seeds with a low viability percentage are considered more toxic and the ones with a high viability percentage are considered less toxic. In addition, as desired, the data can be split into only miRNAs or all sRNAs.

### Output C

(file name starts with “C_binned ”) is generated by aggregating all rows according to the seed toxicity in 1% rounded steps (creating 1% sized bins). Counts are added for all seeds that have toxicity within each 1% bin. These files contain a row for each bin even if the count is 0, allowing for easy plotting of the data. We use Excel to generate the graphs of read counts vs. viability bin, with line smoothing turned on. The resulting plot, the Seed Tox graph, allows one to display abundance of all miRNAs solely based on their seed toxicity. An option at this stage is to plot all sRNAs/miRNAs in the analysis or only the ones significantly deregulated (<0.05 adjusted p-value) between conditions. The Seed Tox graph can then be generated with either all or the differentially expressed sRNAs. Here we show an example for each case (**Figures 2** and **3**). The peaks in the Seed Tox graph can then be manually labeled with the most prominent miRNAs that fall with in that particular viability range. This information can be obtained from Output file B. Most peaks will contain multiple miRNAs and we usually label the ones whose abundance is above a certain threshold (e.g., 1000 reads). We chose to generate this Seed Tox graph over a density plot (**Figure S1**) because it allows to compare the actual read numbers in each peak between samples and different analyses.

### Output D

(file names start with “D_toxAnalysis”). The SPOROS pipeline creates these files from Output B to display the average seed viability in each analyzed group or in individual samples. The file expands each row according to the number of reads in that row after a normalization to 1000 rows (this normalization, i.e., dividing by 1000, prevents files from getting too large and thus increasing computational complexity of the statistical analysis). The data column can be used in any standard statistics program to create a plot. We usually summarize the distributions using a boxplot. A nonparametric rank test can be performed to compare viability between groups. We currently use StatPlus (v. 7.5) and the Wilcoxon ranksum test of the Kruskal-Wallis test if there are more than 2 groups.

### Output E

(file names start with “E_seedAnalysis”). These files are also created from Output B in a similar way as Output D, except instead of expanding each row according to toxicity, the row is expanded according to the 6mer seed. SPOROS uses this output file to generate a custom Weblogo (http://weblogo.threeplusone.com/) to display nucleotide frequencies in each of the 6 seed positions. The Weblogo is generated as pdf and high resolution png file that easily allows the user to extract the image and convert it into a desired format.

### Output F

(file name starts with “F_seedExpand”). Output F is used in the statistical analysis comparing 6mer seed composition between different samples. It is generated from output E by further expanding the rows with average seed counts so that each nucleotide is in a different row for each seed, and rows are indexed by position. As in Output D, seed counts are divided by 1000, and seeds with counts <1000 are omitted from this analysis. In total, Output F contains 6N rows, where N is the total scaled seed count per sample.

To analyze seed composition data, we developed a novel framework using multinomial mixed effects regression models. This approach allows us to compare differences in sequence patterns between groups and positions and provides a statistical framework for both testing and estimation of such differences. Unlike analyses which compare counts between groups, here each seed represents the unit of analysis, and nucleotides are compared between positions and samples. Using terminology from the generalized linear mixed effects models literature [24] each seed can be thought of as a “subject”, and the nucleotides in each position can be thought of as 6 potentially correlated measurements within a “subject”. Correlation between positions could occur, for example, if certain patterns are likely. An example would be the enrichment of Gs versus other nucleotides towards the 5’end of the 6mer seed as reported [16]. The outcome variable in the model is the nucleotide in each position of each seed. We assume that at each position nucleotides follow a multinomial distribution with 4 possible outcomes (A, C, G, U), and, as in logistic regression, we must set one of the levels as the reference (we set A as the reference). Group (e.g., genotype or experimental condition), position and their interaction are included as the “fixed effect” predictors, and the seed itself (i.e., the unique seed id) is included as the “random effect”. Including the seed random effect allows us to account for the potential within-seed correlation between positions. The estimated model provides tests of whether there are group or position differences, and the odds ratios comparing the odds of observing G, C or U vs. A between groups at each position. These models can be fitted using PROC GLIMMIX in SAS statistical software [25].

Outputs A-E can be generated for either total reads (as in Figure 2) or only for those reads that are enriched in one set of samples relative to another (as in Figure 3). In the case of a differential analysis, SPOROS users create a comparisons table setting up one set of samples (denoted with 1) as Group1 and another set of samples (denoted with −1) as Group2. Samples irrelevant to a particular pairwise comparison are denoted in the comma separated table with a 0 (more detail is included with the SPOROS documentation on Github). The R package EdgeR [26] is used to identify reads differentially expressed within each pairwise comparison. Output A file names contain “diff” (adjusted p-value < 0.05 and logFC > 0.585 or < −0.585). These differentially regulated reads are used for Output C-E, see **Figure 3B-D**.

## 5. An example analysis of RISC-bound sRNAs (all sRNAs)

For the first example we chose to analyze RISC-bound sRNAs isolated from wt HCT116 cells and cells deficient for either Drosha or Dicer (**Figure 2**). Neither of these two k.o. cells can produce canonical miRNAs [20]. As previously described [15, 21, 22, 27], we used a GW182 peptide coupled to GST to pull down all four Ago proteins which are critical for RISC formation and function [28]. A principal component analysis (PCA) shows all three genotypes cluster independently with biological replicates for each genotype tightly grouped together (**Figure 2A**). (PCA script in **Appendix A)**. Wild-type cells contained >96% miRNAs and this amount was reduced to ~23% in the Dicer k.o. cells and further reduced to ~13% in Drosha k.o. cells (**Figure 2B**).

When comparing the three genotypes using the Seed Tox graph, it became apparent that in both Dicer and Drosha k.o. cells, most miRNAs in the RISC were replaced by other sRNAs, most notably tRNA fragments (**Figure 2C**). This likely occurred because in contrast to normal cells [29, 30], tumor cells maintain expression of argonaute proteins in the absence of miRNAs [14].

The change in RISC composition in the two mutant cells resulted in a reduction in average seed viability of all reads (**Figure 2D**). This was most prominent for the Drosha k.o. cells which have the lowest amount of miRNAs, suggesting that endogenous miRNAs which carry mostly nontoxic seeds [22] likely protect cells from potentially toxic endogenous sRNAs entering the RISC. These sRNAs (e.g., tRNA or rRNA fragments) often contain C-rich sequences, which is likely the reason why the average 6mer seed composition in the RISC shifted towards C-richness in the mutant cells with sequences being somewhat more G-rich in the Drosha k.o. cell lines (**Figure 2E**). Together, these trends likely account for the more strongly reduced seed viability (**Figure 2D**) in these cells.

Multinomial mixed effects models revealed that differences between the three genotypes differ by position (p<0.0001, interaction term). Model-based odds ratio (OR) estimates comparing G, C and U vs. A between genotypes are graphically summarized in **Figure 2F**, and the majority are statistically significant (dark black outer circles). For example, relative to A, C is more likely to occur in almost all positions of the seed in Dicer k.o. cells than in wt (OR=5.5 to 27.7 in all positions except position 5; p<0.0001, large orange circles). Similarly, relative to A, G is significantly more likely to occur in Dicer k.o. than in wt in positions 2-6, but the differences between genotypes are smaller (OR=1.8 to 4.5; p<0.0001). Intuitively, the odds ratios correspond to the relative frequencies in **Figure 2E**. For example, in the Drosha k.o. weblogo, in position 6 G occurs much more frequently than A, whereas in wt G occurs less frequently than A, resulting in a particularly large odds ratio OR=35.9 (**Figure 2F**).

We previously showed that these Drosha k.o. cells grow slower than their wt counterparts and knocking down Ago2 corrected the growth rate [27], suggesting that endogenous sRNAs with toxic seeds that entered the RISC were causing a growth reduction. Importantly it demonstrated the relevance of including non-canonical sRNAs in the Seed Tox analysis. Non-canonical sRNAs in the RISC also exert RNAi activity.

## 6. An example analysis of total cellular smRNAs (differentially expressed sRNAs)

The data set of the second example on sRNAs from AD brains contains RNAs 16-25 nt in length and had already been assigned RNA species using RNA World [23]. Our analysis now shows that sRNAs significantly enriched in either control or AD brains have a profound shift from mostly miRNAs to mainly tRNA fragments (**Figure 3A**), a phenomenon quite similar to that observed in the HCT116 Drosha k.o. cells in which miRNA biogenesis is impaired. To determine how this shift influences the Seed Tox, we used SPOROS to do a differential analysis of the read counts made available in GEO (GSE63501), starting from a counts table, and running a differential analysis. The Seed Tox graph based on Output B revealed a shift from mostly nontoxic miRNAs in control brains to sRNAs with more toxic seeds in AD patients. Most of the toxic seeds are contributed by tRNAs (**Figure 3**). This shift to seeds with lower viability also became apparent in the average Seed Tox analysis of all reads (**Figure 3C**) and this was mostly due to a significant increase in Gs towards the 5’ end of the 6mer seed of the sRNA in the AD brains (**Figure 3D**). This analysis suggests that in AD brains the repertoire of sRNAs available to function through RNAi are predicted to be more toxic.

## 7. Conclusions

The human genome contains a large number of predicted miRNAs [17]. They have been shown to regulate almost all biological processes and to be deregulated in countless disease states [31]. The state-of-the-art method to analyze miRNAs is by RNA Seq. Almost all RNA Seq data are analyzed with the goal of identifying differentially expressed genes after aligning reads to a genome and employing appropriate normalization to account for transcript length (reads per kilobase gene model) or variation between data sets. These types of analyses, while standard, have major shortcomings and a comprehensive method to normalize these large data sets to identify and study individual miRNAs has not been developed [32]. The analysis of 6mer Seed Tox does not focus on individual miRNAs or canonical miRNA families but treats miRNAs solely based on their position 2-7. This allows for the ranking of all miRNAs into blocks from highly toxic to nontoxic miRNAs and often it is the balance between the sum of toxic versus nontoxic sRNAs bound to the RISC that determines the responses of cells [22]. The analyses in the two examples presented (cancer and AD), were chosen to demonstrate the power and the potential of the SPOROS pipeline to analyze 6mer Seed Tox. In example #1, the data support the view that most miRNAs carry nontoxic seeds and are in part protecting cells from loading of endogenous sRNAs, that by nature are more G/C rich and hence when entering the RISC, exert toxicity. In the AD example #2, the data suggest that in AD brains, equilibrium shifts away from nontoxic miRNAs to more toxic sRNAs, such as tRNA fragments. While this result needs to be validated by performing Ago pull-down experiments with AD patient brains, it is intriguing that a recent study in another neurodegenerative disease, Huntington’s disease (HD), reported that sRNAs isolated from HD brains were toxic when injected into mouse brains [33]. This was shown in part to be due to an increase in tRNAs in the disease brains when compared to normal control brains. The methods we have developed to analyze 6mer Seed Tox will now allow one to study the above and multiple other disease situations.

## Acknowledgement

We would like to thank Dr. Thomas Tuschl for providing the RNA World data sets. This work was funded by grant R35CA197450 to M.E.P., and P30CA060553 to M.E.P. and M.K., and R50CA221848 to E.T.B.

---

**Figure S1: Displaying Seed toxicity in a density plot.**

RISC-bound sRNA Seed Tox data of HCT116 wt, Dicer k.o., and Drosha k.o. cells displayed as density plots. The same source data were used to generate **Figure 2C**.

**Table S1. List of mature miRNAs with the same 6mer seed that overlap between human and mouse.** Colors are the same as in **Figure 4B** and miRNAs labeled in that figure are in bold. miRNA data sets were obtained from miRBase (v. 22.1) and are reflected in **Tables S2** and **S3**.

**Table S2: Data on human mature miRNAs used in the BLAST search.** Data from miRBase supplemented with adapter and artificial sequences to ensure that these are appropriately flagged. This file also contains four HIV-1 coded miRNAs.

**Table S3: Data on mouse mature miRNAs used in the BLAST search.** Data from miRBase supplemented with adapter and artificial sequences to ensure that these are appropriately flagged.

## Appendix A

Authors: Alboukadel Kassambara, Fabian Mundt

Link to package: https://CRAN.R-project.org/package=factoextra

description of use: http://www.sthda.com/english/rpkgs/factoextra

R Script used for PCA plot in Figure 2A adapted from Statistical tools for high-throughput data analysis (http://www.sthda.com/english/wiki/fviz-pca-quick-principal-component-analysis-data-visualization-r-software-and-data-mining)

~~~
library(tidyverse)
library(data.table)
library(factoextra)
setwd(“/Users/api/Desktop/DISE42_for methods”)
#read in data
DISE42 <- fread(“DISE-
042.noAdapter.notUMId.normCounts.table.withTox.withMiRNAandRNAworld.blast.trunc.minSu m6.txt”)
# make new dataframe with just the data from uninfected cells
DISE42_Uninfected <- cbind(DISE42[,1:2], DISE42[,4])
DISE42_Uninfected <- cbind(DISE42_Uninfected, DISE42[,10:11])
DISE42_Uninfected <- cbind(DISE42_Uninfected, DISE42[,16:17])
DISE42_Uninfected <- cbind(DISE42_Uninfected, DISE42[,22:23])
DISE42_Uninfected <- cbind(DISE42_Uninfected, DISE42[,32])
# abbreviate the column names
colnames(DISE42_Uninfected) <- c(“Read”, “Seed”, “Avg_viability”, “DicerKO.rep1”,
“DicerKO.rep2”, “DroshaKO.rep1”, “DroshaKO.rep2”, “WT.rep1”, “WT.rep2”, “RNAworld”)
# This code splits the RNAworld column by the symbols | or _ and takes the first element to extract
# just the RNA name from the alignment result
DISE42_RNAworld <- lapply(strsplit(DISE42_Uninfected$’RNAworld’, “[|_]”), “[“, 1)
DISE42_RNAworld <- as.data.frame(unlist(DISE42_RNAworld))
DISE42_Uninfected <- cbind(DISE42_Uninfected, DISE42_RNAworld)
# Add the read counts for reads mapping to the same RNA
DISE42_Uninfected_agg <- aggregate(DISE42_Uninfected[,4:9], by = list(DISE42_Uninfected$’unlist(DISE42_RNAworld)’),
             FUN = sum, na.rm = TRUE)
# changes all NAs to 0
DISE42_Uninfected_agg[is.na(DISE42_Uninfected_agg)] <- 0
# makes a new dataframe for the PCA analysis
DISE42_Uninfected_agg.names <- DISE42_Uninfected_agg$Group.1
rownames(DISE42_Uninfected_agg) <- DISE42_Uninfected_agg.names
DISE42_Uninfected_agg2 <- DISE42_Uninfected_agg[,2:7]
# compute PCA and save the results in DISE42_Uninfected_agg.pca
DISE42_Uninfected_agg.pca <- prcomp(DISE42_Uninfected_agg2, scale = TRUE)
# visualize the PCA
# how many dimensions explain the most variation
fviz_eig(DISE42_Uninfected_agg.pca)
# which RNAs contribute most to the variation
fviz_pca_ind(DISE42_Uninfected_agg.pca,
        col.ind = “cos2”, # Color by the quality of representation
        gradient.cols = c(“#00AFBB”, “#E7B800”, “#FC4E07”),
        repel = FALSE # Avoid text overlapping
)
# how do the samples group
fviz_pca_var(DISE42_Uninfected_agg.pca, col.var=“steelblue”)+
   theme_minimal()
fviz_pca_var(DISE42_Uninfected_agg.pca, geom = c(“point”, “text”), col.var=“black”, repel = TRUE)+
  theme_minimal()
fviz_pca_var(DISE42_Uninfected_agg.pca, geom = c(“point”, “text”), col.var = “contrib”,
gradient.cols = c(“#00AFBB”, “#E7B800”, “#FC4E07”), repel = TRUE)
library(“corrplot”)
var <- get_pca_var(DISE42_Uninfected_agg.pca)
corrplot(var$cos2, is.corr = FALSE)
# how do the samples group
fviz_pca_var(DISE42_Uninfected_agg.pca,
        col.var = “contrib”, # Color by contributions to the PC
        gradient.cols = c(“#00AFBB”, “#E7B800”, “#FC4E07”),
        repel = TRUE # Avoid text overlapping
)
# overlay the RNA and sample grouping results
fviz_pca_biplot(DISE42_Uninfected_agg.pca, repel = FALSE,
        col.var = “#2E9FDF”, # Variables color
        col.ind = “#696969” # Individuals color
)
~~~

