## Supplementary figures and images for "SPOROS: A pipeline to analyze DISE/6mer seed toxicity"

### Figure S1

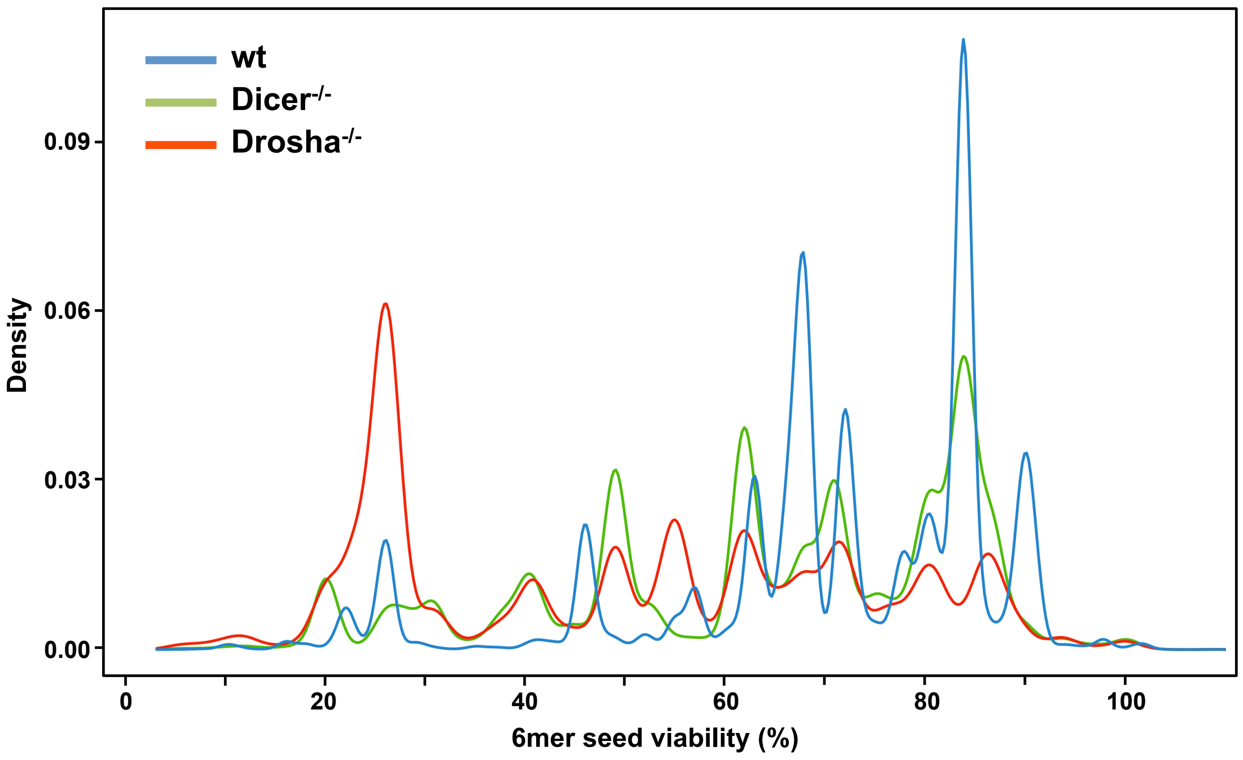
