## Supplementary material for "SPOROS: A pipeline to analyze DISE/6mer seed toxicity": Table S2

**Table S2: Human mature miRNAs (including HIV-1 miRNAs)**

>hsa-miR-519d-3p
CAAAGTGCCTCCCTTTAGAGTG
>hsa-miR-6780b-3p
TCCCTTGTCTCCTTTCCCTAG
>hsa-miR-6884-3p
CCCATCACCTTTCCGTCTCCCCT
>hsa-miR-4296
ATGTGGGCTCAGGCTCA
>hsa-miR-4764-3p
TTAACTCCTTTCACACCCATGG
>hsa-miR-550a-5p
AGTGCCTGAGGGAGTAAGAGCCC
>hsa-miR-514a-5p
TACTCTGGAGAGTGACAATCATG
>hsa-miR-342-3p
TCTCACACAGAAATCGCACCCGT
>hsa-miR-200a-3p
TAACACTGTCTGGTAACGATGT
>hsa-miR-5684
AACTCTAGCCTGAGCAACAG
>hsa-miR-3180
TGGGGCGGAGCTTCCGGAG
>hsa-miR-3142
AAGGCCTTTCTGAACCTTCAGA
>hsa-miR-2116-5p
GGTTCTTAGCATAGGAGGTCT
>hsa-miR-409-3p
GAATGTTGCTCGGTGAACCCCT
>hsa-miR-3661
TGACCTGGGACTCGGACAGCTG
>hsa-miR-4723-5p
TGGGGGAGCCATGAGATAAGAGCA
>hsa-miR-4442
GCCGGACAAGAGGGAGG
>hsa-miR-548g-5p|hsa-miR-548x-5p|hsa-miR-548aj-5p
TGCAAAAGTAATTGCAGTTTTTG
>hsa-miR-7154-3p
AGGAGGACAAGTTGTGGGAT
>hsa-miR-137-3p
TTATTGCTTAAGAATACGCGTAG
>hsa-miR-616-3p
AGTCATTGGAGGGTTTGAGCAG
>hsa-miR-504-3p
GGGAGTGCAGGGCAGGGTTTC
>hsa-miR-1250-3p
ACATTTTCCAGCCCATTCA
>hsa-miR-628-3p
TCTAGTAAGAGTGGCAGTCGA
>hsa-miR-4742-3p
TCTGTATTCTCCTTTGCCTGCAG
>hsa-miR-10400-3p
CTGGGCTCCCGGACGAGGCGGG
>hsa-miR-4259
CAGTTGGGTCTAGGGGTCAGGA
>hsa-miR-10395-3p
ATGTATTCGTACTGTCTGATG
>hsa-miR-1260b
ATCCCACCACTGCCACCAT
>hsa-miR-7161-5p
TAAAGACTGTAGAGGCAACTGGT

>hsa-miR-6737-5p
TTGGGGTGGTCGGCCCTGGAG
>hsa-miR-3193
TCCTGCGTAGGATCTGAGGAGT
>hsa-miR-8072
GGCGGCGGGGAGGTAGGCAG
>hsa-miR-497-5p
CAGCAGCACACTGTGGTTTGT
>hsa-miR-6735-3p
AGGCCTGTGGCTCCTCCCTCAG
>hsa-miR-6516-5p
TTTGCAGTAACAGGTGTGAGCA
>hsa-miR-6509-3p
TTCCACTGCCACTACCTAATTT
>hsa-miR-4791
TGGATATGATGACTGAAA
>hsa-miR-3686
ATCTGTAAGAGAAAGTAAATGA
>hsa-miR-548g-3p
AAAACTGTAATTACTTTTGTAC
>hsa-miR-491-3p
CTTATGCAAGATTCCCTTCTAC
>hsa-miR-4536-3p
TCGTGCATATATCTACCACAT
>hsa-miR-4497
CTCCGGGACGGCTGGGC
>hsa-miR-382-3p
AATCATTCACGGACAACACTT
>hsa-miR-3622b-3p
TCACCTGAGCTCCCGTGCCTG
>hsa-miR-1908-3p
CCGGCCGCCGGCTCCGCCCCG
>hsa-miR-4743-5p
TGGCCGGATGGGACAGGAGGCAT
>hsa-miR-218-1-3p
ATGGTTCCGTCAAGCACCATGG
>hsa-miR-5195-3p
ATCCAGTTCTCTGAGGGGGCT
>hsa-miR-6727-3p
TCCTGCCACCTCCTCCGCAG
>hsa-miR-8074
CTATGGCGAGACTGGCATGTACTC
>hsa-miR-2115-5p
AGCTTCCATGACTCCTGATGGA
>hsa-miR-1-5p
ACATACTTCTTTATATGCCCAT
>hsa-miR-6850-5p
GTGCGGAACGCTGGCCGGGGCG
>hsa-miR-1471
GCCCGCGTGTGGAGCCAGGTGT
>hsa-miR-4750-5p
CTCGGGCGGAGGTGGTTGAGTG
>hsa-miR-6809-5p
TGGCAAGGAAAGAAGAGGATCA
>hsa-miR-30e-3p
CTTTCAGTCGGATGTTTACAGC
>hsa-miR-103a-3p
AGCAGCATTGTACAGGGCTATGA
>hsa-miR-12133
CTTGGCACCATTAAAAAGTACA
>hsa-miR-4267
TCCAGCTCGGTGGCAC

>hsa-miR-4789-3p
CACACATAGCAGGTGTATATA
>hsa-miR-4735-5p
CCTAATTTGAACACCTTCGGTA
>hsa-miR-4482-5p
AACCCAGTGGGCTATGGAAATG
>hsa-miR-2276-5p
GCCCTCTGTCACCTTGCAGACG
>hsa-miR-106b-3p
CCGCACTGTGGGTACTTGCTGC
>hsa-miR-30a-3p
CTTTCAGTCGGATGTTTGCAGC
>hsa-miR-9851-3p
TGGCACCAGCACTGGCGGTGTC
>hsa-miR-4425
TGTTGGGATTCAGCAGGACCAT
>hsa-miR-6759-5p
TTGTGGGTGGGCAGAAGTCTGT
>hsa-miR-6722-5p
AGGCGCACCCGACCACATGC
>hsa-miR-569
AGTTAATGAATCCTGGAAAGT
>hsa-miR-4744
TCTAAAGACTAGACTTCGCTATG
>hsa-miR-128-2-5p
GGGGGCCGATACACTGTACGAGA
>hsa-miR-4522
TGACTCTGCCTGTAGGCCGGT
>hsa-miR-708-5p
AAGGAGCTTACAATCTAGCTGGG
>hsa-miR-376a-2-5p
GGTAGATTTTCCTTCTATGGT
>hsa-miR-6783-5p
TAGGGGAAAAGTCCTGATCCGG
>hsa-miR-4660
TGCAGCTCTGGTGGAAAATGGAG
>hsa-miR-1255b-5p
CGGATGAGCAAAGAAAGTGGTT
>hsa-miR-4682
TCTGAGTTCCTGGAGCCTGGTCT
>hsa-miR-4533
TGGAAGGAGGTTGCCGGACGCT
>hsa-miR-564
AGGCACGGTGTCAGCAGGC
>hsa-miR-149-5p
TCTGGCTCCGTGTCTTCACTCCC
>hsa-miR-4726-3p
ACCCAGGTTCCCTCTGGCCGCA
>hsa-miR-4745-3p
TGGCCCGGCGACGTCTCACGGTC
>hsa-miR-3622b-5p
AGGCATGGGAGGTCAGGTGA
>hsa-miR-190a-3p
CTATATATCAAACATATTCCT
>hsa-miR-378e
ACTGGACTTGGAGTCAGGA
>hsa-miR-9718
TTGCTGACCTGGGTGGTGGT
>hsa-miR-7107-3p
TGGTCTGTTCATTCTCTCTTTTTGGCC
>hsa-miR-7152-5p
TTTCCTGTCCTCCAACCAGACC

>hsa-miR-1208
TCACTGTTCAGACAGGCGGA
>hsa-miR-661
TGCCTGGGTCTCTGGCCTGCGCGT
>hsa-miR-125a-5p
TCCCTGAGACCCTTTAACCTGTGA
>hsa-miR-1238-5p
GTGAGTGGGAGCCCCAGTGTGTG
>hsa-miR-519c-5p|hsa-miR-519b-5p|hsa-miR-523-5p|hsa-miR-518e-5p|hsa-miR-522-5p|hsa-miR-519a-5p
CTCTAGAGGGAAGCGCTTTCTG
>hsa-miR-939-3p
CCCTGGGCCTCTGCTCCCCAG
>hsa-miR-340-5p
TTATAAAGCAATGAGACTGATT
>hsa-miR-6879-5p
CAGGGCAGGGAAGGTGGGAGAG
>hsa-miR-6806-3p
TGAAGCTCTGACATTCCTGCAG
>hsa-miR-219b-5p
AGATGTCCAGCCACAATTCTCG
>hsa-miR-1286
TGCAGGACCAAGATGAGCCCT
>hsa-miR-1203
CCCGGAGCCAGGATGCAGCTC
>hsa-miR-503-5p
TAGCAGCGGGAACAGTTCTGCAG
>hsa-miR-16-5p
TAGCAGCACGTAAATATTGGCG
>hsa-miR-4494
CCAGACTGTGGCTGACCAGAGG
>hsa-miR-4256
ATCTGACCTGATGAAGGT
>hsa-miR-3182
GCTTCTGTAGTGTAGTC
>hsa-miR-10396b-3p
GGCCCCGGGCCCTCGACCGGAC
>hsa-miR-3940-5p
GTGGGTTGGGGCGGGCTCTG
>hsa-miR-548j-3p
CAAAAACTGCATTACTTTTGC
>hsa-miR-1226-3p
TCACCAGCCCTGTGTTCCCTAG
>hsa-miR-513c-5p
TTCTCAAGGAGGTGTCGTTTAT
>hsa-miR-3616-3p
CGAGGGCATTTCATGATGCAGGC
>hsa-miR-6768-3p
CAAAGGCCACATTCTCCTGTGCAC
>hsa-miR-30c-5p
TGTAAACATCCTACACTCTCAGC
>hsa-miR-4784
TGAGGAGATGCTGGGACTGA
>hsa-miR-4771
AGCAGACTTGACCTACAATTA
>hsa-miR-3171
AGATGTATGGAATCTGTATATATC
>hsa-miR-1181
CCGTCGCCGCCACCCGAGCCG
>hsa-miR-10392-3p
CCGGCCCCGCCTCGGCTCCGCACC
>hsa-miR-6753-3p

TGGTCTGTCTCTGCCCTGGCAC
>hsa-miR-423-3p
AGCTCGGTCTGAGGCCCCTCAGT
>hsa-miR-4507
CTGGGTTGGGCTGGGCTGGG
>hsa-miR-1185-5p
AGAGGATACCCTTTGTATGTT
>hsa-miR-5187-5p
TGGGATGAGGGATTGAAGTGGA
>hsa-miR-100-5p
AACCCGTAGATCCGAACTTGTG
>hsa-miR-6771-5p
CTCGGGAGGGCATGGGCCAGGC
>hsa-miR-4758-5p
GTGAGTGGGAGCCGGTGGGGCTG
>hsa-miR-523-3p
GAACGCGCTTCCCTATAGAGGGT
>hsa-miR-4322
CTGTGGGCTCAGCGCGTGGGG
>hsa-miR-29a-5p
ACTGATTTCTTTTGGTGTTCAG
>hsa-miR-4632-3p
TGCCGCCCTCTCGCTGCTCTAG
>hsa-miR-99a-3p
CAAGCTCGCTTCTATGGGTCTG
>hsa-miR-632
GTGTCTGCTTCCTGTGGGA
>hsa-miR-4529-5p
AGGCCATCAGCAGTCCAATGAA
>hsa-miR-4470
TGGCAAACGTGGAAGCCGAGA
>hsa-miR-4427
TCTGAATAGAGTCTGAAGAGT
>hsa-miR-4664-3p
CTTCCGGTCTGTGAGCCCCGTC
>hsa-miR-4778-3p
TCTTCTTCCTTTGCAGAGTTGA
>hsa-miR-4330
CCTCAGATCAGAGCCTTGC
>hsa-miR-5697
TCAAGTAGTTTCATGATAAAGG
>hsa-miR-4764-5p
TGGATGTGGAAGGAGTTATCT
>hsa-miR-4438
CACAGGCTTAGAAAAGACAGT
>hsa-miR-646
AAGCAGCTGCCTCTGAGGC
>hsa-miR-1251-5p
ACTCTAGCTGCCAAAGGCGCT
>hsa-miR-563
AGGTTGACATACGTTTCCC
>hsa-miR-4421
ACCTGTCTGTGGAAAGGAGCTA
>hsa-miR-499b-5p
ACAGACTTGCTGTGATGTTCA
>hsa-miR-4670-5p
AAGCGACCATGATGTAACTTCA
>hsa-miR-8066
CAATGTGATCTTTTGGATGTA
>hsa-miR-6512-5p
TACCATTAGAAGAGCTGGAAGA
>hsa-miR-3605-5p

TGAGGATGGATAGCAAGGAAGCC
>hsa-miR-6742-5p
AGTGGGGTGGGACCCAGCTGTT
>hsa-let-7g-3p
CTGTACAGGCCACTGCCTTGC
>hsa-miR-589-5p
TGAGAACCACGTCTGCTCTGAG
>hsa-miR-3609
CAAAGTGATGAGTAATACTGGCTG
>hsa-miR-6729-3p
TCATCCCCCTCGCCCTCTCAG
>hsa-miR-6507-3p
CAAAGTCCTTCCTATTTTTCCC
>hsa-miR-6796-3p
GAAGCTCTCCCCTCCCCGCAG
>hsa-miR-3620-3p
TCACCCTGCATCCCGCACCCAG
>hsa-miR-624-3p
CACAAGGTATTGGTATTACCT
>hsa-miR-6809-3p
CTTCTCTTCTCTCCTTCCCAG
>hsa-miR-4496
GAGGAAACTGAAGCTGAGAGGG
>hsa-miR-1273h-3p
CTGCAGACTCGACCTCCCAGGC
>hsa-miR-4525
GGGGGGATGTGCATGCTGGTT
>hsa-miR-493-5p
TTGTACATGGTAGGCTTTCATT
>hsa-miR-337-3p
CTCCTATATGATGCCTTTCTTC
>hsa-miR-196a-5p
TAGGTAGTTTCATGTTGTTGGG
>hsa-miR-4277
GCAGTTCTGAGCACAGTACAC
>hsa-miR-1204
TCGTGGCCTGGTCTCCATTAT
>hsa-let-7a-3p
CTATACAATCTACTGTCTTTC
>hsa-miR-221-3p
AGCTACATTGTCTGCTGGGTTTC
>hsa-miR-7161-3p
TAGATCTTTGACTCTGGCAGTCTCCAGG
>hsa-miR-6747-3p
TCCTGCCTTCCTCTGCACCAG
>hsa-miR-4797-3p
TCTCAGTAAGTGGCACTCTGT
>hsa-miR-4700-5p
TCTGGGGATGAGGACAGTGTGT
>hsa-miR-1972
TCAGGCCAGGCACAGTGGCTCA
>hsa-miR-5694
CAGATCATGGGACTGTCTCAG
>hsa-miR-6860
ACTGGGCAGGGCTGTGGTGAGT
>hsa-miR-1178-5p
CAGGGTCAGCTGAGCATG
>hsa-miR-598-3p
TACGTCATCGTTGTCATCGTCA
>hsa-miR-2681-3p
TATCATGGAGTTGGTAAAGCAC
>hsa-miR-1587

TTGGGCTGGGCTGGGTTGGG
>hsa-miR-1257
AGTGAATGATGGGTTCTGACC
>hsa-miR-6764-3p
TCTCTGGTCTTTCCTTGACAG
>hsa-miR-203b-5p
TAGTGGTCCTAAACATTTCACA
>hsa-miR-1247-3p
CCCCGGGAACGTCGAGACTGGAGC
>hsa-miR-455-3p
GCAGTCCATGGGCATATACAC
>hsa-miR-6503-3p
GGGACTAGGATGCAGACCTCC
>hsa-miR-8086
TGCTAGTCTGGACTGATATGGT
>hsa-miR-6772-3p
TTGCTCCTGACTCTGTGCCCACA
>hsa-miR-8069
GGATGGTTGGGGGCGGTCGGCGT
>hsa-miR-6832-5p
AGTAGAGAGGAAAAGTTAGGGTC
>hsa-miR-4774-5p
TCTGGTATGTAGTAGGTAATAA
>hsa-miR-4689
TTGAGGAGACATGGTGGGGGCC
>hsa-miR-320a-5p
GCCTTCTCTTCCCGGTTCTTCC
>hsa-miR-10393-3p
TGGTCAGATTTGAACTCTTCAA
>hsa-miR-7153-5p
TGAGAACTGACAAATGTGGTAGG
>hsa-miR-4717-3p
ACACATGGGTGGCTGTGGCCT
>hsa-miR-130a-5p
GCTCTTTTCACATTGTGCTACT
>hsa-miR-4799-3p
ACTGGCATGCTGCATTTATATA
>hsa-miR-3616-5p
ATGAAGTGCACTCATGATATGT
>hsa-miR-425-5p
AATGACACGATCACTCCCGTTGA
>hsa-miR-4276
CTCAGTGACTCATGTGC
>hsa-miR-3127-3p
TCCCCTTCTGCAGGCCTGCTGG
>hsa-miR-6718-5p
TAGTGGTCAGAGGGCTTATGA
>hsa-miR-378c
ACTGGACTTGGAGTCAGAAGAGTGG
>hsa-miR-6781-3p
TGCCTCTTTTCCACGGCCTCAG
>hsa-miR-6840-5p
ACCCCCGGGCAAAGACCTGCAGAT
>hsa-miR-5047
TTGCAGCTGCGGTTGTAAGGT
>hsa-miR-301a-5p
GCTCTGACTTTATTGCACTACT
>hsa-miR-7153-3p
CACCATGGACGGTTTACC
>hsa-miR-4729
TCATTTATCTGTTGGGAAGCTA
>hsa-miR-3189-3p

CCCTTGGGTCTGATGGGGTAG
>hsa-miR-5590-5p
TTGCCATACATAGACTTTATT
>hsa-miR-5579-5p
TATGGTACTCCTTAAGCTAAC
>hsa-miR-548s
ATGGCCAAAACTGCAGTTATTTT
>hsa-miR-3714
GAAGGCAGCAGTGCTCCCCTGT
>hsa-miR-6745
TGGGTGGAAGAAGGTCTGGTT
>hsa-miR-7704
CGGGGTCGGCGGCGACGTG
>hsa-miR-4457
TCACAAGGTATTGACTGGCGTA
>hsa-miR-7855-5p
TTGGTGAGGACCCCAAGCTCGG
>hsa-miR-4703-5p
TAGCAATACAGTACAAATATAGT
>hsa-miR-611
GCGAGGACCCCTCGGGGTCTGAC
>hsa-miR-7110-3p
TCTCTCTCCCACTTCCCTGCAG
>hsa-miR-190a-5p
TGATATGTTTGATATATTAGGT
>hsa-miR-593-5p
AGGCACCAGCCAGGCATTGCTCAGC
>hsa-miR-6834-5p
GTGAGGGACTGGGATTTGTGG
>hsa-miR-1264
CAAGTCTTATTTGAGCACCTGTT
>hsa-miR-6130
TGAGGGAGTGGATTGTATG
>hsa-miR-4446-5p
ATTTCCCTGCCATTCCCTTGGC
>hsa-miR-6740-3p
TGTCTTCTCTCCTCCCAAACAG
>hsa-miR-3202
TGGAAGGGAGAAGAGCTTTAAT
>hsa-miR-214-3p
ACAGCAGGCACAGACAGGCAGT
>hsa-miR-370-3p
GCCTGCTGGGGTGGAACCTGGT
>hsa-miR-509-3-5p
TACTGCAGACGTGGCAATCATG
>hsa-miR-184
TGGACGGAGAACTGATAAGGGT
>hsa-miR-4447
GGTGGGGGCTGTTGTTT
>hsa-miR-302a-5p
ACTTAAACGTGGATGTACTTGCT
>hsa-miR-10394-3p
TGGGCGCGCCGGGACTGTGAGAC
>hsa-miR-4455
AGGGTGTGTGTGTTTTT
>hsa-miR-6514-3p
CTGCCTGTTCTTCCACTCCAG
>hsa-miR-6893-3p
CCCTGCTGCCTTCACCTGCCAG
>hsa-miR-9-5p
TCTTTGGTTATCTAGCTGTATGA
>hsa-miR-6815-5p

TAGGTGGCGCCGGAGGAGTCATT
>hsa-miR-541-5p
AAAGGATTCTGCTGTCGGTCCCACT
>hsa-miR-6807-3p
CACTGCATTCCTGCTTGGCCCAG
>hsa-miR-6502-5p
AGCTCTAGAAAGATTGTTGACC
>hsa-miR-5196-5p
AGGGAAGGGGACGAGGGTTGGG
>hsa-miR-1225-5p
GTGGGTACGGCCCAGTGGGGGG
>hsa-miR-381-5p
AGCGAGGTTGCCCTTTGTATAT
>hsa-miR-367-5p
ACTGTTGCTAATATGCAACTCT
>hsa-miR-1275
GTGGGGGAGAGGCTGTC
>hsa-miR-4720-5p
CCTGGCATATTTGGTATAACTT
>hsa-miR-7157-3p
TCTGTGCTACTGGATGAAGAGT
>hsa-miR-3667-3p
ACCTTCCTCTCCATGGGTCTTT
>hsa-miR-4672
TTACACAGCTGGACAGAGGCA
>hsa-miR-363-5p
CGGGTGGATCACGATGCAATTT
>hsa-miR-1202
GTGCCAGCTGCAGTGGGGGAG
>hsa-miR-922
GCAGCAGAGAATAGGACTACGTC
>hsa-miR-139-5p
TCTACAGTGCACGTGTCTCCAGT
>hsa-miR-548n
CAAAAGTAATTGTGGATTTTGT
>hsa-miR-23b-3p
ATCACATTGCCAGGGATTACCAC
>hsa-miR-653-5p
GTGTTGAAACAATCTCTACTG
>hsa-miR-1296-3p
GAGTGGGGCTTCGACCCTAACC
>hsa-miR-1268b
CGGGCGTGGTGGTGGGGGTG
>hsa-miR-1270
CTGGAGATATGGAAGAGCTGTGT
>hsa-miR-766-3p
ACTCCAGCCCCACAGCCTCAGC
>hsa-miR-583
CAAAGAGGAAGGTCCCATTAC
>hsa-miR-10b-3p
ACAGATTCGATTCTAGGGGAAT
>hsa-miR-5197-5p
CAATGGCACAAACTCATTCTTGA
>hsa-miR-3197
GGAGGCGCAGGCTCGGAAAGGCG
>hsa-miR-4279
CTCTCCTCCCGGCTTC
>hsa-miR-136-5p
ACTCCATTTGTTTTGATGATGGA
>hsa-miR-4700-3p
CACAGGACTGACTCCTCACCCCAGTG
>hsa-miR-6877-5p

AGGGCCGAAGGGTGGAAGCTGC
>hsa-miR-4724-3p
GTACCTTCTGGTTCAGCTAGT
>hsa-miR-296-5p
AGGGCCCCCCCTCAATCCTGT
>hsa-miR-6862-5p
CGGGCATGCTGGGAGAGACTTT
>hsa-miR-7159-5p
TTCAACAAGGGTGTAGGATGG
>hsa-miR-3150b-3p
TGAGGAGATCGTCGAGGTTGG
>hsa-miR-5002-5p
AATTTGGTTTCTGAGGCACTTAGT
>hsa-miR-4707-5p
GCCCCGGCGCGGGCGGGTTCTGG
>hsa-miR-3124-5p
TTCGCGGGCGAAGGCAAAGTC
>hsa-miR-3679-5p
TGAGGATATGGCAGGGAAGGGGA
>hsa-miR-5000-5p
CAGTTCAGAAGTGTTCCTGAGT
>hsa-miR-1914-3p
GGAGGGGTCCCGCACTGGGAGG
>hsa-miR-6780a-5p
TTGGGAGGGAAGACAGCTGGAGA
>hsa-miR-2110
TTGGGGAAACGGCCGCTGAGTG
>hsa-miR-32-5p
TATTGCACATTACTAAGTTGCA
>hsa-miR-1248
ACCTTCTTGTATAAGCACTGTGCTAAA
>hsa-miR-374b-3p
CTTAGCAGGTTGTATTATCATT
>hsa-miR-662
TCCCACGTTGTGGCCCAGCAG
>hsa-let-7f-1-3p
CTATACAATCTATTGCCTTCCC
>hsa-miR-4499
AAGACTGAGAGGAGGGA
>hsa-miR-4293
CAGCCTGACAGGAACAG
>hsa-miR-4485-3p
TAACGGCCGCGGTACCCTAA
>hsa-miR-3663-3p
TGAGCACCACACAGGCCGGGCGC
>hsa-miR-219a-5p
TGATTGTCCAAACGCAATTCT
>hsa-miR-3177-5p
TGTGTACACACGTGCCAGGCGCT
>hsa-miR-548aq-3p
CAAAAACTGCAATTACTTTTGC
>hsa-miR-6855-3p
AGACTGACCTTCAACCCCACAG
>hsa-miR-5584-3p
TAGTTCTTCCCTTTGCCCAATT
>hsa-miR-208b-5p
AAGCTTTTTGCTCGAATTATGT
>hsa-miR-10226
CCATGTGTCTGGGCTGGGAAAC
>hsa-miR-320c
AAAAGCTGGGTTGAGAGGGT
>hsa-miR-140-5p

CAGTGGTTTTACCCTATGGTAG
>hsa-miR-4767
CGCGGGCGCTCCTGGCCGCCGCC
>hsa-miR-22-5p
AGTTCTTCAGTGGCAAGCTTTA
>hsa-miR-4804-3p
TGCTTAACCTTGCCCTCGAAA
>hsa-miR-4733-3p
CCACCAGGTCTAGCATTGGGAT
>hsa-miR-6808-5p
CAGGCAGGGAGGTGGGACCATG
>hsa-miR-4643
GACACATGACCATAAATGCTAA
>hsa-miR-539-3p
ATCATACAAGGACAATTTCTTT
>hsa-miR-3194-5p
GGCCAGCCACCAGGAGGGCTG
>hsa-miR-8054
GAAAGTACAGATCGGATGGGT
>hsa-miR-544a
ATTCTGCATTTTTAGCAAGTTC
>hsa-miR-6815-3p
TGGCTTCTCTTGCACACCCAG
>hsa-miR-548av-5p
AAAAGTACTTGCGGATTT
>hsa-miR-23a-5p
GGGGTTCCTGGGGATGGGATTT
>hsa-miR-4478
GAGGCTGAGCTGAGGAG
>hsa-miR-3144-5p
AGGGGACCAAAGAGATATATAG
>hsa-miR-6851-3p
TGGCCCTTTGTACCCCTCCAG
>hsa-miR-6515-5p
TTGGAGGGTGTGGAAGACATC
>hsa-miR-3605-3p
CCTCCGTGTTACCTGTCCTCTAG
>hsa-miR-1295b-5p
CACCCAGATCTGCGGCCTAAT
>hsa-miR-3674
ATTGTAGAACCTAAGATTGGCC
>hsa-miR-658
GGCGGAGGGAAGTAGGTCCGTTGGT
>hsa-miR-6773-3p
ACTGTCACTTCTCTGCCCATAG
>hsa-miR-7160-5p
TGCTGAGGTCCGGGCTGTGCC
>hsa-miR-330-3p
GCAAAGCACACGGCCTGCAGAGA
>hsa-miR-9898
TACTTACCTGTCCCCTACCCCA
>hsa-miR-197-5p
CGGGTAGAGAGGGCAGTGGGAGG
>hsa-miR-450a-2-3p
ATTGGGGACATTTTGCATTCAT
>hsa-miR-5004-3p
CTTGGATTTTCCTGGGCCTCAG
>hsa-miR-4704-3p
TCAGTCACATATCTAGTGTCTA
>hsa-miR-3158-5p
CCTGCAGAGAGGAAGCCCTTC
>hsa-miR-5699-3p

TCCTGTCTTTCCTTGTTGGAGC
>hsa-miR-3170
CTGGGGTTCTGAGACAGACAGT
>hsa-miR-4701-5p
TTGGCCACCACACCTACCCCTT
>hsa-miR-6769a-3p
GAGCCCCTCTCTGCTCTCCAG
>hsa-miR-2277-5p
AGCGCGGGCTGAGCGCTGCCAGTC
>hsa-miR-6748-5p
TGTGGGTGGGAAGGACTGGATT
>hsa-miR-370-5p
CAGGTCACGTCTCTGCAGTTAC
>hsa-miR-7856-5p
TTTTAAGGACACTGAGGGATC
>hsa-miR-6849-5p
GAGTGGATAGGGGAGTGTGTGGA
>hsa-miR-920
GGGGAGCTGTGGAAGCAGTA
>hsa-miR-6802-5p
CTAGGTGGGGGGCTTGAAGC
>hsa-miR-4282
TAAAATTTGCATCCAGGA
>hsa-miR-454-5p
ACCCTATCAATATTGTCTCTGC
>hsa-miR-211-5p
TTCCCTTTGTCATCCTTCGCCT
>hsa-miR-5787
GGGCTGGGGCGCGGGGAGGT
>hsa-miR-2278
GAGAGCAGTGTGTGTTGCCTGG
>hsa-miR-1293
TGGGTGGTCTGGAGATTTGTGC
>hsa-miR-22-3p
AAGCTGCCAGTTGAAGAACTGT
>hsa-miR-3123
CAGAGAATTGTTTAATC
>hsa-miR-30d-3p
CTTTCAGTCAGATGTTTGCTGC
>hsa-miR-214-5p
TGCCTGTCTACACTTGCTGTGC
>hsa-miR-3650
AGGTGTGTCTGTAGAGTCC
>hsa-miR-3146
CATGCTAGGATAGAAAGAATGG
>hsa-miR-338-5p
AACAATATCCTGGTGCTGAGTG
>hsa-miR-3065-5p
TCAACAAAATCACTGATGCTGGA
>hsa-miR-6859-5p
GAGAGGAACATGGGCTCAGGACA
>hsa-miR-6165
CAGCAGGAGGTGAGGGGAG
>hsa-miR-4537
TGAGCCGAGCTGAGCTTAGCTG
>hsa-miR-631
AGACCTGGCCCAGACCTCAGC
>hsa-miR-1226-5p
GTGAGGGCATGCAGGCCTGGATGGGG
>hsa-miR-3135a
TGCCTAGGCTGAGACTGCAGTG
>hsa-miR-103a-1-5p

GGCTTCTTTACAGTGCTGCCTTG
>hsa-miR-26b-5p
TTCAAGTAATTCAGGATAGGT
>hsa-miR-4762-5p
CCAAATCTTGATCAGAAGCCT
>hsa-miR-5189-3p
TGCCAACCGTCAGAGCCCAGA
>hsa-miR-525-3p
GAAGGCGCTTCCCTTTAGAGCG
>hsa-miR-9986
TGTGAGGTTGTCATGCCTGC
>hsa-miR-27b-3p
TTCACAGTGGCTAAGTTCTGC
>hsa-miR-373-5p
ACTCAAAATGGGGGCGCTTTCC
>hsa-miR-4661-5p
AACTAGCTCTGTGGATCCTGAC
>hsa-miR-543
AAACATTCGCGGTGCACTTCTT
>hsa-miR-513a-5p
TTCACAGGGAGGTGTCAT
>hsa-miR-2117
TGTTCTCTTTGCCAAGGACAG
>hsa-miR-30e-5p
TGTAAACATCCTTGACTGGAAG
>hsa-miR-3151-5p
GGTGGGGCAATGGGATCAGGT
>hsa-miR-587
TTTCCATAGGTGATGAGTCAC
>hsa-miR-382-5p
GAAGTTGTTCGTGGTGGATTCG
>hsa-miR-516b-5p
ATCTGGAGGTAAGAAGCACTTT
>hsa-miR-30c-2-3p
CTGGGAGAAGGCTGTTTACTCT
>hsa-miR-6084
TTCCGCCAGTCGGTGGCCGG
>hsa-miR-514b-5p
TTCTCAAGAGGGAGGCAATCAT
>hsa-miR-1250-5p
ACGGTGCTGGATGTGGCCTTT
>hsa-miR-19b-2-5p
AGTTTTGCAGGTTTGCATTTCA
>hsa-miR-3688-5p
AGTGGCAAAGTCTTTCCATAT
>hsa-miR-6821-3p
TGACCTCTCCGCTCCGCACAG
>hsa-miR-548u
CAAAGACTGCAATTACTTTTGCG
>hsa-miR-544b
ACCTGAGGTTGTGCATTTCTAA
>hsa-miR-1231
GTGTCTGGGCGGACAGCTGC
>hsa-miR-3614-5p
CCACTTGGATCTGAAGGCTGCCC
>hsa-miR-6875-5p
TGAGGGACCCAGGACAGGAGA
>hsa-miR-503-3p
GGGGTATTGTTTCCGCTGCCAGG
>hsa-miR-6892-5p
GTAAGGGACCGGAGAGTAGGA
>hsa-miR-3976

TATAGAGAGCAGGAAGATTAATGT
>hsa-miR-5088-3p
TCCCTTCTTCCTGGGCCCTCA
>hsa-miR-3666
CAGTGCAAGTGTAGATGCCGA
>hsa-miR-6750-5p
CAGGGAACAGCTGGGTGAGCTGCT
>hsa-miR-2114-5p
TAGTCCCTTCCTTGAAGCGGTC
>hsa-miR-5583-5p
AAACTAATATACCCATATTCTG
>hsa-miR-577
TAGATAAAATATTGGTACCTG
>hsa-miR-4688
TAGGGGCAGCAGAGGACCTGGG
>hsa-miR-1909-3p
CGCAGGGGCCGGGTGCTCACCG
>hsa-miR-675-5p
TGGTGCGGAGAGGGCCCACAGTG
>hsa-miR-495-3p
AAACAAACATGGTGCACTTCTT
>hsa-miR-4726-5p
AGGGCCAGAGGAGCCTGGAGTGG
>hsa-miR-548ab
AAAAGTAATTGTGGATTTTGCT
>hsa-miR-4315
CCGCTTTCTGAGCTGGAC
>hsa-miR-1228-3p
TCACACCTGCCTCGCCCCCC
>hsa-miR-4650-5p
TCAGGCCTCTTTCTACCTT
>hsa-miR-4326
TGTTCCTCTGTCTCCCAGAC
>hsa-miR-98-5p
TGAGGTAGTAAGTTGTATTGTT
>hsa-miR-548bb-5p
AAAAGTAACTATGGTTTTTGCC
>hsa-miR-4633-3p
AGGAGCTAGCCAGGCATATGCA
>hsa-miR-4325
TTGCACTTGTCTCAGTGA
>hsa-miR-4493
AGAAGGCCTTTCCATCTCTGT
>hsa-miR-3690
ACCTGGACCCAGCGTAGACAAAG
>hsa-miR-1180-3p
TTTCCGGCTCGCGTGGGTGTGT
>hsa-miR-607
GTTCAAATCCAGATCTATAAC
>hsa-miR-6765-3p
TCACCTGGCTGGCCCGCCCAG
>hsa-miR-6798-5p
CCAGGGGGATGGGCGAGCTTGGG
>hsa-miR-203a-3p
GTGAAATGTTTAGGACCACTAG
>hsa-miR-433-3p
ATCATGATGGGCTCCTCGGTGT
>hsa-miR-4741
CGGGCTGTCCGGAGGGGTCGGCT
>hsa-miR-3667-5p
AAAGACCCATTGAGGAGAAGGT
>hsa-miR-1909-5p

TGAGTGCCGGTGCCTGCCCTG
>hsa-miR-1299
TTCTGGAATTCTGTGTGAGGGA
>hsa-miR-452-5p
AACTGTTTGCAGAGGAAACTGA
>hsa-miR-3173-3p
AAAGGAGGAAATAGGCAGGCCA
>hsa-miR-203a-5p
AGTGGTTCTTAACAGTTCAACAGTT
>hsa-miR-6803-3p
TCCCTCGCCTTCTCACCCTCAG
>hsa-miR-103b
TCATAGCCCTGTACAATGCTGCT
>hsa-miR-4783-3p
CCCCGGTGTTGGGGCGCGTCTGC
>hsa-miR-3201
GGGATATGAAGAAAAAT
>hsa-miR-1537-5p
AGCTGTAATTAGTCAGTTTTCT
>hsa-miR-380-3p
TATGTAATATGGTCCACATCTT
>hsa-miR-6503-5p
AGGTCTGCATTCAAATCCCCAGA
>hsa-miR-7154-5p
TTCATGAACTGGGTCTAGCTTGG
>hsa-miR-194-3p
CCAGTGGGGCTGCTGTTATCTG
>hsa-miR-369-5p
AGATCGACCGTGTTATATTCGC
>hsa-miR-3919
GCAGAGAACAAAGGACTCAGT
>hsa-miR-4782-3p
TGATTGTCTTCATATCTAGAAC
>hsa-miR-6838-5p
AAGCAGCAGTGGCAAGACTCCT
>hsa-miR-3143
ATAACATTGTAAAGCGCTTCTTTCG
>hsa-miR-369-3p
AATAATACATGGTTGATCTTT
>hsa-miR-297
ATGTATGTGTGCATGTGCATG
>hsa-miR-875-5p
TATACCTCAGTTTTATCAGGTG
>hsa-miR-6789-3p
CGGCGCCCGTGTCTCCTCCAG
>hsa-miR-7848-3p
CTACCCTCGGTCTGCTTACCACA
>hsa-miR-3160-3p
AGAGCTGAGACTAGAAAGCCCA
>hsa-miR-876-5p
TGGATTTCTTTGTGAATCACCA
>hsa-miR-7152-3p
TCTGGTCCTGGACAGGAGGC
>hsa-miR-3188
AGAGGCTTTGTGCGGATACGGGG
>hsa-miR-4757-5p
AGGCCTCTGTGACGTCACGGTGT
>hsa-miR-4475
CAAGGGACCAAGCATTCATTAT
>hsa-miR-199b-5p
CCCAGTGTTTAGACTATCTGTTC
>hsa-miR-330-5p

TCTCTGGGCCTGTGTCTTAGGC
>hsa-miR-548ah-3p
CAAAAACTGCAGTTACTTTTGC
>hsa-miR-515-3p
GAGTGCCTTCTTTTGGAGCGTT
>hsa-miR-183-5p
TATGGCACTGGTAGAATTCACT
>hsa-miR-29b-3p
TAGCACCATTTGAAATCAGTGTT
>hsa-miR-548j-5p
AAAAGTAATTGCGGTCTTTGGT
>hsa-miR-4526
GCTGACAGCAGGGCTGGCCGCT
>hsa-miR-936
ACAGTAGAGGGAGGAATCGCAG
>hsa-miR-494-3p
TGAAACATACACGGGAAACCTC
>hsa-miR-518c-3p
CAAAGCGCTTCTCTTTAGAGTGT
>hsa-miR-6762-5p
CGGGGCCATGGAGCAGCCTGTGT
>hsa-miR-196a-1-3p
CAACAACATTAAACCACCCGA
>hsa-miR-1910-3p
GAGGCAGAAGCAGGATGACA
>hsa-miR-1-3p
TGGAATGTAAAGAAGTATGTAT
>hsa-miR-335-5p
TCAAGAGCAATAACGAAAAATGT
>hsa-miR-4680-3p
TCTGAATTGTAAGAGTTGTTA
>hsa-miR-4718
AGCTGTACCTGAAACCAAGCA
>hsa-let-7e-5p
TGAGGTAGGAGGTTGTATAGTT
>hsa-miR-4435
ATGGCCAGAGCTCACACAGAGG
>hsa-miR-4274
CAGCAGTCCCTCCCCCTG
>hsa-miR-580-5p
TAATGATTCATCAGACTCAGAT
>hsa-miR-3200-5p
AATCTGAGAAGGCGCACAAGGT
>hsa-miR-6886-3p
TGCCCTTCTCTCCTCCTGCCT
>hsa-miR-4781-5p
TAGCGGGGATTCCAATATTGG
>hsa-miR-450a-1-3p
ATTGGGAACATTTTGCATGTAT
>hsa-miR-6080
TCTAGTGCGGGCGTTCCCG
>hsa-miR-6830-3p
TGTCTTTCTTCTCTCCCTTGCAG
>hsa-miR-8082
TGATGGAGCTGGGAATACTCTG
>hsa-miR-6769b-5p
TGGTGGGTGGGGAGGAGAAGTGC
>hsa-miR-1179
AAGCATTCTTTCATTGGTTGG
>hsa-miR-6854-5p
AAGCTCAGGTTTGAGAACTGCTGA
>hsa-miR-200c-3p

TAATACTGCCGGGTAATGATGGA
>hsa-miR-20a-3p
ACTGCATTATGAGCACTTAAAG
>hsa-miR-196b-5p
TAGGTAGTTTCCTGTTGTTGGG
>hsa-miR-6836-5p
CGCAGGGCCCTGGCGCAGGCAT
>hsa-miR-4768-5p
ATTCTCTCTGGATCCCATGGAT
>hsa-miR-525-5p
CTCCAGAGGGATGCACTTTCT
>hsa-miR-1282
TCGTTTGCCTTTTTCTGCTT
>hsa-miR-135b-5p
TATGGCTTTTCATTCCTATGTGA
>hsa-miR-4485-5p
ACCGCCTGCCCAGTGA
>hsa-miR-548ak
AAAAGTAACTGCGGTTTTTGA
>hsa-miR-9-3p
ATAAAGCTAGATAACCGAAAGT
>hsa-miR-147b-5p
TGGAAACATTTCTGCACAAACT
>hsa-miR-34b-5p
TAGGCAGTGTCATTAGCTGATTG
>hsa-miR-6795-3p
ACCCCTCGTTTCTTCCCCCAG
>hsa-miR-6797-5p
AGGAGGGAAGGGGCTGAGAACAGGA
>hsa-miR-4757-3p
CATGACGTCACAGAGGCTTCGC
>hsa-miR-33b-3p
CAGTGCCTCGGCAGTGCAGCCC
>hsa-miR-3914
AAGGAACCAGAAAATGAGAAGT
>hsa-miR-3606-3p
AAAATTTCTTTCACTACTTAG
>hsa-miR-1292-5p
TGGGAACGGGTTCCGGCAGACGCTG
>hsa-miR-193a-3p
AACTGGCCTACAAAGTCCCAGT
>hsa-miR-6889-3p
TCTGTGCCCCCTACTTCCCAG
>hsa-miR-6792-3p
CTCCTCCACAGCCCCTGCTCAT
>hsa-miR-708-3p
CAACTAGACTGTGAGCTTCTAG
>hsa-miR-216a-5p
TAATCTCAGCTGGCAACTGTGA
>hsa-miR-3064-3p
TTGCCACACTGCAACACCTTACA
>hsa-miR-296-3p
GAGGGTTGGGTGGAGGCTCTCC
>hsa-miR-6736-5p
CTGGGTGAGGGCATCTGTGGT
>hsa-miR-6870-5p
TGGGGGAGATGGGGGTTGA
>hsa-miR-548x-3p
TAAAAACTGCAATTACTTTC
>hsa-miR-1322
GATGATGCTGCTGATGCTG
>hsa-miR-216b-3p

ACACACTTACCCGTAGAGATTCTA
>hsa-miR-4301
TCCCACTACTTCACTTGTGA
>hsa-miR-3141
GAGGGCGGGTGGAGGAGGA
>hsa-miR-3187-3p
TTGGCCATGGGGCTGCGCGG
>hsa-miR-1263
ATGGTACCCTGGCATACTGAGT
>hsa-miR-6728-3p
TCTCTGCTCTGCTCTCCCCAG
>hsa-miR-5002-3p
TGACTGCCTCACTGACCACTT
>hsa-miR-501-5p
AATCCTTTGTCCCTGGGTGAGA
>hsa-miR-6819-3p
AAGCCTCTGTCCCCACCCCAG
>hsa-miR-6877-3p
CAGCCTCTGCCCTTGGCCTCC
>hsa-miR-3167
AGGATTTCAGAAATACTGGTGT
>hsa-miR-4787-5p
GCGGGGGTGGCGGCGGCATCCC
>hsa-miR-4754
ATGCGGACCTGGGTTAGCGGAGT
>hsa-miR-10393-5p
AGAATTCTCTTATCCAACATCAACA
>hsa-miR-6758-5p
TAGAGAGGGGAAGGATGTGATGT
>hsa-miR-122b-3p
AAACACCATTGTCACACTCCAC
>hsa-miR-5192
AGGAGAGTGGATTCCAGGTGGT
>hsa-miR-182-3p
TGGTTCTAGACTTGCCAACTA
>hsa-miR-5579-3p
TTAGCTTAAGGAGTACCAGATC
>hsa-miR-4316
GGTGAGGCTAGCTGGTG
>hsa-miR-644a
AGTGTGGCTTTCTTAGAGC
>hsa-miR-515-5p
TTCTCCAAAAGAAAGCACTTTCTG
>hsa-miR-1207-3p
TCAGCTGGCCCTCATTTC
>hsa-miR-374c-3p
CACTTAGCAGGTTGTATTATAT
>hsa-miR-4707-3p
AGCCCGCCCCAGCCGAGGTTCT
>hsa-miR-4436b-3p
CAGGGCAGGAAGAAGTGGACAA
>hsa-miR-6762-3p
TGGCTGCTTCCCTTGGTCTCCAG
>hsa-miR-4323
CAGCCCCACAGCCTCAGA
>hsa-miR-8073
ACCTGGCAGCAGGGAGCGTCGT
>hsa-miR-2113
ATTTGTGCTTGGCTCTGTCAC
>hsa-miR-4501
TATGTGACCTCGGATGAATCA
>hsa-miR-181a-5p

AACATTCAACGCTGTCGGTGAGT
>hsa-miR-1245a
AAGTGATCTAAAGGCCTACAT
>hsa-miR-323b-5p
AGGTTGTCCGTGGTGAGTTCGCA
>hsa-miR-3150a-3p
CTGGGGAGATCCTCGAGGTTGG
>hsa-miR-548f-3p
AAAAACTGTAATTACTTTT
>hsa-miR-6825-5p
TGGGGAGGTGTGGAGTCAGCAT
>hsa-miR-1255a
AGGATGAGCAAAGAAAGTAGATT
>hsa-miR-127-5p
CTGAAGCTCAGAGGGCTCTGAT
>hsa-miR-4655-3p
ACCCTCGTCAGGTCCCCGGGG
>hsa-miR-371b-5p
ACTCAAAAGATGGCGGCACTTT
>hsa-miR-1912-3p
TACCCAGAGCATGCAGTGTGAA
>hsa-miR-302a-3p
TAAGTGCTTCCATGTTTTGGTGA
>hsa-miR-34a-3p
CAATCAGCAAGTATACTGCCCT
>hsa-miR-3126-5p
TGAGGGACAGATGCCAGAAGCA
>hsa-miR-3943
TAGCCCCCAGGCTTCACTTGGCG
>hsa-miR-498-5p
TTTCAAGCCAGGGGGCGTTTTTC
>hsa-miR-548ba
AAAGGTAACTGTGATTTTTGCT
>hsa-miR-6880-3p
CCGCCTTCTCTCCTCCCCCAG
>hsa-miR-6786-3p
TGACGCCCCTTCTGATTCTGCCT
>hsa-miR-4492
GGGGCTGGGCGCGCGCC
>hsa-miR-570-3p
CGAAAACAGCAATTACCTTTGC
>hsa-miR-4766-5p
TCTGAAAGAGCAGTTGGTGTT
>hsa-miR-6861-3p
TGGACCTCTCCTCCCCAG
>hsa-miR-6076
AGCATGACAGAGGAGAGGTGG
>hsa-miR-7847-3p
CGTGGAGGACGAGGAGGAGGC
>hsa-miR-6845-3p
CCTCTCCTCCCTGTGCCCCAG
>hsa-miR-4418
CACTGCAGGACTCAGCAG
>hsa-miR-3655
GCTTGTCGCTGCGGTGTTGCT
>hsa-miR-6841-5p
TAGGGTACTCAGAGCAAGTTGT
>hsa-miR-518b
CAAAGCGCTCCCCTTTAGAGGT
>hsa-miR-3936
TAAGGGGTGTATGGCAGATGCA
>hsa-miR-582-3p

TAACTGGTTGAACAACTGAACC
>hsa-miR-380-5p
TGGTTGACCATAGAACATGCGC
>hsa-miR-4677-5p
TTGTTCTTTGGTCTTTCAGCCA
>hsa-miR-890
TACTTGGAAAGGCATCAGTTG
>hsa-miR-5188
AATCGGACCCATTTAAACCGGAG
>hsa-miR-4788
TTACGGACCAGCTAAGGGAGGC
>hsa-miR-6513-3p
TCAAGTGTCATCTGTCCCTAG
>hsa-miR-6883-3p
TTCCCTATCTCACTCTCCTCAG
>hsa-miR-3163
TATAAAATGAGGGCAGTAAGAC
>hsa-miR-432-5p
TCTTGGAGTAGGTCATTGGGTGG
>hsa-miR-6892-3p
TCCCTCTCCCACCCCTTGCAG
>hsa-miR-7112-5p
ACGGGCAGGGCAGTGCACCCTG
>hsa-miR-7160-3p
CAGGGCCCTGGCTTTAGCAGA
>hsa-miR-2909
GTTAGGGCCAACATCTCTTGG
>hsa-miR-409-5p
AGGTTACCCGAGCAACTTTGCAT
>hsa-miR-5092
AATCCACGCTGAGCTTGGCATC
>hsa-miR-3161
CTGATAAGAACAGAGGCCCAGAT
>hsa-miR-548aj-3p
TAAAAACTGCAATTACTTTTA
>hsa-miR-1269a
CTGGACTGAGCCGTGCTACTGG
>hsa-miR-12129
GATGTACTGAACTGGGTCAGAC
>hsa-miR-584-5p
TTATGGTTTGCCTGGGACTGAG
>hsa-miR-4503
TTTAAGCAGGAAATAGAATTTA
>hsa-miR-597-5p
TGTGTCACTCGATGACCACTGT
>hsa-miR-4654
TGTGGGATCTGGAGGCATCTGG
>hsa-miR-345-3p
GCCCTGAACGAGGGGTCTGGAG
>hsa-miR-6790-5p
GTGAGTGTGGATTTGGCGGGGTT
>hsa-miR-6779-3p
AAGCCCTGTCTCCTCCCATCT
>hsa-miR-3186-5p
CAGGCGTCTGTCTACGTGGCTT
>hsa-miR-216b-5p
AAATCTCTGCAGGCAAATGTGA
>hsa-miR-200b-3p
TAATACTGCCTGGTAATGATGA
>hsa-miR-6872-5p
TCTCGCATCAGGAGGCAAGG
>hsa-miR-6818-5p

```
TTGTGTGAGTACAGAGAGCATC
>hsa-miR-19a-5p
AGTTTTGCATAGTTGCACTACA
>hsa-miR-6885-3p
CTTTGCTTCCTGCTCCCCTAG
>hsa-miR-4480
AGCCAAGTGGAAGTTACTTTA
>hsa-miR-3909
TGTCCTCTAGGGCCTGCAGTCT
>hsa-miR-582-5p
TTACAGTTGTTCAACCAGTTACT
>hsa-miR-345-5p
GCTGACTCCTAGTCCAGGGCTC
>hsa-miR-6802-3p
TTCACCCCTCTCACCTAAGCAG
>hsa-miR-4748
GAGGTTTGGGGAGGATTTGCT
>hsa-miR-4795-5p
AGAAGTGGCTAATAATATTGA
>hsa-miR-6774-5p
ACTTGGGCAGGAGGGACCCTGTATG
>hsa-miR-3910
AAAGGCATAAAACCAAGACA
>hsa-miR-1273c
GGCGACAAAACGAGACCCTGTC
>hsa-miR-3117-5p
AGACACTATACGAGTCATAT
>hsa-miR-629-5p
TGGGTTTACGTTGGGAGAACT
>hsa-miR-509-5p
TACTGCAGACAGTGGCAATCA
>hsa-miR-5691
TTGCTCTGAGCTCCGAGAAAGC
>hsa-miR-4422
AAAAGCATCAGGAAGTACCCA
>hsa-miR-4289
GCATTGTGCAGGGCTATCA
>hsa-miR-668-3p
TGTCACTCGGCTCGGCCCACTAC
>hsa-miR-3150b-5p
CAACCTCGAGGATCTCCCCAGC
>hsa-miR-508-5p
TACTCCAGAGGGCGTCACTCATG
>hsa-miR-4453
GAGCTTGGTCTGTAGCGGTT
>hsa-miR-130b-5p
ACTCTTTCCCTGTTGCACTAC
>hsa-miR-329-3p
AACACACCTGGTTAACCTCTTT
>hsa-miR-4663
AGCTGAGCTCCATGGACGTGCAGT
>hsa-miR-5705
TGTTTCGGGGCTCATGGCCTGTG
>hsa-miR-548w
AAAAGTAACTGCGGTTTTTGCCT
>hsa-miR-3691-3p
ACCAAGTCTGCGTCATCCTCTC
>hsa-miR-301a-3p
CAGTGCAATAGTATTGTCAAAGC
>hsa-miR-4768-3p
CCAGGAGATCCAGAGAGAAT
>hsa-miR-548aq-5p
```

GAAAGTAATTGCTGTTTTTGCC
>hsa-miR-6529-5p
GAGAGATCAGAGGCGCAGAGTG
>hsa-miR-4309
CTGGAGTCTAGGATTCCA
>hsa-miR-2115-3p
CATCAGAATTCATGGAGGCTAG
>hsa-miR-513b-5p
TTCACAAGGAGGTGTCATTTAT
>hsa-miR-7111-5p
TGGGGGAGGAAGGACAGGCCAT
>hsa-let-7f-5p
TGAGGTAGTAGATTGTATAGTT
>hsa-miR-4656
TGGGCTGAGGGCAGGAGGCCTGT
>hsa-miR-610
TGAGCTAAATGTGTGCTGGGA
>hsa-miR-4659a-3p
TTTCTTCTTAGACATGGCAACG
>hsa-miR-3918
ACAGGGCCGCAGATGGAGACT
>hsa-miR-149-3p
AGGGAGGGACGGGGGCTGTGC
>hsa-miR-191-5p
CAACGGAATCCCAAAAGCAGCTG
>hsa-miR-3606-5p
TTAGTGAAGGCTATTTTAATT
>hsa-miR-3973
ACAAAGTACAGCATTAGCCTTAG
>hsa-miR-378a-5p
CTCCTGACTCCAGGTCCTGTGT
>hsa-miR-4732-3p
GCCCTGACCTGTCCTGTTCTG
>hsa-miR-488-3p
TTGAAAGGCTATTTCTTGGTC
>hsa-miR-6808-3p
GTGTGACCACCGTTCCTGCAG
>hsa-miR-548y
AAAAGTAATCACTGTTTTTGCC
>hsa-miR-500a-3p
ATGCACCTGGGCAAGGATTCTG
>hsa-miR-141-3p
TAACACTGTCTGGTAAAGATGG
>hsa-miR-3151-3p
CCTGATCCCACAGCCCACCT
>hsa-miR-137-5p
ACGGGTATTCTTGGGTGGATAAT
>hsa-miR-4319
TCCCTGAGCAAAGCCAC
>hsa-miR-1291
TGGCCCTGACTGAAGACCAGCAGT
>hsa-miR-186-3p
GCCCAAAGGTGAATTTTTTGGG
>hsa-miR-12115
TAGTGGAGCTGGGAGGCAGCTCGG
>hsa-miR-215-5p
ATGACCTATGAATTGACAGAC
>hsa-miR-301b-5p
GCTCTGACGAGGTTGCACTACT
>hsa-miR-6511b-5p
CTGCAGGCAGAAGTGGGGCTGACA
>hsa-miR-6870-3p

GCTCATCCCCATCTCCTTTCAG
>hsa-miR-4723-3p
CCCTCTCTGGCTCCTCCCCAAA
>hsa-miR-507
TTTTGCACCTTTTGGAGTGAA
>hsa-miR-5683
TACAGATGCAGATTCTCTGACTTC
>hsa-miR-1229-5p
GTGGGTAGGGTTTGGGGGAGAGCG
>hsa-miR-6799-5p
GGGGAGGTGTGCAGGGCTGG
>hsa-miR-5681a
AGAAAGGGTGGCAATACCTCTT
>hsa-miR-1183
CACTGTAGGTGATGGTGAGAGTGGGCA
>hsa-miR-4420
GTCACTGATGTCTGTAGCTGAG
>hsa-miR-3670
AGAGCTCACAGCTGTCCTTCTCTA
>hsa-miR-629-3p
GTTCTCCCAACGTAAGCCCAGC
>hsa-miR-6070
CCGGTTCCAGTCCCTGGAG
>hsa-miR-502-3p
AATGCACCTGGGCAAGGATTCA
>hsa-miR-208a-5p
GAGCTTTTGGCCCGGGTTATAC
>hsa-miR-216a-3p
TCACAGTGGTCTCTGGGATTAT
>hsa-miR-520b-3p
AAAGTGCTTCCTTTTAGAGGG
>hsa-miR-10401-3p
ACCTCGCCGTCCCGCCCGCCG
>hsa-miR-6829-3p
TGCCTCCTCCGTGGCCTCAG
>hsa-miR-548am-3p
CAAAAACTGCAGTTACTTTTGT
>hsa-miR-6746-5p
CCGGGAGAAGGAGGTGGCCTGG
>hsa-miR-3940-3p
CAGCCCGGATCCCAGCCCACTT
>hsa-miR-217-5p
TACTGCATCAGGAACTGATTGGA
>hsa-miR-224-3p
AAAATGGTGCCCTAGTGACTACA
>hsa-miR-1468-5p
CTCCGTTTGCCTGTTTCGCTG
>hsa-miR-642b-5p
GGTTCCCTCTCCAAATGTGTCT
>hsa-miR-3935
TGTAGATACGAGCACCAGCCAC
>hsa-miR-429
TAATACTGTCTGGTAAAACCGT
>hsa-miR-518f-3p
GAAAGCGCTTCTCTTTAGAGG
>hsa-miR-6781-5p
CGGGCCGGAGGTCAAGGGCGT
>hsa-miR-6744-3p
GGGCCTCTCTTGTCATCCTGCAG
>hsa-miR-4733-5p
AATCCCAATGCTAGACCCGGTG
>hsa-miR-3913-5p

TTTGGGACTGATCTTGATGTCT
>hsa-miR-598-5p
GCGGTGATCCCGATGGTGTGAGC
>hsa-miR-7850-5p
GTTTGGACATAGTGTGGCTGG
>hsa-miR-888-3p
GACTGACACCTCTTTGGGTGAA
>hsa-miR-4490
TCTGGTAAGAGATTTGGGCATA
>hsa-miR-4264
ACTCAGTCATGGTCATT
>hsa-miR-99b-5p
CACCCGTAGAACCGACCTTGCG
>hsa-miR-4540
TTAGTCCTGCCTGTAGGTTTA
>hsa-miR-372-5p
CCTCAAATGTGGAGCACTATTCT
>hsa-miR-6760-5p
CAGGGAGAAGGTGGAAGTGCAGA
>hsa-miR-937-3p
ATCCGCGCTCTGACTCTCTGCC
>hsa-miR-1324
CCAGACAGAATTCTATGCACTTTC
>hsa-miR-7-2-3p
CAACAAATCCCAGTCTACCTAA
>hsa-miR-502-5p
ATCCTTGCTATCTGGGTGCTA
>hsa-miR-3134
TGATGGATAAAAGACTACATATT
>hsa-miR-524-5p
CTACAAAGGGAAGCACTTTCTC
>hsa-miR-1294
TGTGAGGTTGGCATTGTTGTCT
>hsa-miR-6763-3p
CTCCCCGGCCTCTGCCCCCAG
>hsa-miR-4695-5p
CAGGAGGCAGTGGGCGAGCAGG
>hsa-miR-3942-3p
TTTCAGATAACAGTATTACAT
>hsa-miR-126-5p
CATTATTACTTTTGGTACGCG
>hsa-miR-5583-3p
GAATATGGGTATATTAGTTTGG
>hsa-miR-101-3p
TACAGTACTGTGATAACTGAA
>hsa-miR-5680
GAGAAATGCTGGACTAATCTGC
>hsa-miR-4714-3p
CCAACCTAGGTGGTCAGAGTTG
>hsa-miR-4785
AGAGTCGGCGACGCCGCCAGC
>hsa-miR-378f
ACTGGACTTGGAGCCAGAAG
>hsa-miR-8052
CGGGACTGTAGAGGGCATGAGC
>hsa-miR-6751-3p
ACTGAGCCTCTCTCTCTCCAG
>hsa-miR-937-5p
GTGAGTCAGGGTGGGGCTGG
>hsa-miR-3692-5p
CCTGCTGGTCAGGAGTGGATACTG
>hsa-miR-6814-3p

ACTCGCATCCTTCCCTTGGCAG
>hsa-miR-3145-5p
AACTCCAAACACTCAAAACTCA
>hsa-miR-10396a-5p
GGCGGGGCTCGGAGCCGGG
>hsa-miR-6876-5p
CAGGAAGGAGACAGGCAGTTCA
>hsa-miR-3668
AATGTAGAGATTGATCAAAAT
>hsa-miR-3651
CATAGCCCGGTCGCTGGTACATGA
>hsa-miR-1281
TCGCCTCCTCCTCTCCC
>hsa-miR-6805-5p
TAGGGGGCGGCTTGTGGAGTGT
>hsa-miR-4766-3p
ATAGCAATTGCTCTTTTGGAA
>hsa-miR-6737-3p
TCTGTGCTTCACCCCTACCCAG
>hsa-miR-3129-5p
GCAGTAGTGTAGAGATTGGTTT
>hsa-miR-8083
CAGGACTTGACGGCTGCAACT
>hsa-miR-1283
TCTACAAAGGAAAGCGCTTTCT
>hsa-miR-3618
TGTCTACATTAATGAAAAGAGC
>hsa-miR-498-3p
AAAGCACCTCCAGAGCTTGAAGC
>hsa-miR-8071
CGGTGGACTGGAGTGGGTGG
>hsa-miR-129-1-3p
AAGCCCTTACCCCAAAAAGTAT
>hsa-miR-4671-5p
ACCGAAGACTGTGCGCTAATCT
>hsa-miR-4740-5p
AGGACTGATCCTCTCGGGCAGG
>hsa-miR-6845-5p
CGGGGCCAGAGCAGAGAGC
>hsa-miR-8063
TCAAAATCAGGAGTCGGGGCTT
>hsa-miR-3913-3p
AGACATCAAGATCAGTCCCAAA
>hsa-miR-185-5p
TGGAGAGAAAGGCAGTTCCTGA
>hsa-miR-630
AGTATTCTGTACCAGGGAAGGT
>hsa-let-7e-3p
CTATACGGCCTCCTAGCTTTCC
>hsa-miR-3154
CAGAAGGGGAGTTGGGAGCAGA
>hsa-miR-325
CCTAGTAGGTGTCCAGTAAGTGT
>hsa-miR-4793-3p
TCTGCACTGTGAGTTGGCTGGCT
>hsa-miR-5089-3p
ATGCTACTCGGAAATCCCACTGA
>hsa-miR-4517
AAATATGATGAAACTCACAGCTGAG
>hsa-miR-6743-5p
AAGGGGCAGGGACGGGTGGCCC
>hsa-miR-4659b-5p

TTGCCATGTCTAAGAAGAA
>hsa-miR-3977
GTGCTTCATCGTAATTAACCTTA
>hsa-miR-664b-5p
TGGGCTAAGGGAGATGATTGGGTA
>hsa-miR-4530
CCCAGCAGGACGGGAGCG
>hsa-miR-3622a-5p
CAGGCACGGGAGCTCAGGTGAG
>hsa-miR-6071
TTCTGCTGCCGGCCAAGGC
>hsa-miR-1307-3p
ACTCGGCGTGGCGTCGGTCGTG
>hsa-miR-433-5p
TACGGTGAGCCTGTCATTATTC
>hsa-miR-4261
AGGAAACAGGGACCCA
>hsa-miR-1321
CAGGGAGGTGAATGTGAT
>hsa-miR-556-5p
GATGAGCTCATTGTAATATGAG
>hsa-miR-3187-5p
CCTGGGCAGCGTGTGGCTGAAGG
>hsa-miR-5008-3p
CCTGTGCTCCCAGGGCCTCGC
>hsa-miR-4724-5p
AACTGAACCAGGAGTGAGCTTCG
>hsa-miR-328-5p
GGGGGGGCAGGAGGGGCTCAGGG
>hsa-miR-4727-5p
ATCTGCCAGCTTCCACAGTGG
>hsa-miR-4636
AACTCGTGTTCAAAGCCTTTAG
>hsa-miR-548ar-5p
AAAAGTAATTGCAGTTTTTGC
>hsa-miR-1245b-3p
TCAGATGATCTAAAGGCCTATA
>hsa-miR-422a
ACTGGACTTAGGGTCAGAAGGC
>hsa-miR-19b-1-5p
AGTTTTGCAGGTTTGCATCCAGC
>hsa-miR-6795-5p
TGGGGGGACAGGATGAGAGGCTGT
>hsa-miR-196b-3p
TCGACAGCACGACACTGCCTTC
>hsa-miR-1261
ATGGATAAGGCTTTGGCTT
>hsa-miR-206
TGGAATGTAAGGAAGTGTGTGG
>hsa-miR-1260a
ATCCCACCTCTGCCACCA
>hsa-miR-4284
GGGCTCACATCACCCCAT
>hsa-miR-130a-3p
CAGTGCAATGTTAAAAGGGCAT
>hsa-miR-6805-3p
TTGCTCTGCTCCCCCGCCCCCAG
>hsa-miR-3613-5p
TGTTGTACTTTTTTTTTTGTTC
>hsa-miR-4317
ACATTGCCAGGGAGTTT
>hsa-miR-378g

ACTGGGCTTGGAGTCAGAAG
>hsa-miR-92a-3p
TATTGCACTTGTCCCGGCCTGT
>hsa-miR-603
CACACACTGCAATTACTTTTGC
>hsa-miR-6820-3p
TGTGACTTCTCCCCTGCCACAG
>hsa-miR-3529-5p
AGGTAGACTGGGATTTGTTGTT
>hsa-miR-6719-3p
TCTGACATCAGTGATTCTCCTG
>hsa-miR-4747-3p
AAGGCCCGGGCTTTCCTCCCAG
>hsa-miR-4711-5p
TGCATCAGGCCAGAAGACATGAG
>hsa-miR-4703-3p
TGTAGTTGTATTGTATTGCCAC
>hsa-miR-6887-5p
TGGGGGGACAGATGGAGAGGACA
>hsa-miR-6735-5p
CAGGGCAGAGGGCACAGGAATCTGA
>hsa-miR-5193
TCCTCCTCTACCTCATCCCAGT
>hsa-miR-4715-5p
AAGTTGGCTGCAGTTAAGGTGG
>hsa-miR-3921
TCTCTGAGTACCATATGCCTTGT
>hsa-miR-1245b-5p
TAGGCCTTTAGATCACTTAAA
>hsa-miR-218-5p
TTGTGCTTGATCTAACCATGT
>hsa-miR-3155b
CCAGGCTCTGCAGTGGGA
>hsa-miR-181b-3p
CTCACTGAACAATGAATGCAA
>hsa-miR-219a-1-3p
AGAGTTGAGTCTGGACGTCCCG
>hsa-miR-891b
TGCAACTTACCTGAGTCATTGA
>hsa-miR-7977
TTCCCAGCCAACGCACCA
>hsa-miR-7107-5p
TCGGCCTGGGGAGGAGGAAGGG
>hsa-miR-6873-3p
TTCTCTCTGTCTTTCTCTCTCAG
>hsa-miR-4798-3p
AACTCACGAAGTATACCGAAGT
>hsa-miR-767-5p
TGCACCATGGTTGTCTGAGCATG
>hsa-miR-12130
CGGGTTGTACCTTTTTTGC
>hsa-miR-5004-5p
TGAGGACAGGGCAAATTCACGA
>hsa-miR-3679-3p
CTTCCCCCCAGTAATCTTCATC
>hsa-miR-561-5p
ATCAAGGATCTTAAACTTTGCC
>hsa-miR-187-3p
TCGTGTCTTGTGTTGCAGCCGG
>hsa-miR-4649-3p
TCTGAGGCCTGCCTCTCCCCA
>hsa-miR-7151-5p

GATCCATCTCTGCCTGTATTGGC
>hsa-miR-3923
AACTAGTAATGTTGGATTAGGG
>hsa-miR-548h-5p
AAAAGTAATCGCGGTTTTTGTC
>hsa-miR-7974
AGGCTGTGATGCTCTCCTGAGCCC
>hsa-miR-135a-3p
TATAGGGATTGGAGCCGTGGCG
>hsa-miR-11399
TTCAGGTCTGGGGCTGAAACCT
>hsa-miR-6788-5p
CTGGGAGAAGAGTGGTGAAGA
>hsa-miR-3168
GAGTTCTACAGTCAGAC
>hsa-miR-6796-5p
TTGTGGGGTTGGAGAGCTGGCTG
>hsa-miR-411-5p
TAGTAGACCGTATAGCGTACG
>hsa-miR-222-3p
AGCTACATCTGGCTACTGGGT
>hsa-miR-6818-3p
TTGTCTCTTGTTCCTCACACAG
>hsa-miR-3678-5p
TCCGTACAAACTCTGCTGTG
>hsa-miR-129-5p
CTTTTTGCGGTCTGGGCTTGC
>hsa-miR-616-5p
ACTCAAAACCCTTCAGTGACTT
>hsa-miR-95-5p
TCAATAAATGTCTGTTGAATT
>hsa-miR-4665-5p
CTGGGGGACGCGTGAGCGCGAGC
>hsa-miR-652-5p
CAACCCTAGGAGAGGGTGCCATTCA
>hsa-miR-4730
CTGGCGGAGCCCATTCCATGCCA
>hsa-miR-3654
GACTGGACAAGCTGAGGAA
>hsa-miR-122-5p
TGGAGTGTGACAATGGTGTTTG
>hsa-miR-4508
GCGGGGCTGGGCGCGCG
>hsa-miR-7705
AATAGCTCAGAATGTCAGTTCTG
>hsa-miR-6793-5p
TGTGGGTTCTGGGTTGGGGTGA
>hsa-miR-6510-3p
CACCGACTCTGTCTCCTGCAG
>hsa-miR-2114-3p
CGAGCCTCAAGCAAGGGACTT
>hsa-miR-885-5p
TCCATTACACTACCCTGCCTCT
>hsa-miR-620
ATGGAGATAGATATAGAAAT
>hsa-miR-320a-3p
AAAAGCTGGGTTGAGAGGGCGA
>hsa-miR-6501-5p
AGTTGCCAGGGCTGCCTTTGGT
>hsa-miR-23a-3p
ATCACATTGCCAGGGATTTCC
>hsa-miR-1227-3p

CGTGCCACCCTTTTCCCCAG
>hsa-miR-181a-3p
ACCATCGACCGTTGATTGTACC
>hsa-miR-4659b-3p
TTTCTTCTTAGACATGGCAGCT
>hsa-miR-3680-5p
GACTCACTCACAGGATTGTGCA
>hsa-miR-4538
GAGCTTGGATGAGCTGGGCTGA
>hsa-miR-3175
CGGGGAGAGAACGCAGTGACGT
>hsa-miR-15a-3p
CAGGCCATATTGTGCTGCCTCA
>hsa-miR-421
ATCAACAGACATTAATTGGGCGC
>hsa-miR-655-5p
AGAGGTTATCCGTGTTATGTTC
>hsa-miR-1244
AAGTAGTTGGTTTGTATGAGATGGTT
>hsa-miR-1914-5p
CCCTGTGCCCGGCCCACTTCTG
>hsa-miR-181a-2-3p
ACCACTGACCGTTGACTGTACC
>hsa-miR-448
TTGCATATGTAGGATGTCCCAT
>hsa-miR-4731-3p
CACACAAGTGGCCCCCAACACT
>hsa-miR-1306-5p
CCACCTCCCCTGCAAACGTCCA
>hsa-miR-6770-5p
TGAGAAGGCACAGCTTGCACGTGA
>hsa-miR-4307
AATGTTTTTTCCTGTTTCC
>hsa-miR-4428
CAAGGAGACGGGAACATGGAGC
>hsa-miR-1825
TCCAGTGCCCTCCTCTCC
>hsa-miR-4443
TTGGAGGCGTGGGTTTT
>hsa-miR-6890-3p
CCACTGCCTATGCCCCACAG
>hsa-miR-96-3p
AATCATGTGCAGTGCCAATATG
>hsa-miR-4653-3p
TGGAGTTAAGGGTTGCTTGGAGA
>hsa-miR-155-5p
TTAATGCTAATCGTGATAGGGGTT
>hsa-miR-6894-5p
AGGAGGATGGAGAGCTGGGCCAGA
>hsa-miR-302c-3p
TAAGTGCTTCCATGTTTCAGTGG
>hsa-miR-24-1-5p
TGCCTACTGAGCTGATATCAGT
>hsa-miR-4303
TTCTGAGCTGAGGACAG
>hsa-miR-548m
CAAAGGTATTTGTGGTTTTTG
>hsa-miR-6791-3p
TGCCTCCTTGGTCTCCGGCAG
>hsa-miR-4524a-3p
TGAGACAGGCTTATGCTGCTAT
>hsa-miR-4999-3p

TCACTACCTGACAATACAGT
>hsa-miR-6747-5p
AGGGGTGTGGAAAGAGGCAGAACA
>hsa-miR-150-5p
TCTCCCAACCCTTGTACCAGTG
>hsa-miR-6856-5p
AAGAGAGGAGCAGTGGTGCTGTGG
>hsa-miR-4477a
CTATTAAGGACATTTGTGATTC
>hsa-miR-454-3p
TAGTGCAATATTGCTTATAGGGT
>hsa-miR-6749-5p
TCGGGCCTGGGGTTGGGGGAGC
>hsa-miR-3133
TAAAGAACTCTTAAAACCCAAT
>hsa-miR-1301-5p
CGCTCTAGGCACCGCAGCA
>hsa-miR-9900
ACAGGTCCTAAGAGACTGCAT
>hsa-miR-4695-3p
TGATCTCACCGCTGCCTCCTTC
>hsa-miR-4714-5p
AACTCTGACCCCTTAGGTTGAT
>hsa-miR-8075
TGCTGATGGCAGATGTCGGGTCTG
>hsa-miR-4756-3p
CCAGAGATGGTTGCCTTCCTAT
>hsa-miR-4472
GGTGGGGGGTGTTGTTTT
>hsa-miR-1253
AGAGAAGAAGATCAGCCTGCA
>hsa-miR-6790-3p
CGACCTCGGCGACCCCTCACT
>hsa-miR-657
GGCAGGTTCTCACCCTCTCTAGG
>hsa-miR-3183
GCCTCTCTCGGAGTCGCTCGGA
>hsa-miR-202-3p
AGAGGTATAGGGCATGGGAA
>hsa-miR-939-5p
TGGGGAGCTGAGGCTCTGGGGGTG
>hsa-miR-93-3p
ACTGCTGAGCTAGCACTTCCCG
>hsa-let-7c-5p
TGAGGTAGTAGGTTGTATGGTT
>hsa-miR-4479
CGCGCGGCCGTGCTCGGAGCAG
>hsa-miR-604
AGGCTGCGGAATTCAGGAC
>hsa-miR-548az-5p
CAAAAGTGATTGTGGTTTTTGC
>hsa-miR-3652
CGGCTGGAGGTGTGAGGA
>hsa-miR-4719
TCACAAATCTATAATATGCAGG
>hsa-miR-7109-3p
CAAGCCTCTCCTGCCCTTCCAG
>hsa-miR-101-2-5p
TCGGTTATCATGGTACCGATGC
>hsa-miR-6769b-3p
CCCTCTCTGTCCCACCCATAG
>hsa-miR-200b-5p

CATCTTACTGGGCAGCATTGGA
>hsa-miR-302d-5p
ACTTTAACATGGAGGCACTTGC
>hsa-miR-3683
TGCGACATTGGAAGTAGTATCA
>hsa-miR-676-5p
TCTTCAACCTCAGGACTTGCA
>hsa-miR-198
GGTCCAGAGGGGAGATAGGTTC
>hsa-miR-6869-3p
CGCCGCGCGCATCGGCTCAGC
>hsa-miR-4664-5p
TGGGGTGCCCACTCCGCAAGTT
>hsa-miR-4638-5p
ACTCGGCTGCGGTGGACAAGT
>hsa-miR-548a-5p
AAAAGTAATTGCGAGTTTTACC
>hsa-miR-4513
AGACTGACGGCTGGAGGCCCAT
>hsa-miR-3191-5p
CTCTCTGGCCGTCTACCTTCCA
>hsa-miR-4717-5p
TAGGCCACAGCCACCCATGTGT
>hsa-miR-510-5p
TACTCAGGAGAGTGGCAATCAC
>hsa-miR-548an
AAAAGGCATTGTGGTTTTTG
>hsa-miR-3131
TCGAGGACTGGTGGAAGGGCCTT
>hsa-miR-331-5p
CTAGGTATGGTCCCAGGGATCC
>hsa-miR-4761-3p
GAGGGCATGCGCACTTTGTCC
>hsa-miR-128-3p
TCACAGTGAACCGGTCTCTTT
>hsa-miR-3657
TGTGTCCCATTATTGGTGATT
>hsa-miR-548t-3p|hsa-miR-548aa
AAAAACCACAATTACTTTTGCACCA
>hsa-miR-6772-5p
TGGGTGTAGGCTGGAGCTGAGG
>hsa-miR-188-3p
CTCCCACATGCAGGGTTTGCA
>hsa-miR-4644
TGGAGAGAGAAAAGAGACAGAAG
>hsa-miR-32-3p
CAATTTAGTGTGTGTGATATTT
>hsa-miR-6736-3p
TCAGCTCCTCTCTACCCACAG
>hsa-miR-3138
TGTGGACAGTGAGGTAGAGGGAGT
>hsa-miR-3120-3p
CACAGCAAGTGTAGACAGGCA
>hsa-miR-6732-3p
TAACCCTGTCCTCTCCCTCCCAG
>hsa-miR-6864-5p
TTGAAGGGACAAGTCAGATATGCC
>hsa-miR-520a-5p
CTCCAGAGGGAAGTACTTTCT
>hsa-miR-5011-3p
GTGCATGGCTGTATATATAACA
>hsa-miR-1227-5p

GTGGGGCCAGGCGGTGG
>hsa-miR-412-3p
ACTTCACCTGGTCCACTAGCCGT
>hsa-miR-101-5p
CAGTTATCACAGTGCTGATGCT
>hsa-miR-605-5p
TAAATCCCATGGTGCCTTCTCCT
>hsa-miR-6813-5p
CAGGGGCTGGGGTTTCAGGTTCT
>hsa-miR-3615
TCTCTCGGCTCCTCGCGGCTC
>hsa-miR-8067
CCTAGAAACTGTAAACTTAGTC
>hsa-miR-548v
AGCTACAGTTACTTTTGCACCA
>hsa-miR-21-3p
CAACACCAGTCGATGGGCTGT
>hsa-miR-7106-3p
AGCTCCCTGAATCCCTGTCCCAG
>hsa-miR-4477b
ATTAAGGACATTTGTGATTGAT
>hsa-miR-5690
TCAGCTACTACCTCTATTAGG
>hsa-miR-576-5p
ATTCTAATTTCTCCACGTCTTT
>hsa-miR-4257
CCAGAGGTGGGGACTGAG
>hsa-miR-3140-5p
ACCTGAATTACCAAAAGCTTT
>hsa-miR-4790-5p
ATCGCTTTACCATTCATGTT
>hsa-miR-548ao-3p
AAAGACCGTGACTACTTTTGCA
>hsa-miR-6793-3p
TCCCCAACCCCTGCCCGCAG
>hsa-miR-6128
ACTGGAATTGGAGTCAAAA
>hsa-miR-5580-3p
CACATATGAAGTGAGCCAGCAC
>hsa-miR-574-5p
TGAGTGTGTGTGTGTGAGTGTGT
>hsa-miR-6502-3p
TAGACCATCTTTCTAGAGTAT
>hsa-miR-6833-5p
GTGTGGAAGATGGGAGGAGAAA
>hsa-miR-6716-5p
TGGGAATGGGGGTAAGGGCC
>hsa-miR-204-3p
GCTGGGAAGGCAAAGGGACGT
>hsa-miR-6797-3p
TGCATGACCCTTCCCTCCCCAC
>hsa-miR-4706
AGCGGGGAGGAAGTGGGCGCTGCTT
>hsa-miR-578
CTTCTTGTGCTCTAGGATTGT
>hsa-miR-6814-5p
TCCCAAGGGTGAGATGCTGCCA
>hsa-miR-4775
TTAATTTTTTGTTTCGGTCACT
>hsa-miR-4710
GGGTGAGGGCAGGTGGTT
>hsa-miR-942-5p

TCTTCTCTGTTTTGGCCATGTG
>hsa-miR-548a-3p
CAAAACTGGCAATTACTTTTGC
>hsa-miR-516a-5p
TTCTCGAGGAAAGAAGCACTTTC
>hsa-miR-4801
TACACAAGAAAACCAAGGCTCA
>hsa-miR-654-3p
TATGTCTGCTGACCATCACCTT
>hsa-miR-299-5p
TGGTTTACCGTCCCACATACAT
>hsa-miR-1197
TAGGACACATGGTCTACTTCT
>hsa-miR-34c-5p
AGGCAGTGTAGTTAGCTGATTGC
>hsa-miR-5706
TTCTGGATAACATGCTGAAGCT
>hsa-miR-663b
GGTGGCCCGGCCGTGCCTGAGG
>hsa-miR-185-3p
AGGGGCTGGCTTTCCTCTGGTC
>hsa-miR-147b-3p
GTGTGCGGAAATGCTTCTGCT
>hsa-miR-5693
GCAGTGGCTCTGAAATGAACTC
>hsa-miR-142-5p
CATAAAGTAGAAAGCACTACT
>hsa-miR-5001-3p
TTCTGCCTCTGTCCAGGTCCTT
>hsa-miR-4449
CGTCCCGGGGCTGCGCGAGGCA
>hsa-miR-378a-3p
ACTGGACTTGGAGTCAGAAGGC
>hsa-miR-4727-3p
ATAGTGGGAAGCTGGCAGATTC
>hsa-miR-4436a
GCAGGACAGGCAGAAGTGGAT
>hsa-miR-145-3p
GGATTCCTGGAAATACTGTTCT
>hsa-miR-4772-5p
TGATCAGGCAAAATTGCAGACT
>hsa-miR-3942-5p
AAGCAATACTGTTACCTGAAAT
>hsa-miR-28-5p
AAGGAGCTCACAGTCTATTGAG
>hsa-miR-4504
TGTGACAATAGAGATGAACATG
>hsa-miR-150-3p
CTGGTACAGGCCTGGGGGACAG
>hsa-miR-8057
GTGGCTCTGTAGTAAGATGGA
>hsa-miR-4524b-5p
ATAGCAGCATAAGCCTGTCTC
>hsa-miR-3200-3p
CACCTTGCGCTACTCAGGTCTG
>hsa-miR-485-3p
GTCATACACGGCTCTCCTCTCT
>hsa-miR-4306
TGGAGAGAAAGGCAGTA
>hsa-miR-3185
AGAAGAAGGCGGTCGGTCTGCGG
>hsa-miR-4725-5p

```
AGACCCTGCAGCCTTCCCACC
>hsa-miR-302f
TAATTGCTTCCATGTTT
>hsa-miR-6817-5p
TCTGCCATAGGAAGCTTGGAGTGG
>hsa-miR-3124-3p
ACTTTCCTCACTCCCGTGAAGT
>hsa-miR-6858-5p
GTGAGGAGGGGCTGGCAGGGAC
>hsa-miR-128-1-5p
CGGGGCCGTAGCACTGTCTGAGA
>hsa-miR-4502
GCTGATGATGATGGTGCTGAAG
>hsa-miR-590-3p
TAATTTTATGTATAAGCTAGT
>hsa-miR-6857-5p
TTGGGGATTGGGTCAGGCCAGT
>hsa-miR-4445-3p
CACGGCAAAAGAAACAATCCA
>hsa-miR-6865-3p
ACACCCTCTTTCCCTACCGCC
>hsa-miR-141-5p
CATCTTCCAGTACAGTGTTGGA
>hsa-miR-7110-5p
TGGGGGTGTGGGGAGAGAGAG
>hsa-miR-4500
TGAGGTAGTAGTTTCTT
>hsa-miR-10524-5p
CAGGATGCCAGCATAGT
>hsa-miR-4311
GAAAGAGAGCTGAGTGTG
>hsa-miR-517a-3p|hsa-miR-517b-3p
ATCGTGCATCCCTTTAGAGTGT
>hsa-miR-3116
TGCCTGGAACATAGTAGGGACT
>hsa-miR-4999-5p
TGCTGTATTGTCAGGTAGTGA
>hsa-miR-20b-5p
CAAAGTGCTCATAGTGCAGGTAG
>hsa-miR-4481
GGAGTGGGCTGGTGGTT
>hsa-miR-769-3p
CTGGGATCTCCGGGGTCTTGGTT
>hsa-miR-5191
AGGATAGGAAGAATGAAGTGCT
>hsa-miR-12117
GAAGTGGAGCACATCAGTGA
>hsa-miR-6812-3p
CCGCTCTTCCCCTGACCCCAG
>hsa-miR-3689b-3p|hsa-miR-3689c
CTGGGAGGTGTGATATTGTGGT
>hsa-miR-4305
CCTAGACACCTCCAGTTC
>hsa-miR-548o-3p
CCAAAACTGCAGTTACTTTTGC
>hsa-miR-625-3p
GACTATAGAACTTTCCCCCTCA
>hsa-miR-6516-3p
ATCATGTATGATACTGCAAACA
>hsa-miR-641
AAAGACATAGGATAGAGTCACCTC
>hsa-miR-6511a-5p
```

CAGGCAGAAGTGGGGCTGACAGG
>hsa-miR-466
ATACACATACACGCAACACACAT
>hsa-miR-1304-3p
TCTCACTGTAGCCTCGAACCCC
>hsa-miR-497-3p
CAAACCACACTGTGGTGTTAGA
>hsa-miR-627-5p
GTGAGTCTCTAAGAAAAGAGGA
>hsa-miR-6748-3p
TCCTGTCCCTGTCTCCTACAG
>hsa-miR-451a
AAACCGTTACCATTACTGAGTT
>hsa-miR-651-5p
TTTAGGATAAGCTTGACTTTTG
>hsa-miR-374a-5p
TTATAATACAACCTGATAAGTG
>hsa-miR-3978
GTGGAAAGCATGCATCCAGGGTGT
>hsa-miR-517-5p
CCTCTAGATGGAAGCACTGTCT
>hsa-miR-555
AGGGTAAGCTGAACCTCTGAT
>hsa-miR-1827
TGAGGCAGTAGATTGAAT
>hsa-miR-153-5p
TCATTTTTGTGATGTTGCAGCT
>hsa-miR-6882-5p
TACAAGTCAGGAGCTGAAGCAG
>hsa-miR-6083
CTTATATCAGAGGCTGTGGG
>hsa-miR-3194-3p
AGCTCTGCTGCTCACTGGCAGT
>hsa-miR-6741-5p
GTGGGTGCTGGTGGGAGCCGTG
>hsa-miR-3130-5p
TACCCAGTCTCCGGTGCAGCC
>hsa-miR-1205
TCTGCAGGGTTTGCTTTGAG
>hsa-miR-133a-3p
TTTGGTCCCCTTCAACCAGCTG
>hsa-miR-6722-3p
TGCAGGGGTCGGGTGGGCCAGG
>hsa-miR-6085
AAGGGGCTGGGGGAGCACA
>hsa-miR-4469
GCTCCCTCTAGGGTCGCTCGGA
>hsa-miR-8061
CTTAGATTAGAGGATATTGTT
>hsa-miR-6732-5p
TAGGGGGTGGCAGGCTGGCC
>hsa-miR-4685-3p
TCTCCCTTCCTGCCCTGGCTAG
>hsa-miR-223-5p
CGTGTATTTGACAAGCTGAGTT
>hsa-miR-221-5p
ACCTGGCATACAATGTAGATTT
>hsa-miR-623
ATCCCTTGCAGGGGCTGTTGGGT
>hsa-miR-615-5p
GGGGGTCCCCGGTGCTCGGATC
>hsa-miR-4747-5p

AGGGAAGGAGGCTTGGTCTTAG
>hsa-miR-877-5p
GTAGAGGAGATGGCGCAGGG
>hsa-miR-6787-5p
TGGCGGGGGTAGAGCTGGCTGC
>hsa-miR-6835-3p
AAAAGCACTTTTCTGTCTCCCAG
>hsa-miR-4774-3p
ATTGCCTAACATGTGCCAGAA
>hsa-miR-3678-3p
CTGCAGAGTTTGTACGGACCGG
>hsa-miR-6878-5p
AGGGAGAAAGCTAGAAGCTGAAG
>hsa-miR-365b-5p
AGGGACTTTCAGGGGCAGCTGT
>hsa-miR-6757-3p
AACACTGGCCTTGCTATCCCCA
>hsa-miR-4535
GTGGACCTGGCTGGGAC
>hsa-miR-379-5p
TGGTAGACTATGGAACGTAGG
>hsa-let-7a-2-3p
CTGTACAGCCTCCTAGCTTTCC
>hsa-miR-767-3p
TCTGCTCATACCCCATGGTTTCT
>hsa-miR-6893-5p
CAGGCAGGTGTAGGGTGGAGC
>hsa-miR-4528
TCATTATATGTATGATCTGGAC
>hsa-miR-4253
AGGGCATGTCCAGGGGGT
>hsa-miR-520f-5p
CCTCTAAAGGGAAGCGCTTTCT
>hsa-miR-6866-3p
GATCCCTTTATCTGTCCTCTAG
>hsa-miR-885-3p
AGGCAGCGGGGTGTAGTGGATA
>hsa-miR-4712-3p
AATGAGAGACCTGTACTGTAT
>hsa-miR-7843-3p
ATGAAGCCTTCTCTGCCTTACG
>hsa-miR-3136-5p
CTGACTGAATAGGTAGGGTCATT
>hsa-miR-4446-3p
CAGGGCTGGCAGTGACATGGGT
>hsa-miR-6754-3p
TCTTCACCTGCCTCTGCCTGCA
>hsa-miR-5190
CCAGTGACTGAGCTGGAGCCA
>hsa-miR-6505-5p
TTGGAATAGGGGATATCTCAGC
>hsa-miR-6075
ACGGCCCAGGCGGCATTGGTG
>hsa-miR-4745-5p
TGAGTGGGGCTCCCGGGACGGCG
>hsa-miR-4662a-3p
AAAGATAGACAATTGGCTAAAT
>hsa-miR-378d
ACTGGACTTGGAGTCAGAAA
>hsa-miR-576-3p
AAGATGTGGAAAAATTGGAATC
>hsa-miR-593-3p

TGTCTCTGCTGGGGTTTCT
>hsa-miR-542-5p
TCGGGGATCATCATGTCACGAGA
>hsa-miR-374b-5p
ATATAATACAACCTGCTAAGTG
>hsa-miR-4521
GCTAAGGAAGTCCTGTGCTCAG
>hsa-miR-6787-3p
TCTCAGCTGCTGCCCTCTCCAG
>hsa-miR-12127
TAGAGGTTCTAATGGTCCAGGGTTA
>hsa-miR-4518
GCTCAGGGATGATAACTGTGCTGAGA
>hsa-miR-653-3p
TTCACTGGAGTTTGTTTCAATA
>hsa-miR-509-3p
TGATTGGTACGTCTGTGGGTAG
>hsa-miR-1206
TGTTCATGTAGATGTTTAAGC
>hsa-miR-7845-5p
AAGGGACAGGGAGGGTCGTGG
>hsa-miR-4634
CGGCGCGACCGGCCCGGGG
>hsa-miR-4451
TGGTAGAGCTGAGGACA
>hsa-miR-6734-5p
TTGAGGGGAGAATGAGGTGGAGA
>hsa-miR-12132
TATTACTGTGAGAATTATGATG
>hsa-miR-6792-5p
GTAAGCAGGGGCTCTGGGTGA
>hsa-miR-6868-5p
ACTGGCAGAACACTGAAGCAGC
>hsa-miR-6821-5p
GTGCGTGGTGGCTCGAGGCGGGG
>hsa-miR-146a-3p
CCTCTGAAATTCAGTTCTTCAG
>hsa-miR-1268a
CGGGCGTGGTGGTGGGGG
>hsa-let-7i-5p
TGAGGTAGTAGTTTGTGCTGTT
>hsa-miR-4430
AGGCTGGAGTGAGCGGAG
>hsa-miR-3157-3p
CTGCCCTAGTCTAGCTGAAGCT
>hsa-miR-511-5p
GTGTCTTTTGCTCTGCAGTCA
>hsa-miR-6866-5p
TTAGAGGCTGGAATAGAGATTCT
>hsa-miR-3907
AGGTGCTCCAGGCTGGCTCACA
>hsa-miR-29c-3p
TAGCACCATTTGAAATCGGTTA
>hsa-miR-6746-3p
CAGCCGCCGCCTGTCTCCACAG
>hsa-miR-609
AGGGTGTTTCTCTCATCTCT
>hsa-miR-193b-5p
CGGGGTTTTGAGGGCGAGATGA
>hsa-miR-2682-3p
CGCCTCTTCAGCGCTGTCTTCC
>hsa-miR-3925-3p

ACTCCAGTTTTAGTTCTCTTG
>hsa-miR-508-3p
TGATTGTAGCCTTTTGGAGTAGA
>hsa-miR-4705
TCAATCACTTGGTAATTGCTGT
>hsa-miR-6775-3p
AGGCCCTGTCCTCTGCCCCAG
>hsa-miR-4794
TCTGGCTATCTCACGAGACTGT
>hsa-miR-4681
AACGGGAATGCAGGCTGTATCT
>hsa-miR-4761-5p
ACAAGGTGTGCATGCCTGACC
>hsa-miR-520d-5p
CTACAAAGGGAAGCCCTTTC
>hsa-miR-140-3p
TACCACAGGGTAGAACCACGG
>hsa-miR-4716-5p
TCCATGTTTCCTTCCCCCTTCT
>hsa-miR-15a-5p
TAGCAGCACATAATGGTTTGTG
>hsa-miR-4722-5p
GGCAGGAGGGCTGTGCCAGGTTG
>hsa-miR-7852-3p
TATGTAGTAGTCAAAGGCATTT
>hsa-miR-5587-5p
ATGGTCACCTCCGGGACT
>hsa-miR-6782-5p
TAGGGGTGGGGGAATTCAGGGGTGT
>hsa-miR-5572
GTTGGGGTGCAGGGGTCTGCT
>hsa-miR-4486
GCTGGGCGAGGCTGGCA
>hsa-miR-3689f
TGTGATATCGTGCTTCCTGGGA
>hsa-miR-10399-5p
AATTACAGATTGTCTCAGAGA
>hsa-miR-4749-3p
CGCCCCTCCTGCCCCCACAG
>hsa-miR-6731-3p
TCTATTCCCCACTCTCCCCAG
>hsa-miR-606
AAACTACTGAAAATCAAAGAT
>hsa-miR-549a-3p
TGACAACTATGGATGAGCTCT
>hsa-miR-3085-3p
TCTGGCTGCTATGGCCCCCTC
>hsa-miR-186-5p
CAAAGAATTCTCCTTTTGGGCT
>hsa-miR-649
AAACCTGTGTTGTTCAAGAGTC
>hsa-miR-8068
TGTTTGTTGTAAGGATCGTTGT
>hsa-miR-1278
TAGTACTGTGCATATCATCTAT
>hsa-miR-3691-5p
AGTGGATGATGGAGACTCGGTAC
>hsa-miR-4487
AGAGCTGGCTGAAGGGCAG
>hsa-miR-4462
TGACACGGAGGGTGGCTTGGGAA
>hsa-miR-4776-3p

CTTGCCATCCTGGTCCACTGCAT
>hsa-miR-522-3p
AAAATGGTTCCCTTTAGAGTGT
>hsa-miR-1915-5p
ACCTTGCCTTGCTGCCCGGGCC
>hsa-miR-764
GCAGGTGCTCACTTGTCCTCCT
>hsa-miR-548as-3p
TAAAACCCACAATTATGTTTGT
>hsa-miR-455-5p
TATGTGCCTTTGGACTACATCG
>hsa-miR-5682
GTAGCACCTTGCAGGATAAGGT
>hsa-miR-107
AGCAGCATTGTACAGGGCTATCA
>hsa-miR-4523
GACCGAGAGGGCCTCGGCTGT
>hsa-miR-4452
TTGAATTCTTGGCCTTAAGTGAT
>hsa-miR-5707
ACGTTTGAATGCTGTACAAGGC
>hsa-miR-520f-3p
AAGTGCTTCCTTTTAGAGGGTT
>hsa-miR-940
AAGGCAGGGCCCCCGCTCCCC
>hsa-miR-5588-3p
AAGTCCCACTAATGCCAGC
>hsa-miR-219b-3p
AGAATTGCGTTTGGACAATCAGT
>hsa-miR-6828-3p
ATCTGCTCTCTTGTTCCCAG
>hsa-miR-12116
TTAGGCTTCCCCCTCCTCCTGC
>hsa-miR-7706
TGAAGCGCCTGTGCTCTGCCGAGA
>hsa-miR-3922-5p
TCAAGGCCAGAGGTCCCACAGCA
>hsa-miR-597-3p
TGGTTCTCTTGTGGCTCAAGCGT
>hsa-miR-1292-3p
TCGCGCCCCGGCTCCCGTTC
>hsa-miR-924
AGAGTCTTGTGATGTCTTGC
>hsa-miR-4270
TCAGGGAGTCAGGGGAGGGC
>hsa-miR-545-5p
TCAGTAAATGTTTATTAGATGA
>hsa-let-7d-3p
CTATACGACCTGCTGCCTTTCT
>hsa-miR-4675
GGGGCTGTGATTGACCAGCAGG
>hsa-miR-873-3p
GGAGACTGATGAGTTCCCGGGA
>hsa-miR-377-5p
AGAGGTTGCCCTTGGTGAATTC
>hsa-miR-6716-3p
TCCGAACTCTCCATTCCTCTGC
>hsa-miR-6773-5p
TTGGGCCCAGGAGTAAACAGGAT
>hsa-miR-490-5p
CCATGGATCTCCAGGTGGGT
>hsa-miR-7158-3p

CTGAACTAGAGATTGGGCCCA
>hsa-miR-519e-3p
AAGTGCCTCCTTTTAGAGTGTT
>hsa-miR-146a-5p
TGAGAACTGAATTCCATGGGTT
>hsa-miR-573
CTGAAGTGATGTGTAACTGATCAG
>hsa-miR-1290
TGGATTTTTGGATCAGGGA
>hsa-miR-1258
AGTTAGGATTAGGTCGTGGAA
>hsa-miR-548b-3p
CAAGAACCTCAGTTGCTTTTGT
>hsa-miR-6499-3p
AGCAGTGTTTGTTTTGCCCACA
>hsa-miR-518a-5p|hsa-miR-527
CTGCAAAGGGAAGCCCTTTC
>hsa-miR-10525-3p
TGACTATGATGTGCACCTGAT
>hsa-miR-378i
ACTGGACTAGGAGTCAGAAGG
>hsa-miR-6894-3p
TTGCCTGCCCTCTTCCTCCAG
>hsa-miR-1303
TTTAGAGACGGGGTCTTGCTCT
>hsa-miR-122-3p
AACGCCATTATCACACTAAATA
>hsa-miR-132-5p
ACCGTGGCTTTCGATTGTTACT
>hsa-miR-7113-3p
CCTCCCTGCCCGCCTCTCTGCAG
>hsa-miR-5703
AGGAGAAGTCGGGAAGGT
>hsa-miR-3920
ACTGATTATCTTAACTCTCTGA
>hsa-miR-514a-3p
ATTGACACTTCTGTGAGTAGA
>hsa-miR-6847-5p
ACAGAGGACAGTGGAGTGTGAGC
>hsa-miR-4498
TGGGCTGGCAGGGCAAGTGCTG
>hsa-miR-654-5p
TGGTGGGCCGCAGAACATGTGC
>hsa-miR-1256
AGGCATTGACTTCTCACTAGCT
>hsa-miR-3162-3p
TCCCTACCCCTCCACTCCCCA
>hsa-miR-4280
GAGTGTAGTTCTGAGCAGAGC
>hsa-miR-548b-5p
AAAAGTAATTGTGGTTTTGGCC
>hsa-miR-8053
TGGCGATTTTGGAACTCAATGGCA
>hsa-miR-5581-3p
TTCCATGCCTCCTAGAAGTTCC
>hsa-miR-4505
AGGCTGGGCTGGGACGGA
>hsa-miR-6755-3p
TGTTGTCATGTTTTTTCCCTAG
>hsa-miR-6727-5p
CTCGGGGCAGGCGGCTGGGAGCG
>hsa-miR-4495

AATGTAAACAGGCTTTTTGCT
>hsa-miR-5591-3p
ATACCCATAGCTTAGCTCCCA
>hsa-miR-5588-5p
ACTGGCATTAGTGGGACTTTT
>hsa-miR-4436b-5p
GTCCACTTCTGCCTGCCCTGCC
>hsa-miR-5001-5p
AGGGCTGGACTCAGCGGCGGAGCT
>hsa-miR-5197-3p
AAGAAGAGACTGAGTCATCGAAT
>hsa-miR-26a-2-3p
CCTATTCTTGATTACTTGTTTC
>hsa-miR-18a-3p
ACTGCCCTAAGTGCTCCTTCTGG
>hsa-miR-3174
TAGTGAGTTAGAGATGCAGAGCC
>hsa-miR-10a-5p
TACCCTGTAGATCCGAATTTGTG
>hsa-miR-139-3p
TGGAGACGCGGCCCTGTTGGAGT
>hsa-miR-212-5p
ACCTTGGCTCTAGACTGCTTACT
>hsa-miR-5699-5p
TGCCCCAACAAGGAAGGACAAG
>hsa-miR-8084
GAATACTAAGTAAAAAATCAGTA
>hsa-miR-18b-5p
TAAGGTGCATCTAGTGCAGTTAG
>hsa-miR-4735-3p
AAAGGTGCTCAAATTAGACAT
>hsa-miR-450a-5p
TTTTGCGATGTGTTCCTAATAT
>hsa-miR-4728-3p
CATGCTGACCTCCCTCCTGCCCCAG
>hsa-miR-2682-5p
CAGGCAGTGACTGTTCAGACGTC
>hsa-miR-6134
TGAGGTGGTAGGATGTAGA
>hsa-miR-4313
AGCCCCCTGGCCCCAAACCC
>hsa-miR-29b-1-5p
GCTGGTTTCATATGGTGGTTTAGA
>hsa-miR-30d-5p
TGTAAACATCCCCGACTGGAAG
>hsa-miR-6720-3p
CGCGCCTGCAGGAACTGGTAGA
>hsa-miR-3671
ATCAAATAAGGACTAGTCTGCA
>hsa-miR-5008-5p
TGAGGCCCTTGGGGCACAGTGG
>hsa-miR-6849-3p
ACCAGCCTGTGTCCACCTCCAG
>hsa-miR-760
CGGCTCTGGGTCTGTGGGGA
>hsa-miR-143-5p
GGTGCAGTGCTGCATCTCTGGT
>hsa-miR-3908
GAGCAATGTAGGTAGACTGTTT
>hsa-miR-6825-3p
GCGCTGACCCGCCTTCTCCGCA
>hsa-miR-219a-2-3p

AGAATTGTGGCTGGACATCTGT
>hsa-miR-6798-3p
CTACCCCCCATCCCCCTGTAG
>hsa-miR-4690-5p
GAGCAGGCGAGGCTGGGCTGAA
>hsa-miR-148b-5p
AAGTTCTGTTATACACTCAGGC
>hsa-miR-1180-5p
GGACCCACCCGGCCGGGAATA
>hsa-miR-135a-5p
TATGGCTTTTTATTCCTATGTGA
>hsa-miR-196a-3p
CGGCAACAAGAAACTGCCTGAG
>hsa-miR-1469
CTCGGCGCGGGGCGCGGGCTCC
>hsa-miR-6853-3p
TGTTCATTGGAACCCTGCGCAG
>hsa-miR-3660
ACTGACAGGAGAGCATTTTGA
>hsa-miR-5187-3p
ACTGAATCCTCTTTTCCTCAG
>hsa-miR-6779-5p
CTGGGAGGGGCTGGGTTTGGC
>hsa-miR-4536-5p
TGTGGTAGATATATGCACGAT
>hsa-miR-4255
CAGTGTTCAGAGATGGA
>hsa-miR-6776-5p
TCTGGGTGCAGTGGGGGTT
>hsa-miR-8076
TATATGGACTTTTCTGATACAATG
>hsa-miR-6857-3p
TGACTGAGCTTCTCCCCACAG
>hsa-miR-34b-3p
CAATCACTAACTCCACTGCCAT
>hsa-miR-635
ACTTGGGCACTGAAACAATGTCC
>hsa-miR-651-3p
AAAGGAAAGTGTATCCTAAAAG
>hsa-miR-12126
GACTTGGGGACCAGACCTTTTCTT
>hsa-miR-548ap-3p
AAAAACCACAATTACTTTT
>hsa-miR-6854-3p
TGCGTTTCTCCTCTTGAGCAG
>hsa-miR-4781-3p
AATGTTGGAATCCTCGCTAGAG
>hsa-miR-562
AAAGTAGCTGTACCATTTGC
>hsa-miR-6507-5p
GAAGAATAGGAGGGACTTTGT
>hsa-miR-6786-5p
GCGGTGGGGCCGGAGGGGCGT
>hsa-miR-744-3p
CTGTTGCCACTAACCTCAACCT
>hsa-miR-4670-3p
TGAAGTTACATCATGGTCGCTT
>hsa-miR-106b-5p
TAAAGTGCTGACAGTGCAGAT
>hsa-miR-6813-3p
AACCTTGGCCCCTCTCCCCAG
>hsa-miR-3925-5p

AAGAGAACTGAAAGTGGAGCCT
>hsa-miR-323a-5p
AGGTGGTCCGTGGCGCGTTCGC
>hsa-miR-6724-5p
CTGGGCCCGCGGCGGGCGTGGGG
>hsa-miR-151b
TCGAGGAGCTCACAGTCT
>hsa-miR-4640-5p
TGGGCCAGGGAGCAGCTGGTGGG
>hsa-miR-12121
CTGCCACGAGCGTGCGGGCCT
>hsa-miR-6785-5p
TGGGAGGGCGTGGATGATGGTG
>hsa-miR-4786-3p
TGAAGCCAGCTCTGGTCTGGGC
>hsa-miR-7111-3p
ATCCTCTCTTCCCTCCTCCCAG
>hsa-miR-6131
GGCTGGTCAGATGGGAGTG
>hsa-miR-4655-5p
CACCGGGGATGGCAGAGGGTCG
>hsa-miR-4651
CGGGGTGGGTGAGGTCGGGC
>hsa-miR-4666b
TTGCATGTCAGATTGTAATTCCC
>hsa-miR-933
TGTGCGCAGGGAGACCTCTCCC
>hsa-miR-7162-3p
TCTGAGGTGGAACAGCAGC
>hsa-miR-6504-3p
CATTACAGCACAGCCATTCT
>hsa-miR-9983-3p
TTTTTTGCTGGAACATTTCTGG
>hsa-miR-4295
CAGTGCAATGTTTTCCTT
>hsa-miR-6780a-3p
CTCCTCTGTTTTCTTTCCTAG
>hsa-miR-8070
ATGTGATTGACGGCTGACTCCA
>hsa-miR-99a-5p
AACCCGTAGATCCGATCTTGTG
>hsa-miR-4755-3p
AGCCAGGCTCTGAAGGGAAAGT
>hsa-miR-143-3p
TGAGATGAAGCACTGTAGCTC
>hsa-miR-486-3p
CGGGGCAGCTCAGTACAGGAT
>hsa-miR-891a-3p
AGTGGCACATGTTTGTTGTGAG
>hsa-miR-6843-3p
ATGGTCTCCTGTTCTCTGCAG
>hsa-miR-4275
CCAATTACCACTTCTTT
>hsa-miR-548h-3p|hsa-miR-548z
CAAAAACCGCAATTACTTTTGCA
>hsa-miR-300
TATACAAGGGCAGACTCTCTCT
>hsa-miR-329-5p
GAGGTTTTCTGGGTTTCTGTTTC
>hsa-miR-4709-5p
ACAACAGTGACTTGCTCTCCAA
>hsa-miR-377-3p

ATCACACAAAGGCAACTTTTGT
>hsa-miR-4696
TGCAAGACGGATACTGTCATCT
>hsa-miR-579-3p
TTCATTTGGTATAAACCGCGATT
>hsa-miR-762
GGGGCTGGGGCCGGGGCCGAGC
>hsa-miR-3189-5p
TGCCCCATCTGTGCCCTGGGTAGGA
>hsa-miR-600
ACTTACAGACAAGAGCCTTGCTC
>hsa-miR-3125
TAGAGGAAGCTGTGGAGAGA
>hsa-miR-12118
CAAGGAGGAGCGGGGATTAG
>hsa-miR-4423-5p
AGTTGCCTTTTTGTTCCCATGC
>hsa-miR-181b-2-3p
CTCACTGATCAATGAATGCA
>hsa-miR-6764-5p
TCCCAGGGTCTGGTCAGAGTTG
>hsa-miR-208a-3p
ATAAGACGAGCAAAAAGCTTGT
>hsa-miR-3975
TGAGGCTAATGCACTACTTCAC
>hsa-miR-7703
TTGCACTCTGGCCTTCTCCCAGG
>hsa-miR-3121-3p
TAAATAGAGTAGGCAAAGGACA
>hsa-miR-6733-5p
TGGGAAAGACAAACTCAGAGTT
>hsa-miR-487a-3p
AATCATACAGGGACATCCAGTT
>hsa-miR-99b-3p
CAAGCTCGTGTCTGTGGGTCCG
>hsa-miR-1277-3p
TACGTAGATATATATGTATTTT
>hsa-miR-6754-5p
CCAGGGAGGCTGGTTTGGAGGA
>hsa-miR-489-3p
GTGACATCACATATACGGCAGC
>hsa-miR-12136
GAAAAAGTCATGGAGGCC
>hsa-miR-599
GTTGTGTCAGTTTATCAAAC
>hsa-miR-1249-5p
AGGAGGGAGGAGATGGGCCAAGTT
>hsa-miR-4762-3p
CTTCTGATCAAGATTTGTGGTG
>hsa-miR-190b-5p
TGATATGTTTGATATTGGGTTG
>hsa-miR-2277-3p
TGACAGCGCCCTGCCTGGCTC
>hsa-miR-3617-5p
AAAGACATAGTTGCAAGATGGG
>hsa-miR-4677-3p
TCTGTGAGACCAAAGAACTACT
>hsa-miR-182-5p
TTTGGCAATGGTAGAACTCACACT
>hsa-miR-1298-5p
TTCATTCGGCTGTCCAGATGTA
>hsa-miR-125a-3p

ACAGGTGAGGTTCTTGGGAGCC
>hsa-miR-411-3p
TATGTAACACGGTCCACTAACC
>hsa-miR-1307-5p
TCGACCGGACCTCGACCGGCT
>hsa-miR-514b-3p
ATTGACACCTCTGTGAGTGGA
>hsa-miR-3186-3p
TCACGCGGAGAGATGGCTTTG
>hsa-miR-4266
CTAGGAGGCCTTGGCC
>hsa-miR-6861-5p
ACTGGGTAGGTGGGGCTCCAGG
>hsa-miR-6770-3p
CTGGCGGCTGTGTCTTCACAG
>hsa-miR-205-3p
GATTTCAGTGGAGTGAAGTTC
>hsa-miR-6738-3p
CTTCTGCCTGCATTCTACTCCCAG
>hsa-miR-33a-5p
GTGCATTGTAGTTGCATTGCA
>hsa-miR-203b-3p
TTGAACTGTTAAGAACCACTGGA
>hsa-miR-561-3p
CAAAGTTTAAGATCCTTGAAGT
>hsa-miR-3675-3p
CATCTCTAAGGAACTCCCCCAA
>hsa-miR-200a-5p
CATCTTACCGGACAGTGCTGGA
>hsa-miR-92b-3p
TATTGCACTCGTCCCGGCCTCC
>hsa-miR-4753-5p
CAAGGCCAAAGGAAGAGAACAG
>hsa-miR-7155-3p
TGGCCCAAGACCTCAGACC
>hsa-miR-6777-5p
ACGGGGAGTCAGGCAGTGGTGGA
>hsa-miR-3150a-5p
CAACCTCGACGATCTCCTCAGC
>hsa-miR-5698
TGGGGGAGTGCAGTGATTGTGG
>hsa-miR-495-5p
GAAGTTGCCCATGTTATTTTCG
>hsa-miR-504-5p
AGACCCTGGTCTGCACTCTATC
>hsa-miR-6829-5p
TGGGCTGCTGAGAAGGGGCA
>hsa-miR-6811-3p
AGCCTGTGCTTGTCCCTGCAG
>hsa-miR-1305
TTTTCAACTCTAATGGGAGAGA
>hsa-miR-4448
GGCTCCTTGGTCTAGGGGTA
>hsa-miR-154-5p
TAGGTTATCCGTGTTGCCTTCG
>hsa-miR-376c-3p
AACATAGAGGAAATTCCACGT
>hsa-miR-5696
CTCATTTAAGTAGTCTGATGCC
>hsa-miR-199a-3p|hsa-miR-199b-3p
ACAGTAGTCTGCACATTGGTTA
>hsa-miR-574-3p

CACGCTCATGCACACACCCACA
>hsa-miR-7114-3p
TGACCCACCCCTCTCCACCAG
>hsa-miR-7108-5p
GTGTGGCCGGCAGGCGGGTGG
>hsa-miR-5586-5p
TATCCAGCTTGTTACTATATGC
>hsa-miR-6730-5p
AGAAAGGTGGAGGGGTTGTCAGA
>hsa-miR-134-5p
TGTGACTGGTTGACCAGAGGGG
>hsa-miR-875-3p
CCTGGAAACACTGAGGTTGTG
>hsa-miR-431-3p
CAGGTCGTCTTGCAGGGCTTCT
>hsa-miR-568
ATGTATAAATGTATACACAC
>hsa-miR-8062
CAGTGATTTGAGGATTATTGC
>hsa-miR-6133
TGAGGGAGGAGGTTGGGTA
>hsa-miR-510-3p
ATTGAAACCTCTAAGAGTGGA
>hsa-miR-3928-3p
GGAGGAACCTTGGAGCTTCGGC
>hsa-miR-6753-5p
CACCAGGGCAGAGCAGGGCTGA
>hsa-miR-133a-5p
AGCTGGTAAAATGGAACCAAAT
>hsa-miR-3160-5p
GGCTTTCTAGTCTCAGCTCTCC
>hsa-miR-4445-5p
AGATTGTTTCTTTTGCCGTGCA
>hsa-let-7f-2-3p
CTATACAGTCTACTGTCTTTCC
>hsa-miR-6847-3p
GGCTCATGTGTCTGTCCTCTTC
>hsa-miR-650
AGGAGGCAGCGCTCTCAGGAC
>hsa-miR-1539
TCCTGCGCGTCCCAGATGCCC
>hsa-miR-342-5p
AGGGGTGCTATCTGTGATTGA
>hsa-miR-532-3p
CCTCCCACACCCAAGGCTTGCA
>hsa-miR-921
CTAGTGAGGGACAGAACCAGGATTC
>hsa-miR-1288-3p
TGGACTGCCCTGATCTGGAGA
>hsa-miR-6757-5p
TAGGGATGGGAGGCCAGGATGA
>hsa-miR-320d
AAAAGCTGGGTTGAGAGGA
>hsa-miR-4684-5p
CTCTCTACTGACTTGCAACATA
>hsa-miR-4694-3p
CAAATGGACAGGATAACACCT
>hsa-miR-7156-5p
TTGTTCTCAAACTGGCTGTCAGA
>hsa-miR-552-3p
AACAGGTGACTGGTTAGACAA
>hsa-miR-6529-3p

CCTGTGCCTTTTACTTCTTTAA
>hsa-miR-6755-5p
TAGGGTAGACACTGACAACGTT
>hsa-miR-3074-5p
GTTCCTGCTGAACTGAGCCAG
>hsa-miR-7114-5p
TCTGTGGAGTGGGGTGCCTGT
>hsa-miR-6823-3p
TGAGCCTCTCCTTCCCTCCAG
>hsa-miR-4251
CCTGAGAAAAGGGCCAA
>hsa-miR-548az-3p
AAAAACTGCAATCACTTTTGC
>hsa-miR-4510
TGAGGGAGTAGGATGTATGGTT
>hsa-miR-542-3p
TGTGACAGATTGATAACTGAAA
>hsa-miR-670-5p
GTCCCTGAGTGTATGTGGTG
>hsa-miR-4738-5p
ACCAGCGCGTTTTCAGTTTCAT
>hsa-miR-181c-5p
AACATTCAACCTGTCGGTGAGT
>hsa-miR-4512
CAGGGCCTCACTGTATCGCCCA
>hsa-miR-548f-5p
TGCAAAAGTAATCACAGTTTTT
>hsa-miR-92a-2-5p
GGGTGGGGATTTGTTGCATTAC
>hsa-miR-193b-3p
AACTGGCCCTCAAAGTCCCGCT
>hsa-miR-4423-3p
ATAGGCACCAAAAAGCAACAA
>hsa-miR-6089
GGAGGCCGGGGTGGGGCGGGGCGG
>hsa-miR-4646-3p
ATTGTCCCTCTCCCTTCCCAG
>hsa-miR-3115
ATATGGGTTTACTAGTTGGT
>hsa-miR-3173-5p
TGCCCTGCCTGTTTTCTCCTTT
>hsa-miR-496
TGAGTATTACATGGCCAATCTC
>hsa-miR-4327
GGCTTGCATGGGGGACTGG
>hsa-miR-6836-3p
ATGCCTCCCCCGGCCCCGCAG
>hsa-miR-5701
TTATTGTCACGTTCTGATT
>hsa-miR-4524a-5p
ATAGCAGCATGAACCTGTCTCA
>hsa-miR-4328
CCAGTTTTCCCAGGATT
>hsa-miR-4680-5p
AGAACTCTTGCAGTCTTAGATGT
>hsa-miR-12114
CAGGTGGAGGTGTGAGGTC
>hsa-miR-2355-3p
ATTGTCCTTGCTGTTTGGAGAT
>hsa-miR-376a-5p
GTAGATTCTCCTTCTATGAGTA
>hsa-miR-888-5p

TACTCAAAAAGCTGTCAGTCA
>hsa-miR-1266-5p
CCTCAGGGCTGTAGAACAGGGCT
>hsa-miR-6801-3p
ACCCCTGCCACTCACTGGCC
>hsa-miR-134-3p
CCTGTGGGCCACCTAGTCACCAA
>hsa-miR-500b-5p
AATCCTTGCTACCTGGGT
>hsa-miR-889-5p
AATGGCTGTCCGTAGTATGGTC
>hsa-miR-570-5p|hsa-miR-548ai
AAAGGTAATTGCAGTTTTTCCC
>hsa-miR-7108-3p
ACCCGCCCGTCTCCCCACAG
>hsa-miR-1911-5p
TGAGTACCGCCATGTCTGTTGGG
>hsa-miR-4263
ATTCTAAGTGCCTTGGCC
>hsa-miR-6820-5p
TGCGGCAGAGCTGGGGTCA
>hsa-miR-3620-5p
GTGGGCTGGGCTGGGCTGGGCC
>hsa-miR-5003-5p
TCACAACAACCTTGCAGGGTAGA
>hsa-miR-126-3p
TCGTACCGTGAGTAATAATGCG
>hsa-miR-4797-5p
GACAGAGTGCCACTTACTGAA
>hsa-miR-6874-5p
ATGGAGCTGGAACCAGATCAGGC
>hsa-miR-637
ACTGGGGGCTTTCGGGCTCTGCGT
>hsa-miR-4260
CTTGGGGCATGGAGTCCCA
>hsa-miR-7978
TCTGGTGTATAGCGTTGCTCA
>hsa-miR-520d-3p
AAAGTGCTTCTCTTTGGTGGGT
>hsa-miR-4760-5p
TTTAGATTGAACATGAAGTTAG
>hsa-miR-195-3p
CCAATATTGGCTGTGCTGCTCC
>hsa-miR-4743-3p
TTTCTGTCTTTTCTGGTCCAG
>hsa-miR-6740-5p
AGTTTGGGATGGAGAGAGGAGA
>hsa-miR-4746-5p
CCGGTCCCAGGAGAACCTGCAGA
>hsa-miR-549a-5p
AGCTCATCCATAGTTGTCACTG
>hsa-miR-7854-3p
TGAGGTGACCGCAGATGGGAA
>hsa-miR-6775-5p
TCGGGGCATGGGGGAGGGAGGCTGG
>hsa-miR-4454
GGATCCGAGTCACGGCACCA
>hsa-miR-3937
ACAGGCGGCTGTAGCAATGGGGG
>hsa-miR-6831-5p
TAGGTAGAGTGTGAGGAGGAGGTC
>hsa-miR-191-3p

GCTGCGCTTGGATTTCGTCCCC
>hsa-miR-638
AGGGATCGCGGGCGGGTGGCGGCCT
>hsa-miR-18b-3p
TGCCCTAAATGCCCCTTCTGGC
>hsa-miR-626
AGCTGTCTGAAAATGTCTT
>hsa-miR-4789-5p
GTATACACCTGATATGTGTATG
>hsa-miR-6738-5p
CGAGGGGTAGAAGAGCACAGGGG
>hsa-miR-6852-3p
TGTCCTCTGTTCCTCAG
>hsa-miR-3155a
CCAGGCTCTGCAGTGGGAACT
>hsa-miR-424-5p
CAGCAGCAATTCATGTTTTGAA
>hsa-miR-363-3p
AATTGCACGGTATCCATCTGTA
>hsa-miR-491-5p
AGTGGGGAACCCTTCCATGAGG
>hsa-miR-96-5p
TTTGGCACTAGCACATTTTTGCT
>hsa-miR-452-3p
CTCATCTGCAAAGAAGTAAGTG
>hsa-miR-3157-5p
TTCAGCCAGGCTAGTGCAGTCT
>hsa-miR-6088
AGAGATGAAGCGGGGGGGCG
>hsa-miR-4632-5p
GAGGGCAGCGTGGGTGTGGCGGA
>hsa-miR-4527
TGGTCTGCAAAGAGATGACTGT
>hsa-miR-6763-5p
CTGGGGAGTGGCTGGGGAG
>hsa-miR-3664-3p
TCTCAGGAGTAAAGACAGAGTT
>hsa-miR-1976
CCTCCTGCCCTCCTTGCTGT
>hsa-miR-4690-3p
GCAGCCCAGCTGAGGCCTCTG
>hsa-miR-6869-5p
GTGAGTAGTGGCGCGCGGCGGC
>hsa-miR-765
TGGAGGAGAAGGAAGGTGATG
>hsa-miR-550a-3p
TGTCTTACTCCCTCAGGCACAT
>hsa-miR-4758-3p
TGCCCCACCTGCTGACCACCCTC
>hsa-miR-552-5p
GTTTAACCTTTTGCCTGTTGG
>hsa-miR-6824-3p
TCTCTGGTCTTGCCACCCCAG
>hsa-miR-106a-3p
CTGCAATGTAAGCACTTCTTAC
>hsa-miR-4722-3p
ACCTGCCAGCACCTCCCTGCAG
>hsa-miR-7973
TGTGACCCTAGAATAATTAC
>hsa-miR-5692a
CAAATAATACCACAGTGGGTGT
>hsa-miR-6850-3p

CCCGGCCGGAACGCCGCACT
>hsa-miR-4483
GGGGTGGTCTGTTGTTG
>hsa-miR-6081
AGGAGCAGTGCCGGCCAAGGCGCC
>hsa-miR-1288-5p
GCAGATCAGGACTGTAACTCACC
>hsa-miR-3159
TAGGATTACAAGTGTCGGCCAC
>hsa-miR-30a-5p
TGTAAACATCCTCGACTGGAAG
>hsa-miR-4692
TCAGGCAGTGTGGGTATCAGAT
>hsa-miR-136-3p
CATCATCGTCTCAAATGAGTCT
>hsa-miR-4444
CTCGAGTTGGAAGAGGCG
>hsa-miR-4482-3p
TTTCTATTTCTCAGTGGGGCTC
>hsa-miR-548as-5p
AAAAGTAATTGCGGGTTTTGCC
>hsa-miR-5687
TTAGAACGTTTTAGGGTCAAAT
>hsa-miR-4780
ACCCTTGAGCCTGATCCCTAGC
>hsa-miR-7702
CTTAGACTGCCAGACTCCCTGA
>hsa-miR-617
AGACTTCCCATTTGAAGGTGGC
>hsa-miR-337-5p
GAACGGCTTCATACAGGAGTT
>hsa-miR-571
TGAGTTGGCCATCTGAGTGAG
>hsa-miR-521
AACGCACTTCCCTTTAGAGTGT
>hsa-miR-145-5p
GTCCAGTTTTCCCAGGAATCCCT
>hsa-miR-6875-3p
ATTCTTCCTGCCCTGGCTCCAT
>hsa-miR-3184-5p
TGAGGGGCCTCAGACCGAGCTTTT
>hsa-miR-1267
CCTGTTGAAGTGTAATCCCCA
>hsa-miR-6846-5p
TGGGGGCTGGATGGGGTAGAGT
>hsa-miR-379-3p
TATGTAACATGGTCCACTAACT
>hsa-miR-4281
GGGTCCCGGGGAGGGGGG
>hsa-miR-524-3p
GAAGGCGCTTCCCTTTGGAGT
>hsa-miR-125b-1-3p
ACGGGTTAGGCTCTTGGGAGCT
>hsa-miR-548c-5p|hsa-miR-548o-5p|hsa-miR-548am-5p
AAAAGTAATTGCGGTTTTTGCC
>hsa-miR-10398-5p
TGGCTCCCTTCTCTCCGTCTG
>hsa-miR-5007-5p
TAGAGTCTGGCTGATATGGTTT
>hsa-miR-1266-3p
CCCTGTTCTATGCCCTGAGGGA
>hsa-miR-4765

TGAGTGATTGATAGCTATGTTC
>hsa-miR-484
TCAGGCTCAGTCCCCTCCCGAT
>hsa-miR-5571-5p
CAATTCTCAAAGGAGCCTCCC
>hsa-let-7i-3p
CTGCGCAAGCTACTGCCTTGCT
>hsa-miR-519a-3p
AAAGTGCATCCTTTTAGAGTGT
>hsa-miR-758-5p
GATGGTTGACCAGAGAGCACAC
>hsa-miR-4685-5p
CCCAGGGCTTGGAGTGGGGCAAGGTT
>hsa-miR-6505-3p
TGACTTCTACCTCTTCCAAAG
>hsa-miR-3916
AAGAGGAAGAAATGGCTGGTTCTCAG
>hsa-miR-4777-5p
TTCTAGATGAGAGATATATATA
>hsa-miR-6811-5p
ATGCAGGCCTGTGTACAGCACT
>hsa-miR-4514
ACAGGCAGGATTGGGGAA
>hsa-miR-4799-5p
ATCTAAATGCAGCATGCCAGTC
>hsa-miR-1277-5p
AAATATATATATATATGTACGTAT
>hsa-miR-4473
CTAGTGCTCTCCGTTACAAGTA
>hsa-miR-1236-5p
TGAGTGACAGGGGAAATGGGGA
>hsa-miR-4262
GACATTCAGACTACCTG
>hsa-miR-6878-3p
CTGGCCTCTTCTTTCTCCTAG
>hsa-miR-6778-5p
AGTGGGAGGACAGGAGGCAGGT
>hsa-miR-371a-5p
ACTCAAACTGTGGGGGCACT
>hsa-miR-378j
ACTGGATTTGGAGCCAGAA
>hsa-miR-7846-3p
CAGCGGAGCCTGGAGAGAAGG
>hsa-miR-7158-5p
GGCTCAATCTCTGGTCCTGCAGCC
>hsa-miR-6778-3p
TGCCTCCCTGACATTCCACAG
>hsa-miR-15b-5p
TAGCAGCACATCATGGTTTACA
>hsa-miR-7-1-3p
CAACAAATCACAGTCTGCCATA
>hsa-miR-500a-5p
TAATCCTTGCTACCTGGGTGAGA
>hsa-miR-4657
AATGTGGAAGTGGTCTGAGGCAT
>hsa-miR-3677-5p
CAGTGGCCAGAGCCCTGCAGTG
>hsa-miR-6810-5p
ATGGGGACAGGGATCAGCATGGC
>hsa-miR-6761-5p
TCTGAGAGAGCTCGATGGCAG
>hsa-miR-4290

TGCCCTCCTTTCTTCCCTC
>hsa-miR-3610
GAATCGGAAAGGAGGCGCCG
>hsa-miR-624-5p
TAGTACCAGTACCTTGTGTTCA
>hsa-miR-505-3p
CGTCAACACTTGCTGGTTTCCT
>hsa-miR-6846-3p
TGACCCCTTCTGTCTCCCTAG
>hsa-miR-4649-5p
TGGGCGAGGGGTGGGCTCTCAGAG
>hsa-miR-6862-3p
CCTCACCCAGCTCTCTGGCCCTCT
>hsa-miR-384
ATTCCTAGAAATTGTTCATA
>hsa-miR-3972
CTGCCAGCCCCGTTCCAGGGCA
>hsa-miR-4803
TAACATAATAGTGTGGATTGA
>hsa-miR-6780b-5p
TGGGGAAGGCTTGGCAGGGAAGA
>hsa-miR-3165
AGGTGGATGCAATGTGACCTCA
>hsa-miR-891a-5p
TGCAACGAACCTGAGCCACTGA
>hsa-miR-4633-5p
ATATGCCTGGCTAGCTCCTC
>hsa-miR-4755-5p
TTTCCCTTCAGAGCCTGGCTTT
>hsa-miR-223-3p
TGTCAGTTTGTCAAATACCCCA
>hsa-miR-4691-5p
GTCCTCCAGGCCATGAGCTGCGG
>hsa-miR-1298-3p
CATCTGGGCAACTGACTGAAC
>hsa-miR-5009-5p
TTGGACTTTTTCAGATTTGGGGAT
>hsa-miR-1246
AATGGATTTTTGGAGCAGG
>hsa-miR-518f-5p
CTCTAGAGGGAAGCACTTTCTC
>hsa-miR-3614-3p
TAGCCTTCAGATCTTGGTGTTTT
>hsa-miR-8079
CAGTGATCGTCTCTGCTGGC
>hsa-miR-6837-5p
ACCAGGGCCAGCAGGGAATGT
>hsa-miR-4666a-5p
ATACATGTCAGATTGTATGCC
>hsa-miR-3658
TTTAAGAAAACACCATGGAGAT
>hsa-miR-105-3p
ACGGATGTTTGAGCATGTGCTA
>hsa-miR-758-3p
TTTGTGACCTGGTCCACTAACC
>hsa-miR-6739-3p
ATTGTTCTGTCTTTCTCCCAG
>hsa-miR-3681-3p
ACACAGTGCTTCATCCACTACT
>hsa-miR-4299
GCTGGTGACATGAGAGGC
>hsa-miR-3681-5p

TAGTGGATGATGCACTCTGTGC
>hsa-miR-6884-5p
AGAGGCTGAGAAGGTGATGTTG
>hsa-miR-671-3p
TCCGGTTCTCAGGGCTCCACC
>hsa-miR-3663-5p
GCTGGTCTGCGTGGTGCTCGG
>hsa-miR-10394-5p
TCTGCAGGTCCTGGTGAACGCCAT
>hsa-miR-129-2-3p
AAGCCCTTACCCCAAAAAGCAT
>hsa-miR-2467-5p
TGAGGCTCTGTTAGCCTTGGCTC
>hsa-miR-487a-5p
GTGGTTATCCCTGCTGTGTTCG
>hsa-miR-4458
AGAGGTAGGTGTGGAAGAA
>hsa-miR-6074
GATATTCAGAGGCTAGGTGG
>hsa-miR-3153
GGGGAAAGCGAGTAGGGACATTT
>hsa-miR-449a
TGGCAGTGTATTGTTAGCTGGT
>hsa-miR-6073
GGTAGTGAGTTATCAGCTAC
>hsa-miR-3178
GGGGCGCGGCCGGATCG
>hsa-miR-4285
GCGGCGAGTCCGACTCAT
>hsa-miR-485-5p
AGAGGCTGGCCGTGATGAATTC
>hsa-miR-1304-5p
TTTGAGGCTACAGTGAGATGTG
>hsa-miR-3119
TGGCTTTTAACTTTGATGGC
>hsa-miR-6851-5p
AGGAGGTGGTACTAGGGGCCAGC
>hsa-miR-449b-5p
AGGCAGTGTATTGTTAGCTGGC
>hsa-miR-7976
TGCCCTGAGACTTTTGCTC
>hsa-miR-548av-3p
AAAACTGCAGTTACTTTTGC
>hsa-miR-31-3p
TGCTATGCCAACATATTGCCAT
>hsa-miR-5006-3p
TTTCCCTTTCCATCCTGGCAG
>hsa-miR-5695
ACTCCAAGAAGAATCTAGACAG
>hsa-miR-4725-3p
TGGGGAAGGCGTCAGTGTCGGG
>hsa-miR-619-5p
GCTGGGATTACAGGCATGAGCC
>hsa-miR-1910-5p
CCAGTCCTGTGCCTGCCGCCT
>hsa-miR-33a-3p
CAATGTTTCCACAGTGCATCAC
>hsa-miR-5091
ACGGAGACGACAAGACTGTGCTG
>hsa-miR-193a-5p
TGGGTCTTTGCGGGCGAGATGA
>hsa-miR-5000-3p

TCAGGACACTTCTGAACTTGGA
>hsa-miR-5689
AGCATACACCTGTAGTCCTAGA
>hsa-miR-1237-3p
TCCTTCTGCTCCGTCCCCCAG
>hsa-miR-187-5p
GGCTACAACACAGGACCCGGGC
>hsa-miR-4694-5p
AGGTGTTATCCTATCCATTTGC
>hsa-miR-1255b-2-3p
AACCACTTTCTTTGCTCATCCA
>hsa-miR-4524b-3p
GAGACAGGTTCATGCTGCTA
>hsa-miR-6817-3p
TCTCTCTGACTCCATGGCA
>hsa-miR-1468-3p
AGCAAAATAAGCAAATGGAAAA
>hsa-miR-451b
TAGCAAGAGAACCATTACCATT
>hsa-miR-1262
ATGGGTGAATTTGTAGAAGGAT
>hsa-miR-769-5p
TGAGACCTCTGGGTTCTGAGCT
>hsa-miR-4796-3p
TAAAGTGGCAGAGTATAGACAC
>hsa-miR-29b-2-5p
CTGGTTTCACATGGTGGCTTAG
>hsa-miR-371b-3p
AAGTGCCCCCACAGTTTGAGTGC
>hsa-miR-4779
TAGGAGGGAATAGTAAAAGCAG
>hsa-miR-6729-5p
TGGGCGAGGGCGGCTGAGCGGC
>hsa-miR-4698
TCAAAATGTAGAGGAAGACCCCA
>hsa-miR-3689d
GGGAGGTGTGATCTCACACTCG
>hsa-miR-4759
TAGGACTAGATGTTGGAATTA
>hsa-miR-367-3p
AATTGCACTTTAGCAATGGTGA
>hsa-miR-6090
GGGGAGCGAGGGGCGGGGC
>hsa-miR-124-5p
CGTGTTCACAGCGGACCTTGAT
>hsa-miR-320e
AAAGCTGGGTTGAGAAGG
>hsa-miR-27a-3p
TTCACAGTGGCTAAGTTCCGC
>hsa-miR-12122
TGTCACAGATGGCCGGAAGAGACTC
>hsa-miR-659-5p
AGGACCTTCCCTGAACCAAGGA
>hsa-miR-4650-3p
AGGTAGAATGAGGCCTGACAT
>hsa-miR-4790-3p
TGAATGGTAAAGCGATGTCACA
>hsa-miR-1296-5p
TTAGGGCCCTGGCTCCATCTCC
>hsa-miR-4712-5p
TCCAGTACAGGTCTCTCATTTC
>hsa-miR-873-5p

GCAGGAACTTGTGAGTCTCCT
>hsa-miR-548ad-5p|hsa-miR-548ae-5p
AAAAGTAATTGTGGTTTTTG
>hsa-miR-5692c
AATAATATCACAGTAGGTGTAC
>hsa-miR-9903
TTATCCTCCAGTAGACTAGGGA
>hsa-miR-935
CCAGTTACCGCTTCCGCTACCGC
>hsa-miR-4699-3p
AATTTACTCTGCAATCTTCTCC
>hsa-miR-5093
AGGAAATGAGGCTGGCTAGGAGC
>hsa-miR-1265
CAGGATGTGGTCAAGTGTTGTT
>hsa-miR-4265
CTGTGGGCTCAGCTCTGGG
>hsa-miR-4466
GGGTGCGGGCCGGCGGGG
>hsa-miR-10392-5p
GCGCTTCGACGGGCTGGGCTGTG
>hsa-miR-6506-5p
ACTGGGATGTCACTGAATATGGT
>hsa-miR-5700
TAATGCATTAAATTATTGAAGG
>hsa-miR-621
GGCTAGCAACAGCGCTTACCT
>hsa-miR-633
CTAATAGTATCTACCACAATAAA
>hsa-miR-4662b
AAAGATGGACAATTGGCTAAAT
>hsa-miR-519b-3p
AAAGTGCATCCTTTTAGAGGTT
>hsa-miR-665
ACCAGGAGGCTGAGGCCCCT
>hsa-miR-1243
AACTGGATCAATTATAGGAGTG
>hsa-miR-25-3p
CATTGCACTTGTCTCGGTCTGA
>hsa-miR-938
TGCCCTTAAAGGTGAACCCAGT
>hsa-miR-539-5p
GGAGAAATTATCCTTGGTGTGT
>hsa-miR-1913
TCTGCCCCCTCCGCTGCTGCCA
>hsa-miR-532-5p
CATGCCTTGAGTGTAGGACCGT
>hsa-miR-4793-5p
ACATCCTGCTCCACAGGGCAGAGG
>hsa-miR-9899
CGGGCGCCGCGCTCCCGCCCGC
>hsa-miR-744-5p
TGCGGGGCTAGGGCTAACAGCA
>hsa-miR-5189-5p
TCTGGGCACAGGCGGATGGACAGG
>hsa-miR-5094
AATCAGTGAATGCCTTGAACCT
>hsa-miR-6785-3p
ACATCGCCCCACCTTCCCCAG
>hsa-miR-6500-5p
AGGAGCTATCCACTCCAGGTGTCC
>hsa-miR-6895-3p

TGTCTCTCGCCCTTGGCCTTAG
>hsa-miR-4433a-3p
ACAGGAGTGGGGGTGGGACAT
>hsa-miR-432-3p
CTGGATGGCTCCTCCATGTCT
>hsa-miR-5194
TGAGGGGTTTGGAATGGGATGG
>hsa-miR-6072
TCCTCATCACACTGCACCTTAG
>hsa-miR-7159-3p
TTTCTATGTTAGTTGGAAG
>hsa-miR-3662
GAAAATGATGAGTAGTGACTGATG
>hsa-miR-4669
TGTGTCCGGGAAGTGGAGGAGG
>hsa-miR-4720-3p
TGCTTAAGTTGTACCAAGTAT
>hsa-miR-664a-5p
ACTGGCTAGGGAAAATGATTGGAT
>hsa-miR-375-5p
GCGACGAGCCCCTCGCACAAACC
>hsa-miR-4298
CTGGGACAGGAGGAGGAGGCAG
>hsa-miR-6800-5p
GTAGGTGACAGTCAGGGGCGG
>hsa-miR-122b-5p
TTTAGTGTGATAATGGCGTTTGA
>hsa-let-7g-5p
TGAGGTAGTAGTTTGTACAGTT
>hsa-miR-3121-5p
TCCTTTGCCTATTCTATTTAAG
>hsa-miR-6511a-3p
CCTCACCATCCCTTCTGCCTGC
>hsa-miR-7106-5p
TGGGAGGAGGGGATCTTGGG
>hsa-miR-212-3p
TAACAGTCTCCAGTCACGGCC
>hsa-miR-339-5p
TCCCTGTCCTCCAGGAGCTCACG
>hsa-miR-575
GAGCCAGTTGGACAGGAGC
>hsa-miR-11181-3p
AGGAGGAGGAGGTCAGGC
>hsa-miR-138-5p
AGCTGGTGTTGTGAATCAGGCCG
>hsa-miR-328-3p
CTGGCCCTCTCTGCCCTTCCGT
>hsa-miR-10527-5p
AAAGCAAATGTTGGGTGAACGGC
>hsa-miR-4440
TGTCGTGGGGCTTGCTGGCTTG
>hsa-miR-365a-5p
AGGGACTTTTGGGGGCAGATGTG
>hsa-miR-4635
TCTTGAAGTCAGAACCCGCAA
>hsa-miR-548ao-5p
AGAAGTAACTACGGTTTTTGCA
>hsa-miR-668-5p
TGCGCCTCGGGTGAGCATG
>hsa-miR-8056
CGTGGATTGTCTGGATGCAT
>hsa-miR-4637

TACTAACTGCAGATTCAAGTGA
>hsa-miR-518a-3p
GAAAGCGCTTCCCTTTGCTGGA
>hsa-miR-12124
GAGGAAATGCAGATGCTGGA
>hsa-miR-3148
TGGAAAAAACTGGTGTGTGCTT
>hsa-miR-1252-3p
CAAATGAGCTTAATTTCCTTTT
>hsa-miR-6129
TGAGGGAGTTGGGTGTATA
>hsa-miR-3135b
GGCTGGAGCGAGTGCAGTGGTG
>hsa-miR-4468
AGAGCAGAAGGATGAGAT
>hsa-miR-147a
GTGTGTGGAAATGCTTCTGC
>hsa-miR-6881-5p
TGGGGTAAGGATAGGAGGGTCA
>hsa-miR-660-5p
TACCCATTGCATATCGGAGTTG
>hsa-miR-520e-3p
AAAGTGCTTCCTTTTTGAGGG
>hsa-miR-567
AGTATGTTCTTCCAGGACAGAAC
>hsa-miR-8081
CTTGAGTCGTGCCTTTCTGAATG
>hsa-miR-324-5p
CGCATCCCCTAGGGCATTGGTG
>hsa-miR-3944-5p
TGTGCAGCAGGCCAACCGAGA
>hsa-miR-3672
ATGAGACTCATGTAAAACATCTT
>hsa-miR-6791-5p
CCCCTGGGGCTGGGCAGGCGGA
>hsa-miR-181b-5p
AACATTCATTGCTGTCGGTGGGT
>hsa-miR-3140-3p
AGCTTTTGGGAATTCAGGTAGT
>hsa-miR-6876-3p
AGCTGTCTGTGTTTTCCTTCTCAG
>hsa-miR-4471
TGGGAACTTAGTAGAGGTTTAA
>hsa-miR-6871-5p
CATGGGAGTTCGGGGTGGTTGC
>hsa-miR-642a-3p
AGACACATTTGGAGAGGGAACC
>hsa-miR-1272
GATGATGATGGCAGCAAATTCTGAAA
>hsa-miR-3176
ACTGGCCTGGGACTACCGG
>hsa-miR-381-3p
TATACAAGGGCAAGCTCTCTGT
>hsa-miR-1908-5p
CGGCGGGGACGGCGATTGGTC
>hsa-miR-3166
CGCAGACAATGCCTACTGGCCTA
>hsa-miR-8059
GGGGAACTGTAGATGAAAAGGC
>hsa-miR-4639-5p
TTGCTAAGTAGGCTGAGATTGA
>hsa-miR-6890-5p

CATGGGGTAGGGCAGAGTAGG
>hsa-miR-3059-3p
CCTCTAGGGAAGAGAAGGTTGG
>hsa-miR-10397-5p
TCCTTGACCTGATGCTGTAGGG
>hsa-miR-6840-3p
GCCCAGGACTTTGTGCGGGGTG
>hsa-miR-585-5p
CTAGCACACAGATACGCCCAGA
>hsa-miR-877-3p
TCCTCTTCTCCCTCCTCCCAG
>hsa-miR-425-3p
ATCGGGAATGTCGTGTCCGCCC
>hsa-miR-5702
TGAGTCAGCAACATATCCCATG
>hsa-miR-4782-5p
TTCTGGATATGAAGACAATCAA
>hsa-miR-6500-3p
ACACTTGTTGGGATGACCTGC
>hsa-miR-11181-5p
GTCTGACCAACCTCCTCCCGC
>hsa-miR-483-3p
TCACTCCTCTCCTCCCGTCTT
>hsa-miR-4763-3p
AGGCAGGGGCTGGTGCTGGGCGGG
>hsa-miR-506-3p
TAAGGCACCCTTCTGAGTAGA
>hsa-miR-6079
TTGGAAGCTTGGACCAACTAGCTG
>hsa-miR-1343-3p
CTCCTGGGGCCCGCACTCTCGC
>hsa-miR-6799-3p
TGCCCTGCATGGTGTCCCCACAG
>hsa-miR-608
AGGGGTGGTGTTGGGACAGCTCCGT
>hsa-miR-151a-3p
CTAGACTGAAGCTCCTTGAGG
>hsa-miR-4679
TCTGTGATAGAGATTCTTTGCT
>hsa-miR-1279
TCATATTGCTTCTTTCT
>hsa-miR-4674
CTGGGCTCGGGACGCGCGGCT
>hsa-miR-6823-5p
TCAGGGTTGGTAGGGGTTGCT
>hsa-miR-4300
TGGGAGCTGGACTACTTC
>hsa-miR-671-5p
AGGAAGCCCTGGAGGGGCTGGAG
>hsa-miR-30b-5p
TGTAAACATCCTACACTCAGCT
>hsa-miR-7162-5p
TGCTTCCTTTCTCAGCTG
>hsa-miR-4431
GCGACTCTGAAAACTAGAAGGT
>hsa-miR-8077
GGCTGAGTGGGGTTCTGACTCC
>hsa-miR-6827-5p
TGGGAGCCATGAGGGTCTGTGC
>hsa-miR-556-3p
ATATTACCATTAGCTCATCTTT
>hsa-miR-3117-3p

ATAGGACTCATATAGTGCCAG
>hsa-miR-217-3p
CATCAGTTCCTAATGCATTGCC
>hsa-miR-218-2-3p
CATGGTTCTGTCAAGCACCGCG
>hsa-let-7d-5p
AGAGGTAGTAGGTTGCATAGTT
>hsa-miR-92a-1-5p
AGGTTGGGATCGGTTGCAATGCT
>hsa-miR-6082
GAATACGTCTGGTTGATCC
>hsa-miR-6125
GCGGAAGGCGGAGCGGCGGA
>hsa-miR-8085
TGGGAGAGAGGACTGTGAGGC
>hsa-miR-3680-3p
TTTTGCATGACCCTGGGAGTAGG
>hsa-miR-3682-5p
CTACTTCTACCTGTGTTATCAT
>hsa-miR-4465
CTCAAGTAGTCTGACCAGGGGA
>hsa-miR-1237-5p
CGGGGGCGGGGCCGAAGCGCG
>hsa-miR-4511
GAAGAACTGTTGCATTTGCCCT
>hsa-miR-12123
TTATTCATTCACAAAAGCTTTA
>hsa-miR-9851-5p
CGCTGGCAGTGTTGGTGACACT
>hsa-miR-375-3p
TTTGTTCGTTCGGCTCGCGTGA
>hsa-miR-28-3p
CACTAGATTGTGAGCTCCTGGA
>hsa-miR-647
GTGGCTGCACTCACTTCCTTC
>hsa-miR-652-3p
AATGGCGCCACTAGGGTTGTG
>hsa-miR-1234-3p
TCGGCCTGACCACCCACCCCAC
>hsa-miR-4474-5p
TTAGTCTCATGATCAGACACA
>hsa-miR-4463
GAGACTGGGGTGGGGCC
>hsa-miR-551b-5p
GAAATCAAGCGTGGGTGAGACC
>hsa-miR-4450
TGGGGATTTGGAGAAGTGGTGA
>hsa-miR-3622a-3p
TCACCTGACCTCCCATGCCTGT
>hsa-miR-6766-5p
CGGGTGGGAGCAGATCTTATTGAG
>hsa-miR-103a-2-5p
AGCTTCTTTACAGTGCTGCCTTG
>hsa-miR-1225-3p
TGAGCCCCTGTGCCGCCCCCAG
>hsa-miR-12120
TAAGGAACGCGGGGCCTTGGTAGAGC
>hsa-miR-4286
ACCCCACTCCTGGTACC
>hsa-miR-34c-3p
AATCACTAACCACACGGCCAGG
>hsa-miR-6758-3p

ACTCATTCTCCTCTGTCCAG
>hsa-miR-4441
ACAGGGAGGAGATTGTA
>hsa-miR-6887-3p
TCCCCTCCACTTTCCTCCTAG
>hsa-miR-450b-5p
TTTTGCAATATGTTCCTGAATA
>hsa-miR-4676-5p
GAGCCAGTGGTGAGACAGTGA
>hsa-miR-1287-5p
TGCTGGATCAGTGGTTCGAGTC
>hsa-miR-3127-5p
ATCAGGGCTTGTGGAATGGGAAG
>hsa-miR-3945
AGGGCATAGGAGAGGGTTGATAT
>hsa-miR-7112-3p
TGCATCACAGCCTTTGGCCCTAG
>hsa-miR-2053
GTGTTAATTAAACCTCTATTTAC
>hsa-miR-934
TGTCTACTACTGGAGACACTGG
>hsa-miR-3169
TAGGACTGTGCTTGGCACATAG
>hsa-miR-7851-3p
TACCTGGGAGACTGAGGTTGGA
>hsa-miR-152-3p
TCAGTGCATGACAGAACTTGG
>hsa-miR-586
TATGCATTGTATTTTTAGGTCC
>hsa-miR-24-3p
TGGCTCAGTTCAGCAGGAACAG
>hsa-miR-3939
TACGCGCAGACCACAGGATGTC
>hsa-miR-423-5p
TGAGGGGCAGAGAGCGAGACTTT
>hsa-miR-585-3p
TGGGCGTATCTGTATGCTA
>hsa-miR-215-3p
TCTGTCATTTCTTTAGGCCAATA
>hsa-miR-6804-3p
CGCACCTGCCTCTCACCCACAG
>hsa-miR-3621
CGCGGGTCGGGGTCTGCAGG
>hsa-miR-4802-3p
TACATGGATGGAAACCTTCAAGC
>hsa-miR-2467-3p
AGCAGAGGCAGAGAGGCTCAGG
>hsa-miR-548at-5p
AAAAGTTATTGCGGTTTTGGCT
>hsa-miR-4254
GCCTGGAGCTACTCCACCATCTC
>hsa-miR-6874-3p
CAGTTCTGCTGTTCTGACTCTAG
>hsa-miR-374c-5p
ATAATACAACCTGCTAAGTGCT
>hsa-miR-6888-3p
ATCTGTCTCGATTGTTTCCAG
>hsa-miR-339-3p
TGAGCGCCTCGACGACAGAGCCG
>hsa-miR-3152-5p
ATTGCCTCTGTTCTAACACAAG
>hsa-miR-6771-3p

CAAACCCCTGTCTACCCGCAG
>hsa-miR-5587-3p
GCCCCGGGCAGTGTGATCATC
>hsa-miR-4318
CACTGTGGGTACATGCT
>hsa-miR-618
AAACTCTACTTGTCCTTCTGAGT
>hsa-miR-6720-5p
TTCCAGCCCTGGTAGGCGCCGCG
>hsa-miR-5006-5p
TTGCCAGGGCAGGAGGTGGAA
>hsa-miR-4434
AGGAGAAGTAAAGTAGAA
>hsa-miR-548au-5p
AAAAGTAATTGCGGTTTTTGC
>hsa-miR-10522-5p
AGAAGAATTGGCCTACTCAGG
>hsa-miR-4676-3p
CACTGTTTCACCACTGGCTCTT
>hsa-miR-499a-5p
TTAAGACTTGCAGTGATGTTT
>hsa-miR-181d-3p
CCACCGGGGGATGAATGTCAC
>hsa-miR-331-3p
GCCCCTGGGCCTATCCTAGAA
>hsa-miR-605-3p
AGAAGGCACTATGAGATTTAGA
>hsa-miR-7-5p
TGGAAGACTAGTGATTTTGTTGTT
>hsa-miR-3960
GGCGGCGGCGGAGGCGGGGG
>hsa-miR-4652-3p
GTTCTGTTAACCCATCCCCTCA
>hsa-miR-501-3p
AATGCACCCGGGCAAGGATTCT
>hsa-miR-6858-3p
CAGCCAGCCCCTGCTCACCCCT
>hsa-miR-6891-5p
TAAGGAGGGGGATGAGGGG
>hsa-miR-640
ATGATCCAGGAACCTGCCTCT
>hsa-miR-1247-5p
ACCCGTCCCGTTCGTCCCCGGA
>hsa-miR-10526-3p
AAAAGGGGGCTGAGGTGGAG
>hsa-miR-526a-5p|hsa-miR-520c-5p|hsa-miR-518d-5p
CTCTAGAGGGAAGCACTTTCTG
>hsa-miR-7109-5p
CTGGGGGGAGGAGACCCTGCT
>hsa-miR-3713
GGTATCCGTTTGGGGATGGT
>hsa-miR-3074-3p
GATATCAGCTCAGTAGGCACCG
>hsa-miR-4697-5p
AGGGGGCGCAGTCACTGACGTG
>hsa-miR-3137
TCTGTAGCCTGGGAGCAATGGGGT
>hsa-miR-449c-3p
TTGCTAGTTGCACTCCTCTCTGT
>hsa-miR-199a-5p
CCCAGTGTTCAGACTACCTGTTC
>hsa-miR-5585-3p

CTGAATAGCTGGGACTACAGGT
>hsa-miR-572
GTCCGCTCGGCGGTGGCCCA
>hsa-miR-1537-3p
AAAACCGTCTAGTTACAGTTGT
>hsa-miR-6782-3p
CACCTTTGTGTCCCCATCCTGCA
>hsa-miR-138-1-3p
GCTACTTCACAACACCAGGGCC
>hsa-miR-505-5p
GGGAGCCAGGAAGTATTGATGT
>hsa-miR-9902
CCCAGAAATCTGGTATGCCAGC
>hsa-miR-892c-5p
TATTCAGAAAGGTGCCAGTCA
>hsa-miR-1295a
TTAGGCCGCAGATCTGGGTGA
>hsa-miR-6880-5p
TGGTGGAGGAAGAGGGCAGCTC
>hsa-miR-670-3p
TTTCCTCATATTCATTCAGGA
>hsa-miR-6717-5p
AGGCGATGTGGGGATGTAGAGA
>hsa-miR-615-3p
TCCGAGCCTGGGTCTCCCTCTT
>hsa-miR-124-3p
TAAGGCACGCGGTGAATGCCAA
>hsa-miR-1238-3p
CTTCCTCGTCTGTCTGCCCC
>hsa-miR-1470
GCCCTCCGCCCGTGCACCCCG
>hsa-miR-2276-3p
TCTGCAAGTGTCAGAGGCGAGG
>hsa-miR-6801-5p
TGGTCAGAGGCAGCAGGAAATGA
>hsa-miR-202-5p
TTCCTATGCATATACTTCTTTG
>hsa-miR-374a-3p
CTTATCAGATTGTATTGTAATT
>hsa-miR-6844
TTCTTTGTTTTTAATTCACAG
>hsa-miR-222-5p
CTCAGTAGCCAGTGTAGATCCT
>hsa-miR-3912-5p
ATGTCCATATTATGGGTTAGT
>hsa-miR-6774-3p
TCGTGTCCCTCTTGTCCACAG
>hsa-miR-95-3p
TTCAACGGGTATTTATTGAGCA
>hsa-miR-3158-3p
AAGGGCTTCCTCTCTGCAGGAC
>hsa-miR-3177-3p
TGCACGGCACTGGGGACACGT
>hsa-miR-5011-5p
TATATATACAGCCATGCACTC
>hsa-miR-602
GACACGGGCGACAGCTGCGGCCC
>hsa-miR-619-3p
GACCTGGACATGTTTGTGCCCAGT
>hsa-miR-548t-5p
CAAAAGTGATCGTGGTTTTTG
>hsa-miR-541-3p

TGGTGGGCACAGAATCTGGACT
>hsa-miR-1289
TGGAGTCCAGGAATCTGCATTTT
>hsa-miR-6508-3p
TGGGCCATGCATTTCTAGAACT
>hsa-miR-7975
ATCCTAGTCACGGCACCA
>hsa-miR-4666a-3p
CATACAATCTGACATGTATTT
>hsa-miR-6859-3p
TGACCCCCATGTCGCCTCTGTAG
>hsa-miR-10a-3p
CAAATTCGTATCTAGGGGAATA
>hsa-miR-4693-5p
ATACTGTGAATTTCACTGTCACA
>hsa-miR-6822-5p
CAGGGAACCAGTTGGGGCTT
>hsa-miR-181c-3p
AACCATCGACCGTTGAGTGGAC
>hsa-miR-6827-3p
ACCGTCTCTTCTGTTCCCCAG
>hsa-miR-4531
ATGGAGAAGGCTTCTGA
>hsa-miR-6891-3p
CCCTCATCTTCCCCTCCTTTC
>hsa-miR-5186
AGAGATTGGTAGAAATCAGGT
>hsa-miR-548au-3p
TGGCAGTTACTTTTGCACCAG
>hsa-miR-1182
GAGGGTCTTGGGAGGGATGTGAC
>hsa-miR-892c-3p
CACTGTTTCCTTTCTGAGTGGA
>hsa-miR-27a-5p
AGGGCTTAGCTGCTTGTGAGCA
>hsa-miR-4731-5p
TGCTGGGGGCCACATGAGTGTG
>hsa-miR-520a-3p
AAAGTGCTTCCCTTTGGACTGT
>hsa-miR-4798-5p
TTCGGTATACTTTGTGAATTGG
>hsa-miR-4252
GGCCACTGAGTCAGCACCA
>hsa-miR-4324
CCCTGAGACCCTAACCTTAA
>hsa-miR-4732-5p
TGTAGAGCAGGGAGCAGGAAGCT
>hsa-miR-642a-5p
GTCCCTCTCCAAATGTGTCTTG
>hsa-miR-4645-3p
AGACAGTAGTTCTTGCCTGGTT
>hsa-miR-4662a-5p
TTAGCCAATTGTCCATCTTTAG
>hsa-miR-3684
TTAGACCTAGTACACGTCCTT
>hsa-miR-1200
CTCCTGAGCCATTCTGAGCCTC
>hsa-miR-4711-3p
CGTGTCTTCTGGCTTGAT
>hsa-miR-6515-3p
TCTCTTCATCTACCCCCCAG
>hsa-miR-513a-3p

TAAATTTCACCTTTCTGAGAAGG
>hsa-miR-383-5p
AGATCAGAAGGTGATTGTGGCT
>hsa-miR-1301-3p
TTGCAGCTGCCTGGGAGTGACTTC
>hsa-miR-26a-1-3p
CCTATTCTTGGTTACTTGCACG
>hsa-miR-3934-3p
TGCTCAGGTTGCACAGCTGGGA
>hsa-miR-6826-5p
TCAATAGGAAAGAGGTGGGACCT
>hsa-miR-4734
GCTGCGGGCTGCGGTCAGGGCG
>hsa-miR-499b-3p
AACATCACTGCAAGTCTTAACA
>hsa-miR-148b-3p
TCAGTGCATCACAGAACTTTGT
>hsa-miR-486-5p
TCCTGTACTGAGCTGCCCCGAG
>hsa-miR-155-3p
CTCCTACATATTAGCATTAACA
>hsa-miR-4673
TCCAGGCAGGAGCCGGACTGGA
>hsa-miR-761
GCAGCAGGGTGAAACTGACACA
>hsa-miR-3118
TGTGACTGCATTATGAAAATTCT
>hsa-miR-1199-5p
CCTGAGCCCGGGCCGCGCAG
>hsa-miR-210-5p
AGCCCCTGCCCACCGCACACTG
>hsa-miR-512-3p
AAGTGCTGTCATAGCTGAGGTC
>hsa-miR-1297
TTCAAGTAATTCAGGTG
>hsa-miR-3180-3p
TGGGGCGGAGCTTCCGGAGGCC
>hsa-miR-4770
TGAGATGACACTGTAGCT
>hsa-miR-5089-5p
GTGGGATTTCTGAGTAGCATC
>hsa-miR-6848-5p
TGGGGGCTGGGATGGGCCATGGT
>hsa-miR-548p
TAGCAAAAACTGCAGTTACTTT
>hsa-miR-2054
CTGTAATATAAATTTAATTTATT
>hsa-miR-4708-5p
AGAGATGCCGCCTTGCTCCTT
>hsa-miR-224-5p
TCAAGTCACTAGTGGTTCCGTTTAG
>hsa-miR-6831-3p
TGACTAACTCCCACTCTACAG
>hsa-miR-4437
TGGGCTCAGGGTACAAAGGTT
>hsa-miR-4667-5p
ACTGGGGAGCAGAAGGAGAACC
>hsa-miR-1249-3p
ACGCCCTTCCCCCCCTTCTTCA
>hsa-miR-4639-3p
TCACTCTCACCTTGCTTTGC
>hsa-miR-4509

ACTAAAGGATATAGAAGGTTTT
>hsa-miR-499a-3p
AACATCACAGCAAGTCTGTGCT
>hsa-miR-3617-3p
CATCAGCACCCTATGTCCTTTCT
>hsa-miR-6127
TGAGGGAGTGGGTGGGAGG
>hsa-miR-323b-3p
CCCAATACACGGTCGACCTCTT
>hsa-miR-5586-3p
CAGAGTGACAAGCTGGTTAAAG
>hsa-miR-6077
GGGAAGAGCTGTACGGCCTTC
>hsa-miR-3120-5p
CCTGTCTGTGCCTGCTGTACA
>hsa-miR-5688
TAACAAACACCTGTAAAACAGC
>hsa-miR-192-5p
CTGACCTATGAATTGACAGCC
>hsa-miR-4701-3p
ATGGGTGATGGGTGTGGTGT
>hsa-miR-6734-3p
CCCTTCCCTCACTCTTCTCTCAG
>hsa-miR-1233-5p
AGTGGGAGGCCAGGGCACGGCA
>hsa-miR-8055
CTTTGAGCACATGAGCAGACGGA
>hsa-miR-3675-5p
TATGGGGCTTCTGTAGAGATTTC
>hsa-miR-3685
TTTCCTACCCTACCTGAAGACT
>hsa-miR-6888-5p
AAGGAGATGCTCAGGCAGAT
>hsa-miR-1295b-3p
AATAGGCCACGGATCTGGGCAA
>hsa-miR-1306-3p
ACGTTGGCTCTGGTGGTG
>hsa-miR-4429
AAAAGCTGGGCTGAGAGGCG
>hsa-miR-4519
CAGCAGTGCGCAGGGCTG
>hsa-miR-362-5p
AATCCTTGGAACCTAGGTGTGAGT
>hsa-miR-3929
GAGGCTGATGTGAGTAGACCACT
>hsa-miR-6783-3p
TTCCTGGGCTTCTCCTCTGTAG
>hsa-miR-711
GGGACCCAGGGAGAGACGTAAG
>hsa-miR-4715-3p
GTGCCACCTTAACTGCAGCCAAT
>hsa-miR-93-5p
CAAAGTGCTGTTCGTGCAGGTAG
>hsa-miR-1323
TCAAAACTGAGGGGCATTTTCT
>hsa-miR-622
ACAGTCTGCTGAGGTTGGAGC
>hsa-miR-6895-5p
CAGGGCCAGGCACAGAGTAAG
>hsa-miR-639
ATCGCTGCGGTTGCGAGCGCTGT
>hsa-miR-596

AAGCCTGCCCGGCTCCTCGGG
>hsa-miR-4738-3p
TGAAACTGGAGCGCCTGGAGGA
>hsa-miR-874-3p
CTGCCCTGGCCCGAGGGACCGA
>hsa-miR-4640-3p
CACCCCCTGTTTCCTGGCCCAC
>hsa-miR-6752-3p
TCCCTGCCCCCATACTCCCAG
>hsa-miR-12113
TCTGAACTCTATGTGGGATTAG
>hsa-miR-1252-5p
AGAAGGAAATTGAATTCATTTA
>hsa-miR-6752-5p
GGGGGGTGTGGAGCCAGGGGGC
>hsa-miR-1843
TATGGAGGTCTCTGTCTGGC
>hsa-miR-320b
AAAAGCTGGGTTGAGAGGGCAA
>hsa-miR-4697-3p
TGTCAGTGACTCCTGCCCCTTGGT
>hsa-miR-4310
GCAGCATTCATGTCCC
>hsa-miR-188-5p
CATCCCTTGCATGGTGGAGGG
>hsa-miR-183-3p
GTGAATTACCGAAGGGCCATAA
>hsa-miR-4772-3p
CCTGCAACTTTGCCTGATCAGA
>hsa-miR-33b-5p
GTGCATTGCTGTTGCATTGC
>hsa-miR-4314
CTCTGGGAAATGGGACAG
>hsa-miR-3934-5p
TCAGGTGTGGAAACTGAGGCAG
>hsa-miR-550a-3-5p
AGTGCCTGAGGGAGTAAGAG
>hsa-miR-1199-3p
TGCGGCCGGTGCTCAACCTGC
>hsa-miR-4776-5p
GTGGACCAGGATGGCAAGGGCT
>hsa-miR-301b-3p
CAGTGCAATGATATTGTCAAAGC
>hsa-miR-6828-5p
AGGAAGCAAGAGAACCCTGTGG
>hsa-miR-371a-3p
AAGTGCCGCCATCTTTTGAGTGT
>hsa-miR-6794-5p
CAGGGGGACTGGGGGTGAGC
>hsa-miR-302d-3p
TAAGTGCTTCCATGTTTGAGTGT
>hsa-miR-4304
CCGGCATGTCCAGGGCA
>hsa-miR-6889-5p
TCGGGGAGTCTGGGGTCCGGAAT
>hsa-miR-30b-3p
CTGGGAGGTGGATGTTTACTTC
>hsa-miR-4645-5p
ACCAGGCAAGAAATATTGT
>hsa-miR-6506-3p
TCGTATCAGAGATTCCAGACAC
>hsa-miR-2116-3p

CCTCCCATGCCAAGAACTCCC
>hsa-miR-6832-3p
ACCCTTTTTCTCTTTCCCAG
>hsa-miR-17-3p
ACTGCAGTGAAGGCACTTGTAG
>hsa-miR-642b-3p
AGACACATTTGGAGAGGGACCC
>hsa-miR-4534
GGATGGAGGAGGGGTCT
>hsa-miR-664a-3p
TATTCATTTATCCCCAGCCTACA
>hsa-miR-2861
GGGGCCTGGCGGTGGGCGG
>hsa-miR-4750-3p
CCTGACCCACCCCCTCCCGCAG
>hsa-miR-4647
GAAGATGGTGCTGTGCTGAGGAA
>hsa-miR-376b-3p
ATCATAGAGGAAAATCCATGTT
>hsa-miR-595
GAAGTGTGCCGTGGTGTGTCT
>hsa-miR-4693-3p
TGAGAGTGGAATTCACAGTATTT
>hsa-miR-6765-5p
GTGAGGCGGGGCCAGGAGGGTGTGT
>hsa-let-7b-3p
CTATACAACCTACTGCCTTCCC
>hsa-miR-3064-5p
TCTGGCTGTTGTGGTGTGCAA
>hsa-miR-500b-3p
GCACCCAGGCAAGGATTCTG
>hsa-miR-6776-3p
CAACCACCACTGTCTCTCCCCAG
>hsa-miR-6768-5p
CACACAGGAAAAGCGGGGCCCTG
>hsa-miR-487b-5p
GTGGTTATCCCTGTCCTGTTCG
>hsa-miR-4432
AAAGACTCTGCAAGATGCCT
>hsa-miR-3145-3p
AGATATTTTGAGTGTTTGGAATTG
>hsa-miR-4740-3p
GCCCGAGAGGATCCGTCCCTGC
>hsa-miR-6863
TAGACGTGGTGAAGGATTGAGTG
>hsa-miR-1287-3p
CTCTAGCCACAGATGCAGTGAT
>hsa-miR-204-5p
TTCCCTTTGTCATCCTATGCCT
>hsa-miR-492
AGGACCTGCGGGACAAGATTCTT
>hsa-miR-4520-5p
CCTGCGTGTTTTCTGTCCAA
>hsa-miR-20a-5p
TAAAGTGCTTATAGTGCAGGTAG
>hsa-miR-518c-5p
TCTCTGGAGGGAAGCACTTTCTG
>hsa-miR-5580-5p
TGCTGGCTCATTTCATATGTGT
>hsa-miR-4713-3p
TGGGATCCAGACAGTGGGAGAA
>hsa-miR-324-3p

CCCACTGCCCCAGGTGCTGCTGG
>hsa-miR-548ag
AAAGGTAATTGTGGTTTCTGC
>hsa-miR-6883-5p
AGGGAGGGTGTGGTATGGATGT
>hsa-miR-29c-5p
TGACCGATTTCTCCTGGTGTTC
>hsa-miR-3059-5p
TTTCCTCTCTGCCCCATAGGGTGT
>hsa-miR-6871-3p
CAGCACCCTGTGGCTCCCACAG
>hsa-miR-190b-3p
ACTAAATGTCAAACATATTCT
>hsa-miR-373-3p
GAAGTGCTTCGATTTTGGGGTGT
>hsa-miR-146b-3p
GCCCTGTGGACTCAGTTCTGGT
>hsa-miR-340-3p
TCCGTCTCAGTTACTTTATAGC
>hsa-miR-31-5p
AGGCAAGATGCTGGCATAGCT
>hsa-miR-6715b-3p
CTCAAACCGGCTGTGCCTGTGG
>hsa-miR-3191-3p
TGGGGACGTAGCTGGCCAGACAG
>hsa-miR-1271-3p
AGTGCCTGCTATGTGCCAGGCA
>hsa-miR-6742-3p
ACCTGGGTTGTCCCCTCTAG
>hsa-miR-548ah-5p
AAAAGTGATTGCAGTGTTTG
>hsa-miR-16-1-3p
CCAGTATTAACTGTGCTGCTGA
>hsa-miR-4769-5p
GGTGGGATGGAGAGAAGGTATGAG
>hsa-miR-6069
GGGCTAGGGCCTGCTGCCCCC
>hsa-miR-548aw
GTGCAAAAGTCATCACGGTT
>hsa-miR-4291
TTCAGCAGGAACAGCT
>hsa-miR-12125
TCTCTCACCTGGCATAAGCAAT
>hsa-miR-548e-5p
CAAAAGCAATCGCGGTTTTTGC
>hsa-miR-1185-2-3p
ATATACAGGGGGAGACTCTCAT
>hsa-miR-5196-3p
TCATCCTCGTCTCCCTCCCAG
>hsa-miR-3085-5p
AGGTGCCATTCTGAGGGCCAGGAGT
>hsa-miR-8064
AGCACACTGAGCGAGCGGAC
>hsa-miR-675-3p
CTGTATGCCCTCACCGCTCA
>hsa-miR-3190-3p
TGTGGAAGGTAGACGGCCAGAGA
>hsa-miR-4321
TTAGCGGTGGACCGCCCTGCG
>hsa-miR-625-5p
AGGGGGAAAGTTCTATAGTCC
>hsa-miR-6789-5p

GTAGGGGCGTCCCGGGCGCGCGGG
>hsa-miR-9500
AAGGGAAGATGGTGACCAC
>hsa-miR-520e-5p
CTCAAGATGGAAGCAGTTTCTG
>hsa-miR-7515
AGAAGGGAAGATGGTGAC
>hsa-miR-548l
AAAAGTATTTGCGGGTTTTGTC
>hsa-miR-4506
AAATGGGTGGTCTGAGGCAA
>hsa-miR-4736
AGGCAGGTTATCTGGGCTG
>hsa-miR-6068
CCTGCGAGTCTCCGGCGGTGG
>hsa-miR-6810-3p
TCCCCTGCTCCCTTGTTCCCCAG
>hsa-miR-3682-3p
TGATGATACAGGTGGAGGTAG
>hsa-miR-21-5p
TAGCTTATCAGACTGATGTTGA
>hsa-miR-4272
CATTCAACTAGTGATTGT
>hsa-miR-6816-3p
GAAGGACCTGCACCTTCG
>hsa-miR-6807-5p
GTGAGCCAGTGGAATGGAGAGG
>hsa-miR-5582-5p
TAGGCACACTTAAAGTTATAGC
>hsa-miR-8485
CACACACACACACACACGTAT
>hsa-miR-6842-3p
TTGGCTGGTCTCTGCTCCGCAG
>hsa-miR-548ar-3p
TAAAACTGCAGTTATTTTTGC
>hsa-miR-6804-5p
TGAGGGTGTCAGCAGGTGACG
>hsa-miR-449c-5p
TAGGCAGTGTATTGCTAGCGGCTGT
>hsa-miR-488-5p
CCCAGATAATGGCACTCTCAA
>hsa-miR-518e-3p
AAAGCGCTTCCCTTCAGAGTG
>hsa-miR-520g-5p
TCTAGAGGAAGCACTTTCTGTTT
>hsa-miR-195-5p
TAGCAGCACAGAAATATTGGC
>hsa-miR-19a-3p
TGTGCAAATCTATGCAAAACTGA
>hsa-miR-592
TTGTGTCAATATGCGATGATGT
>hsa-miR-766-5p
AGGAGGAATTGGTGCTGGTCTT
>hsa-miR-628-5p
ATGCTGACATATTTACTAGAGG
>hsa-miR-494-5p
AGGTTGTCCGTGTTGTCTTCTCT
>hsa-miR-3611
TTGTGAAGAAAGAAATTCTTA
>hsa-miR-557
GTTTGCACGGGTGGGCCTTGTCT
>hsa-miR-659-3p

CTTGGTTCAGGGAGGGTCCCCA
>hsa-miR-3646
AAAATGAAATGAGCCCAGCCCA
>hsa-miR-8087
GAAGACTTCTTGGATTACAGGGG
>hsa-miR-4329
CCTGAGACCCTAGTTCCAC
>hsa-miR-4699-5p
AGAAGATTGCAGAGTAAGTTCC
>hsa-miR-5582-3p
TAAAACTTTAAGTGTGCCTAGG
>hsa-miR-10399-3p
CTCTCGGACAAGCTGTAGGTC
>hsa-miR-4646-5p
ACTGGGAAGAGGAGCTGAGGGA
>hsa-miR-6826-3p
CTCCCCTCTCTTTCCTGTTCAG
>hsa-miR-6726-3p
CTCGCCCTGTCTCCCGCTAG
>hsa-miR-3665
AGCAGGTGCGGGGCGGCG
>hsa-miR-3144-3p
ATATACCTGTTCGGTCTCTTTA
>hsa-miR-7150
CTGGCAGGGGGAGAGGTA
>hsa-miR-6800-3p
CACCTCTCCTGGCATCGCCCC
>hsa-miR-4424
AGAGTTAACTCAAAATGGACTA
>hsa-miR-5195-5p
AACCCCTAAGGCAACTGGATGG
>hsa-miR-6514-5p
TATGGAGTGGACTTTCAGCTGGC
>hsa-miR-4294
GGGAGTCTACAGCAGGG
>hsa-miR-92b-5p
AGGGACGGGACGCGGTGCAGTG
>hsa-miR-4659a-5p
CTGCCATGTCTAAGAAGAAAAC
>hsa-miR-6882-3p
TGCTGCCTCTCCTCTTGCCTGCAG
>hsa-miR-346
TGTCTGCCCGCATGCCTGCCTCT
>hsa-miR-5003-3p
TACTTTTCTAGGTTGTTGGGG
>hsa-miR-548at-3p
CAAAACCGCAGTAACTTTTGT
>hsa-miR-1269b
CTGGACTGAGCCATGCTACTGG
>hsa-miR-6819-5p
TTGGGGTGGAGGGCCAAGGAGC
>hsa-miR-4795-3p
ATATTATTAGCCACTTCTGGAT
>hsa-miR-4709-3p
TTGAAGAGGAGGTGCTCTGTAGC
>hsa-miR-3139
TAGGAGCTCAACAGATGCCTGTT
>hsa-miR-4687-5p
CAGCCCTCCTCCCGCACCCAAA
>hsa-miR-943
CTGACTGTTGCCGTCCTCCAG
>hsa-miR-6856-3p

TACAGCCCTGTGATCTTTCCAG
>hsa-miR-1911-3p
CACCAGGCATTGTGGTCTCC
>hsa-miR-6733-3p
TCAGTGTCTGGATTTCCTAG
>hsa-miR-4433a-5p
CGTCCCACCCCCCACTCCTGT
>hsa-miR-6509-5p
ATTAGGTAGTGGCAGTGGAAC
>hsa-miR-15b-3p
CGAATCATTATTTGCTGCTCTA
>hsa-miR-4749-5p
TGCGGGGACAGGCCAGGGCATC
>hsa-miR-513c-3p
TAAATTTCACCTTTCTGAGAAGA
>hsa-miR-887-3p
GTGAACGGGCGCCATCCCGAGG
>hsa-miR-144-5p
GGATATCATCATATACTGTAAG
>hsa-miR-6766-3p
TGATTGTCTTCCCCCACCCTCA
>hsa-miR-2355-5p
ATCCCCAGATACAATGGACAA
>hsa-miR-3974
AAAGGTCATTGTAAGGTTAATGC
>hsa-miR-551a
GCGACCCACTCTTGGTTTCCA
>hsa-miR-4520-3p
TTGGACAGAAAACACGCAGGAA
>hsa-miR-12119
TTCTGAGGGGACGGTAGATTTGGGG
>hsa-miR-545-3p
TCAGCAAACATTTATTGTGTGC
>hsa-miR-6124
GGGAAAAGGAAGGGGGAGGA
>hsa-miR-378h
ACTGGACTTGGTGTCAGATGG
>hsa-miR-6864-3p
GTGAGACTTCTCTCCCTTCAG
>hsa-miR-876-3p
TGGTGGTTTACAAAGTAATTCA
>hsa-miR-4642
ATGGCATCGTCCCCTGGTGGCT
>hsa-miR-887-5p
CTTGGGAGCCCTGTTAGACTC
>hsa-miR-10398-3p
GCCCGGAGAGCTGGGAGCCAG
>hsa-miR-2681-5p
GTTTTACCACCTCCAGGAGACT
>hsa-miR-210-3p
CTGTGCGTGTGACAGCGGCTGA
>hsa-miR-4467
TGGCGGCGGTAGTTATGGGCTT
>hsa-miR-7113-5p
TCCAGGGAGACAGTGTGTGAG
>hsa-miR-1184
CCTGCAGCGACTTGATGGCTTCC
>hsa-miR-4777-3p
ATACCTCATCTAGAATGCTGTA
>hsa-miR-8080
GAAGGACACTGGTGTCAACGGCT
>hsa-miR-664b-3p

TTCATTTGCCTCCCAGCCTACA
>hsa-miR-133b
TTTGGTCCCCTTCAACCAGCTA
>hsa-miR-550b-2-5p
ATGTGCCTGAGGGAGTAAGACA
>hsa-miR-6848-3p
GTGGTCTCTTGGCCCCCAG
>hsa-miR-23c
ATCACATTGCCAGTGATTACCC
>hsa-miR-663a
AGGCGGGGCGCCGCGGGACCGC
>hsa-miR-153-3p
TTGCATAGTCACAAAAGTGATC
>hsa-miR-1302
TTGGGACATACTTATGCTAAA
>hsa-miR-6499-5p
TCGGGCGCAAGAGCACTGCAGT
>hsa-miR-5088-5p
CAGGGCTCAGGGATTGGATGGAGG
>hsa-miR-513b-3p
AAATGTCACCTTTTTGAGAGGA
>hsa-miR-29a-3p
TAGCACCATCTGAAATCGGTTA
>hsa-miR-16-2-3p
CCAATATTACTGTGCTGCTTTA
>hsa-miR-6739-5p
TGGGAAAGAGAAAGAACAAGTA
>hsa-miR-6512-3p
TTCCAGCCCTTCTAATGGTAGG
>hsa-miR-449b-3p
CAGCCACAACTACCCTGCCACT
>hsa-miR-152-5p
AGGTTCTGTGATACACTCCGACT
>hsa-miR-3915
TTGAGGAAAAGATGGTCTTATT
>hsa-miR-581
TCTTGTGTTCTCTAGATCAGT
>hsa-miR-148a-3p
TCAGTGCACTACAGAACTTTGT
>hsa-miR-11401
TCACGTCTGCGGCTGTCACG
>hsa-miR-6855-5p
TTGGGGTTTGGGGTGCAGACATTGC
>hsa-miR-3613-3p
ACAAAAAAAAAAGCCCAACCCTTC
>hsa-miR-4786-5p
TGAGACCAGGACTGGATGCACC
>hsa-miR-4763-5p
CGCCTGCCCAGCCCTCCTGCT
>hsa-miR-548ap-5p
AAAAGTAATTGCGGTCTTT
>hsa-miR-30c-1-3p
CTGGGAGAGGGTTGTTTACTCC
>hsa-miR-12128
TTCAGGGATGGCGCATGAAGAGGAGA
>hsa-miR-3147
GGTTGGGCAGTGAGGAGGGTGTGA
>hsa-miR-548k
AAAAGTACTTGCGGATTTTGCT
>hsa-miR-4258
CCCCGCCACCGCCTTGG
>hsa-miR-5009-3p

TCCTAAATCTGAAAGTCCAAAA
>hsa-miR-3612
AGGAGGCATCTTGAGAAATGGA
>hsa-miR-10401-5p
CGTGTGGGAAGGCGTGGGGT
>hsa-miR-34a-5p
TGGCAGTGTCTTAGCTGGTTGT
>hsa-miR-4658
GTGAGTGTGGATCCTGGAGGAAT
>hsa-miR-4520-2-3p
TTTGGACAGAAAACACGCAGGT
>hsa-miR-4278
CTAGGGGGTTTGCCCTTG
>hsa-miR-412-5p
TGGTCGACCAGTTGGAAAGTAAT
>hsa-miR-148a-5p
AAAGTTCTGAGACACTCCGACT
>hsa-miR-548ax
AGAAGTAATTGCGGTTTTGCCA
>hsa-miR-6873-5p
CAGAGGGAATACAGAGGGCAAT
>hsa-miR-4288
TTGTCTGCTGAGTTTCC
>hsa-miR-6744-5p
TGGATGACAGTGGAGGCCT
>hsa-miR-3664-5p
AACTCTGTCTTCACTCATGAGT
>hsa-miR-192-3p
CTGCCAATTCCATAGGTCACAG
>hsa-miR-483-5p
AAGACGGGAGGAAAGAAGGGAG
>hsa-miR-548ac
CAAAAACCGGCAATTACTTTTG
>hsa-miR-4515
AGGACTGGACTCCCGGCAGCCC
>hsa-miR-4488
AGGGGGCGGGCTCCGGCG
>hsa-miR-6839-3p
TTGGGTTTTCTCTTCAATCCAG
>hsa-miR-302b-5p
ACTTTAACATGGAAGTGCTTTC
>hsa-miR-4756-5p
CAGGGAGGCGCTCACTCTCTGCT
>hsa-miR-3129-3p
AAACTAATCTCTACACTGCTGC
>hsa-miR-27b-5p
AGAGCTTAGCTGATTGGTGAAC
>hsa-miR-135a-2-3p
ATGTAGGGATGGAAGCCATGAA
>hsa-miR-5010-5p
AGGGGGATGGCAGAGCAAAATT
>hsa-miR-4746-3p
AGCGGTGCTCCTGCGGGCCGA
>hsa-miR-4433b-3p
CAGGAGTGGGGGGTGGGACGT
>hsa-miR-516b-3p|hsa-miR-516a-3p
TGCTTCCTTTCAGAGGGT
>hsa-miR-892b
CACTGGCTCCTTTCTGGGTAGA
>hsa-miR-3941
TTACACACAACTGAGGATCATA
>hsa-miR-146b-5p

TGAGAACTGAATTCCATAGGCTG
>hsa-miR-6837-3p
CCTTCACTGTGACTCTGCTGCAG
>hsa-miR-6886-5p
CCCGCAGGTGAGATGAGGGCT
>hsa-miR-372-3p
AAAGTGCTGCGACATTTGAGCGT
>hsa-miR-376c-5p
GGTGGATATTCCTTCTATGTT
>hsa-miR-4661-3p
CAGGATCCACAGAGCTAGTCCA
>hsa-miR-4683
TGGAGATCCAGTGCTCGCCCGAT
>hsa-miR-4516
GGGAGAAGGGTCGGGGC
>hsa-miR-450b-3p
TTGGGATCATTTTGCATCCATA
>hsa-miR-655-3p
ATAATACATGGTTAACCTCTTT
>hsa-miR-3922-3p
TCTGGCCTTGACTTGACTCTTT
>hsa-miR-10396a-3p
GGCCCCGGGCCCTCGACCGGG
>hsa-miR-6508-5p
TCTAGAAATGCATGACCCACC
>hsa-miR-4297
TGCCTTCCTGTCTGTG
>hsa-miR-4484
AAAAGGCGGGAGAAGCCCCA
>hsa-miR-6784-5p
GCCGGGGCTTTGGGTGAGGG
>hsa-miR-10523-5p
GACAATGATGAGAAGACCTGAGGA
>hsa-miR-4320
GGGATTCTGTAGCTTCCT
>hsa-miR-8060
CCATGAAGCAGTGGGTAGGAGGAC
>hsa-miR-3192-3p
CTCTGATCGCCCTCTCAGCTC
>hsa-miR-4302
CCAGTGTGGCTCAGCGAG
>hsa-miR-4460
ATAGTGGTTGTGAATTTACCTT
>hsa-miR-4787-3p
GATGCGCCGCCCACTGCCCCGCGC
>hsa-miR-4491
AATGTGGACTGGTGTGACCAAA
>hsa-miR-10400-5p
CGGCGGCGGCGGCTCTGGGCG
>hsa-miR-512-5p
CACTCAGCCTTGAGGGCACTTTC
>hsa-miR-6867-3p
CTCTCCCTCTTTACCCACTAG
>hsa-miR-526b-5p
CTCTTGAGGGAAGCACTTTCTGT
>hsa-miR-6730-3p
CCTGACACCCCATCTGCCCTCA
>hsa-miR-548e-3p
AAAAACTGAGACTACTTTTGCA
>hsa-miR-302e
TAAGTGCTTCCATGCTT
>hsa-miR-6872-3p

CCCATGCCTCCTGCCGCGGTC
>hsa-miR-1285-5p
GATCTCACTTTGTTGCCCAGG
>hsa-miR-3677-3p
CTCGTGGGCTCTGGCCACGGCC
>hsa-miR-1912-5p
CTCATTGCATGGGCTGTGTATA
>hsa-miR-365a-3p|hsa-miR-365b-3p
TAATGCCCCTAAAAATCCTTAT
>hsa-miR-326
CCTCTGGGCCCTTCCTCCAG
>hsa-miR-4426
GAAGATGGACGTACTTT
>hsa-miR-892a
CACTGTGTCCTTTCTGCGTAG
>hsa-miR-5704
TTAGGCCATCATCCCATTATGC
>hsa-miR-4474-3p
TTGTGGCTGGTCATGAGGCTAA
>hsa-miR-6803-5p
CTGGGGGTGGGGGGCTGGGCGT
>hsa-miR-3689a-5p|hsa-miR-3689b-5p|hsa-miR-3689e
TGTGATATCATGGTTCCTGGGA
>hsa-miR-361-5p
TTATCAGAATCTCCAGGGGTAC
>hsa-miR-6868-3p
TTCCTTCTGTTGTCTGTGCAG
>hsa-miR-6839-5p
TCTGGATTGAAGAGACGACCCA
>hsa-miR-6501-3p
CCAGAGCAGCCTGCGGTAACAGT
>hsa-miR-6741-3p
TCGGCTCTCTCCCTCACCCTAG
>hsa-miR-645
TCTAGGCTGGTACTGCTGA
>hsa-miR-4752
TTGTGGATCTCAAGGATGTGCT
>hsa-miR-4769-3p
TCTGCCATCCTCCCTCCCCTAC
>hsa-miR-554
GCTAGTCCTGACTCAGCCAGT
>hsa-miR-8088
CCTCGGTACTGGAAAGGGGTA
>hsa-miR-23b-5p
TGGGTTCCTGGCATGCTGATTT
>hsa-miR-4760-3p
AAATTCATGTTCAATCTAAACC
>hsa-miR-1233-3p
TGAGCCCTGTCCTCCCGCAG
>hsa-miR-6756-3p
TCCCCTTCCTCCCTGCCCAG
>hsa-miR-1538
CGGCCCGGGCTGCTGCTGTTCCT
>hsa-miR-4678
AAGGTATTGTTCAGACTTATGA
>hsa-miR-6126
GTGAAGGCCCGGCGGAGA
>hsa-miR-4638-3p
CCTGGACACCGCTCAGCCGGCCG
>hsa-miR-4687-3p
TGGCTGTTGGAGGGGGCAGGC
>hsa-miR-506-5p

```
TATTCAGGAAGGTGTTACTTAA
>hsa-miR-4773
CAGAACAGGAGCATAGAAAGGC
>hsa-miR-3181
ATCGGGCCCTCGGCGCCGG
>hsa-miR-151a-5p
TCGAGGAGCTCACAGTCTAGT
>hsa-let-7c-3p
CTGTACAACCTTCTAGCTTTCC
>hsa-miR-144-3p
TACAGTATAGATGATGTACT
>hsa-miR-8058
CTGGACTTTGATCTTGCCATAA
>hsa-miR-4802-5p
TATGGAGGTTCTAGACCATGTT
>hsa-miR-4653-5p
TCTCTGAGCAAGGCTTAACACC
>hsa-miR-3152-3p
TGTGTTAGAATAGGGGCAATAA
>hsa-miR-511-3p
AATGTGTAGCAAAAGACAGA
>hsa-miR-3692-3p
GTTCCACACTGACACTGCAGAAGT
>hsa-miR-1236-3p
CCTCTTCCCCTTGTCTCTCCAG
>hsa-miR-3128
TCTGGCAAGTAAAAAACTCTCAT
>hsa-miR-548ae-3p
CAAAAACTGCAATTACTTTCA
>hsa-miR-6721-5p
TGGGCAGGGGCTTATTGTAGGAG
>hsa-miR-4737
ATGCGAGGATGCTGACAGTG
>hsa-miR-3149
TTTGTATGGATATGTGTGTGTAT
>hsa-miR-676-3p
CTGTCCTAAGGTTGTTGAGTT
>hsa-miR-6838-3p
AAGTCCTGCTTCTGTTGCAG
>hsa-miR-6086
GGAGGTTGGGAAGGGCAGAG
>hsa-miR-376a-3p
ATCATAGAGGAAAATCCACGT
>hsa-miR-493-3p
TGAAGGTCTACTGTGTGCCAGG
>hsa-miR-6715a-3p
CCAAACCAGTCGTGCCTGTGG
>hsa-miR-4271
GGGGGAAGAAAAGGTGGGG
>hsa-miR-6726-5p
CGGGAGCTGGGGTCTGCAGGT
>hsa-miR-548bc
AAAAACTGTGATTACTTTTGC
>hsa-miR-550b-3p
TCTTACTCCCTCAGGCACTG
>hsa-miR-6769a-5p
AGGTGGGTATGGAGGAGCCCT
>hsa-miR-558
TGAGCTGCTGTACCAAAAT
>hsa-miR-520g-3p
ACAAAGTGCTTCCCTTTAGAGTGT
>hsa-miR-125b-5p
```

TCCCTGAGACCCTAACTTGTGA
>hsa-miR-6867-5p
TGTGTGTGTAGAGGAAGAAGGGA
>hsa-miR-6510-5p
CAGCAGGGGAGAGAGAGGAGTC
>hsa-miR-4721
TGAGGGCTCCAGGTGACGGTGG
>hsa-miR-5007-3p
ATCATATGAACCAAACTCTAAT
>hsa-miR-181d-5p
AACATTCATTGTTGTCGGTGGGT
>hsa-miR-4312
GGCCTTGTTCCTGTCCCCA
>hsa-miR-208b-3p
ATAAGACGAACAAAAGGTTTGT
>hsa-miR-1193
GGGATGGTAGACCGGTGACGTGC
>hsa-miR-660-3p
ACCTCCTGTGTGCATGGATTA
>hsa-miR-142-3p
TGTAGTGTTTCCTACTTTATGGA
>hsa-miR-580-3p
TTGAGAATGATGAATCATTAGG
>hsa-miR-6806-5p
TGTAGGCATGAGGCAGGGCCCAGG
>hsa-miR-2052
TGTTTTGATAACAGTAATGT
>hsa-miR-6881-3p
ATCCTCTTTCGTCCTTCCCACT
>hsa-miR-335-3p
TTTTTCATTATTGCTCCTGACC
>hsa-miR-7151-3p
CTACAGGCTGGAATGGGCTCA
>hsa-miR-3199
AGGGACTGCCTTAGGAGAAAGTT
>hsa-miR-6777-3p
TCCACTCTCCTGGCCCCCAG
>hsa-miR-548i
AAAAGTAATTGCGGATTTTGCC
>hsa-miR-4273
GTGTTCTCTGATGGACAG
>hsa-miR-4489
TGGGGCTAGTGATGCAGGACG
>hsa-miR-6842-5p
TGGGGGTGGTCTCTAGCCAAGG
>hsa-miR-6759-3p
TGACCTTTGCCTCTCCCCTCAG
>hsa-miR-1207-5p
TGGCAGGGAGGCTGGGAGGGG
>hsa-miR-614
GAACGCCTGTTCTTGCCAGGTGG
>hsa-miR-20b-3p
ACTGTAGTATGGGCACTTCCAG
>hsa-miR-759
GCAGAGTGCAAACAATTTTGAC
>hsa-miR-3529-3p
AACAACAAAATCACTAGTCTTCCA
>hsa-miR-1285-3p
TCTGGGCAACAAAGTGAGACCT
>hsa-miR-10395-5p
GTGATGGAGAGCAATACC
>hsa-miR-362-3p

AACACACCTATTCAAGGATTCA
>hsa-miR-6834-3p
TATGTCCCATCCCTCCATCA
>hsa-miR-4283
TGGGGCTCAGCGAGTTT
>hsa-miR-944
AAATTATTGTACATCGGATGAG
>hsa-miR-643
ACTTGTATGCTAGCTCAGGTAG
>hsa-miR-3179
AGAAGGGGTGAAATTTAAACGT
>hsa-miR-5590-3p
AATAAAGTTCATGTATGGCAA
>hsa-miR-4268
GGCTCCTCCTCTCAGGATGTG
>hsa-miR-3164
TGTGACTTTAAGGGAAATGGCG
>hsa-miR-4713-5p
TTCTCCCACTACCAGGCTCCCA
>hsa-miR-1224-5p
GTGAGGACTCGGGAGGTGG
>hsa-miR-648
AAGTGTGCAGGGCACTGGT
>hsa-miR-378b
ACTGGACTTGGAGGCAGAA
>hsa-miR-548ay-5p
AAAAGTAATTGTGGTTTTTGC
>hsa-miR-548ay-3p
CAAAACCGCGATTACTCTTGCA
>hsa-miR-424-3p
CAAAACGTGAGGCGCTGCTAT
>hsa-miR-490-3p
CAACCTGGAGGACTCCATGCTG
>hsa-miR-26b-3p
CCTGTTCTCCATTACTTGGCT
>hsa-miR-4804-5p
TTGGACGGTAAGGTTAAGCAA
>hsa-miR-1229-3p
CTCTCACCACTGCCCTCCCACAG
>hsa-miR-3192-5p
TCTGGGAGGTTGTAGCAGTGGAA
>hsa-miR-4439
GTGACTGATACCTTGGAGGCAT
>hsa-miR-4691-3p
CCAGCCACGGACTGAGAGTGCAT
>hsa-miR-6812-5p
ATGGGGTGAGATGGGGAGGAGCAGC
>hsa-miR-553
AAAACGGTGAGATTTTGTTTT
>hsa-miR-205-5p
TCCTTCATTCCACCGGAGTCTG
>hsa-miR-1343-5p
TGGGGAGCGGCCCCCGGGTGGG
>hsa-miR-5589-3p
TGCACATGGCAACCTAGCTCCCA
>hsa-miR-106a-5p
AAAAGTGCTTACAGTGCAGGTAG
>hsa-miR-4739
AAGGGAGGAGGAGCGGAGGGGCCCT
>hsa-miR-5585-5p
TGAAGTACCAGCTACTCGAGAG
>hsa-miR-376b-5p

CGTGGATATTCCTTCTATGTTT
>hsa-miR-26a-5p
TTCAAGTAATCCAGGATAGGCT
>hsa-miR-4704-5p
GACACTAGGCATGTGAGTGATT
>hsa-miR-4652-5p
AGGGGACTGGTTAATAGAACTA
>hsa-miR-10396b-5p
CGGCGGGGCTCGGAGCCGGG
>hsa-miR-613
AGGAATGTTCCTTCTTTGCC
>hsa-miR-551b-3p
GCGACCCATACTTGGTTTCAG
>hsa-miR-4800-3p
CATCCGTCCGTCTGTCCAC
>hsa-miR-5087
GGGTTTGTAGCTTTGCTGGCATG
>hsa-miR-4728-5p
TGGGAGGGGAGAGGCAGCAAGCA
>hsa-miR-4433b-5p
ATGTCCCACCCCCACTCCTGT
>hsa-miR-18a-5p
TAAGGTGCATCTAGTGCAGATAG
>hsa-miR-3917
GCTCGGACTGAGCAGGTGGG
>hsa-miR-3122
GTTGGGACAAGAGGACGGTCTT
>hsa-miR-5581-5p
AGCCTTCCAGGAGAAATGGAGA
>hsa-miR-1276
TAAAGAGCCCTGTGGAGACA
>hsa-miR-8078
GGTCTAGGCCCGGTGAGAGACTC
>hsa-miR-3065-3p
TCAGCACCAGGATATTGTTGGAG
>hsa-miR-5571-3p
GTCCTAGGAGGCTCCTCTG
>hsa-miR-520b-5p|hsa-miR-519a-2-5p
CCTCTACAGGGAAGCGCTTTC
>hsa-miR-3928-5p
TGAAGCTCTAAGGTTCCGCCTGC
>hsa-miR-3132
TGGGTAGAGAAGGAGCTCAGAGGA
>hsa-miR-4648
TGTGGGACTGCAAATGGGAG
>hsa-miR-5591-5p
TGGGAGCTAAGCTATGGGTAT
>hsa-miR-941
CACCCGGCTGTGTGCACATGTGC
>hsa-miR-3649
AGGGACCTGAGTGTCTAAG
>hsa-miR-24-2-5p
TGCCTACTGAGCTGAAACACAG
>hsa-miR-548al
AACGGCAATGACTTTTGTACCA
>hsa-miR-8089
CCTGGGGACAGGGGATTGGGGCAG
>hsa-miR-138-2-3p
GCTATTTCACGACACCAGGGTT
>hsa-miR-526b-3p
GAAAGTGCTTCCTTTTAGAGGC
>hsa-miR-519e-5p

TTCTCCAAAAGGGAGCACTTTC
>hsa-miR-5739
GCGGAGAGAGAATGGGGAGC
>hsa-miR-7156-3p
CTGCAGCCACTTGGGGAACTGGT
>hsa-miR-6794-3p
CTCACTCTCAGTCCCTCCCT
>hsa-miR-612
GCTGGGCAGGGCTTCTGAGCTCCTT
>hsa-miR-601
TGGTCTAGGATTGTTGGAGGAG
>hsa-miR-211-3p
GCAGGGACAGCAAAGGGGTGC
>hsa-miR-4464
AAGGTTTGGATAGATGCAATA
>hsa-miR-4456
CCTGGTGGCTTCCTTTT
>hsa-miR-6835-5p
AGGGGGTAGAAAGTGGCTGAAG
>hsa-miR-517c-3p
ATCGTGCATCCTTTTAGAGTGT
>hsa-miR-12135
TAAAGGTTTGTTTGTAAA
>hsa-miR-489-5p
GGTCGTATGTGTGACGCCATTT
>hsa-miR-3927-5p
GCCTATCACATATCTGCCTGT
>hsa-miR-1185-1-3p
ATATACAGGGGGAGACTCTTAT
>hsa-miR-6767-3p
CCACGTGCTTCTCTTTCCGCAG
>hsa-miR-4753-3p
TTCTCTTTCTTTAGCCTTGTGT
>hsa-miR-548q
GCTGGTGCAAAAGTAATGGCGG
>hsa-miR-4686
TATCTGCTGGGCTTTCTGGTGTT
>hsa-miR-6784-3p
TCTCACCCCAACTCTGCCCCAG
>hsa-miR-3924
ATATGTATATGTGACTGCTACT
>hsa-miR-7843-5p
GAGGGCAGAGCCAGCTTCCTGA
>hsa-miR-9985
TTCACAGTGGCTAAGCTAT
>hsa-miR-3648
AGCCGCGGGGATCGCCGAGGG
>hsa-miR-6078
CCGCCTGAGCTAGCTGTGG
>hsa-miR-874-5p
CGGCCCCACGCACCAGGGTAAGA
>hsa-miR-520h
ACAAAGTGCTTCCCTTTAGAGT
>hsa-miR-3912-3p
TAACGCATAATATGGACATGT
>hsa-miR-1973
ACCGTGCAAAGGTAGCATA
>hsa-miR-889-3p
TTAATATCGGACAACCATTGT
>hsa-miR-6132
AGCAGGGCTGGGGATTGCA
>hsa-miR-383-3p

ACAGCACTGCCTGGTCAGA
>hsa-miR-302b-3p
TAAGTGCTTCCATGTTTTAGTAG
>hsa-miR-487b-3p
AATCGTACAGGGTCATCCACTT
>hsa-let-7a-5p
TGAGGTAGTAGGTTGTATAGTT
>hsa-miR-135b-3p
ATGTAGGGCTAAAAGCCATGGG
>hsa-miR-548bb-3p
CAAAAACCATAGTTACTTTTGC
>hsa-miR-6879-3p
TGTCACCCGCTCCTTGCCCAG
>hsa-miR-6749-3p
CTCCTCCCCTGCCTGGCCCAG
>hsa-miR-25-5p
AGGCGGAGACTTGGGCAATTG
>hsa-miR-12131
TTTGGAGAGGTGTACTCCCA
>hsa-miR-559
TAAAGTAAATATGCACCAAAA
>hsa-miR-6743-3p
AGCCGCTCTTCTCCCTGCCCACA
>hsa-miR-6830-5p
CCAAGGAAGGAGGCTGGACATC
>hsa-miR-6824-5p
GTAGGGGAGGTTGGGCCAGGGA
>hsa-miR-5685
ACAGCCCAGCAGTTATCACGGG
>hsa-let-7b-5p
TGAGGTAGTAGGTTGTGTGGTT
>hsa-miR-6816-5p
TGGGGCGGGGCAGGTCCCTGC
>hsa-miR-298
AGCAGAAGCAGGGAGGTTCTCCCA
>hsa-miR-127-3p
TCGGATCCGTCTGAGCTTGGCT
>hsa-miR-770-5p
TCCAGTACCACGTGTCAGGGCCA
>hsa-miR-4539
GCTGAACTGGGCTGAGCTGGGC
>hsa-miR-634
AACCAGCACCCCAACTTTGGAC
>hsa-miR-4783-5p
GGCGCGCCCAGCTCCCGGGCT
>hsa-miR-6833-3p
TTTCTCTCTCCACTTCCTCAG
>hsa-miR-6760-3p
ACACTGTCCCCTTCTCCCCAG
>hsa-miR-3156-3p
CTCCCACTTCCAGATCTTTCT
>hsa-miR-132-3p
TAACAGTCTACAGCCATGGTCG
>hsa-miR-4708-3p
AGCAAGGCGGCATCTCTCTGAT
>hsa-miR-1915-3p
CCCCAGGGCGACGCGGCGGG
>hsa-miR-519c-3p
AAAGTGCATCTTTTTAGAGGAT
>hsa-miR-1273h-5p
CTGGGAGGTCAAGGCTGCAGT
>hsa-miR-197-3p

TTCACCACCTTCTCCACCCAGC
>hsa-miR-4308
TCCCTGGAGTTTCTTCTT
>hsa-miR-4529-3p
ATTGGACTGCTGATGGCCCGT
>hsa-miR-7844-5p
AAAACTAGGACTGTGTGGTGTA
>hsa-miR-361-3p
TCCCCCAGGTGTGATTCTGATTT
>hsa-miR-9901
CGGTCGCCGCGGTTCGCCGCC
>hsa-miR-1284
TCTATACAGACCCTGGCTTTTC
>hsa-miR-323a-3p
CACATTACACGGTCGACCTCT
>hsa-miR-3156-5p
AAAGATCTGGAAGTGGGAGACA
>hsa-miR-3926
TGGCCAAAAAGCAGGCAGAGA
>hsa-miR-548d-5p
AAAAGTAATTGTGGTTTTTGCC
>hsa-miR-4668-3p
GAAAATCCTTTTTGTTTTTCCAG
>hsa-miR-6767-5p
TCGCAGACAGGGACACATGGAGA
>hsa-miR-3198
GTGGAGTCCTGGGGAATGGAGA
>hsa-miR-3659
TGAGTGTTGTCTACGAGGGCA
>hsa-miR-130b-3p
CAGTGCAATGATGAAAGGGCAT
>hsa-miR-6853-5p
AGCGTGGGATGTCCATGAAGTCAG
>hsa-miR-100-3p
CAAGCTTGTATCTATAGGTATG
>hsa-miR-338-3p
TCCAGCATCAGTGATTTTGTTG
>hsa-miR-4671-3p
TTAGTGCATAGTCTTTGGTCT
>hsa-miR-6715b-5p
ACAGGCACGACTGGTTTGGCA
>hsa-miR-2392
TAGGATGGGGGTGAGAGGTG
>hsa-miR-718
CTTCCGCCCCGCCGGGCGTCG
>hsa-miR-410-3p
AATATAACACAGATGGCCTGT
>hsa-miR-3126-3p
CATCTGGCATCCGTCACACAGA
>hsa-miR-4716-3p
AAGGGGGAAGGAAACATGGAGA
>hsa-miR-17-5p
CAAAGTGCTTACAGTGCAGGTAG
>hsa-miR-942-3p
CACATGGCCGAAACAGAGAAGT
>hsa-miR-6865-5p
TAGGTGGCAGAGGAGGGACTTCA
>hsa-miR-4684-3p
TGTTGCAAGTCGGTGGAGACGT
>hsa-miR-548ad-3p
GAAAACGACAATGACTTTTGCA
>hsa-miR-589-3p

TCAGAACAAATGCCGGTTCCCAGA
>hsa-miR-3927-3p
CAGGTAGATATTTGATAGGCAT
>hsa-miR-802
CAGTAACAAAGATTCATCCTTGT
>hsa-miR-4742-5p
TCAGGCAAAGGGATATTTACAGA
>hsa-miR-4665-3p
CTCGGCCGCGGCGCGTAGCCCCCGCC
>hsa-miR-5681b
AGGTATTGCCACCCTTTCTAGT
>hsa-miR-7853-5p
TCAAATGCAGATCCTGACTTC
>hsa-miR-19b-3p
TGTGCAAATCCATGCAAAACTGA
>hsa-miR-590-5p
GAGCTTATTCATAAAAGTGCAG
>hsa-miR-5090
CCGGGGCAGATTGGTGTAGGGTG
>hsa-miR-5589-5p
GGCTGGGTGCTCTTGTGCAGT
>hsa-miR-7849-3p
GACAATTGTTGATCTTGGGCCT
>hsa-miR-3688-3p
TATGGAAAGACTTTGCCACTCT
>hsa-miR-3136-3p
TGGCCCAACCTATTCAGTTAGT
>hsa-miR-11400
TCGGCTGTGTATCTCTGTGTC
>hsa-miR-1228-5p
GTGGGCGGGGGCAGGTGTGTG
>hsa-miR-591
AGACCATGGGTTCTCATTGT
>hsa-miR-6751-5p
TTGGGGGTGAGGTTGGTGTCTGG
>hsa-miR-6822-3p
AGGCTCTAACTGGCTTTCCCTGCA
>hsa-miR-526a-3p
GAAAGCGCTTCCTTTTAGAGGA
>hsa-miR-10397-3p
CATAGATCTCGTCGCTTACTGGGA
>hsa-miR-519d-5p
CCTCCAAAGGGAAGCGCTTTCTGTT
>hsa-miR-656-3p
AATATTATACAGTCAACCTCT
>hsa-miR-627-3p
TCTTTTCTTTGAGACTCACT
>hsa-miR-3619-5p
TCAGCAGGCAGGCTGGTGCAGC
>hsa-miR-3190-5p
TCTGGCCAGCTACGTCCCCA
>hsa-miR-3911
TGTGTGGATCCTGGAGGAGGCA
>hsa-miR-6731-5p
TGGGAGAGCAGGGTATTGTGGA
>hsa-miR-299-3p
TATGTGGGATGGTAAACCGCTT
>hsa-miR-4668-5p
AGGGAAAAAAAAAAGGATTTGTC
>hsa-miR-636
TGTGCTTGCTCGTCCCGCCCGCA
>hsa-miR-4269

GCAGGCACAGACAGCCCTGGC
>hsa-miR-4667-3p
TCCCTCCTTCTGTCCCCACAG
>hsa-miR-588
TTGGCCACAATGGGTTAGAAC
>hsa-miR-6788-3p
TTCGCCACTTCCCTCCCTGCAG
>hsa-miR-6852-5p
CCCTGGGGTTCTGAGGACATG
>hsa-miR-3938
AATTCCCTTGTAGATAACCCGG
>hsa-miR-579-5p
TCGCGGTTTGTGCCAGATGACG
>hsa-miR-5010-3p
TTTTGTGTCTCCCATTCCCCAG
>hsa-miR-584-3p
TCAGTTCCAGGCCAACCAGGCT
>hsa-miR-302c-5p
TTTAACATGGGGGTACCTGCTG
>hsa-miR-3689a-3p
CTGGGAGGTGTGATATCGTGGT
>hsa-miR-548c-3p
CAAAAATCTCAATTACTTTTGC
>hsa-miR-125b-2-3p
TCACAAGTCAGGCTCTTGGGAC
>hsa-miR-656-5p
AGGTTGCCTGTGAGGTGTTCA
>hsa-miR-105-5p
TCAAATGCTCAGACTCCTGTGGT
>hsa-miR-1178-3p
TTGCTCACTGTTCTTCCCTAG
>hsa-miR-1271-5p
CTTGGCACCTAGCAAGCACTCA
>hsa-miR-5584-5p
CAGGGAAATGGGAAGAACTAGA
>hsa-miR-5692b
AATAATATCACAGTAGGTGT
>hsa-miR-3195
CGCGCCGGGCCCGGGTT
>hsa-miR-3162-5p
TTAGGGAGTAGAAGGGTGGGGAG
>hsa-miR-3196
CGGGGCGGCAGGGGCCTC
>hsa-miR-4751
AGAGGACCCGTAGCTGCTAGAAGG
>hsa-miR-3619-3p
GGGACCATCCTGCCTGCTGTGG
>hsa-miR-6756-5p
AGGGTGGGGCTGGAGGTGGGGCT
>hsa-miR-548d-3p
CAAAAACCACAGTTTCTTTTGC
>hsa-miR-4796-5p
TGTCTATACTCTGTCACTTTAC
>hsa-miR-4476
CAGGAAGGATTTAGGGACAGGC
>hsa-miR-6885-5p
AGGGGGGCACTGCGCAAGCAAAGCC
>hsa-miR-200c-5p
CGTCTTACCCAGCAGTGTTTGG
>hsa-miR-194-5p
TGTAACAGCAACTCCATGTGGA
>hsa-miR-4292

CCCCTGGGCCGGCCTTGG
>hsa-miR-410-5p
AGGTTGTCTGTGATGAGTTCG
>hsa-miR-520c-3p
AAAGTGCTTCCTTTTAGAGGGT
>hsa-miR-8065
TGTAGGAACAGTTGAATTTTGGCT
>hsa-miR-3944-3p
TTCGGGCTGGCCTGCTGCTCCGG
>hsa-miR-6750-3p
GAACTCACCCTCTGCTCCCAG
>hsa-miR-4800-5p
AGTGGACCGAGGAAGGAAGGA
>hsa-miR-6513-5p
TTTGGGATTGACGCCACATGTCT
>hsa-miR-98-3p
CTATACAACTTACTACTTTCCC
>hsa-miR-4778-5p
AATTCTGTAAAGGAAGAAGAGG
>hsa-miR-3184-3p
AAAGTCTCGCTCTCTGCCCCTCA
>hsa-miR-1224-3p
CCCCACCTCCTCTCTCCTCAG
>hsa-miR-6841-3p
ACCTTGCATCTGCATCCCCAG
>hsa-miR-6728-5p
TTGGGATGGTAGGACCAGAGGGG
>hsa-miR-5100
TTCAGATCCCAGCGGTGCCTCT
>hsa-miR-3130-3p
GCTGCACCGGAGACTGGGTAA
>hsa-miR-4287
TCTCCCTTGAGGGCACTTT
>hsa-miR-6511b-3p
CCTCACCACCCCTTCTGCCTGCA
>hsa-miR-1251-3p
CGCTTTGCTCAGCCAGTGTAG
>hsa-miR-6504-5p
TCTGGCTGTGCTGTAATGCAG
>hsa-miR-3180-5p
CTTCCAGACGCTCCGCCCCACGTCG
>hsa-miR-431-5p
TGTCTTGCAGGCCGTCATGCA
>hsa-miR-10b-5p
TACCCTGTAGAACCGAATTTGTG
>hsa-miR-518d-3p
CAAAGCGCTTCCCTTTGGAGC
>hsa-miR-5708
ATGAGCGACTGTGCCTGACC
>hsa-miR-7157-5p
TCAGCATTCATTGGCACCAGAGA
>hsa-miR-4641
TGCCCATGCCATACTTTTGCCTCA
>hsa-miR-154-3p
AATCATACACGGTTGACCTATT
>hsa-miR-7155-5p
TCTGGGGTCTTGGGCCATC
>hsa-miR-6761-3p
TCCTACGCTGCTCTCTCACTCC
>cali_37_rc|Ome_cali|artificial
TTCCGCTTTACGGGTTAATAGA
>cali_23_rc|Ome_cali|artificial

```
CTCGTCTCCGGCTGTATATACC
>cali_25_rc|alt_cali|artificial
AGGGCCCTTTAGGCACTAATAG
>cali_19_rc|Ome_cali|artificial
ATGATTCTCTAACGTCGGCATT
>cali_01_rc|alt_cali|artificial
TCCACGACGTCTCATGTATTTC
>cali_24_rc|alt_cali|artificial
TGCTACTCCGATCTTTAGCCTC
>cali_17_rc|alt_cali|artificial
TCATGAGTCCGTACCTTGATTG
>cali_18_rc|alt_cali|artificial
ATCATTTACGATTCGGAGCTGT
>cali_05_rc|Ome_cali|artificial
CATGGTTGTAAGTCCCGGTAAT
>cali_44_rc|alt_cali|artificial
AGCCGCATTTCGTAGTGATATT
>cali_38_rc|Ome_cali|artificial
GGAGGATACTTAATCCGCTGTG
>cali_08_rc|Ome_cali|artificial
GGATTACTCGGGTTTGAGACAG
>cali_04_rc|alt_cali|artificial
GGGTACCATACCGGTTGTCTTA
>cali_41_rc|Ome_cali|artificial
TGCTAGTCCACGGGAGAAATAT
>cali_40_rc|Ome_cali|artificial
GGAGCCGTGAATACAATCCTAG
>cali_39_rc|Ome_cali|artificial
GGTAATCCAACGTTGATGGTTT
>cali_43_rc|alt_cali|artificial
TCTAGTTGCGTGATGGAGAGAA
>cali_29_rc|Ome_cali|artificial
CGTCGATTTAGACCGTATAGCC
>cali_27_rc|alt_cali|artificial
GTAGCTGTCAGTACGTTCGTGC
>p3adapter|adapter|artificial
TCGTATGCCGTCTTCTGCTTG
>p5adapter|adapter|artificial
GTTCAGAGTTCTACAGTCCGACGATC
>cali_20_rc|alt_cali|artificial
GATAGTTCGGGATCGCTGTAAC
>hiv1-miR-N367
ACTGACCTTTGGATGGTGCTTCAA
>hiv1-miR-TAR-5p
TCTCTCTGGTTAGACCAGATCTGA
>hiv1-miR-TAR-3p
TCTCTGGCTAACTAGGGAACCCA
>hiv1-miR-H1
CCAGGGAGGCGTGCCTGGGC
>cali_37_rc|Ome_cali|artificial
TTCCGCTTTACGGGTTAATAGA
>cali_23_rc|Ome_cali|artificial
CTCGTCTCCGGCTGTATATACC
>cali_25_rc|alt_cali|artificial
AGGGCCCTTTAGGCACTAATAG
>cali_19_rc|Ome_cali|artificial
ATGATTCTCTAACGTCGGCATT
>cali_01_rc|alt_cali|artificial
TCCACGACGTCTCATGTATTTC
>cali_24_rc|alt_cali|artificial
TGCTACTCCGATCTTTAGCCTC
>cali_17_rc|alt_cali|artificial
```

```
TCATGAGTCCGTACCTTGATTG
>cali_18_rc|alt_cali|artificial
ATCATTTACGATTCGGAGCTGT
>cali_05_rc|Ome_cali|artificial
CATGGTTGTAAGTCCCGGTAAT
>cali_44_rc|alt_cali|artificial
AGCCGCATTTCGTAGTGATATT
>cali_38_rc|Ome_cali|artificial
GGAGGATACTTAATCCGCTGTG
>cali_08_rc|Ome_cali|artificial
GGATTACTCGGGTTTGAGACAG
>cali_04_rc|alt_cali|artificial
GGGTACCATACCGGTTGTCTTA
>cali_41_rc|Ome_cali|artificial
TGCTAGTCCACGGGAGAAATAT
>cali_40_rc|Ome_cali|artificial
GGAGCCGTGAATACAATCCTAG
>cali_39_rc|Ome_cali|artificial
GGTAATCCAACGTTGATGGTTT
>cali_43_rc|alt_cali|artificial
TCTAGTTGCGTGATGGAGAGAA
>cali_29_rc|Ome_cali|artificial
CGTCGATTTAGACCGTATAGCC
>cali_27_rc|alt_cali|artificial
GTAGCTGTCAGTACGTTCGTGC
>p3adapter|adapter|artificial
TCGTATGCCGTCTTCTGCTTG
>p5adapter|adapter|artificial
GTTCAGAGTTCTACAGTCCGACGATC
>cali_20_rc|alt_cali|artificial
GATAGTTCGGGATCGCTGTAAC
```
