## Supplementary material for "SPOROS: A pipeline to analyze DISE/6mer seed toxicity": Table S3

**Table S3: Murine mature miRNAs**

```
>mmu-miR-12187-3p
TTCATTCACCTCTGCTCTTGC
>mmu-miR-6994-3p
AACGATCTTCTCCGTCTTTGC
>mmu-miR-6341
CAGTGCAATGATATTGTCACTAT
>mmu-miR-744-3p
CTGTTGCCACTAACCTCAACCT
>mmu-miR-106b-5p
TAAAGTGCTGACAGTGCAGAT
>mmu-miR-3066-5p
TTGGTTGCTGTAGATTAAGTAG
>mmu-miR-5104
CTGTGCTAGTGAGGTGGCTCAGCA
>mmu-miR-669p-3p
CATAACATACACACACACACGTAT
>mmu-miR-3106-5p
TGGCTCATTTAGAAGCAGCCA
>mmu-miR-323-5p
AGGTGGTCCGTGGCGCGTTCGC
>mmu-miR-5625-3p
CTGATTCAAGTGGTCTCTCGTGTCC
>mmu-miR-92a-1-5p
AGGTTGGGATTTGTCGCAATGCT
>mmu-miR-10b-3p
CAGATTCGATTCTAGGGGAATA
>mmu-miR-5618-5p
TACCCTTTACACCGTGTAGT
>mmu-miR-190a-3p
ACTATATATCAAGCATATTCCT
>mmu-miR-7066-3p
TCTACCCATTGCCTGCCTCCCAG
>mmu-miR-6935-5p
TGTGGGAGGCTCAGGGCCCCTG
>mmu-miR-342-3p
TCTCACACAGAAATCGCACCCGT
>mmu-miR-3969
CCCTAAAGTAGAAATCACTA
>mmu-miR-200a-3p
TAACACTGTCTGGTAACGATGT
>mmu-miR-3065-5p
TCAACAAAATCACTGATGCTGG
>mmu-miR-582-3p
TAACCTGTTGAACAACTGAAC
>mmu-miR-409-3p
GAATGTTGCTCGGTGAACCCCT
>mmu-miR-155-5p
TTAATGCTAATTGTGATAGGGGT
>mmu-miR-453
AGGTTGCCTCATAGTGAGCTTGCA
>mmu-miR-383-3p
CCACAGCACTGCCTGGTCAGA
>mmu-miR-5122
CCGCGGGACCCGGGGCTGTG
>mmu-miR-146b-3p
GCCCTAGGGACTCAGTTCTGGT
>mmu-miR-137-3p
TTATTGCTTAAGAATACGCGTAG
>mmu-miR-99a-5p
```

AACCCGTAGATCCGATCTTGTG
>mmu-miR-3110-3p
GCACTCCATCGGAGGCAGACAC
>mmu-miR-486a-3p
CGGGGCAGCTCAGTACAGGAT
>mmu-miR-143-3p
TGAGATGAAGCACTGTAGCTC
>mmu-miR-6358
CAGTGTGGCAGTATACAGCTATA
>mmu-miR-412-3p
TTCACCTGGTCCACTAGCCG
>mmu-miR-6972-5p
GCTGGAGGCACAGGGCAGAACAG
>mmu-miR-126b-5p
ATTATTACTCACGGTACGAGTT
>mmu-miR-6901-5p
TGCAGGAATGCAGAGGACCT
>mmu-miR-1894-3p
GCAAGGGAGAGGGTGAAGGGAG
>mmu-miR-1905
CACCAGTCCCACCACGCGGTAG
>mmu-miR-6402
GAGCAGTTTTCCCAGGAACCCGC
>mmu-miR-6898-5p
TGTAAGGGGAGATGCAGGGAGC
>mmu-miR-7006-5p
TGGGGGTGTTCAAGACTTGGGAACC
>mmu-miR-297b-5p
ATGTATGTGTGCATGAACATGT
>mmu-miR-7238-3p
CTGTCCTTTGTGTTC
>mmu-miR-1258-5p
TGCTGAGCTAATTCCCTAACTG
>mmu-miR-7217-3p
TGAGAACCCACAAGCAAGAAGA
>mmu-miR-3097-5p
CACAGGTGGGAAGTGTGTGTCCA
>mmu-miR-467h
ATAAGTGTGTGCATGTATATGT
>mmu-miR-377-3p
ATCACACAAAGGCAACTTTTGT
>mmu-miR-1271-3p
AGTGCCTACTGTGTGCCAAGACA
>mmu-miR-6993-5p
AGTGGGAGAAACGGGTGTCTGTC
>mmu-miR-6964-3p
TTTCTTGTCTTCCACTCTAG
>mmu-miR-5129-3p
AATGTGCCTGTGCATCTCTTCC
>mmu-miR-8110
AAGCGTGGATTGGGGGGGGG
>mmu-miR-7023-5p
ATGGGGGAGCTGGGATTGGGCGT
>mmu-miR-376b-5p
GTGGATATTCCTTCTATGGTTA
>mmu-miR-147-3p
GTGTGCGGAAATGCTTCTGCTA
>mmu-miR-29b-2-5p
CTGGTTTCACATGGTGGCTTAGATT
>mmu-miR-491-3p
CTTATGCAAGATTCCCTTCTAC
>mmu-miR-3102-3p.2-3p

CTCTACTCCCTGCCCCAGCCA
>mmu-miR-3082-5p
GACAGAGTGTGTGTGTCTGTGT
>mmu-miR-208a-3p
ATAAGACGAGCAAAAAGCTTGT
>mmu-miR-7020-5p
TGGGATGGTGGAGAGGGTGACCAG
>mmu-miR-132-5p
AACCGTGGCTTTCGATTGTTAC
>mmu-miR-6934-3p
ACCTCTGCTCCTGCCCCACCAG
>mmu-miR-345-5p
GCTGACCCCTAGTCCAGTGCTT
>mmu-miR-329-3p
AACACACCCAGCTAACCTTTTT
>mmu-miR-99b-3p
CAAGCTCGTGTCTGTGGGTCCG
>mmu-miR-3099-3p
TAGGCTAGAGAGAGGTTGGGGA
>mmu-miR-880-3p
TACTCCATCCTCTCTGAGTAGA
>mmu-miR-291b-3p
AAAGTGCATCCATTTTGTTTGT
>mmu-miR-19a-5p
TAGTTTTGCATAGTTGCACTAC
>mmu-miR-21a-3p
CAACAGCAGTCGATGGGCTGTC
>mmu-miR-26b-3p
CCTGTTCTCCATTACTTGGCTC
>mmu-miR-190b-5p
TGATATGTTTGATATTGGGTTG
>mmu-miR-30e-3p
CTTTCAGTCGGATGTTTACAGC
>mmu-miR-101c
ACAGTACTGTGATAACTGA
>mmu-miR-103-3p
AGCAGCATTGTACAGGGCTATGA
>mmu-miR-1298-5p
TTCATTCGGCTGTCCAGATGTA
>mmu-miR-7060-5p
GTGAGTGCTGGGTAGAATGGGA
>mmu-miR-344d-2-5p
AGTCTGGTTGCTGGCTATATTCCA
>mmu-miR-7077-3p
CCTTCCATGGCTCCTGGCAG
>mmu-miR-451b
TGGGAGCAGCAAGAGAACCGT
>mmu-miR-6896-3p
TTTCTCTCTCTCACCTACAAAC
>mmu-miR-429-3p
TAATACTGTCTGGTAATGCCGT
>mmu-miR-411-3p
TATGTAACACGGTCCACTAACC
>mmu-miR-125a-3p
ACAGGTGAGGTTCTTGGGAGCC
>mmu-miR-31-3p
TGCTATGCCAACATATTGCCATC
>mmu-miR-468-5p
GACTGATGTACTGATAAGAAACTCAGT
>mmu-miR-7648-3p
AGGGCTGGGCCCGGGACGCGG
>mmu-miR-106b-3p

CCGCACTGTGGGTACTTGCTGC
>mmu-miR-30a-3p
CTTTCAGTCGGATGTTTGCAGC
>mmu-miR-540-5p
CAAGGGTCACCCTCTGACTCTGT
>mmu-miR-7059-5p
ACCGGGGACTGCGGCTTTCAGT
>mmu-miR-33-5p
GTGCATTGTAGTTGCATTGCA
>mmu-miR-3079-3p
CAGGCTCATCAGATGAAAGTC
>mmu-miR-5112
TAGCTCAGCGGGAGAGCAC
>mmu-miR-3073a-3p
TTGATGTCCACTGTGACCATAG
>mmu-miR-92b-3p
TATTGCACTCGTCCCGGCCTCC
>mmu-miR-708-5p
AAGGAGCTTACAATCTAGCTGGG
>mmu-miR-200a-5p
CATCTTACCGGACAGTGCTGGA
>mmu-miR-7225-5p
ACGTAGACTGTGTAGAAGCC
>mmu-miR-3088-3p
TTCATGAGCAGCTGCAAAGGTGT
>mmu-miR-466a-5p
TATGTGTGTGTACATGTACATA
>mmu-miR-6917-3p
GTCACTTCTCTTCCCACCACAG
>mmu-miR-710
CCAAGTCTTGGGGAGAGTTGAG
>mmu-miR-344c-3p
TGATCTAGTCAAAGCCTGACAGT
>mmu-miR-6955-5p
AGGGCCTCAGGACTCAGGAGGTGA
>mmu-miR-504-5p
AGACCCTGGTCTGCACTCTATC
>mmu-miR-3105-3p
ACTGCTTATGAGCTTGCACTCC
>mmu-miR-7651-5p
AAGTGGAAAGCCAGCCTGAGAT
>mmu-miR-149-5p
TCTGGCTCCGTGTCTTCACTCCC
>mmu-miR-489-3p
AATGACACCACATATATGGCAGC
>mmu-miR-9718
TTGCTGACCTGGGTGGTGGT
>mmu-miR-1843b-3p
CCGATCGTTCCCCTCCATAC
>mmu-miR-1906
TGCAGCAGCCTGAGGCAGGGCT
>mmu-miR-154-5p
TAGGTTATCCGTGTTGCCTTCG
>mmu-miR-301b-5p
GCTCTGACTAGGTTGCACTACT
>mmu-miR-125a-5p
TCCCTGAGACCCTTTAACCTGTGA
>mmu-miR-7662-5p
AGGGCTGAGGCCTGGATCCAGC
>mmu-miR-6965-5p
AGAGGGAGGCTATTGGGCAGGAGA
>mmu-miR-574-3p

CACGCTCATGCACACACCCACA
>mmu-miR-199a-3p|mmu-miR-199b-3p
ACAGTAGTCTGCACATTGGTTA
>mmu-miR-1911-5p
TGAGTACAGCCATGTCTGTTGGG
>mmu-miR-497b
CACCACAGTGTGGTTTGGACGTGG
>mmu-miR-6967-5p
CAGGGAGGGATAAAGACCAGT
>mmu-miR-1970c-5p
TGTGTCACTGGGGTTATGCTTTG
>mmu-let-7k
TGAGGTAGGAGGTTGTGTG
>mmu-miR-5101
TTTGTTTGTTTTGCTGATGCAG
>mmu-miR-340-5p
TTATAAAGCAATGAGACTGATT
>mmu-miR-134-5p
TGTGACTGGTTGACCAGAGGGG
>mmu-miR-688
TCGCAGGCGACTACTTATTC
>mmu-miR-568
ATGTATAAATGTATACACAC
>mmu-miR-1247-3p
CGGGAACGTCGAGACTGGAGC
>mmu-miR-431-3p
CAGGTCGTCTTGCAGGGCTTCT
>mmu-miR-219b-5p
AGATGTCCAGCCACAATTCTCG
>mmu-miR-434-3p
TTTGAACCATCACTCGACTCCT
>mmu-miR-3100-5p
TTGGGAACGGGGTGTCTTTGGGA
>mmu-miR-1900
GGCCGCCCTCTCTGGTCCTTCA
>mmu-miR-546
ATGGTGGCACGGAGTC
>mmu-miR-16-5p
TAGCAGCACGTAAATATTGGCG
>mmu-miR-7002-3p
TTGTGCTTCCCCTTGCCAG
>mmu-miR-1961
TGAGGTAGTAGTTAGAA
>mmu-miR-302a-5p
ACTTAAACGTGGTTGTACTTGC
>mmu-miR-7223-5p
TAGCTTTGCTGAGGTGTCGG
>mmu-miR-5617-3p
CAGGCGGCCTCAGCTCTCACT
>mmu-miR-700-5p
TAAGGCTCCTTCCTGTGCTTGC
>mmu-miR-6353
TAGCAGCACGTATTTATCCTATT
>mmu-miR-182-3p
GTGGTTCTAGACTTGCCAACT
>mmu-miR-5098
GTTACATGGTGAAGCCCAGTT
>mmu-miR-592-5p
ATTGTGTCAATATGCGATGATGT
>mmu-miR-7015-5p
TCTGTGCAGTAGGCTGTGGGTTC
>mmu-miR-7683-5p

TTCCGTGTTTCGTCTGACCACT
>mmu-miR-6937-5p
TAGCTGTAAGGGCTGGGTCTGTGT
>mmu-miR-6340
GTCAGCAGCAGCTTCGCTTTGGC
>mmu-miR-144-5p
GGATATCATCATATACTGTAAGT
>mmu-miR-12203-5p
TGCCGGAGGGATCGATTGCTT
>mmu-miR-191-3p
GCTGCACTTGGATTTCGTTCCC
>mmu-miR-532-3p
CCTCCCACACCCAAGGCTTGCA
>mmu-miR-6952-5p
ATGGTGGGCGGTGAAACAGAGA
>mmu-miR-30c-5p
TGTAAACATCCTACACTCTCAGC
>mmu-miR-182-5p
TTTGGCAATGGTAGAACTCACACCG
>mmu-miR-7024-3p
CCAGCAGTCCCTGCTCCCTACAG
>mmu-miR-150-3p
CTGGTACAGGCCTGGGGGATAG
>mmu-miR-6925-5p
TGAGGAAGAGAAGGCTTGGACT
>mmu-miR-423-3p
AGCTCGGTCTGAGGCCCCTCAGT
>mmu-miR-7674-3p
TCCCATTCTCACCGCACCCAG
>mmu-miR-7080-5p
GTAGGAGCTGGAGGTGGGTTTGG
>mmu-miR-3078-3p
TTGCTGGGGTAGTCTTTAGG
>mmu-miR-3572-5p
TGGGGAACAGGGCAAGGTGGAC
>mmu-miR-1897-5p
CTTTGGATGGAGAAAGAGGGGG
>mmu-miR-218-1-3p
AAACATGGTTCCGTCAAGCACC
>mmu-miR-7683-3p
TGGAAAGGTGGAACACGGAAC
>mmu-miR-100-5p
AACCCGTAGATCCGAACTTGTG
>mmu-miR-6967-3p
TCATCTTTATCTCTCCCCAG
>mmu-miR-511-3p
AATGTGTAGCAAAAGACAGGAT
>mmu-miR-6410
TGGCTCAGGGACTACTCTGTGC
>mmu-miR-466n-3p
TATACATGAGAGCATACATAGA
>mmu-miR-3084-5p
GTTGAAGGTTAATTAGCAGAGT
>mmu-miR-802-3p
ACGGAGAGTCTTTGTCACTCAG
>mmu-miR-450a-1-3p
ATTGGGAACATTTTGCATAAAT
>mmu-miR-465c-5p
TATTTAGAATGGCGCTGATCTG
>mmu-miR-743b-5p
TGTTCAGACTGGTGTCCATCA
>mmu-miR-496b

CAACATGGCCAATTCCTTTCAT
>mmu-miR-3473g
CAAAAGTGAGGCTGGGGAGA
>mmu-miR-3105-5p
AGAGCAAGCCCGTAAGCAGCGT
>mmu-miR-24-2-5p
GTGCCTACTGAGCTGAAACAGT
>mmu-miR-12181-3p
TCCAGGCCTCTTGCTCTCTCA
>mmu-miR-8119
GAGGAGAGGGAGCTAGGGTC
>mmu-miR-7660-5p
TTTGCCCTGGCTTCAGATCTCG
>mmu-miR-7041-3p
TGGTTCTCTTCTTCCCTCAG
>mmu-miR-1298-3p
CATCTGGGCAACTGATTGAACT
>mmu-miR-29a-5p
ACTGATTTCTTTTGGTGTTCAG
>mmu-miR-6896-5p
TTTGTAGGCGACGAGAACCCGT
>mmu-miR-542-3p
TGTGACAGATTGATAACTGAAA
>mmu-miR-12183-5p
GCTTCTTCTCTGTCCTGA
>mmu-miR-7067-3p
TGATTGCTTCCCTCCCCACAG
>mmu-miR-181c-5p
AACATTCAACCTGTCGGTGAGT
>mmu-miR-3072-3p
TGCCCCCTCCAGGAAGCCTTCT
>mmu-miR-148b-5p
GAAGTTCTGTTATACACTCAGGCT
>mmu-miR-7056-5p
TGTGGAGGAGGACAGAGAGGTT
>mmu-miR-7009-3p
TCTTTTCCCCTCTCCCTGCAG
>mmu-miR-6982-5p
CTGGAGGATCGCAGGGGTGGCCTGG
>mmu-miR-669h-5p
ATGCATGGGTGTATAGTTGAGTGC
>mmu-miR-23b-5p
GGGTTCCTGGCATGCTGATTT
>mmu-miR-5124a
GGTCCAGTGACTAAGAGCAT
>mmu-miR-1251-5p
ACTCTAGCTGCCAAAGGCGCT
>mmu-miR-3068-5p
TTGGAGTTCATGCAAGTTCTAACC
>mmu-miR-450a-2-3p
ATTGGGGATGCTTTGCATTCAT
>mmu-miR-7649-5p
TGACAGCAGTGCTGTAGACATG
>mmu-miR-496a-3p
TGAGTATTACATGGCCAATCTC
>mmu-miR-6947-5p
CATCCAGTGGAGAGGCTCGTG
>mmu-miR-299b-5p
GGTTTACCGTCCCACATACAT
>mmu-miR-344d-3p
GATATAACCACTGCCAGACTGA
>mmu-miR-598-5p

GCGGTGATGCCGATGGTGCGAGC
>mmu-miR-8097
AACAAGGAAAATGTCCGGGCTG
>mmu-miR-6945-5p
TCCTGGGAGGGGCAGCCCCTGG
>mmu-miR-6925-3p
TCCCCTTTTATCCCTCAG
>mmu-miR-291a-3p
AAAGTGCTTCCACTTTGTGTGC
>mmu-miR-7115-3p
ACTTGGTCCCCTGCCCCCACAG
>mmu-miR-7684-3p
TGCTGACTGGGGCTGGCCTGTG
>mmu-miR-7680-3p
ACTGCTTGTTCACTGGAATAGG
>mmu-miR-770-3p
CGTGGGCCTGACGTGGAGCTGG
>mmu-miR-6940-3p
TTACCTTCCGTGCTTGCCCGCAG
>mmu-miR-207
GCTTCTCCTGGCTCTCCTCCCTC
>mmu-miR-7087-3p
TGCTGGCTCTCTCCCCTGCCAG
>mmu-miR-6909-3p
TGCCTTCCCCGGCCTCCCACAG
>mmu-miR-6980-3p
TGACTTGCCTTCCACCCCAG
>mmu-miR-380-3p
TATGTAGTATGGTCCACATCTT
>mmu-miR-6973b-5p
GTAGGTATGGGGAGAGGACAGGGA
>mmu-miR-196a-5p
TAGGTAGTTTCATGTTGTTGGG
>mmu-miR-682
CTGCAGTCACAGTGAAGTCTG
>mmu-miR-7667-3p
AGGGACTGCAGAGATGGCACTG
>mmu-miR-452-5p
TGTTTGCAGAGGAAACTGAGAC
>mmu-miR-674-3p
CACAGCTCCCATCTCAGAACAA
>mmu-miR-221-3p
AGCTACATTGTCTGCTGGGTTTC
>mmu-miR-6987-5p
TGGGACAGGGTGACAGGGTGAG
>mmu-miR-6338
CTGGTTTCTCTCATTTGTGCT
>mmu-miR-7076-5p
AGGGGAGGTGGTATGGTCTCA
>mmu-miR-3081-3p
TTGCGCTCCGATCTCTGAGCTGG
>mmu-miR-670-3p
TTTCCTCATATCCATTCAGGAGTGT
>mmu-miR-7014-3p
CCTCTTTCATACCAACCACAG
>mmu-miR-126a-3p
TCGTACCGTGAGTAATAATGCG
>mmu-miR-3060-3p
CCATAGCACAGAAGCACTCCCA
>mmu-miR-7239-5p
GCCATCCTGACAAAGCCCTT
>mmu-miR-3473d

CCACTGAGCCACTTTCCAGCCCTT
>mmu-miR-675-3p
CTGTATGCCCTAACCGCTCAGT
>mmu-miR-290b-5p
GCTTAAAACTAGGCGGCACTTT
>mmu-miR-7218-5p
TGCAGGGTTTAGTGTAGAGGG
>mmu-miR-6981-5p
GTGAGGAGAAGGAAGAGGCTGAAGGC
>mmu-miR-6903-3p
TGTGTGTCTGTTTTTCCCAG
>mmu-miR-1968-5p
TGCAGCTGTTAAGGATGGTGGACT
>mmu-miR-195a-3p
CCAATATTGGCTGTGCTGCTCC
>mmu-miR-467c-3p
ATATACATACACACACCTATAC
>mmu-miR-743a-5p
TATTCAGATTGGTGCCTGTCAT
>mmu-miR-3084-3p
TTCTGCCAGTCTCCTTCAGAC
>mmu-miR-28a-3p
CACTAGATTGTGAGCTGCTGGA
>mmu-miR-7073-3p
ATAGCCTTTTCCTCCCCAACAG
>mmu-miR-1969
AAGATGGAGACTTTAACATGGGT
>mmu-miR-320-5p
GCCTTCTCTTCCCGGTTCTTCC
>mmu-miR-7647-5p
AACTACCACCTCCTGGCCTTTT
>mmu-miR-1981-5p
GTAAAGGCTGGGCTTAGACGTGGC
>mmu-miR-20b-3p
ACTGCAGTGTGAGCACTTCTAG
>mmu-miR-679-3p
AGCAAGGTCCTCCTCACAGTAG
>mmu-miR-344-5p
AGTCAGGCTCCTGGCTAGATTCCAGG
>mmu-miR-7050-3p
TCTCAGCCTTTATCTCCCCAG
>mmu-miR-130a-5p
GCTCTTTTCACATTGTGCTACT
>mmu-miR-7210-5p
TAACATTGTAGACAGGCACAAAG
>mmu-miR-425-5p
AATGACACGATCACTCCCGTTGA
>mmu-miR-669k-3p
TATGCATATACACGCATGCAA
>mmu-miR-6994-5p
ACAAAGGTGGGTGAAGCCGGTGA
>mmu-miR-8115
CCGCAATAGCAACAAGGGCTCT
>mmu-miR-7086-3p
TCCGTTCTGATATCTCCTCAG
>mmu-miR-6959-3p
CTGCACCGACCTGCTCTCCCAG
>mmu-miR-1191b-5p
TCAGGCTACAGAGCGAGAACCT
>mmu-miR-6411
TAGCATGTGTAGTACATAATG
>mmu-miR-224-5p

TAAGTCACTAGTGGTTCCGTT
>mmu-miR-1291
ATGGCTCTTACTGAAGACTAGCAGTT
>mmu-miR-693-3p
GCAGCTTTCAGATGTGGCTGTAA
>mmu-miR-434-5p
GCTCGACTCATGGTTTGAACCA
>mmu-miR-6950-3p
CTCTGTCTTGATCCTCTCCAG
>mmu-miR-3104-5p
TAGGGGGCAGGAGCCGGAGCCCTCT
>mmu-miR-301a-5p
GCTCTGACTTTATTGCACTACT
>mmu-miR-3077-5p
CTGCGGACGGGTGGGCGGGCAGGCC
>mmu-miR-3088-5p
GCTATGCAGTTGTTCATGAGCT
>mmu-miR-6540-3p
TCTGAAGCTTGCTTACCTCCA
>mmu-miR-6417
TAGCAAACGACAGCTAGCCACA
>mmu-miR-758-5p
TGGTTGACCAGAGAGCACACG
>mmu-miR-7050-5p
ACAGGAGAAGGGGGTGAGAGA
>mmu-miR-363-3p
AATTGCACGGTATCCATCTGTA
>mmu-miR-325-5p
CCTAGTAGGTGCTCAGTAAGTGT
>mmu-miR-6963-3p
TGCCTCTTGCCTCCATCCCACAG
>mmu-miR-491-5p
AGTGGGGAACCCTTCCATGAGG
>mmu-miR-1839-3p
AGACCTACTTATCTACCAACAGC
>mmu-miR-367-5p
ACTGTTGCTAACATGCAACTC
>mmu-miR-96-5p
TTTGGCACTAGCACATTTTTGCT
>mmu-miR-7001-5p
AGGCAGGGTGTGAGCGTGAGCAT
>mmu-miR-5129-5p
ATGTGGGGGCATTGGTATTTTC
>mmu-miR-6385
GCAAGGATGACAGAGAGGAAG
>mmu-miR-669o-5p
TAGTTGTGTGTGCATGTTTATGT
>mmu-miR-205-3p
GATTTCAGTGGAGTGAAGCTCA
>mmu-miR-1843b-5p
ATGGAGGTCTCTGTCTGACTT
>mmu-miR-1964-3p
CCGACTTCTGGGCTCCGGCTTT
>mmu-miR-684
AGTTTTCCCTTCAAGTCAA
>mmu-miR-471-5p
TACGTAGTATAGTGCTTTTCAC
>mmu-miR-7229-5p
TAGTAGACATCCCTGGATAGC
>mmu-miR-7688-3p
TCTGGCTTATGCCTAGCTTTC
>mmu-miR-6924-5p

AGAGGATGGGGATTTGGCGAAGT
>mmu-miR-190a-5p
TGATATGTTTGATATATTAGGT
>mmu-miR-7016-3p
ACCTGCCTCCTGCTCTCCCCAG
>mmu-miR-6988-5p
GGGGTGGAGAGCTGAGGCCCA
>mmu-miR-690
AAAGGCTAGGCTCACAACCAAA
>mmu-miR-217-5p
TACTGCATCAGGAACTGACTGGA
>mmu-miR-667-3p
TGACACCTGCCACCCAGCCCAAG
>mmu-miR-1198-3p
AAGCTAGCCTCTAACTCATGGC
>mmu-miR-6946-3p
TTTCTTCTCTTCCCTTTCAG
>mmu-miR-295-3p
AAAGTGCTACTACTTTTGAGTCT
>mmu-miR-7241-3p
TAGTAGATACTCATGGGAGAGC
>mmu-miR-6999-3p
CTTCAGCTGTCCTCCTTTCTGT
>mmu-miR-466b-3p|mmu-miR-466c-3p|mmu-miR-466p-3p
ATACATACACGCACACATAAGA
>mmu-miR-7017-3p
ACCCTGCTCCTCTCCCTCCAG
>mmu-miR-344g-3p
CAGGCTCTAGCCAGGGGCTTGA
>mmu-miR-7094-1-5p
TCTGAAGAGGATACAGGGTCTGA
>mmu-miR-291a-5p
CATCAAAGTGGAGGCCCTCTCT
>mmu-miR-6516-3p
TCATGTATGATACTGCAAACAG
>mmu-miR-133c
TTTGGTCCCCTTCAAGGAGTCAG
>mmu-miR-7083-3p
TCTGCTCTCCTGGACCCCAG
>mmu-miR-3098-5p
TCCTAACAGCAGGAGTAGGAGC
>mmu-miR-214-3p
ACAGCAGGCACAGACAGGCAGT
>mmu-miR-7216-3p
TGGAATTTGGTCTTTTCCGGC
>mmu-miR-1898
AGGTCAAGGTTCACAGGGGATC
>mmu-miR-1199-3p
TGCGGCCGGTGCTCAGTCGGC
>mmu-miR-666-3p
GGCTGCAGCGTGATCGCCTGCT
>mmu-miR-370-3p
GCCTGCTGGGGTGGAACCTGGT
>mmu-miR-8100
AGGAGGAAAGGGAGCAAGCAGGT
>mmu-miR-124b-3p
TCAAGGTCCGCTGTGAACACGG
>mmu-miR-324-5p
CGCATCCCCTAGGGCATTGGTGT
>mmu-miR-6986-5p
AGGGGAGCTAGGTAGAAAGCCA
>mmu-miR-6413

TGGCTCAGAAGAGCAGGTAGT
>mmu-miR-7648-5p
CCGCGTTCCGGGCTCGGCGC
>mmu-miR-184-3p
TGGACGGAGAACTGATAAGGGT
>mmu-miR-7b-3p
CAACAAGTCACAGCCAGCCTCA
>mmu-miR-30a-5p
TGTAAACATCCTCGACTGGAAG
>mmu-miR-6951-5p
TTGTATTTGTGTGATTAAAGT
>mmu-miR-3154
CAGAAGGGGAGTCGGGAGCGGA
>mmu-miR-6921-5p
TGAGGGGCATGAGGTAGGAAGC
>mmu-miR-6416-5p
CTCCGTATCATCTGCTCTTCT
>mmu-miR-292a-5p
ACTCAAACTGGGGGCTCTTTTG
>mmu-miR-5123
TGTAGATCCATATGCCATGGTGTG
>mmu-miR-7665-5p
AAGGGAAGGCAGGAGAAAGGCTG
>mmu-miR-495-5p
GAAGTTGCCCATGTTATTTTTCG
>mmu-miR-1971
GTAAAGGCTGGGCTGAGA
>mmu-miR-9-5p
TCTTTGGTTATCTAGCTGTATGA
>mmu-miR-7068-5p
GTGAGGCTCAGTATGGGGTGG
>mmu-miR-6981-3p
CAACAGCCTACTGTCTCCTCAG
>mmu-miR-145a-5p
GTCCAGTTTTCCCAGGAATCCCT
>mmu-miR-18a-3p
ACTGCCCTAAGTGCTCCTTCTG
>mmu-miR-1983
CTCACCTGGAGCATGTTTTCT
>mmu-miR-7686-3p
CTGCTCGGGGCACTGTAAGAGA
>mmu-miR-8106
TGACTCTGTACATGGCATTTAT
>mmu-miR-6905-3p
AAATAACCTTATCTGCCCGAT
>mmu-miR-6407
TGGCTGTCCCTGACAGGGCAAGT
>mmu-miR-668-5p
GTAAGTGTGCCTCGGGTGAGCATG
>mmu-miR-7064-3p
CAGGGCCCTTTATGTCTCTCT
>mmu-miR-500-3p
AATGCACCTGGGCAAGGGTTCA
>mmu-miR-8093
TTGGGGAGCAGAGCATGAGTG
>mmu-miR-7065-3p
TTGGTTTTCCTGTTTGCACAG
>mmu-miR-7029-5p
TCTGGACGCTCGGCTGCCAGCGC
>mmu-miR-5046
AGCTCCCGCCACTGTGACCCCCTT
>mmu-miR-505-3p

CGTCAACACTTGCTGGTTTTCT
>mmu-miR-6953-5p
AAGGGGCAGGGGCAGGGATTCAAGTG
>mmu-miR-379-3p
TATGTAACATGGTCCACTAACT
>mmu-miR-6936-5p
GTGAGCCACTGCCGTGCCCGC
>mmu-miR-5620-3p
ACAGTCATCCCCCTGCCTCAC
>mmu-miR-5125
TCTGCCTGGGATTTCCTTGT
>mmu-miR-3057-5p
ATTGGAGCTGAGATTCTGCGGGAT
>mmu-miR-125b-1-3p
ACGGGTTAGGCTCTTGGGAGCT
>mmu-miR-344-3p
TGATCTAGCCAAAGCCTGACTGT
>mmu-miR-211-5p
TTCCCTTTGTCATCCTTTGCCT
>mmu-miR-6966-3p
TGCTGTGTGCTTCCTCTGCAG
>mmu-miR-12194-3p
TCTGTGGGTCTGTTTGTCCGTC
>mmu-miR-1949
CTATACCAGGATGTCAGCATAGTT
>mmu-miR-1264-5p
AGGTCCTCAATAAGTATTTGTT
>mmu-miR-3083-3p
TCCGAAACATTCCCAGCCTTTA
>mmu-miR-3108-5p
GTCTCTAAAGCTAGACGTTCCGG
>mmu-miR-878-3p
GCATGACACCACACTGGGTAGA
>mmu-miR-6908-3p
ACACTCTCCCTTGTGCTGGCAG
>mmu-miR-484
TCAGGCTCAGTCCCCTCCCGAT
>mmu-miR-222-5p
CTCAGTAGCCAGTGTAGATCC
>mmu-miR-7090-5p
TCTCAGAGTTGGGACACCATGAG
>mmu-miR-7047-5p
TGAGGGAGGAGGGCTGGGTCTGA
>mmu-miR-511-5p
ATGCCTTTTGCTCTGCACTCA
>mmu-miR-8107
TGGAAACTAGTGGCTGGGACA
>mmu-miR-6904-3p
TCTGATCTCTTACCCCAG
>mmu-miR-7673-3p
TTCCTTACCTCACCGTCAAAGGA
>mmu-miR-3060-5p
GGGAGTGCTTCGTGCTTCTG
>mmu-miR-6958-5p
GTGGGAGAAGAGGCTGCTGTGG
>mmu-miR-3070-2-3p
TGGTGCTATGGTCAGGGGTAGA
>mmu-miR-653-5p
GTGTTGAAACAATCTCTACTG
>mmu-let-7i-3p
CTGCGCAAGCTACTGCCTTGCT
>mmu-miR-193a-5p

TGGGTCTTTGCGGGCAAGATGA
>mmu-miR-215-5p
ATGACCTATGATTTGACAGAC
>mmu-miR-7675-3p
TGTTTCATTTGTTCGTGACTC
>mmu-miR-1948-3p
TTTAGGCAGAGCACTCGTACAG
>mmu-miR-540-3p
AGGTCAGAGGTCGATCCTGG
>mmu-miR-6357
TAGCACTGCTCAGGGCTTTCAGG
>mmu-miR-7088-5p
GCAAGTGGTGGTGAGGAGGGGAAAC
>mmu-miR-466a-3p|mmu-miR-466e-3p
TATACATACACGCACACATAAGA
>mmu-miR-7688-5p
TAGCTGGGCATGATCTGATGAGC
>mmu-miR-7657-3p
TTCCTTGCTATTCCCAACTTGGT
>mmu-miR-7650-3p
GTTTTGATATATACAAGAAGGA
>mmu-miR-7062-5p
TGGAGGCCAGCTTGTGGAGTGTCT
>mmu-miR-7054-5p
TAGGAAGGTGGTTGGGCTGAGTACT
>mmu-miR-17-3p
ACTGCAGTGAGGGCACTTGTAG
>mmu-miR-3618-3p
CTACATTAATGAAAAGAGCAAT
>mmu-miR-711
GGGACCCGGGGAGAGATGTAAG
>mmu-miR-7658-3p
ACCACCTTCCTCACCCACGCT
>mmu-miR-3101-5p
GGTACCATTGACTAAAGCTAG
>mmu-miR-667-5p
CGGTGCTGGTGGAGCAGTGAGCACG
>mmu-miR-500-5p
AATCCTTGCTATCTGGGTGCTTAGT
>mmu-miR-201-5p
TACTCAGTAAGGCATTGTTCTT
>mmu-miR-7236-5p
CCTAAAACTGTGTGGTCCAATGCC
>mmu-miR-5623-5p
TGGCAGCTGGCAAGTCAGAATGCA
>mmu-miR-12202-3p
TCTTCTCTTTCCAGTCATCAGC
>mmu-miR-6363
TGTAGACTGGACTGTCCTCAAA
>mmu-miR-7667-5p
GAGCCATCTCTCTAGCCCCTGA
>mmu-miR-714
CGACGAGGGCCGGTCGGTCGC
>mmu-miR-302b-5p
ACTTTAACATGGGAATGCTTTCT
>mmu-miR-296-5p
AGGGCCCCCCCTCAATCCTGT
>mmu-miR-7230-3p
TTGAGTTGAGACTGTCAGTAGGT
>mmu-miR-7030-3p
CTTGCTCTGTTATCTCTCCAG
>mmu-miR-7a-1-3p

CAACAAATCACAGTCTGCCATA
>mmu-miR-15b-5p
TAGCAGCACATCATGGTTTACA
>mmu-miR-6370
GCAGGAACAGCAAAGGGGAAG
>mmu-miR-92a-3p
TATTGCACTTGTCCCGGCCTG
>mmu-miR-432
TCTTGGAGTAGATCAGTGGGCAG
>mmu-miR-7094-3p
TACTCTGCTGTTCTCTCCACA
>mmu-miR-7659-3p
AGGAGTCAGCTTGTCGGTTCATG
>mmu-miR-12201-5p
TGGAAGGATGCCTCTTGCAGA
>mmu-miR-7215-5p
GCTCTTTATGTCCAAGCTGAGCTAC
>mmu-miR-32-5p
TATTGCACATTACTAAGTTGCA
>mmu-miR-152-5p
TAGGTTCTGTGATACACTCCGACT
>mmu-miR-466d-5p
TGTGTGTGCGTACATGTACATG
>mmu-miR-376c-5p
GTGGATATTCCTTCTATGTTTA
>mmu-miR-7027-5p
TGGAAAGGAAGAAACAGCAGAGC
>mmu-miR-664-3p
TATTCATTTACTCCCCAGCCTA
>mmu-miR-1195
TGAGTTCGAGGCCAGCCTGCTCA
>mmu-miR-6541
CACAGGCAAGGACTTCTCAAG
>mmu-miR-12181-5p
TTGAGGGTGCATAAGGACTGGA
>mmu-miR-5709-3p
AAAGTCTTAAGGGTGTGTATTG
>mmu-let-7a-1-3p|mmu-let-7c-2-3p
CTATACAATCTACTGTCTTTCC
>mmu-let-7f-1-3p
CTATACAATCTATTGCCTTCCC
>mmu-miR-7687-5p
AGGCGGGGAACCTGAGGCGCAGGCT
>mmu-miR-361-3p
TCCCCCAGGTGTGATTCTGATTTGT
>mmu-miR-217-3p
CATCAGTTCCTAATGCATTGCCT
>mmu-miR-223-3p
TGTCAGTTTGTCAAATACCCCA
>mmu-miR-34c-3p
AATCACTAACCACACAGCCAGG
>mmu-miR-219a-5p
TGATTGTCCAAACGCAATTCT
>mmu-miR-290a-3p
AAAGTGCCGCCTAGTTTTAAGCCC
>mmu-miR-668-3p
TGTCACTCGGCTCGGCCCACTACC
>mmu-miR-741-5p
TACGTAGATTGGTACCTATCATG
>mmu-miR-669j
TGCATATACTCACATGCAAACA
>mmu-miR-653-3p

TTCACTGGAGTTTGTTTCAGT
>mmu-miR-880-5p
TACTCAGATTGATATGAGTCA
>mmu-miR-3966
AGCTGCCAGCTGTAGAACTGT
>mmu-miR-1950
TCTGCATCTAAGGATATGGTCA
>mmu-miR-7064-5p
TCAGAGGCATAAAGGTCATTTGCT
>mmu-miR-466p-5p
TATGTGTGTGTACATGTACAT
>mmu-miR-140-5p
CAGTGGTTTTACCCTATGGTAG
>mmu-miR-22-5p
AGTTCTTCAGTGGCAAGCTTTA
>mmu-miR-344c-5p
AGTCAGGCTGCTGGCTAGAGTAC
>mmu-miR-697
AACATCCTGGTCCTGTGGAGA
>mmu-miR-12206-5p
TACTATGCCTGGAAGGCACCA
>mmu-miR-101a-5p
TCAGTTATCACAGTGCTGATGC
>mmu-miR-34b-3p
AATCACTAACTCCACTGCCATC
>mmu-miR-449c-5p
AGGCAGTGCATTGCTAGCTGG
>mmu-miR-138-1-3p
CGGCTACTTCACAACACCAGGG
>mmu-miR-6928-5p
AGGGGATTTGCATGGACTGCACT
>mmu-miR-671-3p
TCCGGTTCTCAGGGCTCCACC
>mmu-miR-23a-5p
GGGGTTCCTGGGGATGGGATTT
>mmu-miR-129-2-3p
AAGCCCTTACCCCAAAAAGCAT
>mmu-miR-7058-5p
TAGAGGGGAGGTGACCGTGGAAGT
>mmu-miR-1929-5p
TTCTAGGACTTTATAGAGCAGAG
>mmu-miR-2137
GCCGGCGGGAGCCCCAGGGAG
>mmu-miR-7238-5p
ATGCGGATGGGACGCGGC
>mmu-miR-7036a-3p
CCGTCCTCATCCGCTCCTCCCAG
>mmu-miR-5121
AGCTTGTGATGAGACATCTCC
>mmu-miR-1936
TAACTGACCTGCTGTGAACTGGC
>mmu-miR-28b
AGGAGCTCACAATCTATTTAG
>mmu-miR-449a-5p
TGGCAGTGTATTGTTAGCTGGT
>mmu-miR-3087-3p
TAACTCACTGTCATGTCCTCA
>mmu-miR-344i
AAGTCAGGCTCCTGGCTGGA
>mmu-miR-216c-5p
GAAGAATCTCTACAGGTAAGTGT
>mmu-miR-6960-5p

CAGGATGAGAAGAGTTTGGCTG
>mmu-miR-449c-3p
CAGTTGCTAGCTGCACTACCCT
>mmu-miR-1264-3p
CAAATCTTATTTGAGCACCTGT
>mmu-miR-489-5p
TGTCATATGTGTGATGACACTTTCT
>mmu-miR-485-5p
AGAGGCTGGCCGTGATGAATTC
>mmu-miR-6539
GCACAGTGATGAACTCTGAGGGCT
>mmu-miR-378d
ACTGGCCTTGGAGTCAGAAGGT
>mmu-miR-1958
TAGGAAAGTGGAAGCAGTAAGT
>mmu-miR-12196-5p
TCCTTTCTAGCTCACCTGTGACT
>mmu-miR-6971-5p
TGGGGGAGGGTGTAGAGGCT
>mmu-miR-6997-3p
TCAAACCTTACCCTCCTGTTTCC
>mmu-miR-1945
TCTTCGCGGGTACTGTCGGGAC
>mmu-miR-676-5p
ACTCTACAACCTTAGGACTTGC
>mmu-miR-7689-3p
TTAGAGCCAGACTGCCTGGGTTT
>mmu-miR-5622-3p
CTTAGCTGGGTAGTGGTGGTGC
>mmu-miR-33-3p
CAATGTTTCCACAGTGCATCAC
>mmu-miR-3091-5p
CATGGGTCTGGTTGGGCCCGC
>mmu-miR-216a-3p
CACAGTGGTCTCTGGGATTATG
>mmu-miR-6336
TCTCGGATTTAGTAAGAGATC
>mmu-miR-3100-3p
CTGTGACACACCCGCTCCCAG
>mmu-miR-8096
GGGCACGGTAGCTGAAGAGT
>mmu-miR-8095
AAAGGATTCTGCTGTCTGTCCC
>mmu-miR-6939-3p
CCTCACTTGACCCGCTGCAG
>mmu-miR-362-3p
AACACACCTGTTCAAGGATTCA
>mmu-miR-384-5p
TGTAAACAATTCCTAGGCAATGT
>mmu-miR-7222-5p
GGCACTTGCCAGGCTCTGAGCTGGACT
>mmu-miR-3085-5p
AGGTGCCATTCCGAGGGCCAAGAGT
>mmu-miR-8117
GCTCGTGTGGAACAGAAGGGG
>mmu-miR-7228-5p
TGGCGACCTGAACAGATGT
>mmu-miR-7671-5p
AGGCTCCGCTCTGGCTAGCTTCC
>mmu-miR-12197-3p
GCGGGGCTGGGGACATCG
>mmu-miR-1981-3p

CATCTAACCCTGGCCTTTGAC
>mmu-miR-7012-5p
AAGGAGAGGAGTTGGCAGGGACT
>mmu-miR-467d-3p
ATATACATACACACACCTACAC
>mmu-miR-1843a-5p
TATGGAGGTCTCTGTCTGACT
>mmu-miR-6912-3p
CTACCTGAGGGTCTCCTTCCTGCAG
>mmu-miR-767
TGCACCATGGTTGTCTGAGCA
>mmu-miR-22-3p
AAGCTGCCAGTTGAAGAACTGT
>mmu-miR-16-2-3p
ACCAATATTATTGTGCTGCTTT
>mmu-miR-12205-5p
TGTGTTTCCCAGTTGGTTTGA
>mmu-miR-7074-3p
TAGGCCTTCTCCTCTCCCTGCAG
>mmu-miR-6919-5p
TAGGCCACTGGAGGTGGAATTGT
>mmu-miR-30d-3p
CTTTCAGTCAGATGTTTGCTGC
>mmu-miR-7077-5p
GGCCAAGGTCCCATGGCAGGC
>mmu-miR-7033-5p
TCTCCAGGAGTCTGAGGGGCAGG
>mmu-miR-466f
ACGTGTGTGTGCATGTGCATGT
>mmu-miR-187-5p
AGGCTACAACACAGGACCCGGG
>mmu-miR-214-5p
TGCCTGTCTACACTTGCTGTGC
>mmu-miR-551b-5p
GAAATCAAGCTTGGGTGAGACCT
>mmu-miR-669d-2-3p
ATATACATACACACCCATATAC
>mmu-miR-6987-3p
CACCCATTTCTCCCTGCTTCCAG
>mmu-miR-290a-5p
ACTCAAACTATGGGGGCACTTT
>mmu-miR-338-5p
AACAATATCCTGGTGCTGAGTG
>mmu-miR-6244
GGGGGCGTGGAATTATCGGGT
>mmu-miR-374c-3p
ACTTAGCAGGTTGTATTAT
>mmu-miR-6360
TAGTGTTGCTCAGGCAGCAGGA
>mmu-miR-6997-5p
TAACAGGCTGGAGAGGTGCAGA
>mmu-miR-6946-5p
AAAAGGGAGAGAAAGAAATGC
>mmu-miR-6384
GCTTTCCTACTGTTTCCCTG
>mmu-miR-367-3p
AATTGCACTTTAGCAATGGTGA
>mmu-miR-6998-5p
CTGGGCAGAGGGCAAAGTGACT
>mmu-miR-124-5p
CGTGTTCACAGCGGACCTTGAT
>mmu-miR-27a-3p

TTCACAGTGGCTAAGTTCCGC
>mmu-miR-6974-3p
TCTCCACTCTCTTCTGTCCCAG
>mmu-miR-1955-5p
AGTCCCAGGATGCACTGCAGCTTTT
>mmu-miR-497a-5p
CAGCAGCACACTGTGGTTTGTA
>mmu-miR-292a-3p
AAAGTGCCGCCAGGTTTTGAGTGT
>mmu-miR-103-1-5p
GGCTTCTTTACAGTGCTGCCTTG
>mmu-miR-7213-3p
TACCTCAAGAGAGCCAGTCT
>mmu-miR-26b-5p
TTCAAGTAATTCAGGATAGGT
>mmu-miR-7687-3p
CCACTCAGTCGCCCCGCCACA
>mmu-miR-6983-5p
TTGGAAGGGCATACTGATTCGG
>mmu-miR-873a-5p
GCAGGAACTTGTGAGTCTCCT
>mmu-miR-7650-5p
AATCCTCTTGCAACCCAGAACT
>mmu-miR-27b-3p
TTCACAGTGGCTAAGTTCTGC
>mmu-miR-7656-5p
TGTGGGATCAGATAGCCTGTTA
>mmu-miR-5126
GCGGGCGGGGCCGGGGGCGGGG
>mmu-miR-543-3p
AAACATTCGCGGTGCACTTCTT
>mmu-miR-6943-5p
TGGGGTGAGGTTGGAAGCTA
>mmu-miR-3077-3p
CTGACTCCCTGCTTCTCCGCAG
>mmu-miR-509-3p
TGATTGACATTTCTGTAATGG
>mmu-miR-12184-3p
ACAGACGGGCTCTCATGCTGA
>mmu-miR-186-3p
GCCCTAAGGTGAATTTTTTGGG
>mmu-miR-6364
TCGATCCCCAGCACTCCTGTGA
>mmu-miR-30e-5p
TGTAAACATCCTTGACTGGAAG
>mmu-miR-6335
CTGCATACACAGTGATATTCTG
>mmu-miR-6976-3p
ATCTCCCTCCCAACTCCCCAG
>mmu-miR-382-5p
GAAGTTGTTCGTGGTGGATTCG
>mmu-miR-681
CAGCCTCGCTGGCAGGCAGCT
>mmu-miR-30c-2-3p
CTGGGAGAAGGCTGTTTACTCT
>mmu-miR-6961-3p
TCCTCTTTCTTCCTGGCTCTAG
>mmu-miR-6989-5p
TTGGGGACAGAGGCACAGGATG
>mmu-miR-466d-3p
TATACATACACGCACACATAG
>mmu-miR-7093-3p

TTTCCATCTGTCATCCTGCAG
>mmu-miR-7091-3p
AGTGGCTTCTGTCGTCTCTAG
>mmu-miR-7079-3p
TGACTTCTGTTCTCTTTCCAG
>mmu-miR-7658-5p
TGTGGGCGTGGCGGTGCTGG
>mmu-miR-7013-3p
CCACACTTACTGTTGCCTCTTCCT
>mmu-miR-7000-3p
CACCCACCTGCCTGTCCTCCAG
>mmu-miR-25-3p
CATTGCACTTGTCTCGGTCTGA
>mmu-miR-6910-5p
TGGGGGTAGGGCACCAGTGGGCA
>mmu-miR-503-5p
TAGCAGCGGGAACAGTACTGCAG
>mmu-miR-298-3p
GAGGAACTAGCCTTCTCTCAGCTT
>mmu-miR-539-5p
GGAGAAATTATCCTTGGTGTGT
>mmu-miR-6964-5p
TTGGGGTGTGAGACTGGAAAC
>mmu-miR-466l-3p
TATAAATACATGCACACATATT
>mmu-miR-7684-5p
TCTGGGAAGCCTGGGCAGCAG
>mmu-miR-12178-3p
TTTCTGACTCAGCCACCTGAGA
>mmu-miR-3547-5p
GTGGGAAGAGGGGTGGGGCCCGGGA
>mmu-miR-5615-3p
TGTCTCAGAAAACAACCAAGGA
>mmu-miR-6393
CTGCCCACGAAGCACACTGAGT
>mmu-miR-532-5p
CATGCCTTGAGTGTAGGACCGT
>mmu-miR-495-3p
AAACAAACATGGTGCACTTCTT
>mmu-miR-6959-5p
TGGGAAACCTGTGTCGGGCTGTG
>mmu-miR-6903-5p
TTGGGAAAAGCAGTACTTACCA
>mmu-miR-883b-3p
TAACTGCAACATCTCTCAGTAT
>mmu-miR-744-5p
TGCGGGGCTAGGGCTAACAGCA
>mmu-miR-8120
TGTGCCTTGCAACTTCCGAGCC
>mmu-miR-7656-3p
ACAGGCTGTCTGATCCCACGGT
>mmu-miR-2183
TTGAACCCCTGACCTCCT
>mmu-miR-196a-1-3p
CAACGACATCAAACCACCTGAT
>mmu-miR-3070-5p
AGCCCCTGACCTTGAACCTGGGA
>mmu-miR-466k
TGTGTGTGTACATGTACATGTGA
>mmu-miR-6977-5p
GTGAGGCGCTGTGGGCACGGCTTGG
>mmu-miR-6969-5p

CAGCAGAGTAGGGAGCAAAAGA
>mmu-miR-215-3p
TCTGTCATTCTGTAGGCCAAT
>mmu-miR-5124b
TGGCTCAGTGACTAAGAAGCA
>mmu-miR-98-5p
TGAGGTAGTAAGTTGTATTGTT
>mmu-miR-7070-5p
GTGAGGGGAGCTGAGGCAGGA
>mmu-miR-7663-3p
AGTGTGGATCATAAGGAGCTCC
>mmu-miR-669c-5p
ATAGTTGTGTGTGGATGTGTGT
>mmu-miR-300-3p
TATGCAAGGGCAAGCTCTCTTC
>mmu-miR-297a-5p
ATGTATGTGTGCATGTGCATGT
>mmu-miR-7051-3p
TGAGCCACTGATTCCTGTGTGC
>mmu-miR-7010-5p
CGGAAAGGATTGGGAAACTTGA
>mmu-miR-380-5p
ATGGTTGACCATAGAACATGCG
>mmu-miR-6540-5p
CTAAGGCAGGCAGACTTCAGTG
>mmu-miR-449a-3p
CAGCTAACATGCGACTGCTCTC
>mmu-miR-7219-3p
TATCCGGGTTTCTAACACACT
>mmu-miR-5127
TCTCCCAACCCTTTTCCCA
>mmu-miR-3087-5p
CAGGGCAGGGCAAGAGTTGAG
>mmu-miR-346-5p
TGTCTGCCCGAGTGCCTGCCTCT
>mmu-miR-718
CTTCCGCCCGGCCGGGTGTCG
>mmu-let-7g-5p
TGAGGTAGTAGTTTGTACAGTT
>mmu-miR-203-3p
GTGAAATGTTTAGGACCACTAG
>mmu-miR-433-3p
ATCATGATGGGCTCCTCGGTGT
>mmu-miR-7071-5p
CCCGGGAGCGAGGTGCTGCAGCG
>mmu-miR-206-5p
ACATGCTTCTTTATATCCTCATA
>mmu-miR-7062-3p
ACTAACTTCTCCTGGCCCCACAG
>mmu-miR-6403
GCAAAGGCGCTGTTTCCCAGTG
>mmu-miR-466b-5p|mmu-miR-466o-5p
TGATGTGTGTGTACATGTACAT
>mmu-miR-1953
TGGGAAAGTTCTCAGGCTTCTG
>mmu-miR-7061-3p
TATCCCTTTTATTTCCTTTTAG
>mmu-miR-339-5p
TCCCTGTCCTCCAGGAGCTCACG
>mmu-miR-465d-3p
TGATCAGGGCCTTTCTGAGTAA
>mmu-miR-5133

GCTGGAGCTGCGGCAGCGCAG
>mmu-miR-6984-5p
ACTGAAAGGCAATGAAGGAGGAGC
>mmu-miR-138-5p
AGCTGGTGTTGTGAATCAGGCCG
>mmu-miR-1928
AGCTACATTGCCAGCTC
>mmu-miR-466i-3p
ATACACACACACATACACACTA
>mmu-miR-328-3p
CTGGCCCTCTCTGCCCTTCCGT
>mmu-miR-18b-5p
TAAGGTGCATCTAGTGCTGTTAG
>mmu-miR-324-3p
CCACTGCCCCAGGTGCTGCT
>mmu-miR-7023-3p
TCACCCTGTCTGCGCCCCTCAG
>mmu-miR-7119-5p
CTGGGGGATTGGGTGCCTTCT
>mmu-miR-6989-3p
TGTCCACTGTCTCTGGTCCCAG
>mmu-miR-598-3p
TACGTCATCGTCGTCATCGTTA
>mmu-miR-133a-5p
GCTGGTAAAATGGAACCAAAT
>mmu-miR-7019-3p
TCACCTTGGCCGCCTCTCTGCAG
>mmu-miR-1948-5p
ATATGAGTATTCTGCCTAAAT
>mmu-miR-7069-3p
CCTGCCCTGCCAACCCCTACA
>mmu-miR-488-3p
TTGAAAGGCTGTTTCTTGGTC
>mmu-miR-496a-5p
AGGTTGCCCATGGTGTGTTCA
>mmu-miR-194-2-3p
CCAGTGGGGCTGCTGTTATCTG
>mmu-miR-369-5p
AGATCGACCGTGTTATATTCGC
>mmu-miR-3097-3p
CTCAGACCTTTCTACCTGTCAG
>mmu-miR-7237-3p
CATCCTGTTGAGCTTACCGAGC
>mmu-miR-7011-3p
TCTGCTTCCCTCCTCCTCTAG
>mmu-miR-5136
ATATGCGAGGGAACTACTGG
>mmu-miR-7072-3p
CTAGCTTCTCTTCTTCCCCAG
>mmu-miR-7031-5p
CCTGAGAGGCCTGAAGGGTGGGA
>mmu-miR-6352
TTTGGGAAAAGGACCCCAGCTC
>mmu-miR-365-1-5p
AGGGACTTTTGGGGGCAGATGTG
>mmu-miR-369-3p
AATAATACATGGTTGATCTTT
>mmu-miR-875-5p
TATACCTCAGTTTTATCAGGTG
>mmu-miR-6926-3p
TCACCATCCCTGTCTCCATAG
>mmu-miR-712-3p

TGCGAGTCACCCCCGGGTGTTG
>mmu-miR-483-3p
TCACTCCTCCCCTCCCGTCTT
>mmu-miR-7018-3p
TCACCCTGCTGCCGGCTTGCAG
>mmu-miR-7b-5p
TGGAAGACTTGTGATTTTGTTGTT
>mmu-miR-654-5p
TGGTAAGCTGCAGAACATGTGT
>mmu-miR-669b-5p
AGTTTTGTGTGCATGTGCATGT
>mmu-miR-8101
GGCGAGCGGAGCCGGAGGAGCC
>mmu-miR-330-5p
TCTCTGGGCCTGTGTCTTAGGC
>mmu-miR-669l-5p
AGTTGTGTGTGCATGTATATGT
>mmu-miR-6238
TTATTAGTCAGTGGAGGAAATG
>mmu-miR-350-3p
TTCACAAAGCCCATACACTTTC
>mmu-miR-6948-3p
TGTCCTTCCTGTCTGACCACA
>mmu-miR-183-5p
TATGGCACTGGTAGAATTCACT
>mmu-miR-29b-3p
TAGCACCATTTGAAATCAGTGTT
>mmu-miR-339-3p
TGAGCGCCTCGGCGACAGAGCCG
>mmu-miR-6409
TGCGGGTGAGGATGTGTGCTCCT
>mmu-miR-181b-2-3p
CTCACTGATCAATGAATGCAAA
>mmu-miR-5099
TTAGATCGATGTGGTGCTCC
>mmu-miR-494-3p
TGAAACATACACGGGAAACCTC
>mmu-miR-147-5p
TGGAAACATTTCTGCACAAACTAG
>mmu-miR-1a-3p
TGGAATGTAAAGAAGTATGTAT
>mmu-miR-335-5p
TCAAGAGCAATAACGAAAAATGT
>mmu-miR-5620-5p
ACGAGGCAGGGGCTTTGACTGTG
>mmu-miR-3092-5p
AGGGGAAAATGCCTTTCTCCCA
>mmu-miR-701-3p
TATCTATTAAAGAGGCTAGC
>mmu-let-7e-5p
TGAGGTAGGAGGTTGTATAGTT
>mmu-miR-7015-3p
TCTCACTGTCCTCTGCACTAG
>mmu-miR-5118
AAGGTTAGGCCAGCCTGGT
>mmu-miR-802-5p
TCAGTAACAAAGATTCATCCTT
>mmu-miR-7116-5p
TGAAGACATCAGGAAAAAAAA
>mmu-miR-6976-5p
CAGGGAAGTTGAGAGGAAAATTG
>mmu-miR-6399

TTGCAATGATGGTATTCTGAGG
>mmu-miR-6362
GTGGGCATCTAGGCTAGTTCTG
>mmu-miR-7070-3p
CAGGCTCTCCTTCCCTCCAG
>mmu-miR-8109
GCGCCGCGTGCCGGCCGCGGG
>mmu-miR-136-5p
ACTCCATTTGTTTTGATGATGG
>mmu-miR-6516-5p
TTTGCAGTAACAGGTGTGGACA
>mmu-miR-3059-5p
TTTCCTCTCTGCCCCATAGGGT
>mmu-miR-12203-3p
ATGGAATCTCTCCCTCTGGTCT
>mmu-miR-6990-5p
CCCAGGGTGAGTCAGGGCTCT
>mmu-miR-105
CCAAGTGCTCAGATGCTTGTGGT
>mmu-miR-3083b-5p
AAGGCTGGGAATGTTTCGGAGA
>mmu-miR-381-3p
TATACAAGGGCAAGCTCTCTGT
>mmu-miR-6343
TGTAAAGTAGGAGCTGACTGAC
>mmu-miR-693-5p
CAGCCACATCCGAAAGTTTTC
>mmu-let-7f-2-3p
CTATACAGTCTACTGTCTTTC
>mmu-miR-3059-3p
CCTCTAGGGAAGAGAAGGTTGG
>mmu-miR-706
AGAGAAACCCTGTCTCAAAAAA
>mmu-miR-7682-3p
CCTGTGGGTTGGGTTGGCTTT
>mmu-miR-1894-5p
CTCTCCCCTACCACCTGCCTCT
>mmu-miR-7018-5p
GTGAGCAGACAGGGAGTGGTGGGG
>mmu-miR-6980-5p
GTGGGGGGGGAGGCTAGGTTAG
>mmu-miR-3090-5p
GTCTGGGTGGGGCCTGAGATC
>mmu-miR-200c-3p
TAATACTGCCGGGTAATGATGGA
>mmu-miR-7677-3p
CCGGCTGTGTCCTCTCACCTAGG
>mmu-miR-322-3p
AAACATGAAGCGCTGCAACAC
>mmu-miR-292b-3p
AAGAGCCCCCAGTTTGAGTAT
>mmu-miR-6236
GCCGTCGCCGGCAGTCAGG
>mmu-miR-1947-3p
GCACTGAGCTAGCTCTCCCTCC
>mmu-miR-196b-5p
TAGGTAGTTTCCTGTTGTTGGG
>mmu-miR-741-3p
TGAGAGATGCCATTCTATGTAGA
>mmu-miR-6970-5p
GTAAGTTCAGGGCTGGGAGCAGAGA
>mmu-miR-181a-2-3p

ACCACCGACCGTTGACTGTACC
>mmu-miR-7212-3p
TAACACACACGTCTCCAGGTC
>mmu-miR-7026-3p
TGTGCTTTCTGGTCTTGGCTTAG
>mmu-miR-16-1-3p
CCAGTATTGACTGTGCTGCTGA
>mmu-miR-8102
TCACGCGGGGGAACGAGGAAGA
>mmu-miR-669h-3p
TATGCATATACACACATGCACA
>mmu-miR-135b-5p
TATGGCTTTTCATTCCTATGTGA
>mmu-miR-6999-5p
AAGGAAGGAGAGTCAGCAAGCAC
>mmu-miR-31-5p
AGGCAAGATGCTGGCATAGCTG
>mmu-miR-7042-3p
TGTCCCTTTGTTTTCTCTCAG
>mmu-miR-6995-5p
CTGGGAGTAGAAGGGGGAAACCA
>mmu-miR-9-3p
ATAAAGCTAGATAACCGAAAGT
>mmu-miR-6985-5p
TACTGAGGGGTGGCTGCTATGT
>mmu-miR-6992-5p
TGCCTGTGATGGTTTGGCTGAGT
>mmu-miR-6948-5p
AGTTCAGACAGGACTGTGACAC
>mmu-miR-1970
TGTGTCACTGGGGATAGGCTTTG
>mmu-miR-7063-3p
TGCTCTCTGCCCCTCTTAAG
>mmu-miR-7021-3p
TCTCTGTGCTTCTGCTCCTTAG
>mmu-miR-6966-5p
CTCGGGGATGTGGACAGCACA
>mmu-miR-7657-5p
AATAGTTGGATAGTTAGGTAA
>mmu-miR-6416-3p
GCAAAGAGCAGCAAAAGGAAG
>mmu-miR-5134-3p
ACGGGTGGCCCTCTTTCTGCAG
>mmu-miR-7220-5p
GGTGAGCTCTTGGTACCTTGGC
>mmu-miR-193a-3p
AACTGGCCTACAAAGTCCCAGT
>mmu-miR-671-5p
AGGAAGCCCTGGAGGGGCTGGAG
>mmu-miR-1934-3p
AGGATGACGGTGGGGCTGGTGA
>mmu-miR-708-3p
CAACTAGACTGTGAGCTTCTAG
>mmu-miR-6387
TGGAATGGCTTCCCTTGTGTAT
>mmu-miR-3102-5p.2-5p
GGTGGTGCAGGCAGGAGAGCC
>mmu-miR-6973b-3p
TGCTCTCTTACCCTTCCTAG
>mmu-miR-30b-5p
TGTAAACATCCTACACTCAGCT
>mmu-miR-216a-5p

TAATCTCAGCTGGCAACTGTGA
>mmu-miR-7686-5p
CCTTCCACTGGACCTGGGGCTGGGC
>mmu-miR-296-3p
GAGGGTTGGGTGGAGGCTCTCC
>mmu-miR-7084-3p
GCCTCTGACCCCTGTCCTCTGC
>mmu-miR-6972-3p
TTACTGTACCTGTCTCCATAG
>mmu-miR-6962-5p
AAGGACCTGGTGGGAGATCT
>mmu-miR-758-3p
TTTGTGACCTGGTCCACTA
>mmu-miR-135a-2-3p
TGTAGGGATGGAAGCCATGAA
>mmu-miR-3544-5p
AGAAAGGCATCATATAGGAGCTG
>mmu-miR-7039-5p
AGAGGGATGGAGAGCATCGCGT
>mmu-miR-218-2-3p
CATGGTTCTGTCAAGCACCGCG
>mmu-let-7d-5p
AGAGGTAGTAGGTTGCATAGTT
>mmu-miR-325-3p
TTTATTGAGCACCTCCTATCAA
>mmu-miR-12188-5p
TAGAAGAGCTTTCGCTGGCCCAGA
>mmu-let-7a-2-3p
CTGTACAGCCTCCTAGCTTTC
>mmu-miR-1970b-5p
TATGTCACTGGGCATTGGCTTTGA
>mmu-miR-6940-5p
AGGGCTGGCTGGAAGGTTTTTCG
>mmu-miR-3102-3p
GAGCACCCCATTGGCTACCCACA
>mmu-miR-1930-3p
GGTGCAGTTACTGTGGCTGTGG
>mmu-miR-7659-5p
AGGAGCCGGCTCCAGATTCCTG
>mmu-miR-185-3p
AGGGGCTGGCTTTCCTCTGGT
>mmu-miR-1258-3p
TTAGGGAATTAGCTCAGCAGTA
>mmu-miR-7069-5p
TTGGGGGCCTGGGAGGGTGACAC
>mmu-miR-7024-5p
TTGGGGGATGGGTTGCTTGGC
>mmu-miR-5100
TCGAATCCCAGCGGTGCCTCT
>mmu-miR-7003-5p
TGTGGGGAGAAGCTGCGGCAG
>mmu-miR-96-3p
CAATCATGTGTAGTGCCAATAT
>mmu-miR-376b-3p
ATCATAGAGGAACATCCACTT
>mmu-miR-652-3p
AATGGCGCCACTAGGGTTGTG
>mmu-miR-375-3p
TTTGTTCGTTCGGCTCGCGTGA
>mmu-miR-3473a
TGGAGAGATGGCTCAGCA
>mmu-miR-6939-5p

CAGGCAGCTGGTTGGTGGAAG
>mmu-miR-7664-5p
CAGTTTACCTGTCTATGAGAT
>mmu-miR-468-3p
TATGACTGATGTGCGTGTGTCTG
>mmu-miR-122b-3p
AAACACCATTGTCACACTCCAC
>mmu-miR-6898-3p
AAGTCTCTTTGTACCTTCTCT
>mmu-miR-6337
TGAGGGCTGCTGGGTTGCTAGGT
>mmu-miR-1962
AGAGGCTGGCACTGGGACACAT
>mmu-miR-466h-3p
TACGCACGCACACACACAC
>mmu-miR-2861
GGGGCCTGGCGGCGGGCGG
>mmu-miR-299b-3p
TATGTGGGACGGTAAACC
>mmu-miR-7032-5p
CCCAGGGTGGTCCCCGAGAGGGTGGT
>mmu-miR-669f-3p
CATATACATACACACACACGTAT
>mmu-miR-5619-5p
AGTCACTAATGGCATTTTTGT
>mmu-miR-103-2-5p
AGCTTCTTTACAGTGCTGCCTTG
>mmu-miR-7082-3p
GCCCCCCTTGCTGTCTGTCTCC
>mmu-miR-294-3p
AAAGTGCTTCCCTTTTGTGTGT
>mmu-miR-12188-3p
TGGACAGCAGTACCCCAGCCCCTCTGC
>mmu-miR-7042-5p
TAGAGACAGCAGAAGGGCCAC
>mmu-miR-5107-3p
CAACCTGTGCTTCTCTTCCCAG
>mmu-miR-7685-5p
ACCTTCCGGTTTCTTCAAGTCTCC
>mmu-miR-190b-3p
ACTGAATGTCAAGCATACTCTCA
>mmu-miR-6899-5p
AGCAGAACGCAGCGGGCATGA
>mmu-miR-3470a
TCACTTTGTAGACCAGGCTGG
>mmu-miR-6907-5p
TTCTGGGACACAGGGAGGCTCG
>mmu-miR-7655-3p
TTTATGACGCTCCGTGGGCCTG
>mmu-miR-181a-5p
AACATTCAACGCTGTCGGTGAGT
>mmu-miR-467a-5p
TAAGTGCCTGCATGTATATGCG
>mmu-miR-7218-3p
CTAACCTGTCATTCTAGCCTGGGG
>mmu-miR-6382
TGGAATGTAAAGAGAGCACACAAG
>mmu-miR-7014-5p
TTGGGTGCTGTGGAAGGGACAG
>mmu-miR-712-5p
CTCCTTCACCCGGGCGGTACC
>mmu-miR-1224-3p

CCCCACCTCTTCTCTCCTCAG
>mmu-miR-12178-5p
TATGGGTGGCTGGTCTCAGAAGAG
>mmu-miR-6414
CTAAGTTCTGAGCTGAGTCTTG
>mmu-miR-6998-3p
AGAGCTGCTCTGTGCCCACACA
>mmu-miR-298-5p
GGCAGAGGAGGGCTGTTCTTCCC
>mmu-miR-448-5p
GAACATCCTGCATAGTGCTGCC
>mmu-miR-1271-5p
CTTGGCACCTGGTAAGCACTCA
>mmu-miR-6546-3p
GAGCGAAGCCCAAGCGCCAGA
>mmu-miR-7043-5p
TGTGAAAGCAGAGAGGCATTTTT
>mmu-miR-497a-3p
CAAACCACACTGTGGTGTTAG
>mmu-miR-152-3p
TCAGTGCATGACAGAACTTGG
>mmu-miR-6356
TCCCCAGAGTCCTAACAATGA
>mmu-miR-467b-5p
GTAAGTGCCTGCATGTATATG
>mmu-miR-7053-3p
CTCCTGTGTCTCCTTCCCCAG
>mmu-miR-3109-5p
AATGGATGCGATGGTTCCCATGCT
>mmu-miR-1895
CCCCCGAGGAGGACGAGGAGGA
>mmu-miR-24-3p
TGGCTCAGTTCAGCAGGAACAG
>mmu-let-7j
TGAGGTATTAGTTTGTGCTGTTAT
>mmu-miR-423-5p
TGAGGGGCAGAGAGCGAGACTTT
>mmu-miR-3572-3p
TACACTTGTCCTTCTTTCCCCAG
>mmu-miR-127-5p
CTGAAGCTCAGAGGGCTCTGAT
>mmu-miR-7034-3p
TTGGCCTTCCTTCCCTACCCTACA
>mmu-miR-494-5p
AGGTTGTCCGTGTTGTCTTCTC
>mmu-miR-7075-3p
CAACCATGTCTTCTTTCCCAG
>mmu-miR-2136
CTGGGTGTTGACTGAGATGTG
>mmu-miR-1192
AAACAAACAAACAGACCAAATT
>mmu-miR-7679-3p
CAGGATCCCAGGGTACCAACCAGC
>mmu-miR-302a-3p
TAAGTGCTTCCATGTTTTGGTGA
>mmu-miR-691
ATTCCTGAAGAGAGGCAGAAAA
>mmu-miR-6957-3p
TGGATCTCTTGTTTTCTCTTCC
>mmu-miR-7676-3p
TCCGGTGCTCACTCTGCCCACA
>mmu-miR-7646-5p

ACAAGAGAACTGATCAGTGCTG
>mmu-miR-6932-5p
TTGTAGGTGAAGTAGACTTGGC
>mmu-miR-18b-3p
TACTGCCCTAAATGCCCCTTCT
>mmu-miR-7025-5p
CGTGAGCTGAAGCTGGTGGCTCCC
>mmu-miR-6365
TTAGCCTTTAATATATTTGTTG
>mmu-miR-7235-5p
GGAGGGAGGGGTCTGGGC
>mmu-miR-3961
TGCCCTCAGCTCAGTTGGA
>mmu-miR-1930-5p
ACCTCCATAGTACCTGCAGCGT
>mmu-let-7g-3p
ACTGTACAGGCCACTGCCTTGC
>mmu-miR-6396
GGGGCCTTAGCTCCTAGTCCTGGA
>mmu-miR-466m-3p
TACATACACACATACACACGCA
>mmu-miR-7213-5p
ACTGGGTCTCTTGAGGTAGC
>mmu-miR-7086-5p
AAGAGGAGAAAGGTTTGGGCA
>mmu-miR-195b
TAGCAGCACAGAAATAGTAGAA
>mmu-miR-935
CCCAGTTACCGCTTCCGCTACCGC
>mmu-miR-6951-3p
CTTTTTCTTCACAAATACAG
>mmu-miR-344b-3p
CATTTAGCCAAAGCCTGACTGT
>mmu-miR-3071-3p
ATCATCAAAACAAATGGAGTCC
>mmu-miR-3090-3p
TCCCAGGTGACACCCTGACTCA
>mmu-miR-499-5p
TTAAGACTTGCAGTGATGTTT
>mmu-miR-331-3p
GCCCCTGGGCCTATCCTAGAA
>mmu-miR-6392-3p
GACGCAGGTTTGCCGGGTCCTCT
>mmu-miR-133b-5p
GCTGGTCAAACGGAACCAAGTC
>mmu-miR-677-3p
GAAGCCAGATGCCGTTCCTGAGAAGG
>mmu-miR-7066-5p
TGGGTTGGGAAATGAGTAGAGG
>mmu-miR-6978-3p
ACGGCTTCACTCTCACCCTGCAG
>mmu-miR-466g
ATACAGACACATGCACACACA
>mmu-miR-6969-3p
TGTTGTTCCCATCTCTGCAG
>mmu-miR-669k-5p
TGTGCATGTGTGTATAGTTGTGTGC
>mmu-miR-341-5p
CGGTCGGCCGATCGCTCGGTC
>mmu-miR-3473f
CAAATAGGACTGGAGAGATG
>mmu-miR-5107-5p

TGGGCAGAGGAGGCAGGGACA
>mmu-miR-211-3p
GCAAGGACAGCAAAGGGGGGC
>mmu-miR-8111
ACCGGGCATGGTAGTGTACAC
>mmu-miR-669l-3p
ATATACATACACACCCATATAT
>mmu-miR-92b-5p
AGGGACGGGACGTGGTGCAGTGTT
>mmu-miR-3960
GGCGGCGGCGGAGGCGGGGG
>mmu-miR-297c-5p
ATGTATGTGTGCATGTACATGT
>mmu-miR-7085-5p
CAGTGGGGGCAGGCCAGCAGGA
>mmu-miR-7028-3p
CCTTCTCTTCCCCCTCGGCCAG
>mmu-miR-6965-3p
TCTGTGCTCGGTCTCGCTCCAG
>mmu-miR-8103
TCTCCTGTTCTCTGTTCTCCC
>mmu-miR-7005-5p
CCTGGGGATGGGAGGACCAGCA
>mmu-miR-409-5p
AGGTTACCCGAGCAACTTTGCAT
>mmu-miR-6927-3p
CCTGAGCTGGCTCCCCTGCAG
>mmu-miR-7000-5p
AGGGGACAGGTATCTGGACAGA
>mmu-miR-7002-5p
TTGGCTTCGGGGAGTACGTGG
>mmu-miR-1194
GAATGAGTAACTGCTAGATCCT
>mmu-miR-7080-3p
CAGGCTCACCCCTCCGTTCCCT
>mmu-miR-1247-5p
ACCCGTCCCGTTCGTCCCCGGA
>mmu-miR-6386
CATGCCTTAGGGTGGGATGCAAGT
>mmu-miR-1933-3p
CCAGGACCATCAGTGTGACTAT
>mmu-miR-466j
TGTGTGCATGTGCATGTGTGTAA
>mmu-miR-12202-5p
TGGTGCCTGGATTGGAGGATGAGA
>mmu-miR-695
AGATTGGGCATAGGTGACTGAA
>mmu-miR-7681-5p
ATCCTGTCCTTGCCCTCTCT
>mmu-miR-7048-3p
TCTTCCATTCTCTTTCCCCAG
>mmu-miR-7016-5p
CAGGGAGGGGAGCGAGAGTAG
>mmu-miR-216b-5p
AAATCTCTGCAGGCAAATGTGA
>mmu-miR-7232-5p
TACATGGATGATTGTTTTTCCAAG
>mmu-miR-381-5p
AGCGAGGTTGCCCTTTGTATATT
>mmu-miR-3074-1-3p
GATATCAGCTCAGTAGGCACCG
>mmu-miR-7663-5p

GCTGCTTGGTGATCATCCACTGT
>mmu-miR-1b-3p
TGGGTACATAAAGAAGTATGTGC
>mmu-miR-350-5p
AAAGTGCATGCGCTTTGGG
>mmu-miR-8118
GACAAACATGACTATGCTGACA
>mmu-miR-6379
TCCCTGGAGTTTTACAGCTAGA
>mmu-miR-1964-5p
AGCTGGAGCACAAAAGCCGGTG
>mmu-miR-200b-3p
TAATACTGCCTGGTAATGATGA
>mmu-miR-7008-5p
TGTGGGAGATGCCAGGAGCCGTAC
>mmu-miR-6900-3p
TGGTGATGGGCTCTCTTGTAG
>mmu-miR-6910-3p
TCATGGTTTCCACCCTTTGTCC
>mmu-miR-6907-3p
AACGTGCGTTTCTTTTGTGTCCT
>mmu-miR-199a-5p
CCCAGTGTTCAGACTACCTGTTC
>mmu-miR-6954-3p
TGCAGCCAGCTCTTCCCCTAG
>mmu-miR-194-1-3p
CCAGTGGAGCTGCTGTTACTTC
>mmu-miR-219c-3p
CGAGAATTGCGTTTGGACAATC
>mmu-miR-6375
GAATTGGCTGACTGTATTCCAAA
>mmu-miR-6368
CTGGGAAGCAGTGGAGGGGAG
>mmu-miR-700-3p
CACGCGGGAACCGAGTCCACC
>mmu-miR-493-5p
TTGTACATGGTAGGCTTTC
>mmu-miR-466c-5p
TGATGTGTGTGTGCATGTACATAT
>mmu-miR-5624-3p
TTAAGGCAGAGTTTACAATAGG
>mmu-miR-30b-3p
CTGGGATGTGGATGTTTACGTC
>mmu-miR-8105
AGGCGACTGGCGAGACCCGAAG
>mmu-miR-101b-5p
TCGGTTATCATGGTACCGATGCT
>mmu-miR-7046-5p
TGTAGGGTGAGGCTGGGAGCCAGG
>mmu-miR-5621-5p
AGGAGGTCCTGGGGCCGCCCTGA
>mmu-miR-344e-5p|mmu-miR-344h-5p
CAGGCTTCTGGCTATATTCC
>mmu-miR-881-5p
CAGAGAGATAACAGTCACATCT
>mmu-miR-7115-5p
TCGGGGGTTGTGGTGCCGAATTGT
>mmu-miR-6963-5p
GGTGGGATGGATGGCAGAACCTGG
>mmu-miR-6401
TTACACTCCAGTGGTGTCGGGT
>mmu-miR-719

ATCTCGGCTACAGAAAAATGTT
>mmu-miR-6958-3p
TCAGCTGCCTCTTGCCATCACAG
>mmu-miR-615-3p
TCCGAGCCTGGGTCTCCCTCTT
>mmu-miR-6346
TAGCAGGTGTCTGCAATGCCAA
>mmu-miR-6930-5p
AAGGGGGTGCTTGAGAGGTTTGG
>mmu-miR-471-3p
TGAAAGGTGCCATACTATGTAT
>mmu-miR-1911-3p
CACCAGGCATTGTGGTCTCTGC
>mmu-miR-466e-5p
GATGTGTGTGTACATGTACATA
>mmu-miR-7036b-5p
ACGCGGGGTTCTCTGTGGAAGC
>mmu-miR-3073a-5p
GTGGTCACAGTTGGCGCCAGCC
>mmu-miR-12204-5p
TTGAGTCTTAGATGAGTG
>mmu-miR-7054-3p
TCCAACTTGGCCCCTTCCAG
>mmu-miR-7063-5p
ATGAGAGGTGCAGGCTGAGCACA
>mmu-miR-1933-5p
AGTCATGGTGTTCGGTCTTAGTTT
>mmu-miR-301a-3p
CAGTGCAATAGTATTGTCAAAGC
>mmu-miR-7677-5p
AGTGGTGAGCAGAAAGCAGCCGCGG
>mmu-miR-1896
CTCTCTGATGGTGGGTGAGGAG
>mmu-miR-7214-3p
CTCAGTCCTGACCCCTTGAGCGCA
>mmu-miR-7049-3p
TGCAGCCCCTGTCTGCCTCAG
>mmu-miR-7045-5p
AGCGGGTGGGGGAGGGGGACT
>mmu-miR-6916-5p
CAGGAGGATGGAGATGAGTCA
>mmu-miR-7030-5p
TGGGGAGGTGGCTGTGCTAGTC
>mmu-miR-3057-3p
TCCCACAGGCCCAGCTCATAGC
>mmu-miR-7229-3p
TACACAGACCAGTGACTTTCTGCA
>mmu-miR-592-3p
TCATCACGTGGTGACGCAACAT
>mmu-miR-7044-3p
ATGCAGCCCCGACCCTCACAG
>mmu-miR-6996-5p
TGCACAGGACAGAGCACAGTC
>mmu-miR-674-5p
GCACTGAGATGGGAGTGGTGTA
>mmu-miR-15a-3p
CAGGCCATACTGTGCTGCCTCA
>mmu-let-7f-5p
TGAGGTAGTAGATTGTATAGTT
>mmu-miR-7675-5p
CTTACGAAAGAATGAAGAGCAA
>mmu-miR-10a-3p

CAAATTCGTATCTAGGGGAATA
>mmu-miR-3074-2-3p
TGTTTCAGCTCAGTAGGCAC
>mmu-miR-3075-5p
TGTCTGGGAGCAGCCAAGGAC
>mmu-miR-1187
TATGTGTGTGTGTATGTGTGTAA
>mmu-miR-7033-3p
TGCCTCCATTCTCTTGGGCAG
>mmu-miR-6241
CACGGCGGCTGGAATTCCC
>mmu-miR-5134-5p
TTGGCAGAAAGGGCAGCTGTG
>mmu-miR-3472
TAATAGCCAGAAGCTGGAAGGAACC
>mmu-miR-12201-3p
TCTGCAGAGGCTTCCTTGGCT
>mmu-miR-337-3p
TCAGCTCCTATATGATGCCTTT
>mmu-miR-223-5p
CGTGTATTTGACAAGCTGAGTTG
>mmu-miR-7012-3p
TGACCTGTGGCTCCTCTCCCAG
>mmu-miR-6538
CGCGGGCTCCGGGGCGGCG
>mmu-miR-191-5p
CAACGGAATCCCAAAAGCAGCTG
>mmu-miR-12185-5p
CTTGGGGGCAGAGGTGGCACGGTGT
>mmu-miR-743b-3p
GAAAGACATCATGCTGAATAGA
>mmu-miR-3062-5p
GGAGAATGTAGTGTTACCGTGA
>mmu-miR-669f-5p
AGTTGTGTGTGCATGTGCATGTGT
>mmu-miR-378a-5p
CTCCTGACTCCAGGTCCTGTGT
>mmu-miR-6909-5p
CAGGGGGGCTGGAATGGGCACT
>mmu-miR-1231-3p
TGCCCTGTCTGTTCTGCCCACAG
>mmu-miR-1249-5p
AGGAGGGAGGGGATGGGCCAAGTTC
>mmu-miR-6942-5p
TAGGAGGAGGGGAACATTTGC
>mmu-miR-141-3p
TAACACTGTCTGGTAAAGATGG
>mmu-miR-181b-5p
AACATTCATTGCTGTCGGTGGGTT
>mmu-miR-703
AAAACCTTCAGAAGGAAAGAA
>mmu-miR-27a-5p
AGGGCTTAGCTGCTTGTGAGCA
>mmu-miR-137-5p
ACGGGTATTCTTGGGTGGATAAT
>mmu-miR-3108-3p
CGTCTAGGTCTAGAGTCTCAA
>mmu-miR-7224-3p
TCCACTGAGAGGACCACCCAC
>mmu-miR-3535
TGGATATGATGACTGATTACCTGAGA
>mmu-miR-883b-5p

TACTGAGAATGGGTAGCAGTCA
>mmu-miR-1943-5p
AAGGGAGGATCTGGGCACCTGGA
>mmu-miR-7057-5p
CAGGAGAGGACTCAAGTT
>mmu-miR-6968-3p
ACAAGCGCTGTCTCCCCTCCAG
>mmu-miR-1907
GAGCAGCAGAGGATCTGGAGGT
>mmu-miR-6415
GAGCTTTTGAGGGCCCCATGCA
>mmu-miR-301b-3p
CAGTGCAATGGTATTGTCAAAGC
>mmu-miR-12191-5p
ACCGTGGGGCGAACGGAGCG
>mmu-miR-6371
TTCAAGTGGCTTGAAGCTCTTCT
>mmu-miR-3093-5p
CGCACCCCGCGGAGCTCACACT
>mmu-miR-7219-5p
TGTGTTAGAGCTCAGGGTTGAGA
>mmu-miR-7057-3p
TAATTTGATGTCTTTCTCCACC
>mmu-miR-153-5p
GTCATTTTTGTGACGTTGCAGCT
>mmu-miR-3099-5p
CCAGCTTCCTTCCAGCCCTTG
>mmu-miR-3074-5p
GTTCCTGCTGAACTGAGCCAGT
>mmu-miR-26a-1-3p
CCTATTCTTGGTTACTTGCACG
>mmu-miR-7021-5p
ATAGGAGTTGATTACAGAGCAA
>mmu-miR-6960-3p
AGCCAACCACCTCTCTCCTCAG
>mmu-miR-148b-3p
TCAGTGCATCACAGAACTTTGT
>mmu-miR-3103-3p
TAACCTCTGATCCTTCCCACAG
>mmu-miR-486a-5p|mmu-miR-486b-5p
TCCTGTACTGAGCTGCCCCGAG
>mmu-miR-30f
GTAAACATCCGACTGAAAGCTC
>mmu-miR-208a-5p
GAGCTTTTGGCCCGGGTTATAC
>mmu-miR-761
GCAGCAGGGTGAAACTGACACA
>mmu-miR-344b-5p
AGTCAGGCTCCTGGCTAAAGTTC
>mmu-miR-7660-3p
ATCTGTAAGTGATGGGCAAAGTA
>mmu-miR-9b-5p
TTCGGTTATCTAGCTTTATGA
>mmu-miR-302c-3p
AAGTGCTTCCATGTTTCAGTGG
>mmu-miR-199b-5p
CCCAGTGTTTAGACTACCTGTTC
>mmu-miR-6975-5p
GCTGGGGAGAAAGGGGTTTGGCA
>mmu-miR-6979-5p
GGGGAGGCGCAGAGACTGAG
>mmu-miR-883a-5p

TGCTGAGAGAAGTAGCAGTTAC
>mmu-miR-669n
ATTTGTGTGTGGATGTGTGT
>mmu-miR-23b-3p
ATCACATTGCCAGGGATTACC
>mmu-miR-202-5p
TTCCTATGCATATACTTCTTT
>mmu-miR-12182-3p
TAATTAGGCTCTGGATCTGTGA
>mmu-miR-7079-5p
AGGGCTGAGGCAGTGAGTCCTG
>mmu-miR-2139
AGCTGCGCTGCTCCTGGTAACTGC
>mmu-miR-421-5p
CTCATTAAATGTTTGTTGAAT
>mmu-miR-721
CAGTGCAATTAAAAGGGGGAA
>mmu-miR-1249-3p
ACGCCCTTCCCCCCCTTCTTCA
>mmu-miR-327
ACTTGAGGGGCATGAGGAT
>mmu-miR-3062-3p
CGGGGACACACTTTCTCCT
>mmu-miR-1a-2-5p
ACATACTTCTTTATGTACCCATA
>mmu-miR-669a-3-3p
ACATAACATACACACACATGTAT
>mmu-miR-7234-3p
AAACGTCTTTCTAGGGTAGAAGG
>mmu-miR-6996-3p
CGGTGTCTCTGGTCACTCTGCAG
>mmu-miR-100-3p
ACAAGCTTGTGTCTATAGGTAT
>mmu-miR-542-5p
CTCGGGGATCATCATGTCACGA
>mmu-miR-6938-5p
TTGGAGCTAGCTCCAGAGAGAGT
>mmu-miR-3076-3p
CGCACTCTGGTCTTCCCTTGCAG
>mmu-miR-485-3p
AGTCATACACGGCTCTCCTCTC
>mmu-miR-7676-5p
AGGGCAGTATGATGGCCTCTGAT
>mmu-miR-99b-5p
CACCCGTAGAACCGACCTTGCG
>mmu-miR-701-5p
TTAGCCGCTGAAATAGATGGA
>mmu-miR-192-5p
CTGACCTATGAATTGACAGCC
>mmu-miR-6941-3p
CTAACTCGGCTGCTCCCAAAG
>mmu-miR-3080-3p
TCCTCGGGCAAAGCGCTTGACA
>mmu-miR-7007-3p
CCCATCCACGTTTCTTCT
>mmu-miR-7211-5p
TCTTTCCCTCTGCCACTCCACC
>mmu-miR-7047-3p
TCACCCTCACCCTTCCTCCTTCC
>mmu-miR-7008-3p
TGTGCTTCTTGCCTCTTCTCAG
>mmu-miR-6913-5p

TGGGAGACCAGTCAGGTGTTGG
>mmu-miR-1843a-3p
TCTGATCGTTCACCTCCATACA
>mmu-miR-6923-3p
ACACTCCCTCCTCCTCCCCAG
>mmu-miR-465b-5p
TATTTAGAATGGTGCTGATCTG
>mmu-miR-1306-5p
CACCACCTCCCCTGCAAACGTCC
>mmu-miR-463-5p
TACCTAATTTGTTGTCCATCAT
>mmu-miR-126a-5p
CATTATTACTTTTGGTACGCG
>mmu-miR-93-5p
CAAAGTGCTGTTCGTGCAGGTAG
>mmu-miR-465a-5p
TATTTAGAATGGCACTGATGTGA
>mmu-miR-672-5p
TGAGGTTGGTGTACTGTGTGTGA
>mmu-miR-3106-3p
GCTGCCTATACATGGGCTTTCC
>mmu-miR-382-3p
TCATTCACGGACAACACTTTTT
>mmu-miR-101a-3p
TACAGTACTGTGATAACTGAA
>mmu-miR-7027-3p
TCGGGTTTCTCTCCTCTTCCAG
>mmu-miR-7011-5p
AGGAGGATGGGAGAGGGAGGTGT
>mmu-miR-6361
TGGCTCAGCAGTCACCCTGTGAC
>mmu-miR-12192-5p
TGTGGGGTATAGCTAGAAAGA
>mmu-miR-376a-3p
ATCGTAGAGGAAAATCCACGT
>mmu-miR-874-3p
CTGCCCTGGCCCGAGGGACCGA
>mmu-miR-7088-3p
TTGACCTTCCTCCATTGCTTCC
>mmu-miR-6978-5p
AAGGGGTGAGAGAGAAGCTGGTG
>mmu-miR-1a-1-5p
ACATACTTCTTTATATGCCCATA
>mmu-miR-7089-5p
CCGTGGGACCTACAGACTCCTGGG
>mmu-miR-425-3p
ATCGGGAATGTCGTGTCCGCC
>mmu-miR-1952
TCTCCACCCTCCTTCTG
>mmu-miR-6990-3p
AGCCCTGCCTCTTCCTGGCAG
>mmu-miR-188-5p
CATCCCTTGCATGGTGGAGGG
>mmu-miR-1932
GTTGCGGACAGCGCTAGGTCGG
>mmu-miR-3102-5p
GTGAGTGGCCAGGGTGGGGCTG
>mmu-miR-9769-3p
TAGGGTCTGTTCTGTGTCCTCC
>mmu-miR-183-3p
GTGAATTACCGAAGGGCCATAA
>mmu-miR-6929-5p

AGGGAGGAGCAGCATCTGTGA
>mmu-miR-326-5p
GGGGGCAGGGCCTTTGTGAAGGCG
>mmu-miR-6339
GATGCCTTTGTCCTCAGTGTTAG
>mmu-miR-3474
CCCTGGGAGGAGACGTGGATTC
>mmu-miR-3104-3p
ACGCTCTGCTTTGCTCCCCCAGA
>mmu-miR-6383
TAAAGTGCTGTGTCACGGACA
>mmu-miR-7084-5p
TAGAGGATAGAGGTAGAGAGT
>mmu-miR-5128
CAATTGGGGCTGGCGAGATGGCT
>mmu-miR-302d-3p
TAAGTGCTTCCATGTTTGAGTGT
>mmu-miR-1941-3p
CATCTTAGCAGTATCTCCCAT
>mmu-miR-7068-3p
TCACCCTGGACTGACTCTCAG
>mmu-miR-669e-3p
TGAATATACACACACTTACAC
>mmu-miR-3058-3p
TTCCTGTCAGCCGTGGGTGCC
>mmu-miR-7009-5p
TTGGGGTCAGGGGACCAGAGCT
>mmu-miR-7072-5p
GAGGGGAGACAAGAGGCAAGGC
>mmu-miR-6952-3p
TCTCTGACTCTGCCTCCCACAG
>mmu-miR-6400
TTCTTGCTGCTTGGTGCTCGC
>mmu-miR-1668
CAAAGGCCTCCTTCTCTGCAG
>mmu-miR-7073-5p
TGGGGGAATTGAAAGACTATGA
>mmu-miR-3112-3p
CAGTTTGTCTCTTCTTTGTCT
>mmu-miR-129-1-3p
AAGCCCTTACCCCAAAAAGTAT
>mmu-miR-106a-3p
ACTGCAGTGCCAGCACTTCTTAC
>mmu-miR-664-5p
CTGGCTGGGGAAAATGACTGG
>mmu-miR-6391
TGAGGCACGGGAGGACAATGT
>mmu-miR-3058-5p
TCAGCCACGGCTTACCTGGAAGA
>mmu-miR-467b-3p
ATATACATACACACACCAACAC
>mmu-miR-3061-3p
CTACCTTTGATAGTCCACTGCC
>mmu-miR-3083-5p
AGGCTGGGAATATTTCAGAGAT
>mmu-miR-1982-3p
TCTCACCCTATGTTCTCCCACAG
>mmu-miR-7662-3p
TGGAGCCAGGCGGATCAGCCTTGC
>mmu-miR-6944-5p
GTGAGAGCGGGGGGAGTGGCAAG
>mmu-miR-876-3p

TAGTGGTTTACAAAGTAATTCA
>mmu-miR-670-5p
ATCCCTGAGTGTATGTGGTGAA
>mmu-miR-3109-3p
TAGGGCCATCTCATCCAGATA
>mmu-miR-12205-3p
AAACCAACTGGGAAACACAAAT
>mmu-miR-185-5p
TGGAGAGAAAGGCAGTTCCTGA
>mmu-miR-6938-3p
TCATCTGGGGCTGTCTCCTTAG
>mmu-let-7e-3p
CTATACGGCCTCCTAGCTTTCC
>mmu-miR-1904
GTTCTGCTCCTCTGGAGGGAGG
>mmu-miR-99a-3p
CAAGCTCGTTTCTATGGGTCT
>mmu-miR-326-3p
CCTCTGGGCCCTTCCTCCAGT
>mmu-let-7b-3p
CTATACAACCTACTGCCTTCCC
>mmu-miR-6942-3p
TTATGGTTCTCTCCACCTCTCA
>mmu-miR-7067-5p
TGCAGGGGGTGGTTGCACACGGA
>mmu-miR-3475-5p
AAATCATGTACCCCCACAGAGG
>mmu-miR-6991-3p
GCAGCCCTTTGTTCTTCTCCAG
>mmu-miR-8113
CAGGAGAGTCAGGGGCAAGTAG
>mmu-miR-28c
AGGAGCTCACAGTCTATTGA
>mmu-miR-3091-3p
CGGGCCTGACCAGTCTCAAGAC
>mmu-miR-365-2-5p
AGGGACTTTCAGGGGCAGCTGTG
>mmu-miR-698-5p
TGTGGGTGGGACAGGGATGTT
>mmu-miR-6904-5p
TCCTGGGGTTAGAGTTGAGTGG
>mmu-miR-6918-5p
TGCTGAGGACGGGATTAGGTTCT
>mmu-miR-122-3p
AAACGCCATTATCACACTAAAT
>mmu-miR-433-5p
TACGGTGAGCCTGTCATTATTC
>mmu-miR-6900-5p
TGCCAGGAGAAGCCTAGAGCCGTC
>mmu-miR-669i
TGCATATACACACATGCATAC
>mmu-miR-1938
CGGTGGGACTTGTAGTTCGGTC
>mmu-miR-1839-5p
AAGGTAGATAGAACAGGTCTTG
>mmu-miR-204-5p
TTCCCTTTGTCATCCTATGCCT
>mmu-miR-101b-3p
GTACAGTACTGTGATAGCT
>mmu-miR-1946b
GCCGGGCAGTGGTGGCACATGCTTTT
>mmu-miR-3103-5p

GGAGGGAGGATCTGCTGTTAG
>mmu-miR-129b-5p
GCTTTTTGGGGTAAGGGCTTCC
>mmu-miR-1893
GGCGCGGGCGCTGGACGCCTCG
>mmu-miR-669m-5p|mmu-miR-466m-5p
TGTGTGCATGTGCATGTGTGTAT
>mmu-miR-20a-5p
TAAAGTGCTTATAGTGCAGGTAG
>mmu-miR-6914-5p
TCCTGGGGTGGTGGTGGCCACAGA
>mmu-miR-291b-5p
GATCAAAGTGGAGGCCCTCTCC
>mmu-miR-1892
ATTTGGGGACGGGAGGGAGGAT
>mmu-miR-24-1-5p
GTGCCTACTGAGCTGATATCAGT
>mmu-miR-29c-5p
TGACCGATTTCTCCTGGTGTTC
>mmu-miR-7090-3p
TTTGATGTCTGACTTTGCAG
>mmu-miR-669d-3p
TATACATACACACCCATATAC
>mmu-miR-19b-1-5p
AGTTTTGCAGGTTTGCATCCAGC
>mmu-miR-7058-3p
CTCGTTCCTTCCTTTCTTCCAG
>mmu-miR-7076-3p
TGACTACCACTGTCTCCCCAG
>mmu-miR-196b-3p
TCGACAGCACGACACTGCCTTC
>mmu-miR-6905-5p
ACTGGGCAGGTTGGGTTGAATGA
>mmu-miR-302c-5p
GCTTTAACATGGGGTTACCTGC
>mmu-miR-7010-3p
AGGTTCTCCTTTCCTTTGCAG
>mmu-miR-340-3p
TCCGTCTCAGTTACTTTATAGC
>mmu-miR-374b-3p
GGTTGTATTATCATTGTCCGAG
>mmu-miR-9b-3p
ATACAGCTAGATAACCAAAGA
>mmu-miR-206-3p
TGGAATGTAAGGAAGTGTGTGG
>mmu-miR-1951
GTAGTGGAGACTGGTGTGGCTA
>mmu-miR-1191b-3p
AGACTCACTATGTAGCCCAAGC
>mmu-miR-709
GGAGGCAGAGGCAGGAGGA
>mmu-miR-6956-5p
TGAGTGGGTGGGACTGGCCT
>mmu-miR-130a-3p
CAGTGCAATGTTAAAAGGGCAT
>mmu-miR-686
ATTGCTTCCCAGACGGTGAAGA
>mmu-miR-3094-3p
CCTTTAAATTGTGTCCTCAAG
>mmu-miR-6389
CAGTGCAATGTTAAACTTTGC
>mmu-miR-12199-3p

ACACTGGTTCTCTCTGGCTCT
>mmu-miR-7665-3p
AGCCAGGTCCCTGTCCCCTACA
>mmu-miR-3073b-3p
CTGGCGCCAACTGTGACCACTG
>mmu-miR-3076-5p
CACAGGGGAAGCTCAGTGCCAGCC
>mmu-miR-680
GGGCATCTGCTGACATGGGGG
>mmu-miR-7091-5p
GTAGGGGTGATAGATTCCATGGC
>mmu-miR-7060-3p
TCTACTCTACCTTCTACTCAG
>mmu-miR-3069-3p
TTGGACACTAAGTACTGCCACA
>mmu-miR-12191-3p
CCCATGGAGCTGTAGGAGCCG
>mmu-miR-3965
TGCTTATCAGCCTGATGTT
>mmu-miR-3063-5p
TGGAGATCAGGTTTGCACACT
>mmu-miR-6975-3p
TCTCTCCTTTCTCCTCCTAG
>mmu-miR-3079-5p
TTTGATCTGATGAGCTAAGCTGG
>mmu-miR-7226-3p
TGACACAGCCATTCTCTGAGCAG
>mmu-miR-6995-3p
TGTGTCCCCTTCCTCTCACAG
>mmu-miR-6934-5p
TTGGTGGAGACAGCAAGGGTGG
>mmu-miR-8098
GCTATGGTCGCTGGCTCCCAA
>mmu-miR-6420
ACTAATCCTATAAAATCAAAC
>mmu-miR-6405
GTGGGAAGCAGTGGGAAGCAGTG
>mmu-miR-3963
TGTATCCCACTTCTGACAC
>mmu-miR-763
CCAGCTGGGAAGAACCAGTGGC
>mmu-miR-742-5p
TACTCACATGGTTGCTAATCA
>mmu-miR-764-3p
AGGAGGCCATAGTGGCAACTGT
>mmu-miR-1954
ACTGCAGAGTGAGACCCTGTT
>mmu-miR-5103
TCATCCGGGATCCCCTGAGG
>mmu-miR-669g
TGCATTGTATGTGTTGACATGAT
>mmu-miR-141-5p
CATCTTCCAGTGCAGTGTTGGA
>mmu-miR-6931-3p
CCCAACCTCTTCATACCCCAG
>mmu-miR-21a-5p
TAGCTTATCAGACTGATGTTGA
>mmu-miR-6944-3p
TAACTCTTCCCTTGTGCCCTCAG
>mmu-miR-1963
TGGGACGAGATCATGAGGCCTTC
>mmu-miR-3473e

GGGCTGGAGAGATGGCTCGTA
>mmu-miR-5615-5p
CTTGGTTGTTTTCTGAGACAGA
>mmu-miR-3473c
TCTCTCCAGCCCCCATAATAAG
>mmu-miR-32-3p
CAATTTAGTGTGTGTGATATT
>mmu-miR-7678-3p
TTTCCTTACTTTCTCTTGTAGC
>mmu-miR-218-5p
TTGTGCTTGATCTAACCATGT
>mmu-miR-713
TGCACTGAAGGCACACAGC
>mmu-miR-504-3p
AGGGAGAGCAGGGCAGGGTTTC
>mmu-miR-181b-1-3p
CTCACTGAACAATGAATGCAA
>mmu-miR-1941-5p
AGGGAGATGCTGGTACAGAGGCTT
>mmu-miR-7a-5p
TGGAAGACTAGTGATTTTGTTGT
>mmu-miR-3067-3p
CCAAGCGGCTGCCCTGGGAGAGG
>mmu-miR-7239-3p
TGGCTCTGTCAGGCAGGTAG
>mmu-miR-195a-5p
TAGCAGCACAGAAATATTGGC
>mmu-miR-139-5p
TCTACAGTGCACGTGTCTCCAG
>mmu-miR-19a-3p
TGTGCAAATCTATGCAAAACTGA
>mmu-miR-493-3p
TGAAGGTCCTACTGTGTGCCAGG
>mmu-miR-7217-5p
AACTTGTATCTTGTGAGACAGAAGG
>mmu-miR-203b-3p
TTGAACTGTCAAGAACCACTGG
>mmu-miR-6237
TTAAGGATTGGGTCTTGGAATTC
>mmu-miR-6924-3p
TCCTTAATCCCCACCCTTCAG
>mmu-miR-412-5p
TGGTCGACCAGCTGGAAAGTAAT
>mmu-miR-7036b-3p
TTCCTGGTGGGCCTCGGGGTCC
>mmu-miR-107-5p
AGCTTCTTTACAGTGTTGCCTTG
>mmu-miR-7092-3p
TGTTCTTTGTTTTGTTTGCCTTAG
>mmu-miR-6962-3p
TATCTGCCCTTCCCTGTCCTAT
>mmu-miR-6377
TCATTTGTCTTCGTGTCTGCATC
>mmu-miR-134-3p
CTGTGGGCCACCTAGTCACC
>mmu-miR-187-3p
TCGTGTCTTGTGTTGCAGCCGG
>mmu-miR-762
GGGGCTGGGGCCGGGACAGAGC
>mmu-miR-544-5p
TCTTGTTAAAAAGCAGAGTCT
>mmu-miR-501-5p

AATCCTTTGTCCCTGGGTGAAA
>mmu-miR-5108
GTAGAGCACTGGATGGTTT
>mmu-miR-7041-5p
AGAGGGAATGGAGACTCAGTTGA
>mmu-miR-6927-5p
GTGAGGGGATCCAGCCCAGGCT
>mmu-miR-1193-3p
TAGGTCACCCGTTTTACTATC
>mmu-miR-7654-3p
CGAGCGGGAGCGCGCCTCGTCC
>mmu-miR-6240
CCAAAGCATCGCGAAGGCCCACGGCG
>mmu-miR-7083-5p
TCGGGGCTGGACAAGCAGAGA
>mmu-miR-7081-5p
AGAGGAGGGTGCTCGGCCGGGGATTA
>mmu-miR-5135
AGGTCTAGGTGGCAAGGGCGTCCT
>mmu-miR-135a-1-3p
TATAGGGATTGGAGCCGTGGCG
>mmu-miR-1956
AGTCCAGGGCTGAGTCAGCGGA
>mmu-miR-9768-3p
ACTGCCTTCCTTTGTGTGGCCCAG
>mmu-miR-3072-5p
AGGGACCCCGAGGGAGGGCAGG
>mmu-miR-7052-5p
TGTGGGAAGGTGGAGGCCTTT
>mmu-miR-6984-3p
TACTTTCTTTCCTGTCTTTCT
>mmu-miR-687
CTATCCTGGAATGCAGCAATGA
>mmu-miR-341-3p
TCGGTCGATCGGTCGGTCGGT
>mmu-miR-6956-3p
TGACCGGCCTATCCTCTCAG
>mmu-miR-1231-5p
TCTGGGCAGAGCTGCAGGAGAGA
>mmu-miR-411-5p
TAGTAGACCGTATAGCGTACG
>mmu-miR-6366
AGCTAAGGGGCCCGGGGAGCCA
>mmu-miR-702-5p
GTGAGTGGGGTGGTTGGCATG
>mmu-miR-501-3p
AATGCACCCGGGCAAGGATTTG
>mmu-miR-8104
CAGGATGAGGTGGAGTGCAGCG
>mmu-miR-873b
ACAAGTTCCTGCAAATGCACAC
>mmu-miR-129-5p
CTTTTTGCGGTCTGGGCTTGC
>mmu-miR-3070-3p
TGGTGCTACCGTCAGGGGTAGA
>mmu-miR-6902-3p
CCATGTGATGTGTGGGTTCAG
>mmu-miR-7669-5p
AGTACCACCATACACAGCTTTTG
>mmu-miR-7661-5p
AAGAAAGAAACCTGGAGTTTAAAGT
>mmu-miR-5709-5p

ATACGCACACTTAAGACTTTAG
>mmu-miR-7031-3p
AACCCTCTTGCCCTCTCCTAG
>mmu-miR-6961-5p
TGACTGGGAAGGAAATGGCC
>mmu-miR-344d-1-5p
AGTCAGGCTGCTGGCTATACACCA
>mmu-miR-146b-5p
TGAGAACTGAATTCCATAGGCT
>mmu-miR-122-5p
TGGAGTGTGACAATGGTGTTTG
>mmu-miR-5120
TTTGGGGCTGTGGTGCCACCAGC
>mmu-miR-702-3p
TGCCCACCCTTTACCCCGCTCC
>mmu-miR-6394
TCCCTGAGTGGGGCCAGGTCT
>mmu-miR-1188-3p
TCCGAGGCTCCCCACCACACCCTGC
>mmu-miR-320-3p
AAAAGCTGGGTTGAGAGGGCGA
>mmu-miR-7649-3p
TCTCTCCAGCACTGCTTGTTTCT
>mmu-miR-23a-3p
ATCACATTGCCAGGGATTTCC
>mmu-miR-212-3p
TAACAGTCTCCAGTCACGGCCA
>mmu-miR-6957-5p
CAGAAGAGAGACAGGATCTAG
>mmu-miR-15b-3p
CGAATCATTATTTGCTGCTCTA
>mmu-miR-669a-3p|mmu-miR-669o-3p
ACATAACATACACACACACGTAT
>mmu-miR-6992-3p
TCTCACTGATGATCATTTGCAG
>mmu-miR-7026-5p
TTCTGAGACCATGGGGTATAT
>mmu-miR-6381
CAGGATGCTGGTGACATGAGT
>mmu-miR-181a-1-3p
ACCATCGACCGTTGATTGTACC
>mmu-miR-1b-5p
TACATACTTCTTTACATTCCA
>mmu-miR-7065-5p
AGTGTAGCAGGTAAACCAGGA
>mmu-miR-6406
CGCGACCTCAGGCTGACTCATG
>mmu-miR-7211-3p
TGGAGTGACTGTAGGGAGGATGC
>mmu-miR-6913-3p
TCTCTACTGATTTGTCTCCTCAG
>mmu-miR-155-3p
CTCCTACCTGTTAGCATTAAC
>mmu-miR-12200-5p
AAACAAACCAGAGGCTCACACT
>mmu-miR-669c-3p
TACACACACACACACAAGTAAA
>mmu-miR-421-3p
ATCAACAGACATTAATTGGGCGC
>mmu-miR-8099
CCAAGGCGTGGGAAAGGTTGT
>mmu-miR-669a-5p|mmu-miR-669p-5p

AGTTGTGTGTGCATGTTCATGTCT
>mmu-miR-1306-3p
ACGTTGGCTCTGGTGGTGATG
>mmu-miR-448-3p
TTGCATATGTAGGATGTCCCAT
>mmu-miR-509-5p
TACTCCAGAATGTGGCAATCAT
>mmu-miR-300-5p
TTGAAGAGAGGTTATCCTTTGT
>mmu-miR-3065-3p
TCAGCACCAGGATATTGTTGGGG
>mmu-miR-6917-5p
TGTGGAAGGGGAGTTGTGCCG
>mmu-miR-5616-5p
TTTCCTCTCATCACAAGTTGA
>mmu-miR-463-3p
TGATAGACACCATATAAGGTAG
>mmu-miR-466q
GTGCACACACACACATACGT
>mmu-miR-210-3p
CTGTGCGTGTGACAGCGGCTGA
>mmu-miR-6920-3p
CACTGCCCTTTTCCATTGCCT
>mmu-miR-3110-5p
TTCTGCCTCCCCTGAAGGCTC
>mmu-miR-7039-3p
CCTGTTCTGCCCTCCCTTCCAG
>mmu-miR-6920-5p
ACACAATGGAAAGACTGCTTGT
>mmu-miR-133b-3p
TTTGGTCCCCTTCAACCAGCTA
>mmu-miR-1970c-3p
AAGCCGGGCCTAGTGACTCTCT
>mmu-miR-3064-3p
TGCCACACTGCAACACCTTACA
>mmu-miR-203b-5p
AGTGGTCCTAAACATTTCAC
>mmu-miR-3080-5p
TGAAGCGCCTGTTCTTGGAT
>mmu-miR-7087-5p
AGGCAGGTGTGGAGCTGGTCT
>mmu-miR-142b
TCCATAAAGTAGGAAACACT
>mmu-miR-153-3p
TTGCATAGTCACAAAAGTGATC
>mmu-miR-483-5p
AAGACGGGAGAAGAGAAGGGAG
>mmu-miR-7242-3p
TGGCAGCTCGGCTCTACTCTGG
>mmu-miR-1967
TGAGGATCCTGGGGAGAAGATGC
>mmu-miR-6350
TAGCAGGTGCATTATAAATAT
>mmu-miR-3620-5p
CTGTGGGCTGGGCTGGGAAGCA
>mmu-miR-6537-5p
GTGAGTTTCTCCCACTGGTGAC
>mmu-miR-29a-3p
TAGCACCATCTGAAATCGGTTA
>mmu-miR-1927
GACCTCTGGATGTTAGGGACTGA
>mmu-miR-3970

GAGGTTGTAGTTTGTGCTTT
>mmu-miR-665-5p
AGGGGCCTCTGCCTCTATCCAGGATT
>mmu-miR-193b-3p
AACTGGCCCACAAAGTCCCGCT
>mmu-miR-378c
ACTGGACTTGGAGTCAGAAGC
>mmu-miR-150-5p
TCTCCCAACCCTTGTACCAGTG
>mmu-miR-7666-3p
GATGCAGCGCACGGGCGAG
>mmu-miR-6418-5p
TCAGGGGAAGGGAAGAGATGC
>mmu-miR-6954-5p
TGGGGCAGTTCTGGGGGCAGAT
>mmu-miR-6715-5p
ACAGGCACCACAGGTTTGAGCAT
>mmu-miR-7671-3p
AAGCGGGCAGGCGGAGAACCAC
>mmu-miR-7212-5p
TCTGGGGGCTTGTGTGGTAGG
>mmu-miR-145b
GTCCAGTTTTCCCAGGAGACT
>mmu-miR-3075-3p
ATCCTTGGCCTTCCTAGGTGT
>mmu-miR-7004-5p
TTCCGTGGATGAGGCCTGTGCCGT
>mmu-miR-148a-3p
TCAGTGCACTACAGAACTTTGT
>mmu-miR-7044-5p
GTGTGGTGGTGGTGGCGGC
>mmu-miR-5131
CGGCGCCCCACGGAGCCCCGAGC
>mmu-miR-6344
GTTTTCCTACTGTTTCCCTTTT
>mmu-miR-6943-3p
CTGCTTCCATACCCCTACCCAG
>mmu-miR-3618-5p
TGTGATTTCCAATAATTGAGGC
>mmu-miR-6546-5p
AGGCGCCTCTGAGCTTGTGCTTGT
>mmu-miR-7233-5p
AGTTAGGGACAGATAGATG
>mmu-miR-7226-5p
GCCAGGGAAGTTGATTGTGTGAAGGG
>mmu-miR-6899-3p
TTGTCCTTCTGTGTCTTCTGCAG
>mmu-miR-210-5p
AGCCACTGCCCACCGCACACTG
>mmu-miR-429-5p
GTCTTACCAGACATGGTTAGA
>mmu-miR-7036a-5p
AGCGGGGTTCGGTGGGGAAGAGA
>mmu-miR-5619-3p
CGAAAAGCTCATTGGTGATTTC
>mmu-miR-7035-5p
GGCTGGGAGAGTGGCTCAGAA
>mmu-miR-30c-1-3p
CTGGGAGAGGGTTGTTTACTCC
>mmu-miR-6983-3p
TGAATCAAAGTCTGTTGTTCCAG
>mmu-miR-7032-3p

ATCCTCTCGGTACCGCCCTGCA
>mmu-miR-6392-5p
TCTGGCGCCCTGGTCCTGTCAGC
>mmu-miR-93-3p
ACTGCTGAGCTAGCACTTCCCG
>mmu-miR-7681-3p
AGAAAGGGCACTGACAGGATAG
>mmu-let-7c-5p
TGAGGTAGTAGGTTGTATGGTT
>mmu-miR-7093-5p
CAGGATGACAGAAGGAAAACCT
>mmu-miR-7040-3p
CGCCACCCTCTCTCTCACGTAG
>mmu-miR-9769-5p
AGGCACAGAATTAGGCCCGACC
>mmu-miR-7231-5p
TTGGGGAACACTGGGGCATACC
>mmu-miR-8114
TCACCCATCTCCTCTCCGCCT
>mmu-miR-6397
TGGAGACTTTCAGTGGAGTGA
>mmu-miR-676-3p
CCGTCCTGAGGTTGTTGAGCT
>mmu-miR-6982-3p
TGGCCCCTCTGCCCCTTCCAG
>mmu-miR-7022-3p
ACAAGCCTGACCTCTGCCCCCA
>mmu-miR-384-3p
ATTCCTAGAAATTGTTCACAAT
>mmu-miR-34a-5p
TGGCAGTGTCTTAGCTGGTTGT
>mmu-miR-707
CAGTCATGCCGCTTGCCTACG
>mmu-miR-877-3p
TGTCCTCTTCTCCCTCCTCCCA
>mmu-miR-5113
ACAGAGGAGGAGAGAGATCCTGT
>mmu-miR-125b-2-3p
ACAAGTCAGGTTCTTGGGACCT
>mmu-miR-1960
CCAGTGCTGTTAGAAGAGGGCT
>mmu-miR-1903
CCTTCTTCTTCTTCCTGAGACA
>mmu-miR-12200-3p
TGTGAATCTCCGGCGCGTTT
>mmu-miR-148a-5p
AAAGTTCTGAGACACTCCGACT
>mmu-miR-7220-3p
CCAGGAACAAGGTCTCAGTCCGA
>mmu-miR-200b-5p
CATCTTACTGGGCAGCATTGGA
>mmu-miR-7a-2-3p
CAACAAGTCCCAGTCTGCCACA
>mmu-miR-136-3p
ATCATCGTCTCAAATGAGTCTT
>mmu-miR-3967
AGCTTGTCTGACTGATGTTG
>mmu-miR-192-3p
CTGCCAATTCCATAGGTCACAG
>mmu-miR-5130
CTGGAGCGCGCGGGCGAGGCAGGC
>mmu-miR-1191a

CAGTCTTACTATGTAGCCCTA
>mmu-miR-7685-3p
AGACTTGGCTTCCGGCTGGAG
>mmu-miR-486b-3p
CGGGGCAGCTCAGTACAGGA
>mmu-miR-6949-5p
AAAGTGGGTCTAGAAAGAGGTGA
>mmu-miR-196a-2-3p
TCGGCAACAAGAAACTGCCTGA
>mmu-miR-25-5p
AGGCGGAGACTTGGGCAATTGC
>mmu-miR-7243-5p
TGCTGTCAGCTGTTCTGAGT
>mmu-miR-470-5p
TTCTTGGACTGGCACTGGTGAGT
>mmu-miR-330-3p
GCAAAGCACAGGGCCTGCAGAGA
>mmu-miR-21c
TAGCTTATCAGACTGGTACAA
>mmu-miR-1198-5p
TATGTGTTCCTGGCTGGCTTGG
>mmu-miR-6977-3p
AAGGCGTTGCCTGACCCTGACC
>mmu-miR-139-3p
TGGAGACGCGGCCCTGTTGGAG
>mmu-miR-3470b
TCACTCTGTAGACCAGGCTGG
>mmu-miR-694
CTGAAAATGTTGCCTGAAG
>mmu-miR-505-5p
GGGAGCCAGGAAGTATTGATGTT
>mmu-miR-7230-5p
AGACTGCTGTTTCTCTGAGTGGC
>mmu-miR-27b-5p
AGAGCTTAGCTGATTGGTGAAC
>mmu-miR-331-5p
CTAGGTATGGTCCCAGGGATCC
>mmu-miR-878-5p
TATCTAGTTGGATGTCAAGACA
>mmu-miR-874-5p
CGGCCCCACGCACCAGGGTAAG
>mmu-miR-128-3p
TCACAGTGAACCGGTCTCTTT
>mmu-miR-873a-3p
GAGACTGACAAGTTCCCGGGA
>mmu-miR-3066-3p
CACTTAATAGACCGCAACCTGC
>mmu-miR-6906-3p
GAATTCCGGTCTCCTCTCCAG
>mmu-miR-188-3p
CTCCCACATGCAGGGTTTGCA
>mmu-miR-6537-3p
TTATCAGGTGGAAGAACTCAGCT
>mmu-miR-344f-3p
CTCTAGCCAGGACCTGACTAC
>mmu-miR-5114
ACTGGAGACGGAAGCTGCAAGA
>mmu-miR-3069-5p
TTGGCAGTCAAGATATTGTTTAGC
>mmu-miR-743a-3p
GAAAGACACCAAGCTGAGTAGA
>mmu-miR-6979-3p

TTGTGTCTGTCTGGCTCCCAG
>mmu-miR-8112
TCTCCGCCACCTCCACCGCA
>mmu-miR-1199-5p
TCTGAGTCCCGGTCGCGCGG
>mmu-miR-6968-5p
AGGGGGCGAGGGGGCCCTGTGG
>mmu-miR-5616-3p
AACTTGTGATGAGGTGAGACAG
>mmu-miR-216b-3p
ACACTTACCTGTAGAGATTCTT
>mmu-miR-12184-5p
CTTGCATGTGGGCCTGTGTGCT
>mmu-miR-7055-5p
TCAGTGGGTTATCTGAGTTGG
>mmu-miR-128-2-5p
GGGGGCCGATGCACTGTAAGAGA
>mmu-miR-6919-3p
GCTCCTCCTCCTGGCCCACAG
>mmu-miR-582-5p
ATACAGTTGTTCAACCAGTTAC
>mmu-miR-669d-5p
ACTTGTGTGTGCATGTATATGT
>mmu-miR-7231-3p
CTTGCTTCTTTGTTTCCCCAGAA
>mmu-miR-3473b
GGGCTGGAGAGATGGCTCAG
>mmu-miR-12180-3p
AGGAGCTGTGGACACTTG
>mmu-miR-337-5p
CGGCGTCATGCAGGAGTTGATT
>mmu-miR-7228-3p
AGTGTTCGGTCTCCAGT
>mmu-miR-7004-3p
CCCACTCCCTCCTTCACGCAG
>mmu-miR-1190
TCAGCTGAGGTTCCCCTCTGTC
>mmu-miR-692
ATCTCTTTGAGCGCCTCACTC
>mmu-miR-450b-3p
ATTGGGAACATTTTGCATGCAT
>mmu-miR-6914-3p
TCTGTCCACCTCTTCCCAG
>mmu-miR-544-3p
ATTCTGCATTTTTAGCAAGCTC
>mmu-miR-6408
CACATGGGCTTGCATGGGATGGTC
>mmu-miR-872-3p
TGAACTATTGCAGTAGCCTCCT
>mmu-miR-130b-5p
ACTCTTTCCCTGTTGCACTACT
>mmu-miR-6993-3p
CTTGCGTTTGTTTCCTCACAG
>mmu-miR-12190-3p
TGGAGAACTCTGGGGGAG
>mmu-miR-683
CCTGCTGTAAGCTGTGTCCTC
>mmu-miR-221-5p
ACCTGGCATACAATGTAGATTTCTGT
>mmu-miR-3569-3p
TCAGTCTGCGCTCCTCTCCAGC
>mmu-miR-7664-3p

TCTCTTAGGCTGGGTTAACTAAT
>mmu-miR-12179-5p
TCTCTGTCCTCCAGTTCTG
>mmu-miR-7117-3p
TGTCCTTCTCTTTCCCCCAG
>mmu-miR-365-3p
TAATGCCCCTAAAAATCCTTAT
>mmu-miR-6481
CACTGAAAATGCTTAGATG
>mmu-miR-6345
TAGCACTGCTCAGGGTGAACT
>mmu-miR-344g-5p
AGTCAGGCTCCTGGCAGGAGT
>mmu-miR-1912-3p
CACAGAACATGCAGTGAGAACT
>mmu-miR-574-5p
TGAGTGTGTGTGTGTGAGTGTGT
>mmu-miR-7117-5p
TCTGGGGGCTCAGCTGAGGATA
>mmu-miR-1968-3p
ACCACCTCTGCAGCTGTTAAG
>mmu-miR-7669-3p
GAGCTGGGTGTGGTGGCACATGC
>mmu-miR-5618-3p
CTATACCATGTAAAGGGT
>mmu-miR-6955-3p
ACACCTGTCTCCTTTGCCCACA
>mmu-miR-344e-3p
GATATAACCAAAGCCTGACTAT
>mmu-miR-204-3p
GCTGGGAAGGCAAAGGGACGT
>mmu-miR-7074-5p
TGGGAAGAGGCTAGGGCTCCAGT
>mmu-miR-126b-3p
CGCGTACCAAAAGTAATAATGTG
>mmu-miR-1966-3p
TTTCTGACTCAACTCTCCCTTAG
>mmu-miR-8094
AACTGAAGGACAACGAGAAGA
>mmu-miR-361-5p
TTATCAGAATCTCCAGGGGTAC
>mmu-miR-449b
AGGCAGTGTTGTTAGCTGGC
>mmu-miR-376c-3p
AACATAGAGGAAATTTCACGT
>mmu-miR-6372
AGAACTGGGGAAACTTCCAGCCT
>mmu-miR-7029-3p
TGTGGCTCTGTGTGTCCAGATC
>mmu-miR-6351
TAGCGTTGCTCAGGGAGCACATG
>mmu-miR-654-3p
TATGTCTGCTGACCATCACCTT
>mmu-miR-3067-5p
AGTTCTCAGGCCCGCTGTGGTGT
>mmu-miR-3095-5p
AAGCTTTCTCATCTGTGACACT
>mmu-miR-34a-3p
AATCAGCAAGTATACTGCCCT
>mmu-miR-299a-5p
TGGTTTACCGTCCCACATACAT
>mmu-miR-5104b-5p

TGGGCCATCTCTTTAGTCCAGA
>mmu-miR-1197-3p
TAGGACACATGGTCTACTTCT
>mmu-miR-34c-5p
AGGCAGTGTAGTTAGCTGATTGC
>mmu-miR-6367
TCCCTGAGACCCTGGTTCAGGA
>mmu-miR-1193-5p
TGGTAGACCGGTGACGTACA
>mmu-miR-7038-3p
CACTGCTCCTGCCTTCTTACAG
>mmu-miR-3095-3p
TGGACACTGGAGAGAGAGCTTTT
>mmu-miR-1897-3p
TCAACTCGTTCTGTCCGGTGAG
>mmu-miR-7043-3p
ACTGTGCCTCTCTGTTTTCAG
>mmu-miR-362-5p
AATCCTTGGAACCTAGGTGTGAAT
>mmu-miR-7666-5p
GTAGCCCGGGCGGTGCTTCCCC
>mmu-miR-6418-3p
ACTGCAACCTCCTTTCTCCAGG
>mmu-miR-142a-5p
CATAAAGTAGAAAGCACTACT
>mmu-miR-12182-5p
ACAGCGCCAGCTGCCTAATTGA
>mmu-miR-9768-5p
TGGCACTCGGAGGACGGCACTGT
>mmu-miR-7678-5p
TTACAAGGGCTGGATAGGAAT
>mmu-miR-328-5p
GGGGGGCAGGAGGGGCTCAGGG
>mmu-miR-7679-5p
AGGCTGGCACCAGGATCCCT
>mmu-miR-6380
TGTAAGTGCTTTTAACTGCTGAGC
>mmu-miR-467d-5p
TAAGTGCGCGCATGTATATGCG
>mmu-miR-1943-3p
CAGGTGCCAGCTCCTCCCTTC
>mmu-miR-6390
GAATTGATCAGGATGTAACCA
>mmu-miR-675-5p
TGGTGCGGAAAGGGCCCACAGT
>mmu-miR-466h-5p
TGTGTGCATGTGCTTGTGTGTA
>mmu-miR-7214-5p
TGTTTTCTGGGTTGGAATGAGAA
>mmu-miR-6911-5p
TGGGCTGAGTGGGTCTGTGGCAT
>mmu-miR-6395
CTGGCCCTCTCTGCCCTGTTTA
>mmu-miR-151-5p
TCGAGGAGCTCACAGTCTAGT
>mmu-miR-28a-5p
AAGGAGCTCACAGTCTATTGAG
>mmu-miR-7006-3p
TTTCTGACCTGGATCCCCAG
>mmu-miR-6897-3p
CGGTCTGTGTTCTTCCCCCAG
>mmu-let-7c-1-3p

CTGTACAACCTTCTAGCTTTCC
>mmu-miR-144-3p
TACAGTATAGATGATGTACT
>mmu-miR-696
GCGTGTGCTTGCTGTGGG
>mmu-miR-7578
CATGGCTCTGTCTTCTGCCTCAGA
>mmu-miR-322-5p
CAGCAGCAATTCATGTTTTGGA
>mmu-miR-6901-3p
GACCTTCTGTGTTCTTGCAG
>mmu-miR-7653-5p
TAAGGGGCGGACAGACAGACGG
>mmu-miR-879-3p
GCTTATGGCTTCAAGCTTTCGG
>mmu-miR-12189-3p
ATGGCCTGTACTCTCACACT
>mmu-miR-302d-5p
ACTTTAACATGGAGGCACTTGCT
>mmu-miR-669b-3p
CATATACATACACACAAACATAT
>mmu-miR-742-3p
GAAAGCCACCATGCTGGGTAAA
>mmu-miR-3068-3p
GGTGAATTGCAGTACTCCAACA
>mmu-miR-6715-3p
CCAAACCAGGCGTGCCTGTGG
>mmu-miR-5626-3p
CAGCAGTTGAGTGATGTGACAC
>mmu-miR-466l-5p
TTGTGTGTACATGTACATGTAT
>mmu-miR-1902
AGAGGTGCAGTAGGCATGACTT
>mmu-miR-203-5p
AGTGGTTCTTGACAGTTCAACA
>mmu-miR-7223-3p
GTCTCGAACAAGCTAGG
>mmu-miR-6915-3p
CAACTTCCTTCCTCGTGCCCAG
>mmu-miR-219a-1-3p
AGAGTTGCGTCTGGACGTCCCG
>mmu-miR-7673-5p
TTTGACTGAGAGGTAAGGAATG
>mmu-miR-106a-5p
CAAAGTGCTAACAGTGCAGGTAG
>mmu-miR-3544-3p
ACTCCTGCATGACGCCGTTCCC
>mmu-miR-329-5p
AGAGGTTTTCTGGGTCTCTGTT
>mmu-miR-6342
CAGCAGCAATCTGGTCTTTGAG
>mmu-miR-293-5p
ACTCAAACTGTGTGACATTTTG
>mmu-miR-3471
TGAGATCCAACTGTAAGGCATT
>mmu-miR-6355
CACAATGCTCCAATGTGATAT
>mmu-miR-5132-3p
CTGATGTTCCCCATCCCGCAG
>mmu-miR-7670-5p
ATTCAGATGGGCAGATTGGGAA
>mmu-miR-7221-3p

TGACTGTGGGCTGGGGACTGG
>mmu-miR-7654-5p
GGAGTCGCGCTCCCGCTCGCGC
>mmu-miR-7215-3p
TGCTCTGAGAGGCAGAAAAGCGG
>mmu-miR-19b-2-5p
AGTTTTGCAGATTTGCAGTTCAGC
>mmu-miR-590-3p
TAATTTTATGTATAAGCTAGT
>mmu-miR-363-5p
CAGGTGGAACACGATGCAATTT
>mmu-miR-3113-5p
GTCCTGGCCCTGGTCCGGGTCC
>mmu-miR-128-1-5p
CGGGGCCGTAGCACTGTCTGA
>mmu-miR-677-5p
TTCAGTGATGATTAGCTTCTGA
>mmu-miR-3089-5p
TGAGTTCAGGGACAGCGTGTCT
>mmu-miR-7059-3p
TGTTTGCTGTTTGTCCTCCCAG
>mmu-miR-6937-3p
TCACAGGTTCCCCCTTGCACTAG
>mmu-miR-12195-3p
CAGACAAGACTGTTATACCC
>mmu-miR-3061-5p
CAGTGGGCCGTGAAAGGTAGCC
>mmu-miR-293-3p
AGTGCCGCAGAGTTTGTAGTGT
>mmu-miR-129b-3p
CAAGCCCAGACCGCAAAAAGATT
>mmu-miR-20b-5p
CAAAGTGCTCATAGTGCAGGTAG
>mmu-miR-376a-5p
GGTAGATTCTCCTTCTATGAGT
>mmu-miR-6928-3p
AAGCTGTTCCTGTACTCTCCAG
>mmu-miR-125b-5p
TCCCTGAGACCCTAACTTGTGA
>mmu-miR-7051-5p
TCACCAGGAGGAAGTTGGGTCA
>mmu-miR-7235-3p
TCTGACTTCTTGCTTCTCTCCTC
>mmu-miR-5622-5p
TTCACCACACCCAGCTTAAAGA
>mmu-miR-7233-3p
TATTGTCTGCCTTTAGGTCTAC
>mmu-miR-7118-3p
TAACTCTACGTCCTCCTCCACA
>mmu-miR-6933-5p
TGGCGGCAGTCAGGATACCTGT
>mmu-miR-12185-3p
TCAGCCACTGTGTCCCCTCCTT
>mmu-miR-181d-5p
AACATTCATTGTTGTCGGTGGGT
>mmu-miR-5110
GGAGGAGGTAGAGGGTGGTGGAATT
>mmu-miR-7227-3p
TCTGAAAAGGGTCATCGGCAGA
>mmu-miR-6404
GTGGGAAGTGACAAGGGGTAG
>mmu-miR-3475-3p

TCTGGAGGCACATGGTTTGAA
>mmu-miR-466o-3p
TACATACATGCACACATAAGAC
>mmu-miR-208b-3p
ATAAGACGAACAAAAGGTTTGT
>mmu-miR-3112-5p
ACATAGAAAAGGCAGTCTGCA
>mmu-miR-142a-3p
TGTAGTGTTTCCTACTTTATGGA
>mmu-miR-466f-5p
TACGTGTGTGTGCATGTGCATG
>mmu-miR-7089-3p
TCAGTGTCTGCTGTCTCCACAG
>mmu-miR-7075-5p
CGGGGAGGAGGACATGGTTTT
>mmu-miR-8090
GAAGCGCAGTGGAGGTCTGT
>mmu-miR-7005-3p
CTGTGCCTGCTACCCATCCCTTGCA
>mmu-miR-7672-3p
TCGGCTGCCAGGTTCATGTGG
>mmu-miR-7082-5p
TACGGGCAGGAGGAGGGGAGG
>mmu-miR-6398
GTCAATGAACCTCAGGTAAGG
>mmu-miR-6945-3p
TCTGAGCTCTGCCCTTCCCAT
>mmu-miR-335-3p
TTTTTCATTATTGCTCCTGACC
>mmu-miR-652-5p
CAACCCTAGGAGGGGGTGCCATTC
>mmu-miR-8116
AAGGATCCGGGGCTAGTTGGC
>mmu-miR-467a-3p
CATATACATACACACACCTACA
>mmu-miR-451a
AAACCGTTACCATTACTGAGTT
>mmu-miR-146a-3p
CCTGTGAAATTCAGTTCTTCAG
>mmu-miR-666-5p
AGCGGGCACAGCTGTGAGAGCC
>mmu-miR-1901
CCGCTCGTACTCCCGGGGGTCC
>mmu-miR-6991-5p
TGGAGAACAGGGACTGGGCCTG
>mmu-miR-122b-5p
TTTAGTGTGATAATGGCGTTTG
>mmu-miR-6769b-3p
CATCTTCCCTGTCCCACCCAG
>mmu-miR-1224-5p
GTGAGGACTGGGGAGGTGGAG
>mmu-miR-503-3p
GAGTATTGTTTCCACTGCCTGG
>mmu-miR-705
GGTGGGAGGTGGGGTGGGCA
>mmu-miR-12196-3p
TCACAGGTGAGCTAGAAAGTAT
>mmu-miR-378b
CTGGACTTGGAGTCAGAAGA
>mmu-miR-673-5p
CTCACAGCTCTGGTCCTTGGAG
>mmu-miR-488-5p

CCCAGATAATAGCACTCTCAA
>mmu-miR-181d-3p
CCCACCGGGGGATGAATGTCA
>mmu-miR-3081-5p
GACTGGAGCTTGGAGCGGTGAGC
>mmu-miR-344d-3-5p
AGTCAGGCTAGTGGTTATACTCC
>mmu-miR-7227-5p
TCTGATGACTTTTTCATTGA
>mmu-miR-539-3p
CATACAAGGATAATTTCTTTTT
>mmu-miR-29b-1-5p
GCTGGTTTCATATGGTGGTTTA
>mmu-miR-7013-5p
TATGAAGAGGCCAGTGTTGTAG
>mmu-miR-26a-2-3p
CCTGTTCTTGATTACTTGTTTC
>mmu-miR-6374
AAGTTATAAGCTAGTTGGTGCT
>mmu-miR-5132-5p
GCGTGGGGTGGTGGACTCAGG
>mmu-miR-759
GCAGAGTGCAAACAATTTTGAC
>mmu-miR-133a-3p
TTTGGTCCCCTTCAACCAGCTG
>mmu-miR-465d-5p
TATTTAGAATGGTACTGATGTGA
>mmu-miR-5116
TTTGATAGGAACCCCGCCTGA
>mmu-miR-7653-3p
CCCCTACTGTTCCACCCCGCCT
>mmu-miR-12195-5p
TGGGTATAACAGTCTTGGCTGG
>mmu-miR-8092
GCCATCTGGGACTCGCAGGCA
>mmu-miR-6349
TGGAATGTGGGAGGGGCATGCG
>mmu-miR-343
TCTCCCTTCATGTGCCCAGA
>mmu-miR-3086-3p
CCCAATGAGCCTACAGTCTAAG
>mmu-miR-7094b-2-5p
TCTGGAGAGGATACAGGGTCTGA
>mmu-miR-487b-5p
TGGTTATCCCTGTCCTCTTCG
>mmu-miR-615-5p
GGGGGTCCCCGGTGCTCGGATC
>mmu-miR-6936-3p
CGGCGCGCGCGGTCGCTCACC
>mmu-miR-6419
CAGCAGCAATCTGACACATGGA
>mmu-miR-344h-3p
GGTATAACCAAAGCCCGACTGT
>mmu-miR-6986-3p
GTTTTCACCTTCCTCCCAG
>mmu-miR-877-5p
GTAGAGGAGATGGCGCAGGG
>mmu-miR-7003-3p
CCCCGGGTTTTCCCCACAG
>mmu-miR-7061-5p
TATGGAAATAGGAGGTGATTTT
>mmu-miR-7240-5p

TTGGAGAGGACCGCCGTCGGA
>mmu-miR-379-5p
TGGTAGACTATGGAACGTAGG
>mmu-miR-370-5p
CAGGTCACGTCTCTGCAGTT
>mmu-miR-679-5p
GGACTGTGAGGTGACTCTTGGT
>mmu-miR-490-3p
CAACCTGGAGGACTCCATGCTG
>mmu-miR-7652-3p
TCTGGCTTTCCTGTGGCTTCT
>mmu-miR-202-3p
AGAGGTATAGCGCATGGGAAGA
>mmu-miR-21b
TAGTTTATCAGACTGATATTTCC
>mmu-miR-7222-3p
TCCAGGACAGTGGGCAGGAGCAG
>mmu-miR-7052-3p
GCTCTGCCCCCTCCTTCCCAG
>mmu-miR-6950-5p
TCCTGGAGTGGAGGACAGTCAGA
>mmu-miR-6929-3p
CAGGTGCTGTCTTCTTCTTCCAG
>mmu-miR-224-3p
AAATGGTGCCCTAGTGACTACA
>mmu-miR-672-3p
ACACACAGTCACTATCTTCGA
>mmu-miR-717
CTCAGACAGAGATACCTTCTCT
>mmu-miR-470-3p
AACCAGTACCTTTCTGAGAAGA
>mmu-miR-466f-3p
CATACACACACACATACACAC
>mmu-miR-1197-5p
CGGTTGACCATGGTGTGTACG
>mmu-miR-7085-3p
TAGCTGGCCTCTCCCCACCTTC
>mmu-miR-6935-3p
TGTTCTTTGTCTTTCCACAG
>mmu-miR-6922-5p
TGTGGGAGGGGACTGTAGAGAGG
>mmu-miR-5627-5p
AGAGGGTGCGCCGGGCCCTGCG
>mmu-miR-149-3p
GAGGGAGGGACGGGGGCGGTGC
>mmu-miR-205-5p
TCCTTCATTCCACCGGAGTCTG
>mmu-miR-98-3p
CTATACAACTTACTACTTTCCT
>mmu-miR-292b-5p
ACTCAAAACCTGGCGGCACTTTT
>mmu-miR-5106
AGGTCTGTAGCTCAGTTGGCAGA
>mmu-miR-26a-5p
TTCAAGTAATCCAGGATAGGCT
>mmu-miR-599
TTGTGTCAGTTTATCAAAC
>mmu-miR-6921-3p
TGACTACTCCTTGCCTCTCAG
>mmu-miR-3569-5p
TCGGAGGAGAGCAGACCGGTG
>mmu-miR-467c-5p

TAAGTGCGTGCATGTATATGTG
>mmu-miR-7668-3p
CGGTGCAGGGGATTCAGAGGAG
>mmu-miR-374b-5p
ATATAATACAACCTGCTAAGTG
>mmu-miR-551b-3p
GCGACCCATACTTGGTTTCAG
>mmu-miR-7243-3p
TCAGAACAGCTGACAGCAGT
>mmu-miR-7019-5p
ATCTGGGAGGGGCTGTCAGGAGT
>mmu-miR-5624-5p
TATTGTAAACTCTGCCTT
>mmu-miR-7237-5p
ATGGTAGCTTGTGACAGGACTGGC
>mmu-miR-5627-3p
ACAGGGCTCTCCGGCGCCCCTCGT
>mmu-miR-18a-5p
TAAGGTGCATCTAGTGCAGATAG
>mmu-miR-467e-5p
ATAAGTGTGAGCATGTATATGT
>mmu-miR-6988-3p
TGACCTCTGTATCTCCTGCCAG
>mmu-miR-6930-3p
CTGGCCTCTCCATACTCCCAG
>mmu-miR-8091
AGGAGTGAGTGGTAAGCTGGTG
>mmu-miR-7242-5p
ATTGGGCTTGGGAGGTGCTTGT
>mmu-miR-1188-5p
TGGTGTGAGGTTGGGCCAGGA
>mmu-miR-467f
ATATACACACACACACCTACA
>mmu-let-7i-5p
TGAGGTAGTAGTTTGTGCTGTT
>mmu-miR-184-5p
CCTTATCACTTTTCCAGCCAGC
>mmu-miR-5625-5p
CCCGGAAGTTCTTGAGTAGGA
>mmu-miR-1947-5p
AGGACGAGCTAGCTGAGTGCTG
>mmu-miR-1912-5p
TGCTCATTGCATGGGCTGTGTA
>mmu-miR-374c-5p
ATAATACAACCTGCTAAGTG
>mmu-miR-145a-3p
ATTCCTGGAAATACTGTTCTTG
>mmu-miR-7232-3p
TGGTTGAATTCGACTTTGGGGC
>mmu-miR-7668-5p
CACTCTGAATAACCTGCTCTCTGT
>mmu-miR-6949-3p
TTCCCCTTTTTCTATCCACAG
>mmu-miR-29c-3p
TAGCACCATTTGAAATCGGTTA
>mmu-miR-290b-3p
AAGTGCCCCCATAGTTTGAGTA
>mmu-miR-6933-3p
AGGTGTTTCCTCTGCCGTCA
>mmu-miR-193b-5p
CGGGGTTTTGAGGGCGAGATGA
>mmu-miR-201-3p

TGAACAGTGCCTTTCTGTGTAGG
>mmu-miR-665-3p
ACCAGGAGGCTGAGGTCCCT
>mmu-miR-6985-3p
CTCACAGCCACCTCCTCACAG
>mmu-miR-6906-5p
ATGGATGGGCAGACGGACAGACAG
>mmu-miR-208b-5p
AAGCTTTTTGCTCGCGTTATGT
>mmu-miR-34b-5p
AGGCAGTGTAATTAGCTGATTGT
>mmu-miR-7674-5p
TGAGGTGTGGGCAGCATGAGGACT
>mmu-miR-7118-5p
TGGGGAAGGCGGGAGAGGGAAC
>mmu-miR-543-5p
AAGTTGCCCGCGTGTTTTTCG
>mmu-miR-3093-3p
TGTGGACACCGTGGGAGGTTGG
>mmu-miR-7234-5p
TTGTTTTCTCCAAAGACGTTTCT
>mmu-miR-140-3p
TACCACAGGGTAGAACCACGG
>mmu-miR-342-5p
AGGGGTGCTATCTGTGATTGAG
>mmu-miR-299a-3p
TATGTGGGACGGTAAACCGCTT
>mmu-miR-6915-5p
AGGTCAGAGCAGATGGAGGTTCGT
>mmu-miR-3971
CTCCCCACCCCTGTACCAGTGA
>mmu-miR-15a-5p
TAGCAGCACATAATGGTTTGTG
>mmu-miR-3113-3p
TCCTGGCCCTGGTCCTGGTC
>mmu-miR-7034-5p
TCCGGGAGGGATGGATGTGCT
>mmu-miR-5623-3p
GTTCTGGCTTTCAGCTGTCACC
>mmu-miR-7078-5p
TGTGGGTGGTAGGAGACGCT
>mmu-miR-7017-5p
AGAGGGTTGTGAGACTAGGGCTGT
>mmu-miR-7001-3p
CGCTCACACTCCCTCTGCAG
>mmu-miR-6932-3p
TCTGTTCTGCTTCCTCTGCAG
>mmu-miR-467e-3p
ATATACATACACACACCTATAT
>mmu-miR-467g
TATACATACACACACATATAT
>mmu-miR-3064-5p
TCTGGCTGTTGTGGTGTGCAAA
>mmu-miR-6918-3p
TGAGCCTGTGCCCTTCTCACAG
>mmu-miR-186-5p
CAAAGAATTCTCCTTTTGGGCT
>mmu-miR-3085-3p
TCTGGCTGCTATGGCCCCCTC
>mmu-miR-6369
TGGCTCAGTTCAGTGTTGGTGAG
>mmu-miR-302b-3p

TAAGTGCTTCCATGTTTTAGTAG
>mmu-miR-1899
AGCGATGGCCGAATCTGCTTCC
>mmu-miR-3473h-5p
TAGGGGCTGGAAAGGTGACT
>mmu-miR-487b-3p
AATCGTACAGGGTCATCCACTT
>mmu-miR-135b-3p
ATGTAGGGCTAAAAGCCATGGG
>mmu-let-7a-5p
TGAGGTAGTAGGTTGTATAGTT
>mmu-miR-6239
TAGCGTTGGATCACTCGGTG
>mmu-miR-3620-3p
CTGCTCAGCCCCGCCCACCCCA
>mmu-miR-7035-3p
TCTGAGCCGCTGTCCCTGCAG
>mmu-miR-7670-3p
TTCCCTTTCTGAATATCTGTAGA
>mmu-miR-7655-5p
CGGCCCACGGAGCTCCTAAAGA
>mmu-miR-455-5p
TATGTGCCTTTGGACTACATCG
>mmu-miR-3086-5p
TAGATTGTAGGCCCATTGGA
>mmu-miR-12198-3p
GACTCAGGAGGACTGCCGG
>mmu-miR-5710
TCTTGGGACATAGTGTAAGGCA
>mmu-miR-879-5p
AGAGGCTTATAGCTCTAAGCC
>mmu-miR-6347
CAGTGCGGCTGGCTGGAGATGC
>mmu-miR-7672-5p
CGGTGGACTTGACAGCGGGCGA
>mmu-miR-351-3p
GGTCAAGAGGCGCCTGGGAAC
>mmu-let-7b-5p
TGAGGTAGTAGGTTGTGTGGTT
>mmu-miR-3968
CGAATCCCACTCCAGACACCA
>mmu-miR-107-3p
AGCAGCATTGTACAGGGCTATCA
>mmu-miR-1982-5p
TTGGGAGGGTCCTGGGGAGG
>mmu-miR-7682-5p
AGGCACGACTCTTGCCACAGAAAC
>mmu-miR-6926-5p
TCAGTGGGGTGAGGGATGGTGA
>mmu-miR-7092-5p
AAAGGCAAACTCACAGAATCATT
>mmu-miR-127-3p
TCGGATCCGTCTGAGCTTGGCT
>mmu-miR-6911-3p
TCAATCACACTCTGTCCACAG
>mmu-miR-6902-5p
TTCCTCACATTGTCACATGAAG
>mmu-miR-547-5p
CACTTGAGGATGTACCACCCA
>mmu-miR-7048-5p
CGGGGCTGAGAGGTGAGGAAGC
>mmu-miR-6947-3p

GCAGCCTCTTTCCCTGGCATCA
>mmu-miR-5621-3p
TGGGCCCTCCAGACCTCATGC
>mmu-miR-132-3p
TAACAGTCTACAGCCATGGTCG
>mmu-miR-344f-5p
AGTCAGTCTCCTGGCTGGAGTC
>mmu-miR-7046-3p
TGAACCACCCATCCCCCTACAG
>mmu-miR-883a-3p
TAACTGCAACAGCTCTCAGTAT
>mmu-miR-875-3p
CCTGAAAATACTGAGGCTATG
>mmu-miR-5626-5p
GCCCCATCGATTAACTGCTTCC
>mmu-miR-7053-5p
TGGGGAAAGGCAGGCTACTGG
>mmu-miR-450b-5p
TTTTGCAGTATGTTCCTGAATA
>mmu-miR-7025-3p
CAGCACAACTCCAGCTCACCAG
>mmu-miR-219b-3p
AGAATTGCGTTTGGACAATCAGT
>mmu-miR-6348
TCAGCCTTTATAAGGTGTGTGT
>mmu-miR-3073b-5p
ATGGTCACAGTGGACATCAACC
>mmu-miR-323-3p
CACATTACACGGTCGACCTCT
>mmu-miR-7078-3p
TACTTTTTTTATCATCCACAG
>mmu-miR-6974-5p
GTGAGGCAGCAAGAGATTGGGGGT
>mmu-miR-6359
CGATGTTGCCCAGGGTCAGAA
>mmu-miR-295-5p
ACTCAAATGTGGGGCACACTTC
>mmu-miR-20a-3p
ACTGCATTACGAGCACTTAAAG
>mmu-miR-7241-5p
ACTTGCATGAGTTAATACCCACC
>mmu-miR-92a-2-5p
AGGTGGGGATTGGTGGCATTAC
>mmu-miR-7236-3p
TTGGACCGCACAGATTTAGACC
>mmu-let-7d-3p
CTATACGACCTGCTGCCTTTCT
>mmu-miR-7116-3p
TTTTTTTCCTTTGCCTTCTCAG
>mmu-miR-7020-3p
AACCCCTCTCTTCTCTCCCAG
>mmu-miR-130b-3p
CAGTGCAATGATGAAAGGGCAT
>mmu-miR-7651-3p
TCAGTTTGGTTTTTTTCCCCTTGA
>mmu-miR-130c
CAGTGCAATGTTCCAAGGTGTG
>mmu-miR-181c-3p
ACCATCGACCGTTGAGTGGACC
>mmu-miR-377-5p
AGAGGTTGCCCTTGGTGAATTC
>mmu-miR-338-3p

TCCAGCATCAGTGATTTTGTTG
>mmu-miR-6908-5p
TTGCCTGTCAGGGAGAGCCCCA
>mmu-miR-490-5p
CCATGGATCTCCAGGTGGGT
>mmu-miR-410-3p
AATATAACACAGATGGCCTGT
>mmu-miR-378a-3p
ACTGGACTTGGAGTCAGAAGG
>mmu-miR-17-5p
CAAAGTGCTTACAGTGCAGGTAG
>mmu-miR-345-3p
CCTGAACTAGGGGTCTGGAGAC
>mmu-miR-6388
TGCAGATGGCACAGTGAGGCC
>mmu-miR-146a-5p
TGAGAACTGAATTCCATGGGTT
>mmu-miR-7689-5p
CCGAGGCAGCCTGGCTTAG
>mmu-miR-7055-3p
TTGCTACTTTGATACCCCACCAG
>mmu-miR-7028-5p
TGGGCTGAGGCTTGGGTCAGGAG
>mmu-miR-5119
CATCTCATCCTGGGGCTGG
>mmu-miR-12186-5p
AGGGACTCAGGCCTCTGTTGG
>mmu-miR-698-3p
CATTCTCGTTTCCTTCCCT
>mmu-miR-669e-5p
TGTCTTGTGTGTGCATGTTCAT
>mmu-miR-8108
TCTGGGGAGGAGCGTAGTTACA
>mmu-miR-7037-3p
TGTCTCCTAATGTCACCTGCAG
>mmu-miR-1946a
AGCCGGGCAGTGGTGGCACACACTTTT
>mmu-miR-6376
CAGTGTGGCCAATTGGAGATTA
>mmu-miR-1929-3p
CAGCTCATGGAGACCTAGGTGG
>mmu-miR-6973a-5p
TACGGTGGGAGGGGTGGAGTTG
>mmu-miR-673-3p
TCCGGGGCTGAGTTCTGTGCACC
>mmu-miR-6923-5p
GTGAGGGCAGGAGGATTGGGGTGT
>mmu-miR-6916-3p
CTTGGTCTCCCTCTTCCTACAG
>mmu-miR-19b-3p
TGTGCAAATCCATGCAAAACTGA
>mmu-miR-7240-3p
CGCCGTTGGTCCTCTCCAACA
>mmu-miR-455-3p
GCAGTCCACGGGCATATACAC
>mmu-miR-151-3p
CTAGACTGAGGCTCCTTGAGG
>mmu-miR-499-3p
GAACATCACAGCAAGTCTGTGCT
>mmu-miR-3092-3p
GAATGGGGCTGTTTCCCCTCC
>mmu-miR-590-5p

GAGCTTATTCATAAAAGTGCAG
>mmu-miR-3552
AGGCTGCAGGCCCACTTCCCT
>mmu-miR-871-5p
TATTCAGATTAGTGCCAGTCATG
>mmu-miR-466i-5p
TGTGTGTGTGTGTGTGTGTG
>mmu-miR-5617-5p
GTAAGTGAGGGCAAGCCTTCTGG
>mmu-miR-871-3p
TGACTGGCACCATTCTGGATAAT
>mmu-miR-7056-3p
CCTCACCTGTCTCCCACCACAG
>mmu-miR-760-5p
CCCCTCAGGCCACCAGAGCCCGG
>mmu-miR-678
GTCTCGGTGCAAGGACTGGAGG
>mmu-miR-7007-5p
TCAGAAGAGGCAGTGGAGGAGAT
>mmu-miR-7221-5p
CGTCTCTTGGTCACAGGTGGG
>mmu-miR-7071-3p
CTGACGTCTCGGTTCCCGCAG
>mmu-miR-7049-5p
TCTGAAGGCAGTCAGGTAGGCGG
>mmu-miR-3964
ATAAGGTAGAAAGCACTAAA
>mmu-miR-466n-5p
GTGTGTGCGTACATGTACATGT
>mmu-miR-7225-3p
CTTCTAGACAGCCTCTGCAGCA
>mmu-miR-7224-5p
GGGTAGGCCCCTCAGTGAAGA
>mmu-miR-12186-3p
AGCAGCTGGCCTGTGTCCCTG
>mmu-miR-3547-3p
TGAGCACCACCCCTCTCTCAG
>mmu-miR-547-3p
CTTGGTACATCTTTGAGTGAG
>mmu-miR-375-5p
GCGACGAGCCCCTCGCACAAAC
>mmu-miR-294-5p
ACTCAAAATGGAGGCCCTATCT
>mmu-miR-764-5p
GGTGCTCACATGTCCTCCT
>mmu-miR-3101-3p
TAGCTTTGGTGGATGGTCTTT
>mmu-miR-6973a-3p
CACTCTAACCCTACCTACCCAT
>mmu-miR-351-5p
TCCCTGAGGAGCCCTTTGAGCCTG
>mmu-miR-12193-5p
TGGGTGACTGGAAAGGACAAGC
>mmu-miR-7119-3p
AAAAAACACCGTTTCCTCCAG
>mmu-miR-3078-5p
CAAAGCCTAGACTGCAGCTACCT
>mmu-miR-346-3p
AGGCAGGGGCTGGGCCTGCAGC
>mmu-miR-3063-3p
TGAGGAATCCTGATCTCTCGCC
>mmu-miR-3083b-3p

TCTCTGAAATATTCCCAGCCTT
>mmu-miR-7045-3p
TCTCCCCCCTCCCCCGCACAG
>mmu-miR-10a-5p
TACCCTGTAGATCCGAATTTGTG
>mmu-miR-452-3p
TCAGTCTCATCTGCAAAGAGGT
>mmu-miR-143-5p
GGTGCAGTGCTGCATCTCTGG
>mmu-miR-212-5p
ACCTTGGCTCTAGACTGCTTACT
>mmu-miR-872-5p
AAGGTTACTTGTTAGTTCAGG
>mmu-miR-7040-5p
CATACGGAGGGAGATGGAGGCG
>mmu-miR-6941-5p
CTGGGGGACTGACGGGTAGCT
>mmu-miR-124-3p
TAAGGCACGCGGTGAATGCC
>mmu-miR-3094-5p
TGTTGGGGACATTTTTAAAGC
>mmu-miR-7661-3p
TTGACTCCCAGTTTCTCTCTGC
>mmu-miR-450a-5p
TTTTGCGATGTGTTCCTAATAT
>mmu-miR-704
AGACATGTGCTCTGCTCCTAG
>mmu-miR-1931
ATGCAAGGGCTGGTGCGATGGC
>mmu-miR-6971-3p
ACAGCCTCTGCTTCTTCTCAG
>mmu-miR-3071-5p
ACTCATTTGAGACGATGATGGA
>mmu-miR-7037-5p
AAGGTGGCCACAGGAGATCATGGT
>mmu-miR-7647-3p
AAGTCCAGGAGGTGGTAGTAC
>mmu-miR-7216-5p
TGGAGAGCTGGCAGAGGACCCAGA
>mmu-miR-30d-5p
TGTAAACATCCCCGACTGGAAG
>mmu-miR-3962
AGGTAGTAGTTTGTACATTT
>mmu-miR-6953-3p
CTGACCCAGCCTCTATGTCCCCAG
>mmu-miR-1955-3p
GAGCATTGCATGCTGGGACAT
>mmu-miR-6769b-5p
CCTGGTGGGTGGGGAAGAGC
>mmu-miR-6931-5p
TGGGGTGGGGAGTGGGGGACT
>mmu-miR-12187-5p
TAAGGGGAGAGGTGTGGTGTAG
>mmu-miR-876-5p
TGGATTTCTCTGTGAATCACTA
>mmu-miR-760-3p
CGGCTCTGGGTCTGTGGGGA
>mmu-miR-138-2-3p
GCTATTTCACGACACCAGGGT
>mmu-miR-219a-2-3p
AGAATTGTGGCTGGACATCTGT
>mmu-miR-3089-3p

AGCATCTGCTGATCCTGAGCTGT
>mmu-miR-6354
TGCCCTGGGGATCAGGTCTCT
>mmu-miR-770-5p
AGCACCACGTGTCTGGGCCACG
>mmu-miR-7646-3p
TCACTGGTCGCTCTCTTTCAC
>mmu-miR-7038-5p
TGTAGAAGGAAGGGCTGCTGTG
>mmu-miR-135a-5p
TATGGCTTTTTATTCCTATGTGA
>mmu-miR-200c-5p
CGTCTTACCCAGCAGTGTTTGG
>mmu-miR-194-5p
TGTAACAGCAACTCCATGTGGA
>mmu-miR-7680-5p
ATTCCAGTGAACAAGCAGTT
>mmu-miR-410-5p
AGGTTGTCTGTGATGAGTTCG
>mmu-miR-541-5p
AAGGGATTCTGATGTTGGTCACACT
>mmu-miR-3082-3p
CACATGGCACTCAACTCTGCAG
>mmu-miR-6912-5p
TACAGGGAGGGTGCTCAGGCAG
>mmu-miR-6373
TATGGGAATATCTAGGAGGTGA
>mmu-miR-216c-3p
TTTGCCTGCAGAGATTTCCCAGT
>mmu-miR-6378
TGGTTCACGGGAGGACACACGC
>mmu-miR-1966-5p
AAGGGAGCTGGCTCAGGAGAGAGTC
>mmu-miR-1942
TCAGATGTCTTCATCTGGTTG
>mmu-miR-669m-3p
ATATACATCCACACAAACATAT
>mmu-miR-7022-5p
CCTGGCAAGGAAGGGCTCTGTTT
>mmu-miR-6897-5p
TGGGGGCAGTGTCACACGCTGTT
>mmu-miR-465a-3p|mmu-miR-465b-3p|mmu-miR-465c-3p
GATCAGGGCCTTTCTAAGTAGA
>mmu-miR-7652-5p
TAAGGGCACAGGGATTTCAGGC
>mmu-miR-882
AGGAGAGAGTTAGCGCATTAGT
>mmu-miR-219c-5p
GGACGTCCAGACGCAACTCTCG
>mmu-miR-7081-3p
TTCCCCTGGTATCCTCCTCACA
>mmu-miR-3098-3p
TTCTGCTGCCTGCCTTTAGGA
>mmu-miR-881-3p
AACTGTGTCTTTTCTGAATAGA
>mmu-miR-12193-3p
ACCGGTCCTTTCTGTCCCCAG
>mmu-miR-7210-3p
TTATATGTAAGTAACAATGTGGCT
>mmu-miR-1251-3p
CGCTTTGCTCAGCCAGTGTAG
>mmu-miR-222-3p

```
AGCTACATCTGGCTACTGGGTCT
>mmu-miR-431-5p
TGTCTTGCAGGCCGTCATGCA
>mmu-miR-10b-5p
TACCCTGTAGAACCGAATTTGTG
>mmu-miR-541-3p
TGGCGAACACAGAATCCATACT
>mmu-miR-383-5p
AGATCAGAAGGTGACTGTGGCT
>mmu-miR-297b-3p|mmu-miR-297a-3p|mmu-miR-297c-3p
TATACATACACACATACCCATA
>mmu-miR-6412
TCGAAACCATCCTCAGCTACTA
>mmu-miR-6970-3p
TCACGCCACCCACCCTGTGCT
>mmu-miR-804
TGTGAGTTGTTCCTCACCTGGA
>mmu-miR-1934-5p
TCTGGTCCCCTGCTTCGTCCTCT
>mmu-miR-154-3p
AATCATACACGGTTGACCTATT
>mmu-miR-6922-3p
TGTTCTGCTCACCCTTCTCACAG
>cali_37_rc|Ome_cali|artificial
TTCCGCTTTACGGGTTAATAGA
>cali_23_rc|Ome_cali|artificial
CTCGTCTCCGGCTGTATATACC
>cali_25_rc|alt_cali|artificial
AGGGCCCTTTAGGCACTAATAG
>cali_19_rc|Ome_cali|artificial
ATGATTCTCTAACGTCGGCATT
>cali_01_rc|alt_cali|artificial
TCCACGACGTCTCATGTATTTC
>cali_24_rc|alt_cali|artificial
TGCTACTCCGATCTTTAGCCTC
>cali_17_rc|alt_cali|artificial
TCATGAGTCCGTACCTTGATTG
>cali_18_rc|alt_cali|artificial
ATCATTTACGATTCGGAGCTGT
>cali_05_rc|Ome_cali|artificial
CATGGTTGTAAGTCCCGGTAAT
>cali_44_rc|alt_cali|artificial
AGCCGCATTTCGTAGTGATATT
>cali_38_rc|Ome_cali|artificial
GGAGGATACTTAATCCGCTGTG
>cali_08_rc|Ome_cali|artificial
GGATTACTCGGGTTTGAGACAG
>cali_04_rc|alt_cali|artificial
GGGTACCATACCGGTTGTCTTA
>cali_41_rc|Ome_cali|artificial
TGCTAGTCCACGGGAGAAATAT
>cali_40_rc|Ome_cali|artificial
GGAGCCGTGAATACAATCCTAG
>cali_39_rc|Ome_cali|artificial
GGTAATCCAACGTTGATGGTTT
>cali_43_rc|alt_cali|artificial
TCTAGTTGCGTGATGGAGAGAA
>cali_29_rc|Ome_cali|artificial
CGTCGATTTAGACCGTATAGCC
>cali_27_rc|alt_cali|artificial
GTAGCTGTCAGTACGTTCGTGC
>p3adapter|adapter|artificial
```

```
TCGTATGCCGTCTTCTGCTTG
>p5adapter|adapter|artificial
GTTCAGAGTTCTACAGTCCGACGATC
>cali_20_rc|alt_cali|artificial
GATAGTTCGGGATCGCTGTAAC
```
